# Aging-Related Cellular and Molecular Changes Reveal a Role for *Bglap3+* Neurons in MEC Spatial Cognition

**DOI:** 10.1101/2024.06.27.600930

**Authors:** Qichen Cao, Guge Suri, Sihui Cheng, Yuanjing Liu, Jiangyuan Liu, Shidan Wen, Meng Zhong, Jun Hao, Shuyang Yao, Yikai Yang, Shuai Xu, Hong Jin, Xinyu Yan, Weixiang Guo, Chenglin Miao

## Abstract

Aging-related cognitive decline, including impairments in spatial orientation and memory, has been linked to dysfunction within the hippocampus and medial entorhinal cortex (MEC). However, the cellular and molecular alterations underlying MEC aging remain poorly understood. Here we show that old mice exhibit a reduced proportion of grid cells and decreased spatial stability. Transcriptomic profiling of the MEC using 10x Genomics Visium technology reveals 1664 differentially expressed genes between young and old mice. Among these, *Bglap3* is identified as a marker gene for a subpopulation of Layer III neurons that showed an age-associated reduction in cell number. Chemogenetic silencing of *Bglap3*+ neurons in young mice impairs spatial tuning of MEC neurons. Together, these findings elucidate cellular and molecular mechanisms contributing to age-related MEC dysfunction and suggest that the loss of *Bglap3*+ neurons may underlie spatial coding deficits in aging.

## Introduction

Cognitive decline, particularly deficits in spatial memory, is a major challenge for aging populations ^1,2^. Age-related impairments in spatial learning and memory are observed across species ^3,4^, including nonhuman primates ^5^, dogs ^6^, rats ^7^, and mice ^8^, making animal models essential for understanding age-related cognitive decline in humans. The medial entorhinal cortex (MEC), a key structure in the mammalian spatial navigation network ^9,10^, is one of the first regions to show dysfunction in aging-related cognitive decline ^11,12^. MEC lesions impair spatial learning and memory ^13^, and disrupted grid cell coding hinders path integration in mice^14^. Defining the cellular and molecular changes in the aging MEC is therefore critical for uncovering the mechanisms that drive spatial memory deficits in older adults.

Grid cells were the first functional cell type identified in the MEC ^15^; in an open environment they fire at locations that form a hexagonal grid spanning the entire space. They operate within a broader network of specialized MEC cell types, including head direction (HD) cells ^16^, border cells ^17,18^, and speed cells ^19,20^, which interact with neighboring regions such as lateral entorhinal cortex cells representing objects ^21,22^, hippocampal place cells ^23^ and hippocampal cells representing goals ^24,25^ to form a dynamic representation of space that underpins spatial navigation. Place cells, the primary neuronal type encoding spatial location in the hippocampus ^23^, become increasingly unstable across experimental sessions in aged rats ^7^, and similar spatial coding deficits occur in hippocampal and parahippocampal regions of older humans during virtual navigation ^3^. These findings suggest that hippocampal dysfunction plays a critical role in age-related declines in spatial learning and memory.

Alterations in grid cell coding within the MEC, one synapse upstream of the hippocampus, have been observed during normal aging in humans using fMRI ^26^. Impairments in grid cells and spatial memory have also been documented in transgenic mice expressing mutant human tau ^27^, and grid cell-like activity is impaired in young adults carrying the APOE-ε4 allele, a genetic risk factor for Alzheimer’s disease (AD) ^28^. However, these studies have not clarified whether grid cell impairment arises from a reduction in the number of grid cells, from altered intrinsic firing properties such as periodic tuning and stability, or from both. It also remains unclear whether other populations, such as HD cells, border cells and speed cells, are similarly affected, particularly since aging does not affect neural populations uniformly, with some becoming deficient while others remain intact ^29^.

Aging is accompanied by transcriptomic changes that are not uniform across the heterogeneous cell populations of the brain ^30^. Given the diverse neuronal subtypes of the MEC, each defined by unique molecular markers and distinct projection patterns within the MEC and to the hippocampus ^15,31^, it is essential to determine whether age-related alterations in specific subtypes or subregions contribute to deficits in cognitive functions such as spatial navigation. Bulk and single-cell RNA sequencing have provided extensive insight into age-related transcriptomic changes across brain regions ^32–34^, but do not capture in situ changes within specific regions, and spatial transcriptomics has yet to be applied to the MEC in the context of aging ^35–38^.

Here we compared the activity of different neural populations in the MEC between young (3– 4 months) and old (>28 months) mice using extracellular recordings in freely behaving animals. We also applied the 10x Genomics Visium platform, which minimizes tissue dissociation-induced cell death and efficiently recovers RNA from aged brain tissue, to profile MEC gene expression during aging. Combining chemogenetic silencing with two-photon calcium imaging, we identified a *Bglap3+* neuronal population linked to age-related cognitive decline. These findings provide insight into the changes in spatial coding, molecular profiles and circuit mechanisms that underlie normal aging in the MEC.

## Results

### Recording spatial coding cells in young and old mouse MEC

To determine whether normally aged mice exhibit changes in MEC spatial coding, we performed tetrode recordings in 8 young (3–4 months) and 10 old (>28 months) C57BL/6J mice, with all tetrode traces located in the dorsomedial MEC (Fig. 1a and Supplementary Fig. 1). Neither running speed (young: 23.44 ± 0.42 cm/s vs. old: 26.25 ± 0.41 cm/s; P = 0.1328; Supplementary Fig. 2a) nor coverage of the box (young: 88.57 ± 0.26% vs. old: 89.41 ± 0.35%; P = 0.4289; Supplementary Fig. 2b) differed between age groups. To exclude an effect of impaired vision on navigation, we trained the mice in a visual cue discrimination test; old mice showed no significant decline in performance (young: 70.39% vs. old: 67.20%; two-way ANOVA, age: F (1, 5) = 2.064, P = 0.2103; Supplementary Fig. 2c, d), indicating intact visual capacity. Unless stated otherwise, group comparisons used linear mixed-effects models with mouse identity as a random intercept, and full statistics are reported in the figure legends.

**Fig. 1.**
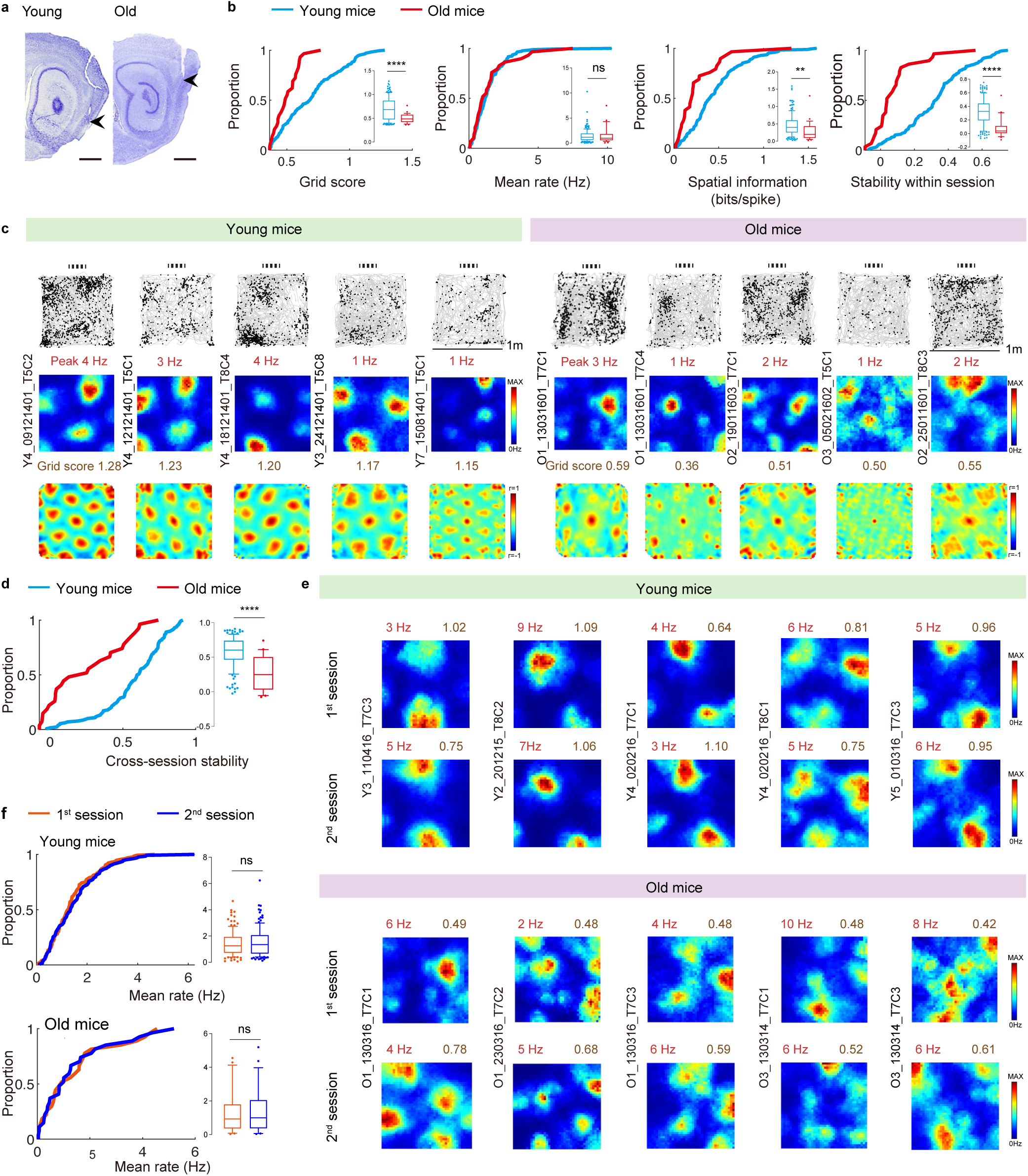
Impaired grid coding and instability across sessions in old mice. **a** Histology showing tetrode traces in MEC (sagittal sections). Left, 3–4-month-old (young) mouse; right, 28-month-old (old) mouse; arrows indicate tetrode locations. Scale bar, 1 mm. **b** Cumulative frequency diagrams and box-whisker plots showing significant decreases in grid score, spatial information, and spatial stability in old vs. young mice (young, n = 169; old, n = 29; linear mixed-effects model: grid score, SE = 0.044, t (196) =-4.34, P = 2.25×10⁻⁵; mean rate, SE = 0.26, t (196) = 0.50, P = 0.62; spatial information, SE = 0.060, t (196) =-3.18, P = 1.74×10⁻³; within-session stability, SE = 0.037, t (196) =-6.23, P = 1.70×10⁻⁹). **c** Trajectory maps (top), color-coded rate maps (middle), and autocorrelograms (bottom) of grid cells in young (left) and old (right) mice. Peak rates (red) and grid scores (brown) shown above each rate map/autocorrelogram; animal and cell numbers indicated left of each triad. **d** Cumulative frequency diagrams and box-whisker plots showing a significant decrease in across-session spatial stability in old vs. young mice (young, n = 128; old, n = 27; linear mixed-effects model, SE = 0.067, t (153) =-4.92, P = 2.23×10⁻⁶). **e** Color-coded rate maps of grid cells in session 1 and session 2; peak rates and grid scores shown above each rate map. **f** Mean firing rate of grid cells across sessions, young (top) and old (bottom) mice (linear mixed-effects model with session as fixed effect; young, n = 128, SE= 0.12, t (254) = 0.93, P = 0.35; old, n = 27, SE = 0.37, t (52) =-0.10, P = 0.92). Data are median (center line), 25th–75th percentiles (box bounds), and 10th–90th percentiles (whiskers). P-values for fixed effects in the linear mixed-effects models were derived from two-sided t-tests. **P < 0.01; ****P < 0.0001; ns, not significant. Source data are provided as a Source Data file.

A total of 1022 cells were recorded, 485 from young and 537 from old mice. Grid cells, HD cells, border cells and speed cells were defined as cells whose grid score, mean vector length (MVL), border score or speed score, respectively, exceeded chance levels determined by repeated shuffling of the experimental data (see Methods). We identified 169 and 29 grid cells, 109 and 84 HD cells, 31 and 40 border cells, and 82 and 37 speed cells in young and old mice, respectively (Supplementary Fig. 3a, c, e, g), with no difference in spike-sorting quality between age groups (Supplementary Fig. 3b, d, f, h).

Analysis of spatial and directional tuning across age groups and within individual mice showed that the proportions of pure grid cells (without directional tuning) and grid+HD conjunctive cells (G+HD) were significantly lower in old mice (young vs. old, two-sided Mann-Whitney test, G: P = 9.14 × 10^-5^, G+HD: P = 3 × 10^-3^), whereas the proportions of pure HD cells (HD), pure border cells (B), border+HD (B+HD), pure speed cells (S) and speed+HD (S+HD) cells were unchanged (Supplementary Fig. 4a–i).

### Spatial tuning of grid cells was impaired in old mice

The grid score of grid cells was significantly lower in old than in young mice (young: 0.69 ± 0.018 vs. old: 0.50 ± 0.018; *P* = 2.25 × 10^-5^; Fig. 1b, c and Supplementary Fig. 5a, b). Mean firing rate was unchanged (young: 1.45 ± 0.094 Hz vs. old: 1.58 ± 0.30 Hz; *P* = 0.6170; Fig. 1b), although firing rate correlated with running speed in young but not old mice (Supplementary Fig. 5c). Spatial information was lower in old mice (young: 0.47 ± 0.024 vs. old: 0.27 ± 0.049; *P* = 1.74 × 10^-3^; Fig. 1b), and the spatial stability of grid fields was markedly reduced (young: 0.32 ± 0.015 vs. old: 0.082 ± 0.025; *P* = 1.70 × 10^-9^; Fig. 1b, c). Spatial stability correlated with grid score in young but not old mice (Supplementary Fig. 5d), indicating that the reduced firing stability of grid cells in old mice cannot be fully explained by their lower grid scores, or vice versa. Across two sessions in the same context, 64.8% of grid cells in young mice (83/128) but only 22.2% in old mice (6/27) retained grid properties (Fisher’s exact test, *P* = 6.98 × 10⁻⁵; Supplementary Fig. 6a, b), and the cross-session correlation of grid fields was significantly lower in old mice (young: 0.57 ± 0.019 vs. old: 0.25 ± 0.05; *P* = 2.23 × 10⁻⁶; Fig. 1d, e). Mean firing rates were stable across sessions in both groups (young, 1^st^: 1.43 ± 0.083 Hz vs. 2^nd^: 1.54 ± 0.092 Hz, *P* = 0.35; old, 1^st^: 1.45 ± 0.26 Hz vs. 2^nd^: 1.41 ± 0.27 Hz, *P* = 0.92; Fig. 1f), as was spike waveform similarity measured by the centre of mass (COM) shift of clusters across sessions (Supplementary Fig. 6c), indicating that grid cell instability in old mice cannot be attributed to changes in firing rate or to errors in unit detection.

### HD and border cells are unimpaired in old mice

Because spatial navigation also requires information about direction and environmental geometry to accurately track the animal’s changing position, we next asked whether the firing patterns of HD cells and border cells recorded simultaneously with grid cells were affected in old mice. HD cells showed no significant differences between young and old mice in MVL (0.34 ± 0.013 vs. 0.33 ± 0.01; *P* = 0.6849), mean firing rate (1.23 ± 0.13 Hz vs. 1.16 ± 0.16 Hz; *P* = 0.7252), angular stability (0.38 ± 0.033 vs. 0.38 ± 0.033; *P* = 0.5928), or the angular difference in peak direction across two sessions (50.54 ± 5.45 vs. 60.48 ± 6.05; *P* = 0.7942) (Fig. 2a–c and Supplementary Fig. 7a, c). The mean firing rate of HD cells was not correlated with running speed in either age group (Supplementary Fig. 7b).

**Fig. 2.**
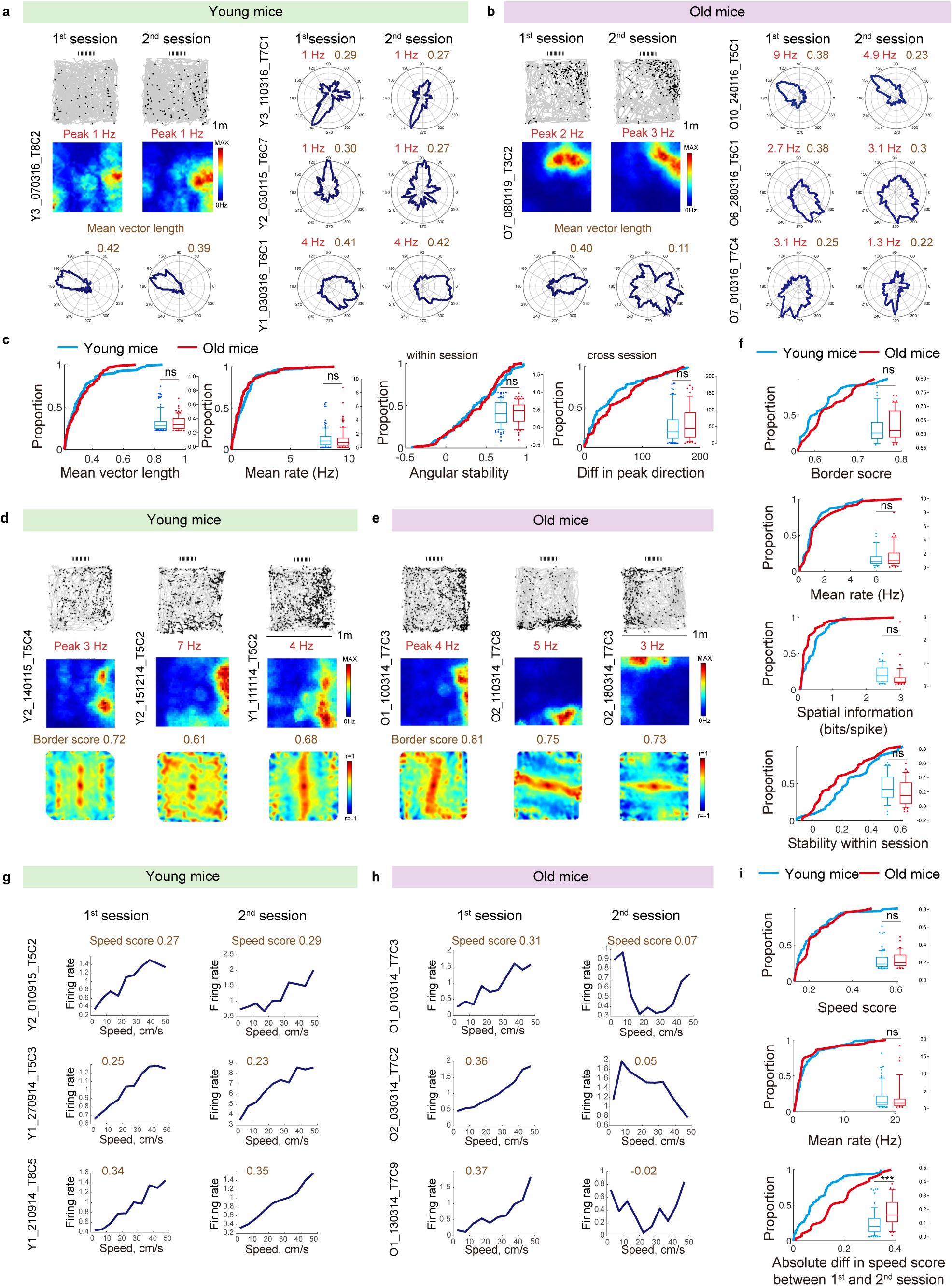
Reduced stability in speed cells, but not in HD and border cells, in old mice. **a** Left, trajectory, rate maps, and polar plots of HD cells in young mice; cell numbers at left. Peak firing rate (red), MVL (brown). **b**, As in **a**, for old mice. **c** MVL, mean rate, and within-session angular stability of HD cells (young, n = 109; old, n = 84; linear mixed-effects model: MVL, SE = 0.017, t (191) =-0.41, P = 0.68; mean rate, SE = 0.20, t (191) =-0.35, P = 0.73; angular stability, SE = 0.081, t (191) = 0.54, P = 0.59); difference in peak direction across sessions (young, n = 86; old, n = 65; SE = 13.33, t (149) = 0.26, P = 0.79). **d** Trajectory, rate maps, and autocorrelograms of border cells in young mice. Border score (brown). **e** As in **d**, for old mice. **f** Border score, mean rate, spatial information, and spatial stability of border cells (young, n = 31; old, n = 40; linear mixed-effects model: border score, SE = 0.013, t (69) = 1. 06, P = 0.29; mean rate, SE = 0.35, t (69) = 0.82, P = 0.41; spatial information, SE = 0.10, t (69) =-1.25, P = 0.21; spatial stability, SE = 0.044, t (69) =-1.67, P = 0.09). **g** Speed–rate relationship plots of speed cells across two sessions; speed scores (brown). **h** As in **g**, for old mice. **i** Speed score and mean rate of speed cells (young, n = 82; old, n = 37; linear mixed-effects model: speed score, SE = 0.026, t (117) = 1.41, P = 0.16; mean rate, SE = 0.71, t (117) =-0.059, P = 0.95); absolute difference in speed scores across sessions (young, n = 67; old, n = 31; SE = 0.020, t (96) =-3.93, P = 1.59×10⁻⁴). Data are median (center line), 25th–75th percentiles (box bounds), and 10th–90th percentiles (whiskers). P-values for fixed effects in the linear mixed-effects models were derived from two-sided t-tests. ***P < 0.001; ns, not significant. Source data are provided as a Source Data file.

Border cells likewise showed no detectable changes in border score (young: 0.62 ± 0.01 vs. old: 0.63 ± 0.01; *P* = 0.2944), mean firing rate (1.38 ± 0.23 vs. 1.67 ± 0.26; *P* = 0.4145), spatial information (0.49 ± 0.064 vs. 0.36 ± 0.075; *P* = 0.2147) or within-session spatial stability (0.26 ± 0.034 vs. 0.19 ± 0.03; *P* = 0.099) (Fig. 2d–f), and this stability persisted across sessions in both groups (Supplementary Fig. 8a). The functional identity of border cells in old mice was confirmed by the formation of a new border field after insertion of a freestanding wall in the recording box (Supplementary Fig. 8b).

### Speed cells show reduced speed-rate stability in old mice

We previously found that impaired grid coding after inactivation of parvalbumin (PV) interneurons in the MEC was coupled with disrupted speed coding of speed cells ^39^, raising the possibility that the impaired spatial selectivity of grid cells in old mice reflects impaired integration of speed information. Speed scores did not differ between age groups (young: 0.22 ± 0.011 vs. old: 0.23 ± 0.015; *P* = 0.16; Fig. 2g–i), nor did the mean firing rates of speed cells (young: 3.03 ± 0.37 Hz vs. old: 2.98 ± 0.69 Hz; *P* = 0.95; Fig. 2i). However, the absolute difference in speed score between the second session and baseline was significantly larger in old mice (young: 0.10 ± 0.011 vs. old: 0.18 ± 0.019; *P* = 1.59 × 10^-4^; Fig. 2i), indicating that speed-rate relationships in old mice were unstable (Fig. 2h and Supplementary Fig. 9a).

### Theta and slow Gamma power were decreased in old mice

The spatial periodicity of grid cells is often correlated with theta oscillations (6-12 Hz) ^40,41^. To examine whether the local field potential (LFP) in the MEC changes during aging, we analysed theta (6–12 Hz) and slow gamma (20–30 Hz) power in young (211 sessions) and old (133 sessions) mice, using a linear mixed-effects model with age as a fixed effect, running speed as a covariate and mouse identity as a random intercept (Theta Power ∼ Age + Running Speed + (1│Mouse)). Old mice exhibited significant reductions in theta power (young: 956.95 ± 52.26 vs. old: 554.05 ± 49.90; *P* = 1.43 × 10^-12^) and slow gamma power (young: 88.40 ± 7.67 vs. old: 72.18 ± 5.85; *P* = 0.025) (Supplementary Fig. 9b–d). Thus, aged mice exhibit reduced LFP power independent of running speed, particularly in the theta band, which is critical for grid cell firing ^41^.

### Aging alters gene expression uniquely in the MEC

To examine gene expression changes in the MEC during aging, we compared young and old C57BL/6J mice using 10x Genomics Visium technology (Fig. 3a). Spatial transcriptomic analyses were performed on 20 sagittal brain sections spanning the lateral to middle MEC from 2 old (>22 months) and 2 young (3–4 months) mice (Fig. 3b and Supplementary Fig. 10a), yielding 50752 spots (25493 young, 25259 old) with a median of 5398 genes per spot (Supplementary Fig. 10b). After batch correction, spots from all 4 mice were uniformly distributed by UMAP (Fig. 3c). Within the MEC dataset, 4033 spots (1633 young, 2400 old) with 5719 ± 20.16 (median ± SEM) genes per spot were identified (Supplementary Fig. 10c).

**Fig. 3.**
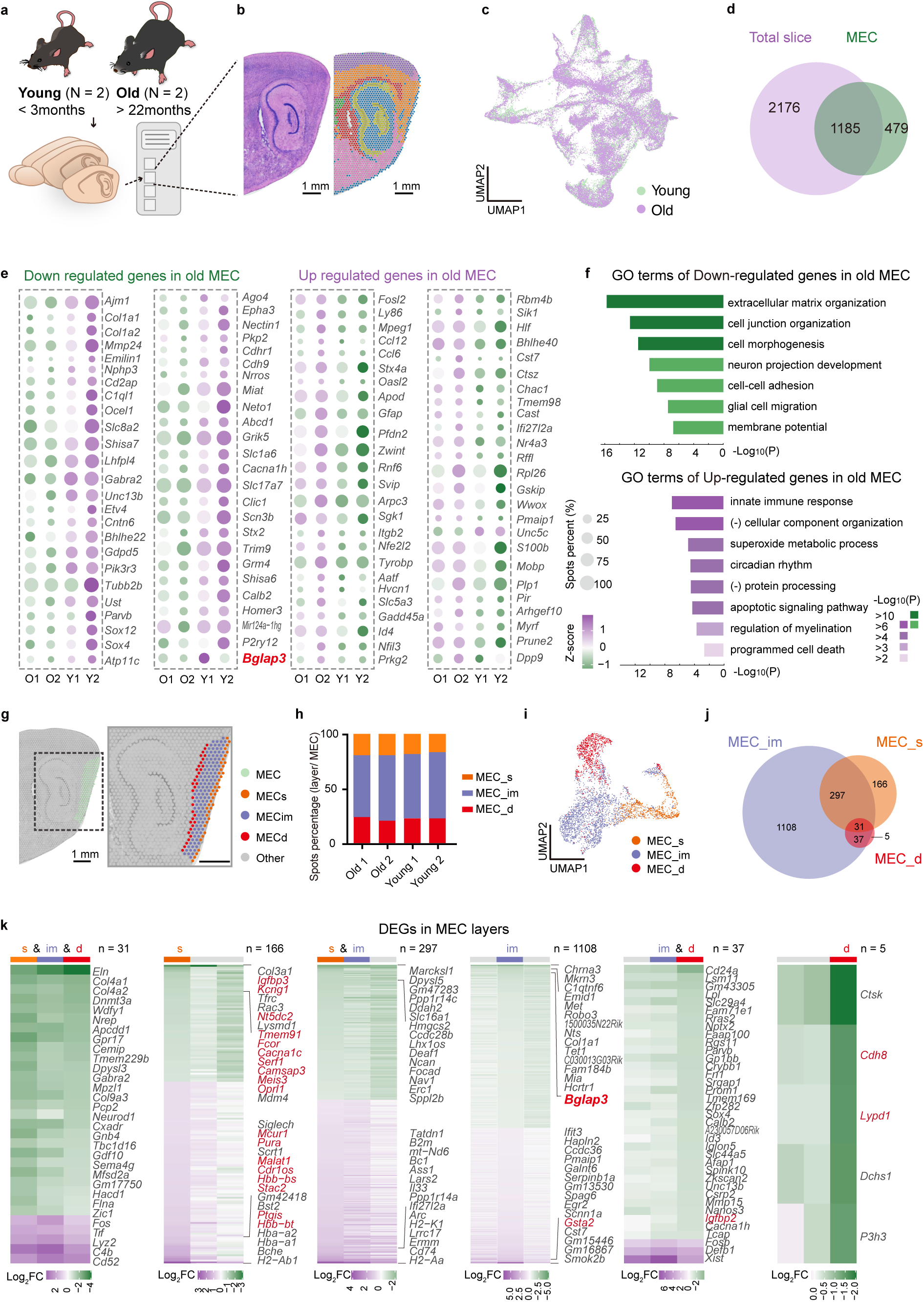
Changes in MEC gene expression profiles during aging using 10x Genomics Visium technology. **a** Streamline schematic of the 10x Genomics Visium experiment. **b** Sagittal sections of young mice brain overlaid with spatial clusters; Scale bar: 1 mm. **c** Uniform manifold approximation and projection (UMAP) map of the factor activities colored by mice from different ages. **d** Venn diagram illustrating the overlap of DEGs between young and old mice across the entire slice and the MEC. **e** 50 selected down or up-regulated genes in old MEC are displayed in dot plot. Dot size shows percentages of spots where that specific gene is expressed in whole MEC spots. Expression levels of genes are indicated by Z-score. **f** Bar graph showing the enrichment GO terms of down and up-regulated genes in old MEC, using a discrete color scale to represent statistical significance (-Log10(P)). A one-sided (cumulative) hypergeometric test to identify significantly enriched GO/pathway terms. **g** Example of manually defining MEC and its sublayers based on H&E staining brain slice: MEC_s, MEC_im and MEC_d. Representative images from n = 20 independent experiments with similar results. **h** Percentage bar plot showing distributions of spots from three sublayers of MEC among all the MEC spots in four mice samples. **i** UMAP of MEC spatial spots colored by their assigned regions. **j** Venn diagram showing the number of DEGs of old and young mice discovered in three sublayers of MEC. **k** DEGs are found from only one, two or both layers in MEC. The first, fifth and sixth columns show the whole genes and the rest show the top 15 genes from both up and down-regulated DEGs. Genes that were not discovered in the entire MEC DEGs lists are highlighted in red. Lower scores of Log2 Fold Change represent a lower expression level of DEGs in old mice. Differential expression analysis was performed using DESeq2, with statistical significance assessed by two-sided Wald tests. Source data are provided as a Source Data file.

Comparing old and young mice, we detected 3361 differentially expressed genes (DEGs) across the entire brain sections and 1664 DEGs in the MEC, of which 479 were differentially expressed in the MEC only (Fig. 3d and Supplementary Table 1). Bulk RNA sequencing of the MEC from 3 old and 3 young mice yielded 573 DEGs consistent with the Visium data (Supplementary Fig. 10d, e and Supplementary Table 2). The 50 most up-and downregulated MEC genes are shown in Fig. 3e, and 28 of these DEGs (8 up, 20 down) were verified by reverse transcription quantitative PCR (RT-qPCR) (Supplementary Fig. 11). Several genes upregulated in old mice have reported protective functions: *Sgk1* and *Apod* maintain nerve integrity and rescue the neural cytoskeleton ^42,43^, and *Bhlhe40* is a transcriptional regulator that enhances synaptic plasticity ^44^. Among the downregulated genes, *Tubb2b*, *Neto1* and *Calb2* have been associated with cognitive deficits, and their mutation or knockdown impairs long-term potentiation and neural excitability ^45–47^.

GO enrichment analysis of the 1664 DEGs (754 up and 910 downregulated) (Fig. 3f) revealed elevated immune and oxidative stress response pathways, indicating imbalance and cumulative damage in the central nervous system during aging ^48^, together with upregulated signaling related to the positive regulation of cell death and apoptosis. In contrast, signaling related to extracellular matrix organization, cell morphogenesis and cell junction organization was downregulated in the MEC of old mice, suggesting that dysfunction of these pathways is related to impaired MEC function during aging.

To examine gene expression across MEC layers, we delineated superficial (MEC_s), intermediate (MEC_im) and deep (MEC_d) layers based on the anatomical structure revealed by hematoxylin and eosin (H&E) staining (Fig. 3g, h and Supplementary Fig. 12a). UMAP dimensional reduction of spots from old and young MEC revealed three continuous sequential sublayer clusters (Fig. 3i and Supplementary Fig. 12b). To validate this assignment, we mapped known MEC markers including *Calb1* and *Reln* (layer II), *Oxr1* (layer III) and *Cux2* (upper cortical layers), based on previous *in vivo* functional studies ^49–52^, together with deep layer markers (*Plch1*, *Rmst*, *Cplx3* and *Cobll1*) reported in previous scRNA-seq studies ^31,53^ (Supplementary Fig. 12c). Comparing old and young mice, we discovered 166, 1108 and 5 layer-specific DEGs in MEC_s, MEC_im and MEC_d, respectively, along with 31 genes differentially expressed across all three layers, 297 genes common to MEC_s and MEC_im, and 37 genes shared between MEC_im and MEC_d (Fig. 3j and Supplementary Table 3). This layer-resolved analysis revealed DEGs absent from the whole-MEC list (highlighted in red, Fig. 3k), indicating sublayer-specific molecular changes during aging.

### *Bglap3*+ neurons are critical for spatial coding of the MEC

Through the 10x Genomics Visium experiment, we identified *Bglap3* as a gene specifically expressed in the MEC (Fig. 3e, k and Fig. 4a), with significantly fewer *Bglap3*+ spots in old mice (young: 68.36 ± 19.80 vs. old: 30.58 ± 2.72; Linear mixed-effects model, SE = 12.05, t (18) = 3.08, P = 6.43 × 10^-3^; Fig.4b). *In situ* hybridization (ISH) confirmed the unique regional expression of *Bglap3* in the MEC (Fig. 4c, d), particularly in layer III (Fig. 4e), and within NeuN+ populations (Fig. 4f–h). Mouse single-cell RNA-sequencing (scRNA-seq) data from Yao et al. (2023) ^54^, showed that *Bglap3* is predominantly expressed in the L2/3 IT ENT glutamatergic (Glut) subclass and superclass populations (Supplementary Fig. 14a–e). Using a detection threshold of UMI ≥ 2, we identified 28623 ENT cells expressing *Oxr1* and 113 cells expressing *Bglap3*; all *Bglap3*+ cells co-expressed *Oxr1* (Supplementary Fig. 14f, g), indicating that *Bglap3* defines a small subpopulation within the *Oxr1*+ population. *Bglap3* showed partial overlap with the layer II marker *Tmem132e*, greater overlap with the layer III marker *Cd44*, and no overlap with the deep layer marker *Nxph4* (Supplementary Fig. 14h). Together, these findings indicate that *Bglap3* marks a subpopulation of layer III glutamatergic neurons in the MEC and is more specific than *Oxr1*, which is broadly expressed across PV and SST interneurons, oligodendrocytes, OPCs and microglia (Supplementary Fig. 14f).

**Fig. 4.**
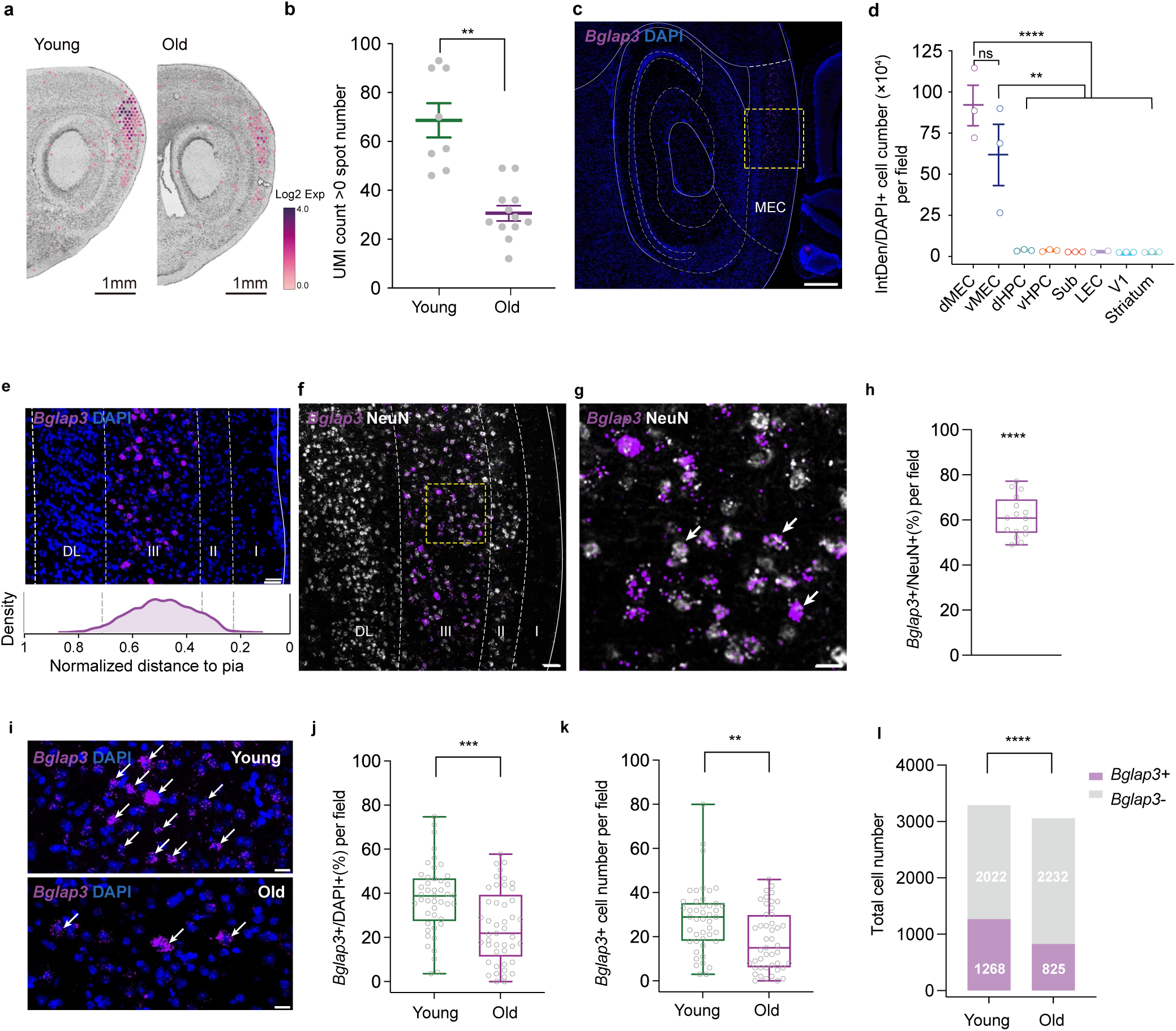
The number of *Bglap3*+ cells is decreased in old mice. **a** Expression of *Bglap3* in MEC. **b** Number of *Bglap3*+ spots (n = 8 fov, N = 2 young mice; n = 12 fov, N = 2 old mice; linear mixed-effects model, P = 6.43×10⁻³). **c** Expression of *Bglap3* in MEC by ISH. Pink, *Bglap3*; blue, DAPI. **d** *Bglap3* expression across regions (d/v, dorsal/ventral; Sub, subiculum; LEC, lateral entorhinal cortex; n = 3 fov, N = 3 young mice; one-way ANOVA, F (7,15) = 18.19, P = 2.86 × 10^-6^; two-sided Tukey’s multiple comparisons: dMEC vs. vMEC, P = 0.2288; dMEC vs. dHPC/vHPC/Sub/LEC/V1/Striatum, P < 0.0001; vMEC vs. dHPC/vHPC/Sub/LEC/V1/Striatum, P < 0.01). **e** Expression of *Bglap3* in MEC (top); *Bglap3*+ cell location relative to pia (n = 952 cells, N = 3 mice) (bottom). DL, deep layer; I–III, layers 1–3. **f** ISH of *Bglap3* co-stained with NeuN in MEC layer III; amplified view in **g**. Pink, *Bglap3*; white, NeuN. **h** Percentage of *Bglap3*+ cells among neurons (n = 16 fov, N = 3 mice; median = 60.81; two-sided Wilcoxon signed-rank test, W = 136, P= 3.05 × 10^-5^). **i** *Bglap3*+ cells in MEC of old and young mice; white arrows, *Bglap3*+ cells. **j** Percentage of *Bglap3*+ cells in old vs. young mouse MEC (n = 45 fov, N = 4 young mice; n = 45 fov, N = 4 old mice; linear mixed-effects model, P = 2.68×10⁻⁴). **k** Number of *Bglap3*+ cells in old vs. young MEC (linear mixed-effects model, P = 1.70×10⁻³). **l** Ratio of *Bglap3*+/*Bglap3*− cells (Pearson’s chi-square test, χ² (1, N = 6347) = 95.71, P = 1.33 ×10⁻^22^). Data in **b** and d are mean ± s.e.m.; data in **h, j**, and **k** are median with min-to-max range. P-values for fixed effects in the linear mixed-effects models were derived from two-sided t-tests. *P < 0.05; **P < 0.01; ***P < 0.001; ****P < 0.0001; ns, not significant. Scale bars: **a**, 1 mm; **c**, 500 µm; **e**, **f**, **g**, 50 µm; **i**, 20 µm. Source data are provided as a Source Data file.

We next compared the percentage of MEC cells expressing *Bglap3* between young (3-4 months) and old (>22 months) mice. The percentage of *Bglap3*+ cells per field of view was significantly decreased in old mice (young: 37.80 ± 2.37% vs. old: 24.94 ± 2.47%; Linear mixed-effects model, SE = 3.39, t (88) =-3.80, P = 2.68 × 10^-4^; Fig.4i, j), accompanied by an absolute decrease in the number of *Bglap3*+ cells (young: 28.18 ± 2.28 vs. old: 18.33 ± 2.05; Linear mixed-effects model, SE = 3.04, t (88) =-3.24, P = 1.70 × 10^-3^; Fig.4k), whereas the total number of DAPI+ cells per field of view remained unchanged (young: 73.11 ± 2.71 vs. old: 67.93 ± 2.3; Linear mixed-effect model, SE = 3.52, t (88) =-1.47, P = 0.1455;

Supplementary Fig.15a, b). The relative proportion of *Bglap3*+ and *Bglap3*-cells was significantly influenced by age (Pearson’s chi-square test, P = 1.33 ×10⁻^22^; Fig.4l). Deconvolution of the Visium data using Cell2location ^55^ likewise revealed a significant decrease in the L2/3 IT ENT Glut subclass fraction in the MEC of old mice (Supplementary Fig. 15c), the subclass with predominant *Bglap3* expression (Supplementary Fig. 14). Altogether, these results demonstrate that *Bglap3*+ neurons are localized to the MEC layer III neuronal population and decrease in number during aging, implying a potential relationship between age-related changes in *Bglap3*+ neurons and MEC dysfunction.

To further investigate the function of *Bglap3*+ neurons, we generated a *Bglap3-Cre* gene knock-in mouse line (Supplementary Fig. 16a). ISH showed co-localization of *Bglap3* with *Cre* (Supplementary Fig. 16b, c), and Cre-driven reporter fluorescence was present only in the MEC and not in neighboring cortex (Supplementary Fig. 16d), confirming the validity of the line. Injecting AAV carrying DIO-mCherry into the MEC of *Bglap3-Cre* mice and co-staining with NeuN and GAD67 showed that only 0.1% of NeuN+ cells expressed both mCherry and GAD67 (Supplementary Fig. 16e, f), excluding inhibitory neurons as a source of *Bglap3* expression. Having established that *Bglap3* is expressed in excitatory neurons (Supplementary Fig. 14d, e), we injected AAV2/9-hSyn-DIO-Synaptophysin-EGFP into the MEC of *Bglap3-Cre* mice to label axon terminals (Supplementary Fig. 17a); *Bglap3*+ neurons projected mainly to the ipsi-and contralateral hippocampus, subiculum and MEC (Supplementary Fig. 17b). We then labelled *Bglap3*+ cells with Cre-dependent TVA and G proteins (AAV2/8-DIO-EGFP-TVA and AAV2/8-DIO-oRVG) followed by rabies virus injection (EnvA-RVdG-mCherry) in the MEC (Supplementary Fig. 17c, d). Presynaptic neurons projecting to *Bglap3*+ neurons were mainly localized in the ipsilateral MEC, parasubiculum, postsubiculum and subiculum, and only sparsely in the hippocampus, retrosplenial cortex and contralateral MEC (Supplementary Fig. 17e), with a higher probability of recurrent connections among presynaptic cells located in superficial MEC (Supplementary Fig. 17f).

To investigate the *in vivo* function of *Bglap3*+ cells and their role in the MEC microcircuit, we conducted two-photon calcium imaging using a miniaturized two-photon microscope (TPM) according to recently published methods ^56^. We injected a mixture of AAV2/9-CaMKIIa-DIO-hM4D(Gi)-mCherry and AAV2/9-hSyn-GCaMP6s into the right MEC of *Bglap3-Cre* mice (one mouse received only AAV2/9-EF1α-DIO-GCaMP6s) (Fig. 5a, b and Supplementary Fig. 18a, b). Mice freely explored an 80 cm × 80 cm open field equipped with spatial cues during two baseline sessions (BL1 and BL2) and one session following deschloroclozapine (DCZ) injection on Day 1, with two recovery sessions (RC1 and RC2) recorded on Day 2 (Supplementary Fig. 19a). All mice showed average trajectory coverage above 90% and no anxiety-related behavior across the five sessions (Supplementary Fig. 19b-h). By imaging *Bglap3*+ cells tagged with Cre-dependent GCaMP6s or aligning mCherry-expressing *Bglap3*+ cells with GCaMP6s-expressing cells (see Methods; Fig. 5c, d and Supplementary Fig. 20), we identified 278 *Bglap3*+ and 669 *Bglap3*-neurons. Both populations contained mixed spatial coding cell types (*Bglap3*+ vs. *Bglap3*-: 1.80% vs. 11.66% grid cells, 8.63% vs. 9.72% border cells, 10.79% vs. 16.44% HD cells, 17.27% vs. 15.40% spatial cells, 38.12% vs. 25.41% unstable cells and 23.38% vs. 21.38% unclassified cells), with decreased percentages of grid and HD cells and an increased percentage of unstable cells within the *Bglap3*+ population (*Bglap3+* vs. *Bglap3-*; grid cell: P = 1.02 × 10^-6^, border cell: P = 0.60, HD cell: P = 0.02, spatial cell: P = 0.47, unstable cell: P = 8.77 × 10^-5^, unclassified cell: P = 0.49, Pearson’s chi-squared test; Fig.5e, f). Cell type definitions were restricted to cells that were stable between the two baseline sessions (see Methods). *Bglap3*+ and *Bglap3*-cells did not differ in baseline grid scores, in the correlation of HD tuning curves, or in the spatial information of spatial tuning cells (SPT); however, *Bglap3*-spatial tuning cells showed a higher correlation of rate maps across baseline sessions (Supplementary Fig. 21a-c).

**Fig. 5.**
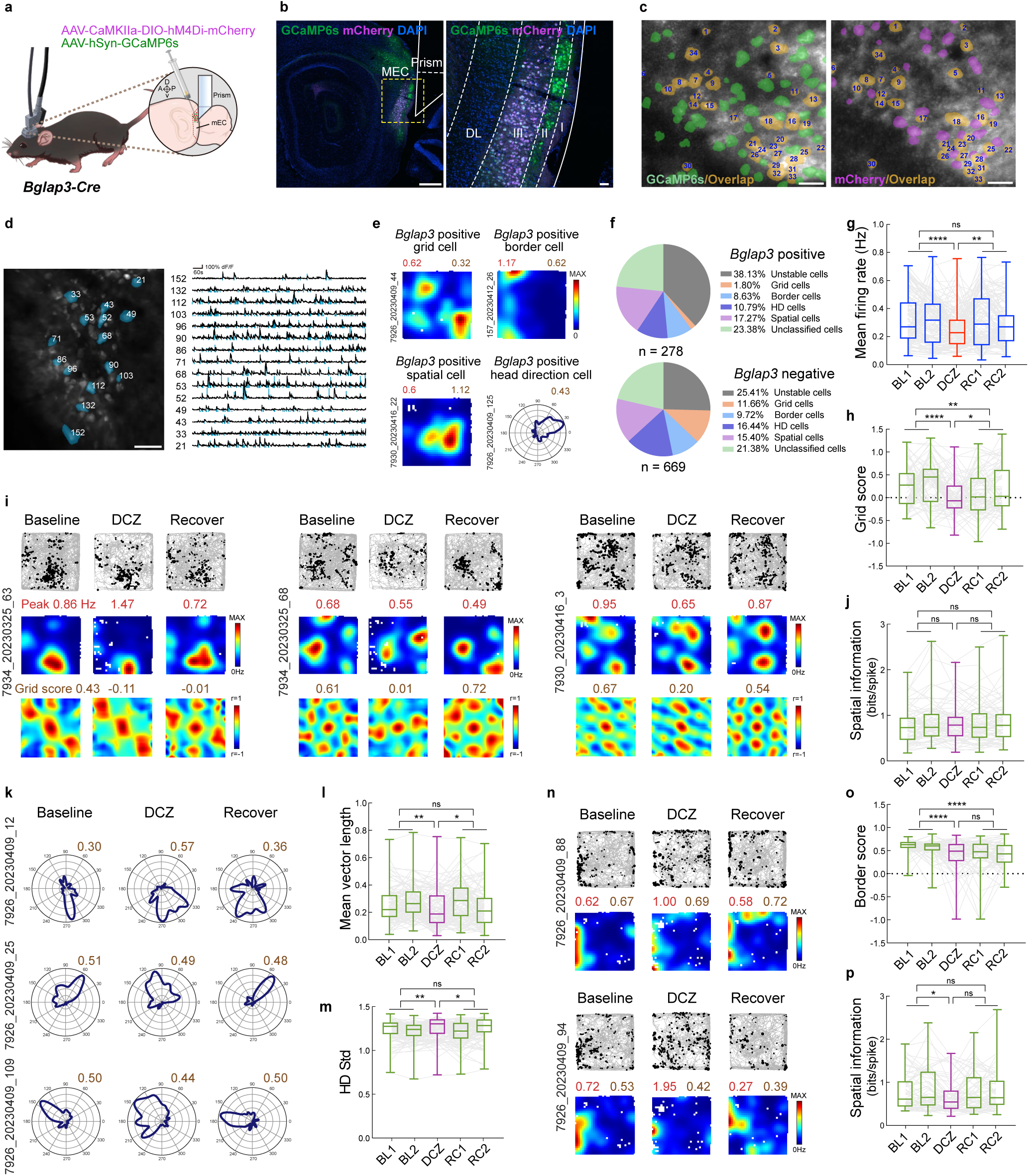
Inactivation of *Bglap3+* cells impaired grid and spatial coding in MEC. **a** 2-photon imaging of *Bglap3*+ cells in MEC. **b** GCaMP6s (green) and mCherry (magenta) expression in MEC; LI–LIII, layers 1– 3; DL, deep layer. **c** FOV of TPM; green, GCaMP6s+; pink, mCherry+; orange (numbered), overlapping cells. **d** FOV of Ca²⁺ imaging and ΔF/F traces. **e** Representative grid, border, HD, and unclassified cells among *Bglap3*+ cells. **f** Proportion of functional cell types in *Bglap3*+ vs. *Bglap3*− populations. **g** Mean firing rate of *Bglap3*+ cells (n = 132 cells, N = 5 mice; P = 2.89×10⁻⁴; BL–DCZ, P = 6.25×10⁻⁵; DCZ–RC, P = 2.20×10⁻³; BL–RC, P = 0.2425). **h** *Bglap3*− grid scores after *Bglap3*+ inhibition (n = 77 cells, N = 5 mice; P = 1.92×10⁻⁵; BL–DCZ, P = 8.20×10⁻⁶; DCZ–RC, P = 0.047; BL–RC, P = 2.03×10⁻³). **i** Trajectory maps, rate maps, and autocorrelograms of *Bglap3*− grid cells; peak rates (red) and grid scores (brown); cell numbers at left. **j** Spatial information of *Bglap3*− grid cells (P = 0.9028; BL–DCZ, P = 0.8126; DCZ– RC, P = 0.8976; BL–RC, P = 0.6542). **k** *Bglap3*− HD cells; MVL (brown). **l, m** MVL and circular s.d. of *Bglap3*− HD cells (n = 110 cells, N = 6 mice; MVL, P = 5.50×10⁻³ [BL–DCZ, P = 1.39×10⁻³; DCZ–RC, P = 0.014; BL–RC, P = 0.3584]; Std: P = 0.01 [BL–DCZ, P = 3.29×10⁻³; DCZ–RC, P = 0.016; BL–RC, P = 0.4993]). **n-p** Rate maps, border scores, and spatial information of *Bglap3*− border cells (n = 65 cells, N = 6 mice; border score, P = 7.74×10⁻⁸ [BL–DCZ, P = 2.10×10⁻⁵; DCZ–RC, P = 0.9306; BL–RC, P = 1.33×10⁻⁷]; spatial information, P = 0.0365 [BL–DCZ, P = 0.0102; DCZ–RC, P = 0.0706; BL–RC, P = 0.3464]). Data, median with min-to-max whiskers; P-values, two-sided linear mixed-effects ANOVA (fixed-effect t-tests) with pairwise post-hoc contrasts. *P<0.05; **P<0.01; ****P<0.0001; ns, not significant. Scale bars: **b**, 500 µm; **c**, 50 µm; **d**, 20 µm. Source data are provided as a Source Data file.

After DCZ injection, the mean firing rate of *Bglap3*+ cells was significantly decreased (BL: 0.32 ± 0.010 Hz vs. DCZ: 0.26 ± 0.012 Hz vs. RC: 0.31 ± 0.010 Hz; linear mixed-effects ANOVA, F (2, 657) = 8.25, *P* = 2.89 × 10^-4^; BL vs. DCZ: SE = 0.016, t (657) =-4.03, P = 6.25 × 10^-5^, DCZ vs. RC: SE = 0.016, t (657) = 3.07, P = 2.20 × 10^-3^; Fig. 5g and Supplementary Fig. 21d, e), indicating successful inactivation, and it recovered to at least baseline level after 24 hours (BL vs. RC: SE = 0.013, t (657) =-1.17, P = 0.2425, Linear mixed-effect ANOVA with pairwise post-hoc contrasts). We observed no significant decrease in the mean firing rate of *Bglap3*-neurons (Supplementary Fig. 21f, g). c-Fos immunostaining of brain slices from DCZ-and saline-injected mice revealed a significant reduction in c-Fos intensity in *Bglap3*+ neurons and in layer II *Bglap3*-neurons, but not in layer III *Bglap3*-neurons (Supplementary Fig. 22a–d).

Characterizing the firing properties of *Bglap3*-cells after silencing *Bglap3*+ neurons, we observed a significant decrease in the grid scores of *Bglap3*-grid cells (BL: 0.29 ± 0.039 vs. DCZ: 0.21 ± 0.046 vs. RC: 0.14 ± 0.039, Linear mixed-effect ANOVA: F (2, 382) = 11.17, P = 1.92 × 10^-5^, with pairwise post-hoc contrasts: BL vs. DCZ: SE = 0.059, t (382) =-4.52, P = 8.20 × 10^-6^, DCZ vs. RC: SE = 0.059, t (382) = 1.98, P = 0.047, BL vs. RC: SE = 0.049, t (382) =-3.11, P = 2.03 × 10^-3^; Fig. 5h). This effect persisted when running speed was included as a covariate (Grid score ∼ Condition + Running Speed + (1│Mouse); condition: F (2, 381) = 5.50, *P* = 4.4 × 10^-3^; Grid score BL vs. DCZ: pairwise post hoc contrasts, t (381) = −3.08, *P* = 2.2 × 10^-3^), whereas running speed itself was not a significant predictor (F (1, 381) = 0.0031, *P* = 0.9554; Supplementary Fig. 19d). Grid scores remained consistent between the two 40-minute baseline sessions (BL1 vs. BL2: *P* > 0.9999, one-way ANOVA), indicating that the decrease was not a result of temporal drift. In firing rate maps and autocorrelograms, the regular hexagonally arranged firing fields were impaired, with certain grids disappearing or becoming irregular after DCZ injection (Fig. 5i and Supplementary Fig. 23a) and a significant drop in the correlation of firing rate maps between DCZ and baseline sessions (Supplementary Fig. 23b). The spatial information of *Bglap3*-grid cells was unchanged (BL: 0.85 ± 0.033 vs. DCZ: 0.86 ± 0.046 vs. RC: 0.86 ± 0.034, Linear mixed-effect ANOVA: F (2, 382) = 0.10, P = 0.9028; Fig.5j). Random subsampling of grid cells with 100 repeated statistical tests showed that the effect became statistically significant at a sample size of approximately 20 cells, supporting the robustness of these findings across the grid cell population (Supplementary Fig. 24a, b).

Following DCZ injection, the MVL of *Bglap3*-HD cells decreased (BL: 0.26 ± 0.0082 vs. DCZ: 0.22 ± 0.013 vs. RC: 0.25 ± 0.0098, Linear mixed-effect ANOVA: F (2, 547) = 5.25, P = 5.50 × 10^-3^, with pairwise post-hoc contrasts: BL vs. DCZ: SE = 0.013, t (547) =-3.21, P = 1.39 × 10^-3^, DCZ vs. RC: SE = 0.013, t (547) = 2.46, P = 0.014, BL vs. RC: SE = 0.011, t (547) = - 0.92, P = 0.3584; Fig.5 k-l) and the dispersion around their preferred direction increased (BL: 1.21 ± 0.0072 vs. DCZ: 1.24 ± 0.011 vs. RC: 1.22 ± 0.0085, Linear mixed-effect ANOVA: F (2, 547) = 4.52, P = 0.011, with pairwise post-hoc contrasts: BL vs. DCZ: SE = 0.011, t (547) = 2.95, P = 3.29 × 10^-3^, DCZ vs. RC: SE = 0.011, t (547) =-2.40, P = 0.016, BL vs. RC: SE = 0.0092, t (547) = 0.68, P = 0.4993; Fig.5m and Supplementary Fig. 25a, b), indicating a decline in directional selectivity. For *Bglap3*-border cells, firing event locations became more chaotic with less boundary selectivity, resulting in a significant decrease in border scores (BL: 0.57 ± 0.014 vs. DCZ: 0.40 ± 0.040 vs. RC: 0.39 ± 0.028, Linear mixed-effect ANOVA: F (2, 322) = 17.23, P = 7.74 × 10^-8^, with pairwise post-hoc contrasts: BL vs. DCZ: SE = 0.039, t (322) =-4.32, P = 2.10 × 10^-5^, DCZ vs. RC: SE = 0.039, t (322) =-0.087, P = 0.9306, BL vs. RC: SE = 0.032, t (322) =-5.39, P = 1.33 × 10^-7^; Fig.5n, o and Supplementary Fig.25c) and reduced spatial information relative to baseline (BL: 0.80 ± 0.040 vs. DCZ: 0.67 ± 0.048 vs. RC: 0.76 ± 0.037, Linear mixed-effect ANOVA: F (2, 322) = 3.34, P = 0.0365, with pairwise post-hoc contrasts: BL vs. DCZ: SE = 0.050, t (322) =-2.58, P = 0.0102, DCZ vs. RC: SE = 0.050, t (322) = 1.81, P = 0.0706, BL vs. RC: SE = 0.041, t (322) =-0.94, P = 0.3464; Fig.5p). The spatial information of *Bglap3*-spatial cells was unchanged, although their rate map correlation decreased after DCZ injection (Supplementary Fig. 25d–f).

To test whether the decrease in spatial tuning during the DCZ session reflected a drug effect, we performed calcium imaging in three mice co-injected with AAV2/9-CaMKIIa-DIO-mCherry and AAV2/9-hSyn-GCaMP6s, following the same imaging procedures (Supplementary Fig. 26a, b). The mean firing rates of both *Bglap3*+ and *Bglap3*-cells showed no significant change after DCZ administration (Supplementary Fig. 26c, d), and no significant alterations were detected in grid, HD, border or spatial tuning indices for either cell type (Supplementary Fig. 26e–h). Collectively, these data indicate a pivotal function of *Bglap3*+ neurons in the spatial circuits of the MEC, and suggest that changes in these neurons may relate to the impaired spatial cognition that occurs during aging.

## Discussion

In this study, the proportion of grid cells in old mice was significantly lower than in young mice, and reduced grid scores were associated with less consistent coding across sessions. Speed cells exhibited reduced stability in old mice, whereas HD and border cells did not. Gene expression analysis identified candidate biological processes related to impaired MEC function during aging and revealed a population of MEC layer III cells expressing *Bglap3* whose number was significantly decreased in old mice. Selectively labeling and inactivating *Bglap3*+ neurons disrupted the tuning of spatial cells, indicating an important role for *Bglap3*+ cells in spatial coding and a potential contribution to the unstable firing of spatial coding cells during aging.

Spatial learning and memory are dramatically impaired during aging across numerous species ^1,2^, which may be caused by impaired function of the MEC and hippocampus. Grid cells, as key components of the MEC circuit, play an essential role in spatial computation and memory ^9^; their unstable firing in old mice could underlie age-related navigational deficits, because inaccurate grid coding causes errors in path integration and impairs spatial computation ^14^. Our LFP recordings revealed a significant decrease in theta and slow gamma power in old mice, indicating a network-level change that is correlated with the irregular firing of grid cells. Mean running speed did not differ between age groups, and the reduction in oscillatory power persisted after controlling for speed, excluding the possibility that reduced theta power simply reflects slower running. Inactivation of the medial septum (MS) reduces theta rhythm and disrupts grid firing in the MEC ^40,41^. Although we cannot exclude neuronal loss and other chronic changes in the aged MS that could eventually affect grid firing ^57,58^, the unstable firing of other spatial coding cells such as speed cells, which are often less affected by theta rhythm, supports the conclusion that local changes occur in the MEC during aging. The reduction in slow gamma power may also explain impaired spatial cognition in aged rodents through decreased coherence between the hippocampus and MEC ^59^, since gamma coordination between these regions is crucial for memory coding and retrieval ^60^. It remains to be determined whether age-related changes in grid cells precede those in place cells or vice versa, and whether changes in the theta and gamma bands contribute similarly to both populations. Overall, altered spatial representations in the hippocampus and MEC, including the instability of place cells and grid cells, are likely major reasons for reduced spatial learning and memory in old age.

Unlike grid and speed cells, HD cells and border cells in old mice did not show instability across or within sessions (Fig. 2c, f). Notably, a previous study using the J20 transgenic mouse model of familial AD reported decreased stability in grid cells but no deficits in HD cells ^61^, and our findings in healthy aging mice align with this observation. During development, border cells show adult-like fields from the outset, whereas grid cells fire irregularly until four weeks of age ^62,63^. A clear representation of borders depends more on tactile and visual input than on the speed and idiothetic input that grid cells and HD cells require for path-or angular-path integration ^64,65^, which may help explain why border cells are less vulnerable to aging than other spatial coding cells in the MEC.

Grid cell coding is influenced by various factors, and changes in external input can affect grid computation ^66^. Impaired grid coding may also result from local changes inside the MEC, as many signaling pathways were altered during aging (Fig. 3f). Multiple structural, molecular and cellular changes have been identified in the neurons of old animals ^67^: long-term potentiation (LTP) is difficult to induce and decays easily ^68,69^, whereas long-term depression (LTD) is activated normally ^69^. This is consistent with our gene expression data showing decreased expression of *Tubb2b*, *Neto1* and *Calb2*, whose mutation or knockdown impairs LTP and neural excitability ^45–47^. The instability we observed in grid cells of old mice could therefore be caused by declining LTP, since a shifted LTP/LTD balance would impair the consolidation of spatial memory and produce inaccurate spatial coding, leading to accumulated navigational errors through dysfunctional path integration ^70,71^.

Aged transgenic mice expressing mutant human tau in the entorhinal cortex show impairments in grid coding as well as spatial memory ^27^, and young human adults at risk for developing AD who carry the APOE-ε4 allele exhibit impaired grid cell-like activity in the MEC ^28^, raising the possibility of including grid cells in the early diagnosis of AD. In our study, we found no notable changes in the expression of several known AD-related genes previously reported in a study of AD patients ^72^ (Supplementary Fig. 13), indicating that the changes in grid coding in our system were not likely caused by AD-related symptoms. Our data support the possibility that changes in grid cells occur during normal aging regardless of the development of AD, which is consistent with results from healthy older humans ^26^.

In our gene expression analysis, most DEGs upregulated in old mice were related to cell injury and cell death, consistent with earlier reports ^67^, whereas downregulated DEGs were related to neural development and circuit maturation, indicating impaired homeostasis in the neural circuits of old mice. One of these DEGs, *Bglap3*, was selectively expressed in layer III of the MEC and marked a population whose cell number decreased during aging, pointing to a potential role for MEC layer III in aging.

In our axon tagging studies, *Bglap3*+ neurons projected to ipsi-and contralateral MEC superficial layers, subiculum and hippocampus, consistent with previous histological studies of MEC layer III neurons ^51,73–75^. Few studies have examined upstream projections to MEC layer III ^76,77^; in our rabies virus tracing experiment, *Bglap3*+ neurons received inputs from the ipsilateral MEC and other parahippocampal regions such as parasubiculum and postsubiculum, identifying candidate sources of the information these neurons encode. Our data also support recurrent connectivity in MEC superficial layers, which is believed to be crucial for HD cells and grid cells in forming attractor networks^78–81^. *In vivo* electrophysiological results show that most layer III cells in the MEC are non-spatial or spatially irregular, with a smaller percentage of grid cells ^73^, consistent with our findings in *Bglap3*+ neurons (Fig. 5f). Our combined chemogenetic silencing and two-photon calcium imaging revealed that inactivating *Bglap3*+ neurons caused irregular firing of grid cells and decreased the directional tuning of HD cells and the border tuning of border cells, indicating that *Bglap3*+ cells have a modulatory function in the local spatial network of the MEC. *In vitro* electrophysiological studies have found that layer III cells project more commonly to layer II stellate cells than to layer II pyramidal cells, and that layer II contains the highest number of grid cells ^82^. Our calcium imaging data therefore provide *in vivo* evidence for connections between layer III *Bglap3*+ neurons and MEC spatial coding cells, especially grid cells, which are most likely located in layer II ^83^. Layer III neurons are also necessary and sufficient to generate local up-down states (UDS), which are crucial for memory encoding, synaptic transmission and plasticity ^84,85^; our *in vivo* silencing of layer III *Bglap3*+ neurons provides further evidence supporting the pivotal role of MEC layer III neurons in MEC microcircuits. Given the decrease in *Bglap3*+ neurons during aging, it will be important to determine whether the irregular and unstable tuning of grid cells in aged mice is due to the loss of these neurons, and how *Bglap3*+ projections to hippocampal CA1 influence place cells, which also become unstable during aging. A limitation of our study is that it remains unclear whether *Bglap3*+ neurons represent a specific population within MEC layer III that selectively influences grid cells, or whether their effects reflect broader layer III activity. Nevertheless, layer III of the MEC comprises diverse neurons with distinct electrophysiological properties and axonal arborization patterns ^86^, suggesting that subpopulations may have specialized roles in the superficial MEC circuitry. It also remains unclear whether *Bglap3*+ neurons contact grid cells directly or indirectly, as silencing them may influence activity along the MEC III-CA1-MEC Vb-MEC II/III pathways ^87–89^. To further elucidate the role of *Bglap3*+ neurons within the MEC microcircuit, it will be important to investigate the molecular functions of *Bglap3* at both the gene and protein levels and to define the mechanisms by which these neurons influence spatial coding cells during aging.

## Methods

### Animals

Experiments were performed according to the Animal Welfare/Ethics guidelines in Peking University. The experiments were approved by the Animal Research Authorities of Peking University.The main text of the manuscript was initially written by the authors and subsequently revised with the assistance of AI (ChatGPT).

JAX^®^ C57BL/6J mice (The Jackson Laboratory, USA) were used in this study with the following characteristics: 21 young mice and 22 old mice, without sex bias, and with a weight of 22–35 g at implantation. 8 young (3-4 months) and 10 old (> 28 months) mice were used for tetrode recordings and for the 10x Genomics Visium analysis; 4 sections per mouse from the lateral to middle part of the MEC were collected from 2 old (>22 months old) and 2 young mice (3-4 months). For bulk RNA-seq analysis, 3 young (3-4 months) and 3 old (>22 months old) mice were analyzed. For ISH, 4 young (3-4 months) and 4 old (>22 months old) mice were used. Visual discrimination learning task was done on 3 old (>22 months old) and 4 young (3-4 months) mice. *Bglap3*-Cre knock-in mice were provided by Cyagen US Inc. by inserting P2A-Cre cassette downstream of *Bglap3*. In total, 22 *Bglap3*-Cre mice were used; among them, 3 mice were used for ISH and Cre expression verification, 9 mice were used for histology studies, 10 mice were used for *in vivo* calcium imaging studies. All mice were housed individually in transparent Plexiglass cages (35 cm × 30 cm × 30 cm) in a humidity-and temperature-controlled environment. All animals were maintained at 90% of free-feeding body weight on a 12-h light/12-h dark schedule. Testing occurred in the dark phase. *Bglap3*-Cre mice were assessed by PCR using primer sequences below: KI-F: 5’-CTCTTGTACAGTGTGGGAAGAGGAT-3’; KI-R: 5’-ATGTCCATCAGGTTCTTGCGA-3’; WT-F: 5’-GTTGTTCTGGGGTAGTCTCTATGACC-3’; WT-R: 5’-CTAAATTAACACAGGTACCAGGCG-3’.

### Surgery, virus injection, and electrode implantation

Before surgery, the animals were anesthetized with isoflurane (airflow: 0.8–1.0 L/min, 0.5–3% isoflurane, adjusted according to physiological condition). Isoflurane was gradually reduced from 3% to 1%. Depth of anesthesia was examined by testing tail and pinch reflexes as well as breathing.

Upon induction of anesthesia, the animal was fixed in a Kopf stereotaxic frame for implantation of tetrodes. Holes were drilled in the skull above the right MEC. For virus injection in MEC, a 10 μL NanoFil syringe (World Precision Instruments, Sarasota, FL, USA) and a 33-gauge beveled metal needle were used to inject viruses in the MEC with the injection site: 0.3 mm anterior to the transverse sinus, 3.1–3.6 mm from the midline (ML axis), 1.8–1.5, 1.5–1.3 mm below the dura. For primary somatosensory barrel cortex (S1BF) virus injection, the injection site was 2.49 mm from the midline (ML axis), 1.0 mm posterior to Bregma and 0.35 mm below the dura. Injection volume (0.4–0.8 μL at each location) and flow rate (0.08 μL/min) were controlled with a Micro4™ Microsyringe Pump Controller (World Precision Instruments). After injection, the needle was left in place for 10 min before being slowly withdrawn.

For electrode implantation, each mouse was implanted with a VersaDrive-4 microdrive (NeuraLynx, Bozeman, MT, USA), connected to 4 tetrodes. The tetrodes were made of 17 µm polyimide-coated platinum-iridium (90%–10%) wire. The electrode tips were plated with platinum to reduce electrode impedances to approximately 100–250 kΩ at 1 Hz. The tetrodes were inserted 0.35–0.40 mm anterior of the transverse sinus, 3.2–3.5 mm lateral to the midline in the right hemisphere, and 0.8–1.2 mm below the dura, at a 5° angle in the sagittal plane, with electrode tips pointing in the posterior direction. The microdrives were secured to the skull with jewelers’ screws and dental cement. Two front screws were connected to ground.

### Electrode turning and recording procedures

Turning of the tetrodes started 2–3 d after the surgery. Data collection began within 3 weeks. Testing of control animals was alternated with testing of experimental animals. Before each recording trial, the mouse rested on a towel in a large flowerpot on a pedestal. The mouse was connected to the recording equipment via AC-coupled unity-gain operational amplifiers close to the head and a counterbalanced cable that allowed the animal to move freely. Over the course of 20–60 d, the tetrodes were lowered in steps ≤50 μm until well-separated single neurons could be recorded. Data were collected when the signal amplitudes exceeded four times the noise level (20–30 μV) and single units were stable for more than 1 h.

Recorded signals were amplified 8000–25,000 times and band-pass filtered between 0.8 and 6.7 kHz. Triggered spikes were stored to disk at 48 kHz (50 samples per waveform, 8 bits/sample) with a 32-bit time stamp (clock rate at 96 kHz). Electroencephalograms (EEGs) were recorded single-ended from one of the electrodes. The local field potential was amplified 3000–10,000 times, low-pass filtered at 500 Hz, sampled at 250 Hz, and stored with the unit data. Through a video camera, the recording system obtained the position of two light-emitting diodes (LEDs) on the headstage of the mouse. The LEDs were tracked individually at a rate of 50 Hz. The two LEDs were separated by 4 cm and aligned with the body axis of the mice.

Over the course of 3–6 weeks following surgery, the mice were first trained to run in a 1 m square black aluminum enclosure polarized by a cue card with white and black stripes. In mice with putative border cells, these trials were occasionally succeeded by a test in the same box with a 50 cm long and 50 cm high wall insert in the center of the box. Each recording session lasted 15 min. The interval between two recording sessions was 3-4 h. Running was motivated by scattering crumbs of chocolate at random locations in the box at 10–15 s intervals.

### Histological procedures and electrode positions

The mice received an overdose of pentobarbital and were subsequently perfused intracardially with saline followed by either 4% formaldehyde or 4% freshly depolymerized paraformaldehyde (PFA) in phosphate buffer. The brains were extracted and stored in the same fixative, and frozen sagittal sections (10 µm) were cut on a cryostat. Every 10^th^ section was stained with cresyl violet. All sections of the infected and implanted part of the MEC were collected. Tetrodes were identified and the tip of each electrode was found by comparison with adjacent sections.

### Tissue preparation for 10x Genomics Visium technology

The 10x Genomics Visium experiment was performed according to the company’s protocol (https://www.10Xgenomics.com/platforms/Visium) with the simplified streamline of experiments shown in Fig. 3a. Fresh frozen brain tissues were embedded, sectioned, and placed onto the Capture Area of the gene expression slide. Each Capture Area has thousands of barcoded spots containing millions of capture oligonucleotides with spatial barcodes unique to that spot. We utilized standard H&E fixation and staining techniques using a brightfield microscope to visualize tissue sections on slides. The tissue was permeabilized to release mRNA from the cells, which bound to the spatially barcoded oligonucleotides present on the spots. Reverse transcription was then performed to produce cDNA from the captured mRNA. The barcoded cDNA was then pooled for downstream processing to generate a sequencing-ready library. For formalin-fixed paraffin-embedded (FFPE) tissues, the tissue was permeabilized to release ligated probe pairs from the cells, which bound to the spatially barcoded oligonucleotides present on the spots. Spatial barcodes were added via an extension reaction. The barcoded molecules were then pooled for downstream processing to generate a sequencing-ready library. The resulting 10× barcoded library was processed with standard NGS short-read sequencing on Illumina sequencers for massive transcriptional profiling of entire tissue sections.

### Bulk RNA sequencing

Brain tissues were extracted from mice that were anesthetized with isoflurane gas. Next, MEC tissue was isolated and homogenized in TRIzol, followed by phase separation with chloroform. RNA was precipitated using isopropanol, washed with ethanol, and air-dried before resuspension in DEPC-treated water. The quality and concentration of RNA were measured. All samples with RNA Integrity Number (RIN) > 8 were used as input to library construction.

### RT-qPCR

The procedure was the same as mentioned above. The quality and concentration of RNA were measured, followed by cDNA synthesis. qPCR was performed using SYBR Green to quantify target gene expression levels, with *GAPDH* as the reference gene.

Primer sequences:

*Bglap3* (F): CTGACAAAGCCTTCATGTCC, (R): TCAAGCTCACATAGCTCCC;

*GAPDH* (F): AGGTCGGTGTGAACGGATTTG, (R): TGTAGACCATGTAGTTGAGGTCA.

### RNAscope in situ hybridization

Brain tissues were collected from mice anesthetized with isoflurane gas. The tissue was then placed in a container embedded with optical cutting temperature (OCT) Compound (4583, Sakura, Tokyo, Japan), placed on dry ice for freezing, and transferred to a cryostat pre-cooled to-20 °C for 1 h. Once the tissue temperature equilibrated, brain slices of 20 μm thickness were cut in a cryostat chamber with temperature adjusted to-10 °C following standard cryosectioning procedures, and mounted onto glass slides. The slides were placed at-20 °C for 20 min and then subjected to the following steps: fixation in 10% neutral formalin solution at 4 °C, dehydration, hydrophobic circle drawing, and treatment with hydrogen peroxide and proteinase IV. The slices were then processed using RNAscope^®^ Multiplex Fluorescent Reagent Kit V2 (323100, Advanced Cell Diagnostics, Inc., Newark, CA, USA). according to manufacturer’s instructions, which includes probe hybridization; AMP1, AMP2, and AMP3 incubation; and Opal 570 and Opal 520 fluorescence labeling. Finally, the slices were then DAPI-stained. A cover slip with a mounting medium containing anti-fading agent was placed on top, and the slides were immediately examined or stored at 4 °C. Subsequent confocal imaging was conducted on the Nikon A1R confocal microscope with a 60× oil immersion objective lens. Image analysis was performed using HALO software (Indica Labs Inc., Albuquerque, NM, USA).

### Axonal labeling and retrograde viral tracing

For axonal labeling, 0.4 μL of AAV2/9-hSyn-DIO-Synaptophysin-EGFP (2.98 × 10^12^ VG/mL; BrainVTA (Wuhan) Co., Ltd., Wuhan, China) was injected in the MEC (ML: +3.1-3.6; AP: sinus + 0.3; DV:-1.8-1.5,-1.5-1.3) with a Nanoliter 2010 Microinjection Pump (World Precision Instruments). For rabies tracing, 0.4 μL mixture of AAV2/8-DIO-EGFP-TVA and AAV2/8-DIO-oRVG (2.73 × 10^12^ VG/mL and 5.10 × 10^12^;BrainVTA (Wuhan) Co., Ltd., Wuhan) was injected in the MEC as the same location mentioned above. After three weeks, 0.8 μL of rabies virus EnvA-RVdG-mCherry (gift from Weixiang Guo Lab) was injected at the same location. Brain was collected one week after rabies virus injection.

### Immunohistochemistry

Mice were administered a 0.5 mL intraperitoneal injection of 2.5% tribromoethanol and securely fixed on an operating table once they showed no pain reflexes. The abdominal cavity was opened from below the sternum, and the chest was exposed by cutting the diaphragm. The right atrium was excised with scissors, followed by the insertion of a perfusion needle into the left ventricle. A total of 20 mL of physiological saline was perfused using a perfusion pump at a rate of 5 mL/min until the liver lost its natural color. Subsequently, 20 mL of 4% PFA was perfused at the same flow rate. The mouse’s head was then severed from the neck, the skull was carefully dissected using fine forceps, and the brain tissue was removed and immersed in PFA for 24 h for fixation. The brain tissue was subsequently dehydrated in a 30% sucrose solution and left to soak for >24 h until the brain settled to the bottom.

Brain tissue sections were prepared using a cryostat microtome. The fixed and dehydrated mouse brain was carefully removed from the surface sucrose solution and excess liquid was blotted. The brain was then secured onto a metal mounting platform embedded with OCT Compound (4583, Sakura, Tokyo, Japan). A layer of OCT was applied to the surface of the brain tissue, and the prepared tissue was placed in the cryostat for freezing. The frozen tissue was then sectioned at a thickness of 30 μm. Using a fine brush, the desired brain sections were flattened on the sample stage. A glass slide was brought close to the brain sections, and the temperature differential was used to adhere the brain sections onto the glass slide. The cryostat chamber temperature, as well as the temperatures of the blade and sample stage, were maintained at - 20 °C throughout the sectioning process.

Brain sections with fluorescence were dried under open air, rinsed with PBST (0.3% Triton-X100) for 10 min, stained with DAPI (1:1000) for 10 min, and sealed with a sealing medium. For immunofluorescence staining, brain slices were incubated in blocking solution (10% normal goat serum PBST, 1% Triton-X100) freely floating for two hours at room temperature, followed by incubation with primary antibodies NeuN (ab190565; 1:500) and GAD67 (ab75712; 1:100) in the blocking solution for three nights at 4 °C. After washing in PBST 10 mins for three times, the secondary antibody (A11039, Invitrogen; 1:500) was added and incubated for 2 hours at room temperature. Then, by washing for another 10 mins for three times, the brain tissues were mounted on the slices and the slices were sealed with a sealing medium. Imaging was processed using a Nikon A1R confocal microscope with a 20× objective lens. Fluorescence cells were counted manually.

### Prism implantation for 2-photon calcium imaging

A 0.8 μL mixture of AAV2/9-CaMKIIa-DIO-hM4D(Gi)-mCherry-WPRE-hGH or AAV2/9-CaMKIIa-DIO-mCherry-WPRE-hGH with AAV2/9-hSyn-GCaMP6s-WPRE-hGH (5.01 × 10^12^ VG/mL, 4.32 × 10^12^ VG/mL, 5.27 × 10^12^ VG/mL; BrainVTA (Wuhan) Co., Ltd., Wuhan, China) was injected in the MEC (ML: +3.1∼3.6; AP: sinus + 0.3; DV:-1.8∼1.5,-1.5∼1.3) with a Nanoliter 2010 Microinjection Pump (World Precision Instruments). Three weeks after injection, mice were anesthetized with isoflurane, and a small square cranial window (2.5 × 2.5 mm^2^) was made and centered over the targeted cortex. The dura was carefully removed, and a small glass prism (3.5 × 2.5 mm^2^) was placed along the dorsoventral surface of the MEC between the cerebrum and cerebellum. After recovery for one week, the headstage was mounted to the skull of each mouse. To increase the imaging stability, 1.5% low-melting-point agarose was added to the objective of the brain tissue. After adaptation, we restrained each mouse with a body holder and used the benchtop TPM to confirm the region of viral infection (ROI). We then mounted the fast, high-resolution, miniaturized TPM (FHIRM-TPM) to the ROI designated by the benchtop TPM. Next, we glued a headstage to the skull during craniotomy preparation and protected the attachment with dental cement. During imaging trials, mice were trained in an 80-cm square black aluminum enclosure with a single white cue card for spatial orientation. Calcium signals were recorded using a miniaturized two-photon microscope (TPM), following previously published protocols ^56^. For experiments with AAV-DIO-hM4D-mCherry injections, trial recordings lasted 40 min, whereas recordings for the AAV-DIO-mCherry control group lasted 30 min. Running behavior was encouraged by scattering cookie crumbs at random locations in the enclosure every 10–15 s. Mice were allowed to rest for 3–5 min between Baseline 1 and 2 sessions, as well as between Recovery 1 and 2 sessions. Following Baseline 2, mice received an intraperitoneal injection of Deschloroclozapine (DCZ, 0.1 μg/kg), and DCZ sessions commenced at least 10 min after the injection. A single Bglap3-Cre mouse (P08) received an MEC injection of AAV2/9-EF1α-DIO-GCaMP6s GCaMP6s (BrainVTA (Wuhan) Co., Ltd., Wuhan, China) for cell-type classification. This animal was excluded from DCZ treatment and the 5-session recording protocol.

### c-Fos staining and image processing

c-Fos immunostaining was performed under the same procedure mentioned previously. Primary antibody c-Fos (226008, Synaptic System; 1:1000) was incubated at 4 °C for one night, followed by secondary antibody (A11034, Invitrogen; 1:500) incubation for one hour at room temperature.

c-Fos+ cells were identified with ImageJ/Fiji plugin StarDist ^90^ using the Versatile (fluorescent nuclei) model. c-Fos+ cell whose Feret’s Diameter was between 8 and 16 µm was selected, and the fluorescence intensity of individual c-Fos+ cells was measured by their Mean Gray Value in arbitrary units. *Bglap3*-positive or negative cells were identified based on whether the Mean Gray Value of mCherry signals was higher or lower than 1000.

### Visual discrimination learning task

The task was done in an operation box that is 40 × 35 × 40 cm, with a touch screen placed at one of the interior walls. In the pre-learning stage, a random image appeared on the touch screen every 25 seconds. Once the mice touches the image, it will be rewarded with chocolate beans in the opposite food trough. This encouraged the mice to actively touch the image, establishing an association between touching the picture and receiving a reward. A chocolate reward for a mouse is counted as one trial. Each mouse does 30 trials or 30 minutes per day. In the learning stage, two distinct images appear on the touch screen at the same time. By touching the correct one the mouse receives a chocolate reward; otherwise, by touching the wrong picture, the mouse receives a short light penalty. Each mouse performs 30 trials or 60 minutes every day, and the test is continued for 1 to 2 months, with each period of 10 days considered as one learning stage. The success of each mouse was assessed on a daily basis, and the accuracy rate was calculated. If a mouse achieved an accuracy rate exceeding 70% for three consecutive days, it was deemed to have successfully learned the task.

## Data analysis

### Spike Waveform Analysis

#### Feature Extraction

For each spike, we extracted the raw waveform from four channels of the tetrode (ch5–ch8), with 50 sampling points per waveform. The energy of each waveform was computed as the sum of squared amplitudes across time. To normalize spike shape independent of amplitude, waveforms were divided by their energy, resulting in energy-normalized waveforms for each channel. Principal component analysis (PCA) was then applied separately to the normalized waveforms of each channel, and the first principal component (PC1) from each was retained. Thus, for each spike, an 8-dimensional feature vector was constructed: four energy values and four PC1 scores (one per channel).

#### Quality control by calculating Mahalanobis Distance and Cluster Metrics

Let *X* ∈ *R^N^*^×8^ denote the feature matrix of all *N* spikes recorded in a session. Given a cluster of interest (i.e., a candidate neuron) consisting of a subset of spikes indexed by *I*_cluster_, we computed the Mahalanobis distance between every spike and the multivariate distribution defined by *X_I_*_cluster,:_, the feature vectors of the cluster. This distance incorporates the full covariance structure of the cluster, providing a robust measure of proximity in feature space.

The following three metrics were then derived:

Isolation Distance: Defined as the squared Mahalanobis distance to the N-th closest non-cluster spike, where N is the number of spikes in the cluster. It represents the radius of a hypersphere around the cluster such that adding more spikes would introduce contamination.

L-ratio: A measure of cluster contamination. For each spike outside the cluster, the squared Mahalanobis distance to the cluster centroid is converted into a p-value under a chi-squared distribution with degrees of freedom equal to the feature dimension. The L-ratio is calculated as the sum of these p-values (i.e., likelihood of belonging to the cluster) normalized by the number of spikes in the cluster. Mean Mahalanobis Distance: As a general indicator of cluster compactness, we also computed the average Mahalanobis distance among spikes within the cluster to their own centroid. COM shift (Center of mass shift): Cluster stability across consecutive sessions was estimated by comparing the shift of the center of mass point for corresponding clusters in two sessions. Specifically, the length of the vector between the two COMs in an 8-dimensional space was calculated and normalized to the length of the vector from the origin (0,0) to the COM point in the first session.

To ensure numerical stability, we excluded spikes with zero-valued waveforms on any channel (which result in undefined normalized waveforms and PCA failure), and standardized all feature vectors (z-scoring) before distance calculations.

### Rate maps, firing fields, and spatial information

Position estimates were based on tracking of LEDs on the headstage connected to the microdrive. Tracked positions were smoothed with a 15 point mean filter offline. To characterize firing fields, the position data were sorted into bins of 2.5 × 2.5 cm^2^ and the firing rate was determined for each bin. A spatial smoothing algorithm was used ^91^. The average rate in any bin *x* was estimated as:

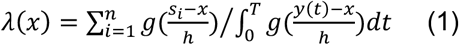

Where *g* is a smoothing kernel, *h* is a smoothing factor, *n* is the number of spikes, *si* the location of the *i*-th spike, *y(t)* is the location of the mice at time *t*, and [*0*, *T*] is the recording period. A Gaussian kernel was used with *g* and *h* = 3. In order to avoid error from extrapolation, we considered positions more than 3 cm away from the tracked path as unvisited.

The cell’s peak rate was estimated as the highest firing rate observed in any bin of the smoothed rate map. For each cell, the spatial information content in bits per spike ^92^was calculated as:

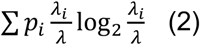

where *λ*i is the mean firing rate of a unit in the *i-*th bin, *λ* is the overall mean firing rate, and *pi* is the probability of the animal being in the *i-*th bin (occupancy in the *i-*th bin / total recording time).

### Analysis of grid cells

Our methods for grid cell analysis were adapted from those described in previous studies ^93,94^. For all cells with more than 100 spikes, we calculated the spatial autocorrelation for each smoothed rate map. Autocorrelograms were based on Pearson’s product moment correlation coefficient with corrections for edge effects and unvisited locations. With *λ (x, y)* denoting the average rate of a cell at location *(x, y)*, the autocorrelation between the fields with spatial lags of *τ_x_* and *τ_y_* was estimated as:

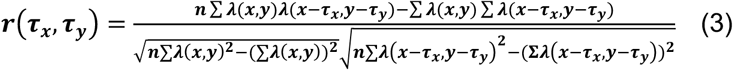

where the summation is over all *n* pixels in *λ (x, y)* for which the rate was estimated for both *λ (x, y)* and *λ (x – τx, y – τy)*. Autocorrelations were not estimated for lags of *τx, τy* where *n* < 20.

The degree of spatial periodicity (‘grid score’) was determined for each recorded cell by taking a circular sample of the autocorrelogram, centered on but excluding the central peak, and comparing rotated versions of this sample. The Pearson correlation of this circle with its rotation in α degrees was obtained for angles of 60° and 120° on one side and 30°, 90°, and 150° on the other. The cell’s grid score was defined as the minimum difference between any of the elements in the first group and any of the elements in the second. Shuffling was performed for each cell individually with 1000 permutations for each cell. In each permutation, the entire sequence of the spikes or the cell’s deconvolved calcium activity was time-shifted along the animal’s path by a random interval between 30 s on one side and 30 s on the other side less than the length of the session, with the end of the session wrapped to the beginning. The grid score of each data shuffle was calculated according to previously described methods ^94^. If the grid score from the recorded data was larger than the 99^th^ percentile of grid scores in the distribution from the 1000 permutations of shuffled data of the same cell, the cell was defined as a grid cell. In calcium imaging analysis, stable grid cells were defined as those with a spatial correlation greater than 0.3 between the two baseline sessions, and a grid score exceeding the 95th percentile of the shuffle distribution in at least one baseline session.

### Analysis of HD cells

Our methods for analyzing HD cells have been reported in previous studies ^94,95^. The mouse’s HD was calculated for each tracker sample from the projection of the relative position of the two LEDs onto the horizontal plane. The directional-tuning function for each cell was obtained by plotting firing rate as a function of the mouse’s directional heading, divided into bins of 3° and smoothed with a 30° mean window filter (five bins on each side). The preferred firing direction was defined by the mean vector of the directional-tuning function.

HD-modulated cells were defined as cells with mean vector lengths significantly exceeding the degree of directional tuning that would be expected by chance. Chance values were determined by a shuffling procedure performed in the same way as for grid cells and using a similar number of permutations per cell. Cells were defined as a HD cell (HD) if the mean vector from the recorded data was larger than the 99^th^ percentile of the mean vector in the distribution from the 1000 permutations of shuffled data of the same cell. In calcium imaging analysis, stable HD cells were defined as those with an HD tuning correlation greater than 0.3 between the two baseline sessions, a difference in preferred angles of no more than 30 degrees, and a mean vector length exceeding the 95th percentile of the shuffle distribution in at least one baseline session, while not having a grid score surpassing that of the shuffle distribution.

### Analysis of border cells

Our methods for analyzing border cells are based on previous reports ^18,63^. Border cells were identified by the score resulting from the following computation: For each cell with an average rate above 0.2 Hz, the border score was the difference between the maximal length of a wall touching on any single firing field of the cell and the average distance of the field from the nearest wall, divided by the sum of those values. Border scores thus ranged from-1 for cells with infinitely small central fields to +1 for cells with infinitely narrow fields that lined up perfectly along the entire wall.

Border cells were defined as cells with border scores significantly exceeding chance levels determined by a shuffling procedure performed in the same way as for grid cells, using a similar number of permutations per cell, but with the addition of a spatial information criterion ^63^. Cells were defined as border cells if two conditions were met: (i) The border score from the recorded data was larger than the 99^th^ percentile of border scores in the distribution from the 1000 permutations of shuffled data of the same cell, and (ii) the spatial information content in the recorded data was higher than the 99^th^ percentile value for spatial information in the shuffled data. In calcium imaging analysis, stable border cells were defined as those with a spatial correlation greater than 0.3 between the two baseline sessions, and a border score exceeding the 95th percentile of the shuffle distribution in at least one baseline session.

Spatial cells were defined as those with a spatial correlation greater than 0.3 between the two baseline sessions, and spatial information content in both baseline sessions exceeding the 95th percentile of the shuffle distribution, excluding grid, border and HD cells.

Spatial tuning cells (SPT) were defined as those with a spatial correlation greater than 0.3 between the two baseline sessions, and spatial information content in both baseline sessions exceeded the 95th percentile of the shuffle distribution. It may intersect with grid, border, or HD cells.

### Analysis of speed cells

Our methods for analyzing speed cells are similar to those reported in a previous study ^19^. The instantaneous firing rate was obtained by dividing the whole trial into 20 ms bins, coinciding with the frames of the tracking camera. A temporal histogram of spiking was then smoothed with a 250 ms-wide Gaussian filter. The speed score for each cell was defined as the Pearson product-moment correlation between the cell’s instantaneous firing rate and the mouse’s instantaneous running speed, on a scale from-1 to 1. A cell was defined as a speed cell if the speed score from the recorded data was larger than the 99^th^ percentile of speed scores in the distribution from the 1000 permutations of shuffled data of the same cell. Shuffling was performed in the same way as for grid cells and HD cells.

### LFP power spectrum analysis

EEG Power Spectrum and Band-Power Quantification Signal extraction and preprocessing: For each recording session, EEG time series were read. The sampling frequency was fixed at 250 Hz for all analyses. No additional filtering, rereferencing, or artifact rejection was applied prior to spectral estimation.

Power spectral density (PSD) estimation (multitaper): PSD was estimated with the multitaper method implemented in MNE-Python (psd_array_multitaper, v1.10.1), which uses discrete prolate spheroidal (DPSS/Slepian) tapers and Thomson’s adaptive weighting. Parameters were: analysis band: 0.1–30 Hz (fmin=0.1, fmax=30); bandwidth=1.0 (DPSS half-bandwidth in Hz, controlling spectral smoothing); adaptive weighting: on (adaptive=True); normalization:’full’ (PSD in power-per-Hz in the signal’s native units); other options at MNE defaults (including removal of DC offset).psd_array_multitaper returns the frequency vector *f* and the one-sided PSD *S_xx_*(*f*). Band-limited power:Two physiologically relevant bands were quantified: theta: 6–12 Hz, gamma: 20–30 Hz.

For a band [*f_min_*, *f_max_*], band power was defined as the arithmetic mean of the PSD samples whose frequencies fall within the band:

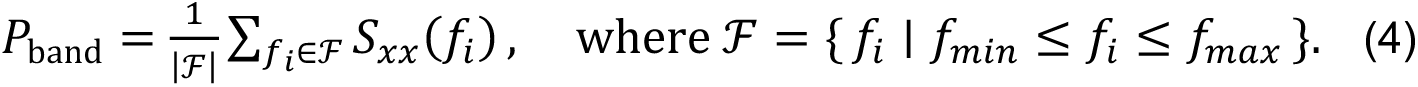

“theta power” and “gamma power” correspond to the mean PSD across 6–12 Hz and 20–30 Hz, respectively. (Band-power averaging is performed directly on the native PSD frequency grid returned by MNE; no integration or additional weighting is applied.)

### 10x Genomics Visium data analysis

Raw Illumina sequencing data was processed with Space Ranger (v2.1) for sample demultiplexing, image alignment, barcode processing, and gene counting. The output file could be displayed Loupe Browser (v7) software to visualize gene expression patterns across brain slices. MEC and its subregions were assigned based on Paxinos & Franklin’s *The Mouse Brain in Stereotaxic Coordinates, Third Edition* ^96^ and the spatial barcodes were extracted for further analysis. Feature-barcode matrices from different mice were merged in R using Seurat package ^97^ following Harmony for batch correction ^98^, and UMAP visualization of dimensional reduction and cluster displaying. After extracting the data based on barcodes from entire MEC or sublayers of MEC, normalization and differential expression of MEC or sublayers between old and young mice was done with DEseq2 in R ^99^, with a threshold that was set to absolute Log2 Fold-Change > 0.5, adjusted P-value < 0.05. Enrichment network analysis of differentially expressed genes was done with Metascape ^100^. Differential expressions of genes presented in Supplementary Fig.13 were calculated in Loupe Browser 7 using algorithms sSeq ^101^and edgeR ^102^.

### 10x Genomics Visium image analysis

Boundaries of the MEC with neighboring regions, such as the postrhinal cortex, parasubiculum, presubiculum, lateral entorhinal cortex, and subiculum, were defined based on Franklin’s *The Mouse Brain in Stereotaxic Coordinates, Third Edition* ^96^. Unfolded MEC maps were prepared for each sagittal brain section by mapping the dorsal border of the MEC onto a straight line. For each section, the surface length of the MEC was measured using Image-Pro Plus software (Media Cybernetics, Rockville, MD, USA) and subsequently mapped onto a straight line perpendicular to the line that represents the dorsal border. All borders were established using cytoarchitectonic criteria that can reliably be established irrespective of the plane of sectioning, as described in detail for the rat brain ^103^. These borders, as defined in the rat, can be reliably applied to the mouse brain ^104^.

### Bulk RNA sequencing data analysis

RNA raw data was first analyzed by Fastqc (http://www.bioinformatics.babraham.ac.uk/projects/fastqc) for quality control, then we used Trimmomatic (v0.39) ^105^ to filter low-quality reads and trim the linker sequence. Reads were aligned to the GRCm39 reference genome using STAR (2.7.5c) ^106^. After comparing with the reference genome, featureCounts was conducted for gene quantification to get the raw counts matrix for downstream analysis. The differential expression analysis was done by DESeq2 ^99^ in R setting the threshold as: absolute Log2 Fold-Change > 0.5, adjusted P-value < 0.05.

### Analysis of scRNA-seq data

Data from Yao et al. (2023)^54^ were downloaded and analyzed in Seurat V4 ^97^. Cells labeled with supertypes from the ‘ENT’ region were extracted for further analysis. SCTransform was used for data normalization and scaling, followed by PCA and UMAP to visualize the distribution of subclasses or supertypes. Default parameters were applied for all analyses. To assess the fraction of *Bglap3*+ cells, we defined cells with more than one read count of *Bglap3* as *Bglap3*+ cells.

Deconvolution was performed on our 10x Visium spatial transcriptomic data using Cell2location ^55^, with a cell type reference derived from scRNA-seq data from Yao et al., applying default parameters (N_cells_per_location set to 5, and detection_alpha set to 20). Read count matrices from single-cell and spatial transcriptomes, along with cell type information on subclasses, were imported into Scanpy (v1.9.8) ^107^ in Python (v3.9). The resulting cell-type compositions were normalized and subsequently transformed using a centered log-ratio (CLR) method, as implemented in the [“compositions” R package for analyzing compositional data]. These transformed data were then used to analyze cell type differences between young and old mice. Average fractions of subclasses across each slide are calculated, followed by the comparison of the average fractions between young and old with two-sided Wilcoxon tests.

### Calcium imaging data analysis

Calcium imaging data were preprocessed using Suite2p ^108^(https://github.com/MouseLand/suite2p), a popular open-source pipeline for calcium imaging data analysis. This involved motion correction to counteract minor drifts in the imaging plane over the session, automated region of interest (ROI) extraction to identify individual neuronal cells based on their spatial footprints and temporal activity patterns, and extracting fluorescence signals (*F_ra_*_w_) from the ROIs. We manually curated and removed ROIs with poor morphological characteristics or subpar fluorescence signals. Custom MATLAB (R2020b) scripts were used for further analysis. For each detected neuron, the baseline fluorescence (*F*_0_) was gauged from the lower 8th percentile of the trace, with the ΔF/F signal calculated as (*F_ra_*_w_ − *F*_0_)/*F*_0_correspondingly. The ΔF/F signal underwent wavelet denoising using the ‘wdenoise’ function to eliminate noise while preserving salient features of the signal. Subsequently, we employed the OASIS ^109^(https://github.com/zhoupc/OASIS_matlab) algorithm for deconvolution to infer the underlying spiking activity from the denoised ΔF/F signal. To further refine our detection of significant calcium events, we defined a time window in the ΔF/F signal as a’significant term’ if its amplitude exceeded twice the local standard deviation. Only the deconvolved calcium events falling within this designated significant term were retained for subsequent analyses. Neuronal activity was correlated with the animal’s position and movement in the arena to identify spatially-modulated cells.

### Cell cross-session alignment

To effectively distinguish between *Bglap3* positive and *Bglap3*-negative neurons in MEC, we employed custom MATLAB scripts to align the scanned red channel field of view with the green channel field of view. In image processing, 2D cross-correlation serves as a robust method to compute similarity between two images. We computed cross-correlations between maximum intensity projection images outputted by Suite2p from different channels and sessions. The xy displacements corresponding to the maximum cross-correlation between two fields of view were employed as translation offsets for alignment corrections. If further alignment was required, the translationally-aligned image stacks were subjected to secondary alignment using MultiStackReg ^110^. MultiStackReg is an open-source ImageJ batch alignment plugin developed on top of Turboreg, which offers five versatile transformation registration modalities and is widely applied in image registration. From this process, we obtained the transformation matrices. Each session’s cellular footprints were then subjected to transformations. Using the transformed and aligned cell footprints as input to CellReg ^111^(https://github.com/zivlab/CellReg), a published neuron cross-day alignment tool, we identified cells that consistently appeared across imaging days. These cells were then classified into *Bglap3* positive or *Bglap3* negative categories based on their presence in the red channel.

### Populational dynamic of *Bglap3*-Grid cells

To compare the grid cell attractor network topology of *Bglap3-*neurons before and after silencing *Bglap3+* neurons. We employed the persistent cohomology framework described in Gardner et al. (Nature, 2022)^112^ to characterize the high-dimensional topological features of the grid cell attractor network. During our analysis, we constructed a population vector for grid cells. Unlike Gardner et al., which used instantaneous firing rates, we utilized denoised ΔF/F signals from calcium imaging to capture relative changes in neuronal activity. Data preprocessing followed the methodology outlined in Gardner et al. To ensure an adequate number of grid cells for analysis, we selected data from mouse #7934, which allowed the simultaneous recording of up to 41 grid cells meeting the inclusion criteria.

### Statistics & Reproducibility

Randomization was applied to the histology experiments: *Bglap3-Cre* mice of the same age were randomly assigned to receive DCZ or saline injection. Randomization was not applicable to the young vs. old comparisons, as age is a predetermined animal characteristic rather than an experimentally assigned variable.

All experiments were independently repeated at least three times, with similar results obtained each time. No data that met the quality-control criteria described above were excluded from analysis. Investigators were not blinded to group allocation during data collection or outcome assessment, except for histology analysis, which was performed blind to group allocation. Sex was not considered as a biological variable in the analyses.

All data were acquired and analyzed using GraphPad Prism v10 and MATLAB 2020b. Data are presented as individual values with mean ± standard error of the mean (SEM), unless otherwise stated. For data visualized with box and whisker plots, values are represented as median ± minimum to maximum range, 1.5 × IQR, or 10th - 90th percentile range.

All statistical analyses involving multi-subject repeated measurements were performed using linear mixed-effects models. Unless otherwise specified, the fixed effect was *age* (young vs. old) and the random intercept was mouse_id for electrophysiological and histological data. For two-photon data involving comparisons across more than two conditions (Baseline vs. DCZ vs. Recovery), an ANOVA with a linear mixed-effects model and post-hoc contrasts was used. Fixed-effect inferences were derived from the ANOVA table of the model. Pairwise post-hoc comparisons between condition levels were conducted as linear contrasts of fixed effects using MATLAB’s coefTest function, with F-tests and the [Satterthwaite] approximation for denominator degrees of freedom.

For data that did not involve repeated measurements within subjects, comparisons between two groups were conducted using two-sided t-tests when data were normally distributed. If the assumption of normality was not met, a two-sided Mann–Whitney U test (non-parametric) was applied. For comparisons involving more than two conditions within a single subject, the Friedman test (for repeated or matched values) was used, followed by Dunn’s multiple comparisons test for post hoc pairwise analyses, unless stated otherwise. Any specific statistical approaches applied in particular analyses are described in the respective Results sections.

## Data availability

The 10x Visium spatial transcriptomic data generated in this study are available in the Gene Expression Omnibus (GEO) under accession number GSE347650. The electrophysiological recording data, bulk RNA-seq data, and processed two-photon calcium imaging data have been deposited in the Zenodo database under accession code https://doi.org/10.5281/zenodo.21914598. The DEG list generated in this study is provided in the Supplementary Information file. Data shown in the figures are provided as a Source Data file.

## Code availability

All open-source code and software utilized in this study are listed in the Methods section. Custom lab code for 10x Visium differential analysis can be accessed on GitHub at: https://github.com/SushangLi/DEGs_for10xTrans/ ^113^.

## Acknowledgments

We thank Jialing Zhao and Julien Rousseau for proofreading the manuscript. We thank the Core Facilities of the School of Life Sciences, the State Key Laboratory of Membrane Biology at Peking University for assistance with confocal microscopy and Miniaturized Two-Photon Microscope.

## Author contributions Statement

Q.C. and C.M. planned and designed the initial experiments, with later input from G.S. Q.C. collected, and C.M. and G.S. analyzed the tetrode recording data. G.S., S.C., and J.L. performed 10X Genomics Visium experiments and analyzed the data. Y.L., S.W., J.H., X.Y., and M.Z. performed surgeries and collected the calcium imaging data. Y.L. analyzed the calcium imaging data. G.S. performed and analyzed histology and viral tracing data. Y.Y. and S.Y. provided code for behavior analysis. W.G. provided the rabies for tracing. S.X. and H.J. assisted with the revisions. C.M. supervised the project and provided funding. G.S. and Y.L. made the figures. G.S. and C.M. wrote the manuscript with input from all authors.

