## Supplemental Information File for "Aging-Related Cellular and Molecular Changes Reveal a Role for *Bglap3+* Neurons in MEC Spatial Cognition"

### Supplementary Figures

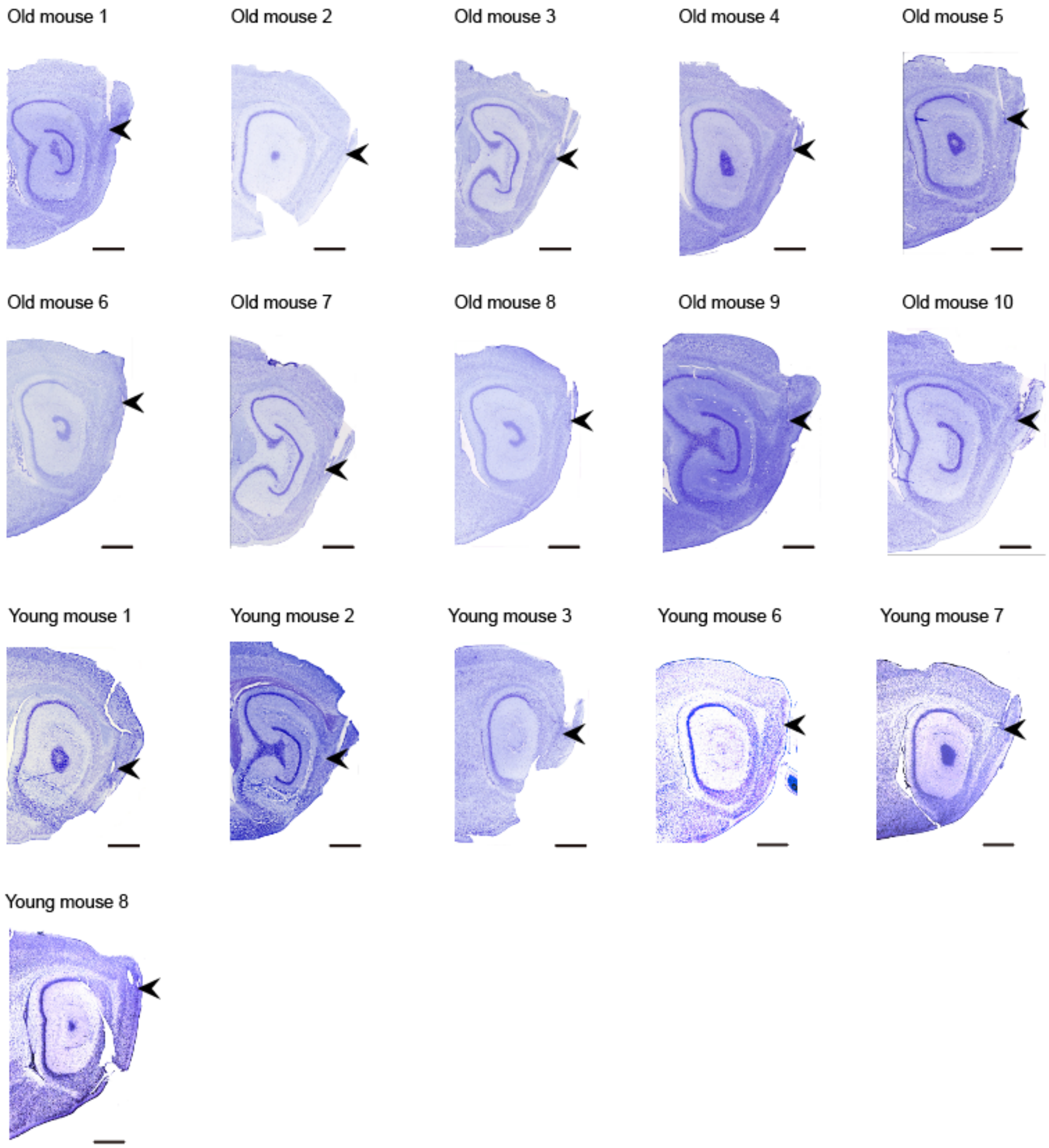

**Supplementary Fig.1 Sagittal brain sections showing locations of tetrodes in the MEC. Related to Fig.1.**

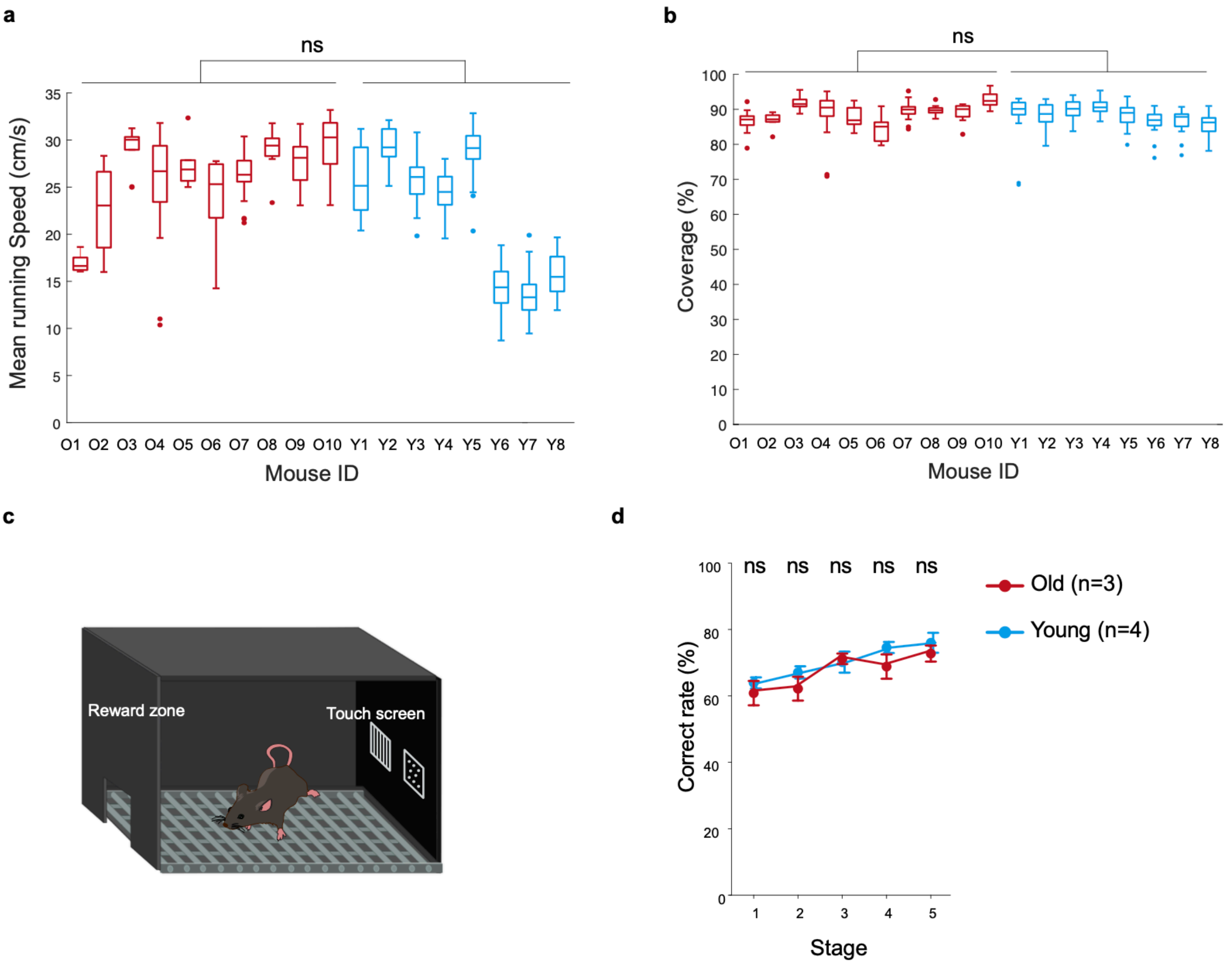

**Supplementary Fig.2 Behavioral performance of young and old mice.**

**a** Mean running speed during electrophysiology recordings of young (Y1-Y8) and old mice (O1-O10) (young n = 211 sessions vs. old n = 133 sessions; young: 23.44 ± 0.42 cm/s vs. old: 26.25 ± 0.41 cm/s; Linear mixed-effect model, SE = 2.34, t (342) = -1.51, P = 0.1328). **b** Trajectory coverage of open field during electrophysiology recordings of young (Y1-Y8) and old mice (O1-O10) (young n = 211 sessions vs. old n = 133 sessions; young: 88.57 ± 0.26% vs. old: 89.41 ± 0.35%; Linear mixed-effect model, SE = 0.95, t (342) = -0.79, P = 0.4289). **c** Schematic of visual cue discrimination learning task. **d** Percentage of correct trials across five stages of visual discrimination learning (10 days per stage). (young: N = 4 vs. old: N = 3. Two-way ANOVA followed by Sidak multiple comparisons: Stage 1, P = 0.9710; Stage 2, P = 0.8420; Stage 3, P = 0.9995; Stage 4, P = 0.7730; Stage 5, P = 0.9460). Data in **a** and **b** are median (center line), 25th–75th percentiles (box bounds), and 10th–90th percentiles (whiskers); data in **d** are presented as mean ± SEM; P-values for fixed effects in the linear mixed-effects models were derived from two-sided t-tests. ns, not significant. Source data are provided as a Source Data file. Related to Fig.1.

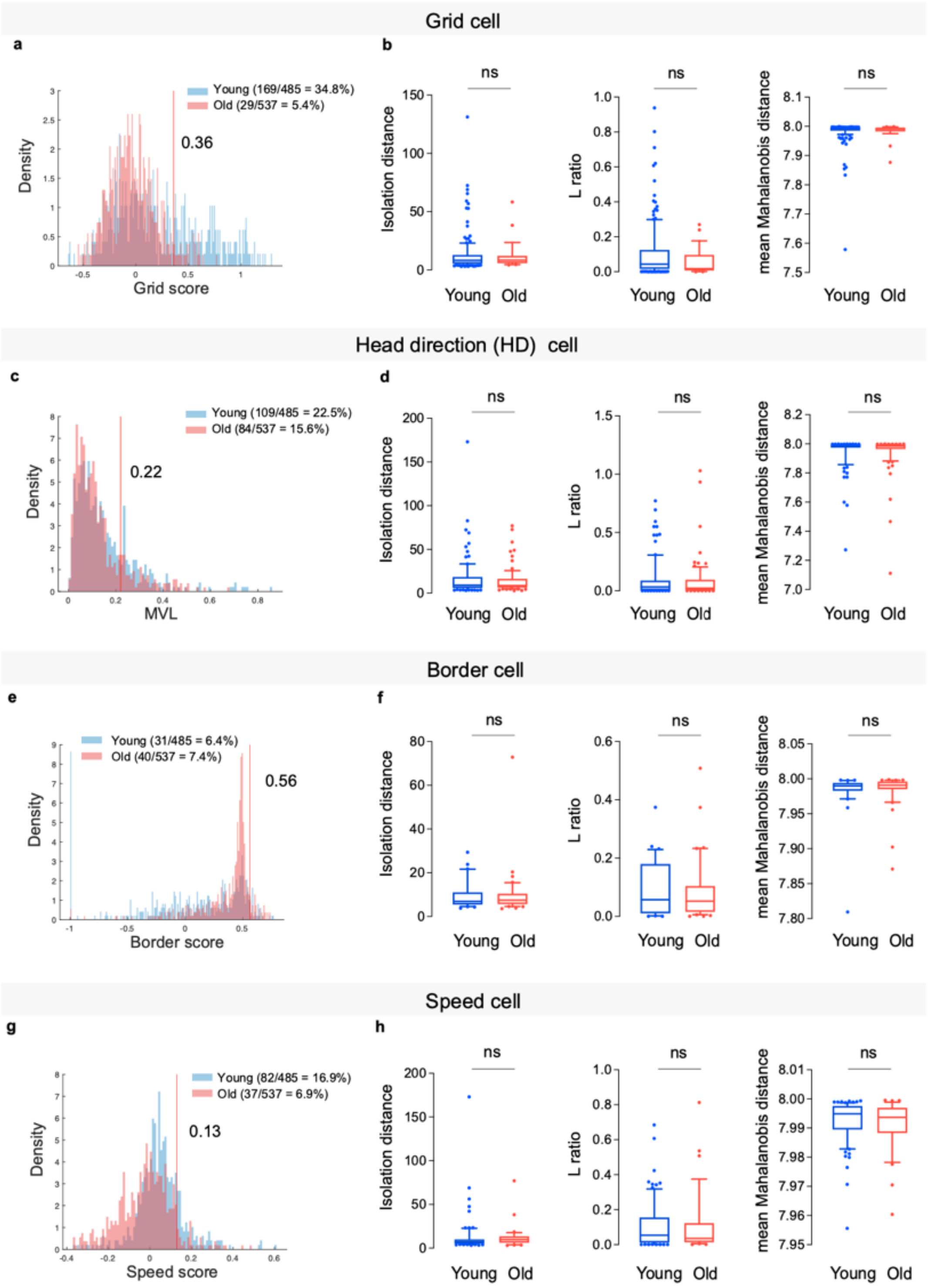
Grid cell
a
b
Young (169/485 = 34.8%)
Old (29/537 = 5.4%)
0.36
Density
Grid score
ns
Isolation distance
L ratio
mean Mahalanobis distance
Young
Old
Head direction (HD) cell
c
d
Young (109/485 = 22.5%)
Old (84/537 = 15.6%)
0.22
MVL
Border cell
e
f
Young (31/485 = 6.4%)
Old (40/537 = 7.4%)
0.56
Border score
Speed cell
g
h
Young (82/485 = 16.9%)
Old (37/537 = 6.9%)
0.13
Speed score

**Supplementary Fig.3 Assignment and quality control of spatial coding cells in young and old mice.**

**a**, **c**, **e**, **g** Histogram density plots of grid, MVL, border, and speed scores, respectively, for all recorded cells in young and old mice; red lines indicate thresholds (99th percentile of the corresponding shuffled distribution; see Methods).

**b**, **d**, **f**, **h** Quality control of grid, HD, border, and speed cells, respectively, assessed by isolation distance, L-ratio, and mean Mahalanobis distance (linear mixed-effects model, age as fixed effect). Values below are SE, t(df), P, in that order for each metric.

Grid cells (young n = 169, old n = 29): isolation distance, 2.95, t(196) = -0.26, P = 0.7915; L-ratio, 0.029, t(196) = -1.49, P = 0.1360; Mahalanobis distance, 0.0074, t(196) = 0.16, P = 0.8714.

HD cells (young n = 109, old n = 84): isolation distance, 2.67, t(191) = -0.87, P = 0.3881; L-ratio, 0.023, t(191) = -0.35, P = 0.7297; Mahalanobis distance, 0.016, t(191) = -0.28, P = 0.7835.

Border cells (young n = 31, old n = 40): isolation distance, 4.08, t(69) = 0.22, P = 0.8297; L-ratio, 0.024, t(69) = -0.34, P = 0.7343; Mahalanobis distance, 0.0068, t(69) = 0.34, P = 0.7386.

Speed cells (young n = 82, old n = 37): isolation distance, 5.16, t(117) = -0.52, P = 0.6007; L-ratio, 0.030, t(117) = 0.24, P = 0.8089; Mahalanobis distance, 0.0022, t(117) = -0.60, P = 0.5467.

Data are median (center line), 25th–75th percentiles (box bounds), and 10th–90th percentiles (whiskers). P-values for fixed effects in the linear mixed-effects models were derived from two-sided t-tests. ns, not significant. Source data are provided as a Source Data file. Related to Fig.1.

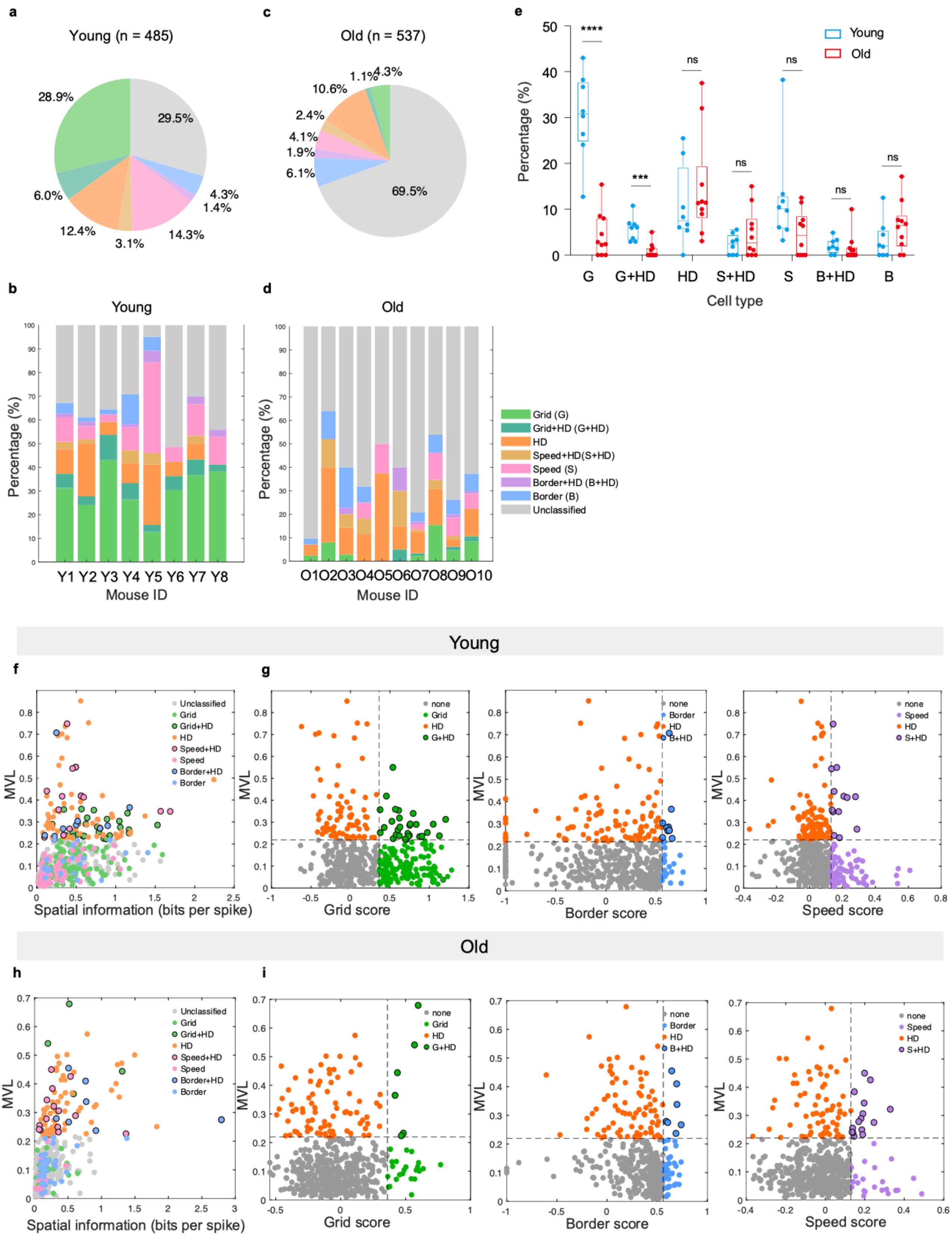
a
Young (n = 485)
28.9%
29.5%
4.3%
1.4%
14.3%
3.1%
12.4%
6.0%
c
Old (n = 537)
1.1%
4.3%
10.6%
2.4%
4.1%
1.9%
6.1%
69.5%
e
****
***
ns
Young
Old
Percentage (%)
G
G+HD
HD
S+HD
S
B+HD
B
Cell type
b
Young
Percentage (%)
Y1 Y2 Y3 Y4 Y5 Y6 Y7 Y8
Mouse ID
d
Old
Percentage (%)
O1O2O3O4O5O6O7O8O9O10
Mouse ID
Grid (G)
Grid+HD (G+HD)
HD
Speed+HD(S+HD)
Speed (S)
Border+HD (B+HD)
Border (B)
Unclassified
Young
f
MVL
Spatial information (bits per spike)
Unclassified
Grid
Grid+HD
HD
Speed+HD
Speed
Border+HD
Border
g
none
Grid
HD
G+HD
Grid score
none
Border
HD
B+HD
Border score
none
Speed
HD
S+HD
Speed score
Old
h
MVL
Spatial information (bits per spike)
i
Grid score
Border score
Speed score

**Supplementary Fig.4 Major cell types and conjunctive cell percentages in young and old mice.** **a** Pie chart showing the distribution of cell types in young mice: pure grid cells (G, without directional tuning, light green), pure border cells (B, without directional tuning, blue), pure speed cells (S, without directional tuning, pink), pure head direction (HD) cells (without spatial or speed tuning, orange), conjunctive grid + HD cells (G+HD, dark green), conjunctive speed + HD cells (S+HD, light brown), conjunctive border + HD cells (B+HD, purple), and unclassified cells (grey), with percentage values indicated. **b** Bar plot showing the percentages of each pure and conjunctive cell type for individual young mice. **c** Pie chart showing the same cell type distribution as in **a** for old mice, with percentage values. **d** Bar plot showing the percentages of each pure and conjunctive cell type for individual old mice. **e** Comparison of different cell types between young and old mice (Mann-Whitney test, G: $U = 79$, $P = 9.14 \times 10^{-5}$, G+HD: $U = 77$, $P = 3 \times 10^{-3}$, HD: $U = 28$, $P = 0.3154$, S+HD: $U = 31.5$, $P = 0.4628$, S: $U = 60$, $P = 0.0766$, B+HD: $U = 54$, $P = 0.2219$, B: $U = 24$, $P = 0.1679$). **f** Scatter plot of all cells from young mice showing spatial information versus MVL. **g** Scatter plots of cells from young mice showing grid versus directional tuning (left), border versus directional tuning (middle), and speed versus directional tuning (right). **h** Same as f but for old mice. **i** Same as g but for old mice. Data are presented as median with whiskers from min to max; **P<0.01; ***P<0.001; ****P<0.0001; ns, not significant. Source data are provided as a Source Data file. Related to Fig.1.

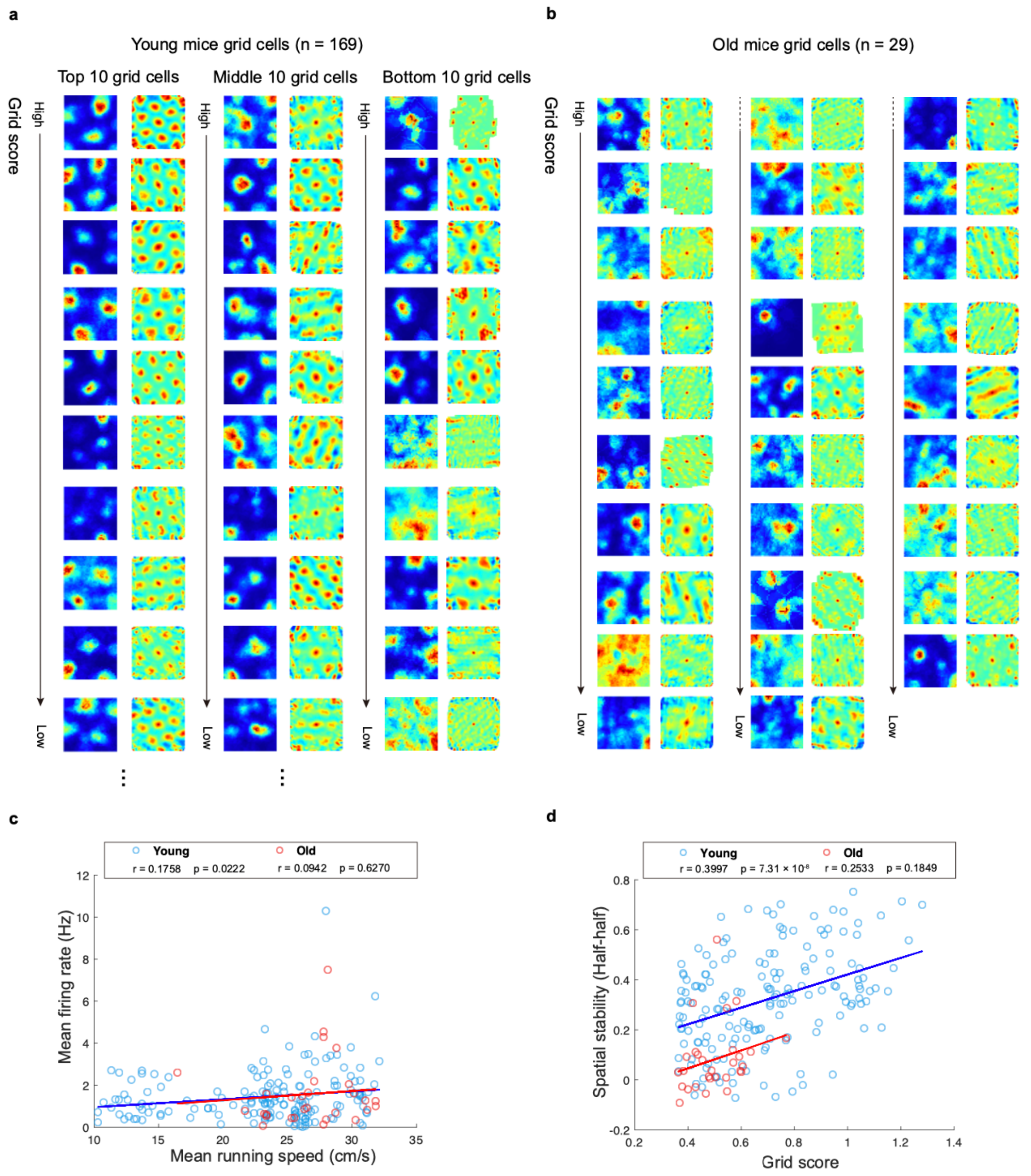

**Supplementary Fig.5 Grid cells in young and old mice.** **a** Color-coded rate maps (left) and autocorrelogram (right) of top, middle, and bottom 10 grid cells of young mice, ordered by their grid scores. **b** Color-coded rate maps (left) and autocorrelogram (right) of all grid cells recorded in old mice, ordered by their grid scores. **c** Correlation between mean firing rate of grid cells and running speed. Grid cells from young mice are shown as blue circles, and those from old mice as red circles; regression lines are shown in the corresponding colors. Pearson' s correlation coefficient (*r*) and *p*-values are indicated. **d** Correlation between spatial stability of grid cells and grid scores. Young grid cells are shown as blue circles and old grid cells as red circles, with regression lines in the corresponding colors. Pearson' s correlation coefficient (*r*) and *p*-values are indicated. Source data are provided as a Source Data file. Related to Fig.1.

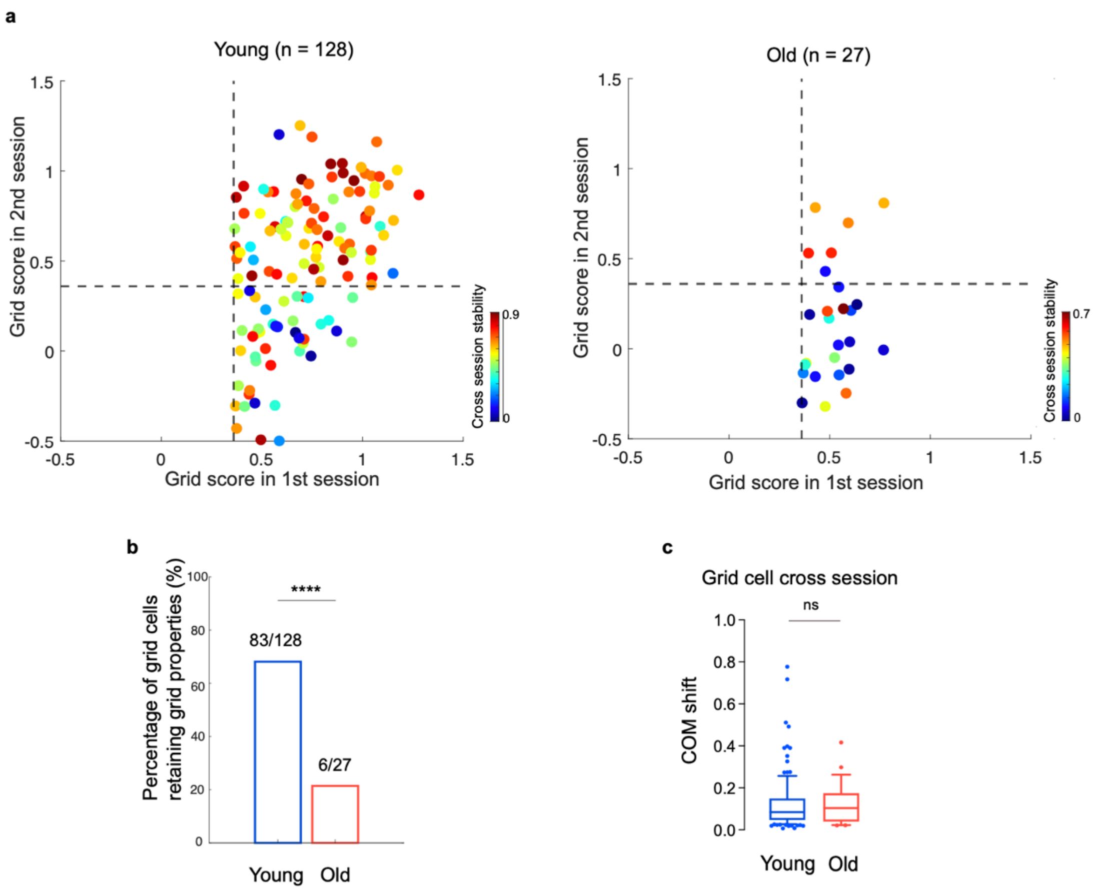

**Supplementary Fig.6 Grid cells recorded across two sessions. a** Scatter plots showing grid scores of grid cells in the first and second recording sessions in young (left) and old (right) mice. Each dot represents a cell, color-coded according to its cross-session correlation value. The dashed line indicates the threshold for grid score (0.36).**b** Bar plots showing the percentage of grid cells that retained grid properties across the two sessions (Fisher' s exact test, $P = 6.98 \times 10^{-5}$). **c** Cluster stability of grid cells across sessions, measured by center of mass shift, in young and old mice (young, n = 128; old, n = 27; Linear mixed-effects model with [age] as fixed effect, SE = 0.025, t (153) = -0.045, P = 0.9644). Data are median (center line), 25th–75th percentiles (box bounds), and 10th–90th percentiles (whiskers). P-values for fixed effects in the linear mixed-effects models were derived from two-sided t-tests. ****P<0.0001; ns, not significant. Source data are provided as a Source Data file. Related to Fig.1.

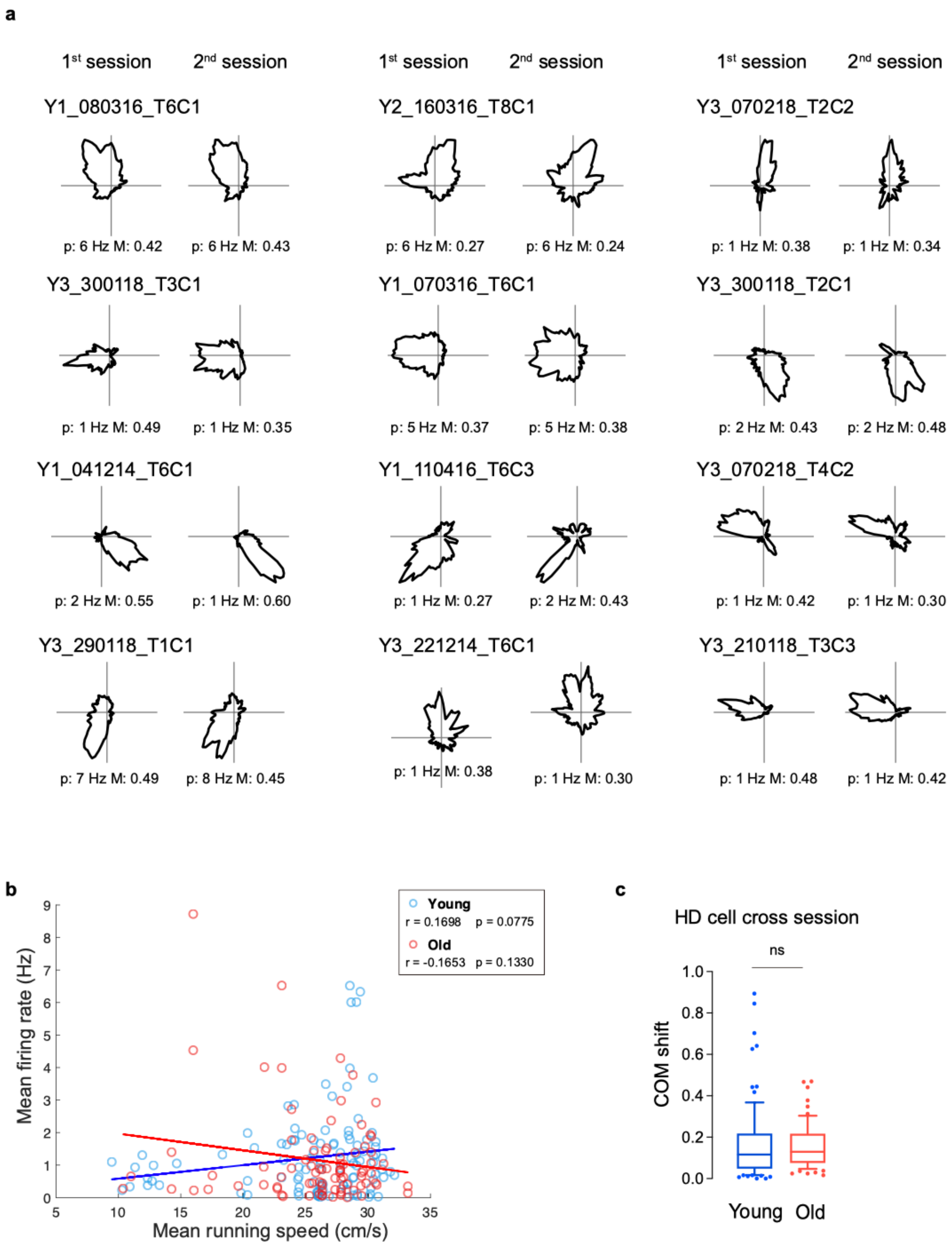
a
1st session
2nd session
Y1_080316_T6C1
p: 6 Hz M: 0.42
p: 6 Hz M: 0.43
Y2_160316_T8C1
p: 6 Hz M: 0.27
p: 6 Hz M: 0.24
Y3_070218_T2C2
p: 1 Hz M: 0.38
p: 1 Hz M: 0.34
Y3_300118_T3C1
p: 1 Hz M: 0.49
p: 1 Hz M: 0.35
Y1_070316_T6C1
p: 5 Hz M: 0.37
p: 5 Hz M: 0.38
Y3_300118_T2C1
p: 2 Hz M: 0.43
p: 2 Hz M: 0.48
Y1_041214_T6C1
p: 2 Hz M: 0.55
p: 1 Hz M: 0.60
Y1_110416_T6C3
p: 1 Hz M: 0.27
p: 2 Hz M: 0.43
Y3_070218_T4C2
p: 1 Hz M: 0.42
p: 1 Hz M: 0.30
Y3_290118_T1C1
p: 7 Hz M: 0.49
p: 8 Hz M: 0.45
Y3_221214_T6C1
p: 1 Hz M: 0.38
p: 1 Hz M: 0.30
Y3_210118_T3C3
p: 1 Hz M: 0.48
p: 1 Hz M: 0.42
b
Young
r = 0.1698 p = 0.0775
Old
r = -0.1653 p = 0.1330
Mean firing rate (Hz)
Mean running speed (cm/s)
c
HD cell cross session
ns
COM shift
Young
Old

**Supplementary Fig.7 Stable coding of head direction cells in young mice across sessions. a** Polar plots indicating firing rate as a function of head direction (right) across sessions for MEC Layers II and III in young mice. Peak rates (p) and mean vector length (M) are indicated under each rate map. Animal and cell numbers are indicated to the left of each triad of rate maps. **b** Correlation between mean firing rate of HD cells and running speed. HD cells from young mice are shown as blue circles, and those from old mice as red circles; regression lines are shown in the corresponding colors. Pearson's correlation coefficient (*r*) and *p*-values are indicated. **c** Cluster stability of HD cells across sessions, measured by center of mass shift, in young and old mice (young n = 86, old n = 65; Linear mixed-effects model with [age] as fixed effect, SE = 0.047, t (149) = -0.40, P = 0.6899). Data are median (center line), 25th–75th percentiles (box bounds), and 10th–90th percentiles (whiskers). P-values for fixed effects in the linear mixed-effects models were derived from two-sided t-tests. ns, non-significant. Source data are provided as a Source Data file. Related to Fig.2.

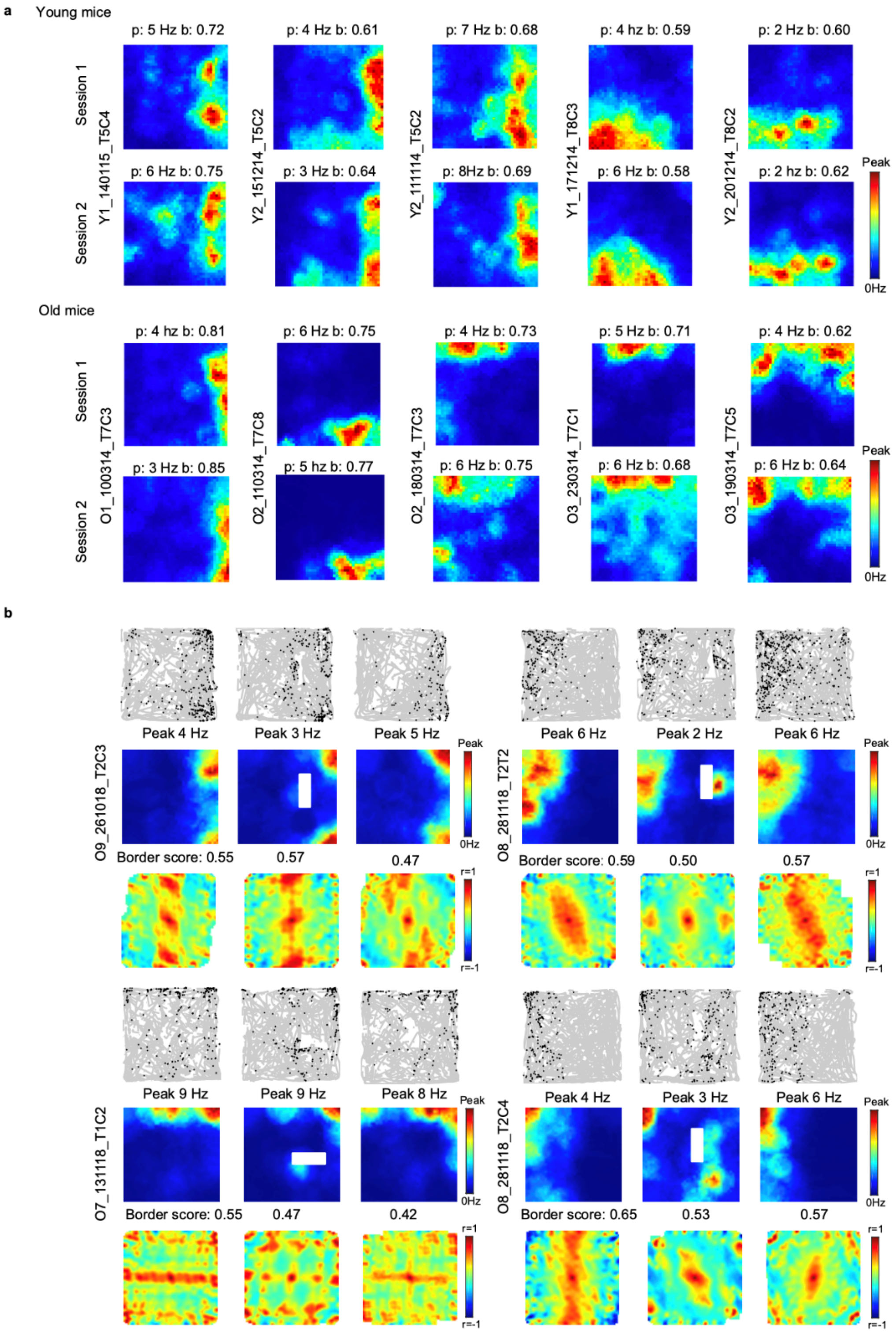

**Supplementary Fig.8 Stable coding of border cells across sessions in old mice. a** Color-coded rate maps across sessions from border cells in MEC Layers II and III in young and old mice. Peak rates and border scores are indicated above each rate map. Animal and cell numbers are indicated to the left of each triad of rate maps. **b** Introducing a standalone wall causes a new border field to appear in the MEC border cells of old mice. Related to Fig.2.

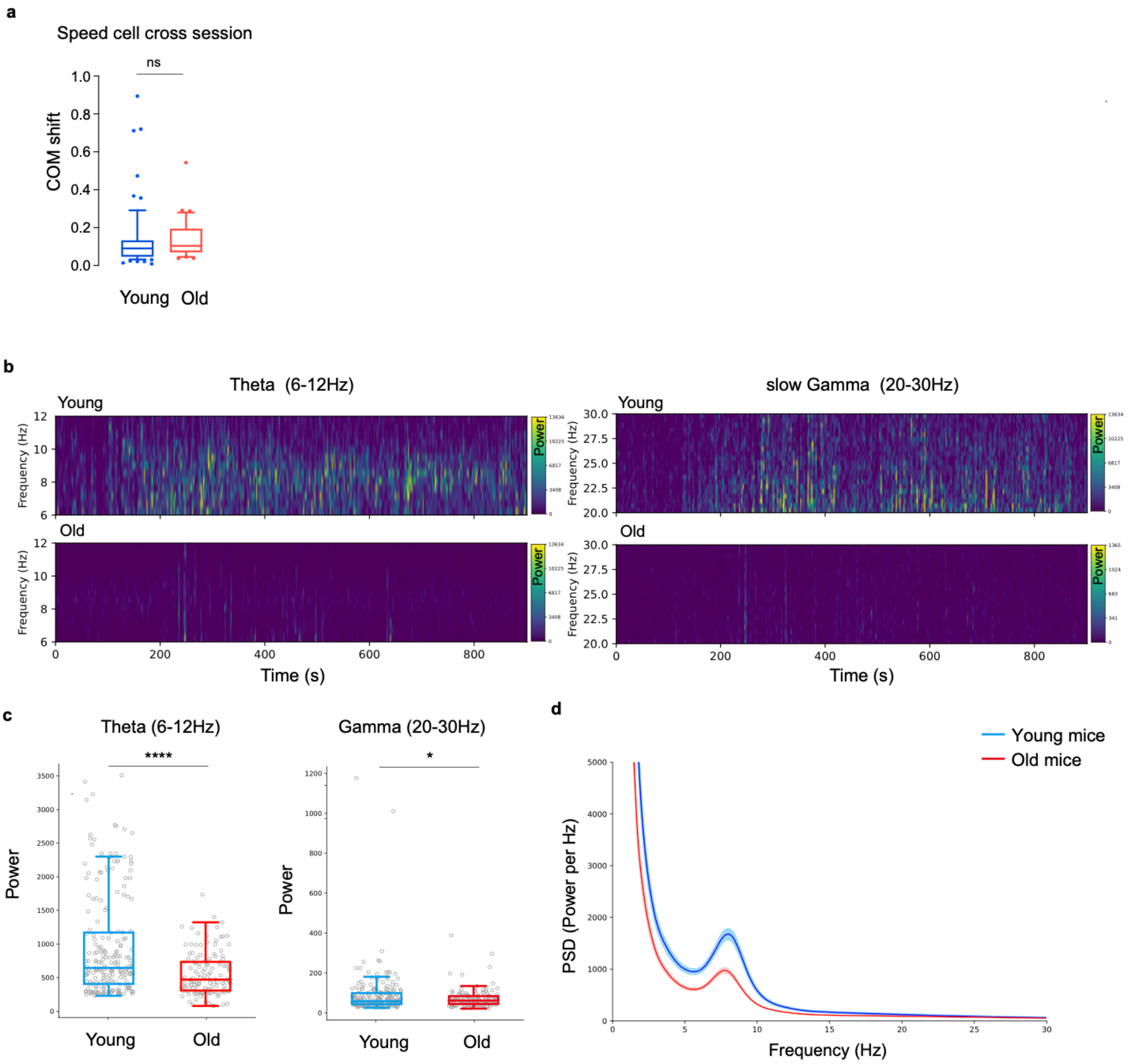

a
Speed cell cross session
ns
COM shift
1.0
0.8
0.6
0.4
0.2
0.0
Young
Old
b
Theta (6-12Hz)
slow Gamma (20-30Hz)
Young
Old
Frequency (Hz)
Power
Time (s)
c
Theta (6-12Hz)
Gamma (20-30Hz)
****
*
Power
Young
Old
d
Young mice
Old mice
PSD (Power per Hz)
Frequency (Hz)

**Supplementary Fig.9 Theta and Gamma power in young and old mice. a** Stability of clusters of speed cells across sessions, measured by center of mass (COM) shift, in young and old mice (young mice n = 67, old mice n = 31; Linear mixed-effects model with [age] as fixed effect, SE = 0.044, t (96) = 0.45, P = 0.6525). **b** Spectrograms of theta (6 -12 Hz) and slow gamma (20 – 30 Hz) in young (top) and old (bottom) mouse from one recording session. **c** Mean theta and gamma power in young and old mice (young mice n = 211 sessions, old mice n = 133, theta power, young: 956.95 ± 52.27, old: 554.05 ± 27.60; gamma power, young: 88.40 ± 7.67, old: 72.18 ± 4.14). Linear mixed-effects models with Age and Running speed as fixed effects and MouseID as a random intercept (Power ~ Age + Running Speed + (1|MouseID)). Theta power: F (1, 341) = 54.10, SE = 67.83, t (341) = 7.36, P = $1.43 \times 10^{-12}$: slow Gamma power: F (1, 341) = 5.00, SE = 10.36, t (341) = 2.24, P = 0.02595; Running speed was a significant covariate in both models (theta: SE = 5.74, t (341) = 5.96, P = $6.27 \times 10^{-9}$; slow gamma: SE = 0.88, t (341) = 2.83, P = $4.96 \times 10^{-3}$). **d** Power spectrum density plot showing power distribution across all frequencies. Data are median (center line), 25th–75th percentiles (box bounds), and 10th–90th percentiles (whiskers). P-values for fixed effects in the linear mixed-effects models were derived from two-sided t-tests. *P < 0.05; ****P < 0.0001; ns: non-significant. Source data are provided as a Source Data file.

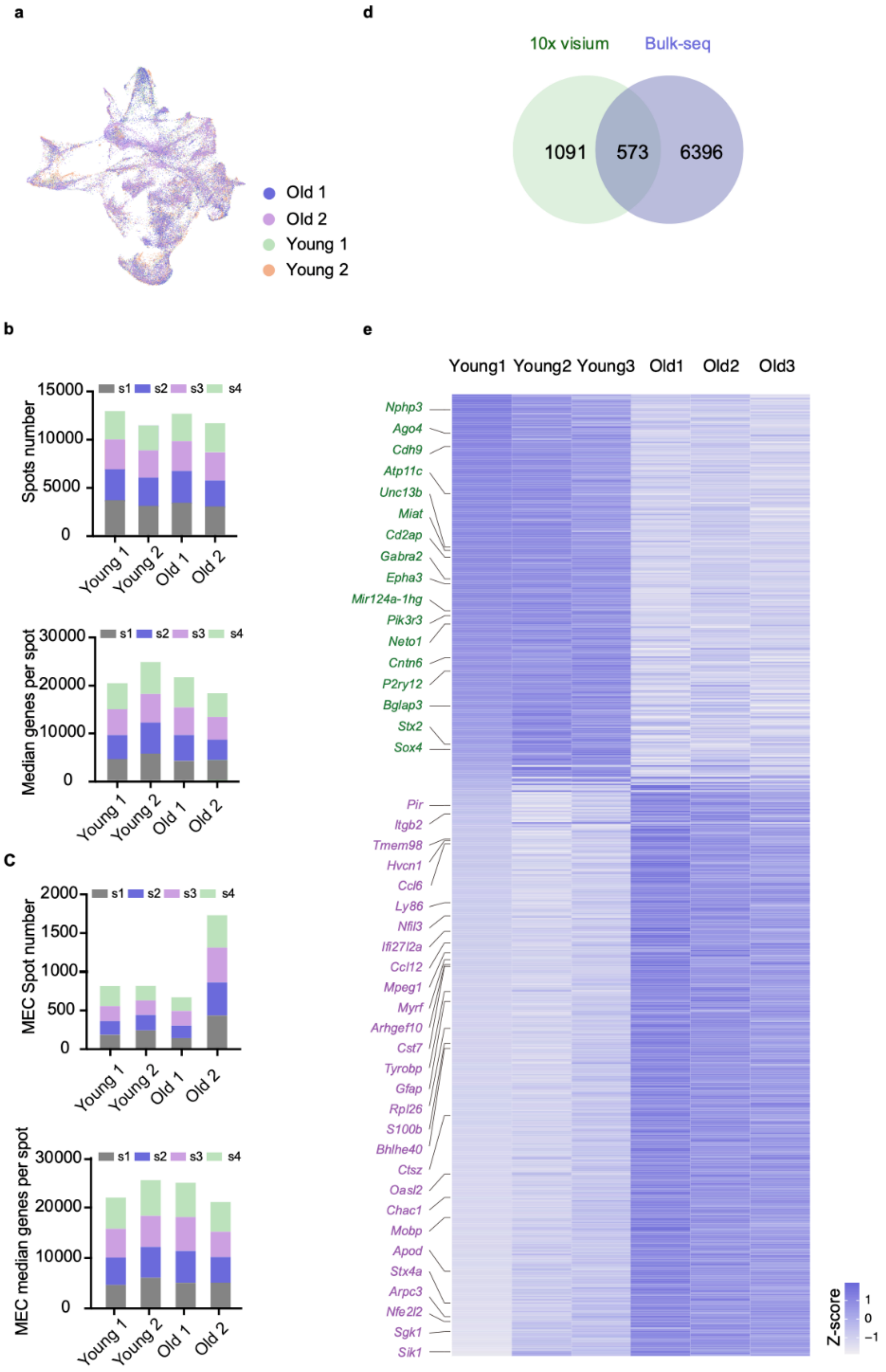

a
Old 1
Old 2
Young 1
Young 2
d
10x visium
Bulk-seq
1091
573
6396
b
s1 s2 s3 s4
Spots number
15000
10000
5000
0
Young 1
Young 2
Old 1
Old 2
Median genes per spot
30000
20000
10000
0
C
MEC Spot number
2000
1500
1000
500
0
MEC median genes per spot
e
Young1 Young2 Young3 Old1 Old2 Old3
Nphp3
Ago4
Cdh9
Atp11c
Unc13b
Miat
Cd2ap
Gabra2
Epha3
Mir124a-1hg
Pik3r3
Neto1
Cntn6
P2ry12
Bglap3
Stx2
Sox4
Pir
Itgb2
Tmem98
Hvcn1
Ccl6
Ly86
Nfil3
Ifi27l2a
Ccl12
Mpeg1
Myrf
Arhgef10
Cst7
Tyrobp
Gfap
Rpl26
S100b
Bhlhe40
Ctsz
Oasl2
Chac1
Mobp
Apod
Stx4a
Arpc3
Nfe2l2
Sgk1
Sik1
Z-score
1
0
−1

**Supplementary Fig.10 Spatial transcriptomic and bulk RNA-sequencing of old and young MEC. a** UMAP presentation of all spatial spots from 2 old and 2 young mice brain slices showing uniformity of samples. **b** Spot numbers (top) and median genes per spot (bottom) of total sagittal slices from young and old mice. s1-4 indicates sagittal slice order from medial to lateral MEC. **c** Spot numbers (top) and median genes per spot (bottom) of MEC spots from young and old mice. s1-4 indicates sagittal slice order from medial to lateral MEC. **d** Venn diagram illustrating the overlap of DEGs from MEC using 10x visium and bulk RNA-sequencing data. **e** Heatmap showing the 573 overlapped DEGs (341 up and 232 down-regulated in old mice) in MEC between 10x visium and bulk RNA-seq data. Genes that are noted are down (green) and up- (purple) regulated genes in old MEC are consistent with genes that are shown in Fig.3e. Gene expression is shown in Z-score. Source data are provided as a Source Data file. Related to Fig.3.

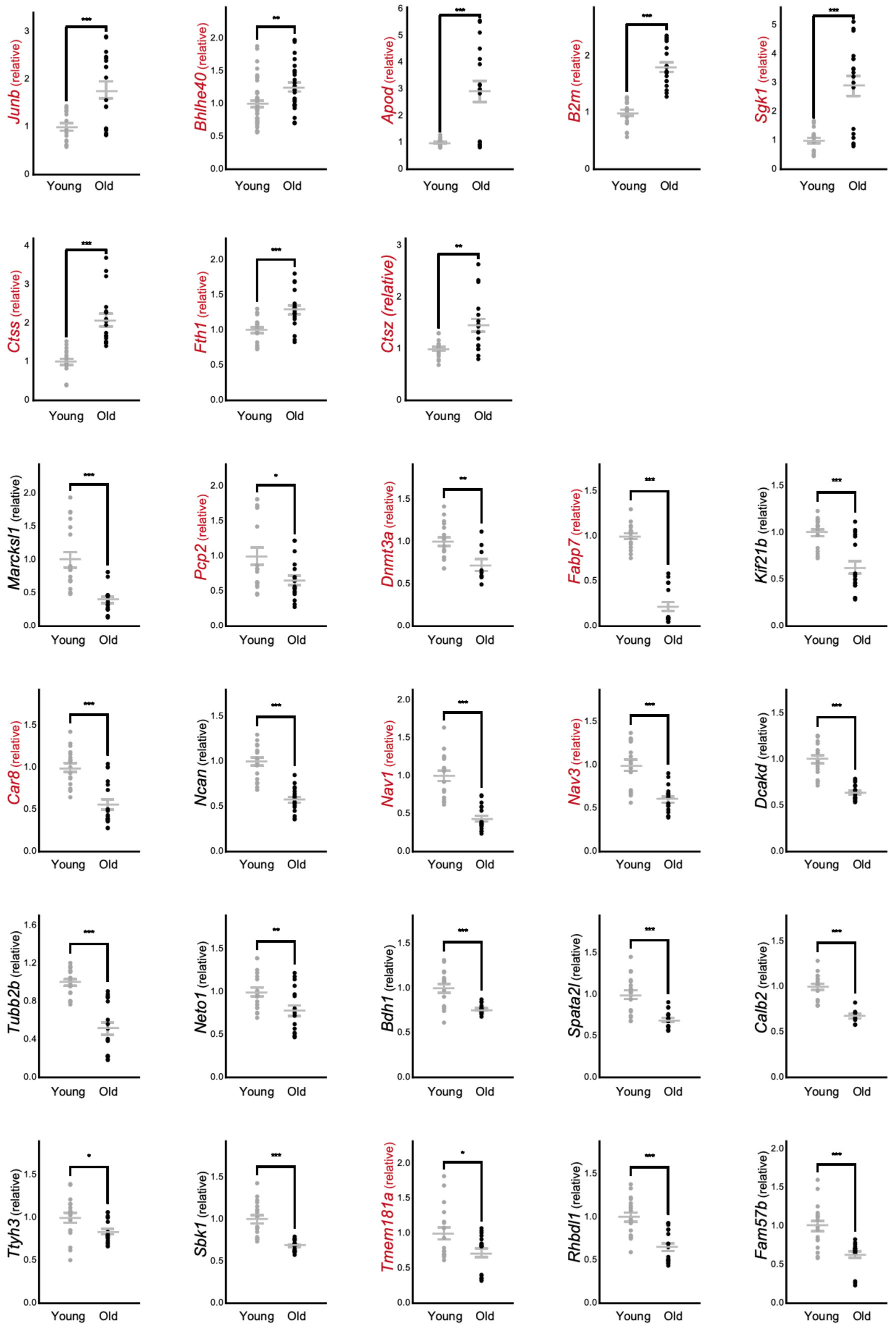

**Supplementary Fig.11 Confirmation of the 10x Genomics Visium experiment results using RT-qPCR for 8 of the top 50 upregulated genes and 20 of the top 50 downregulated genes.** Genes that are found in bulk RNA-seq data are highlighted in red. Data are presented as mean ± SEM; two-sided Mann–Whitney U-test, *P < 0.05; **P < 0.01; ***P < 0.001; ns, non-significant. Related to Fig.3.

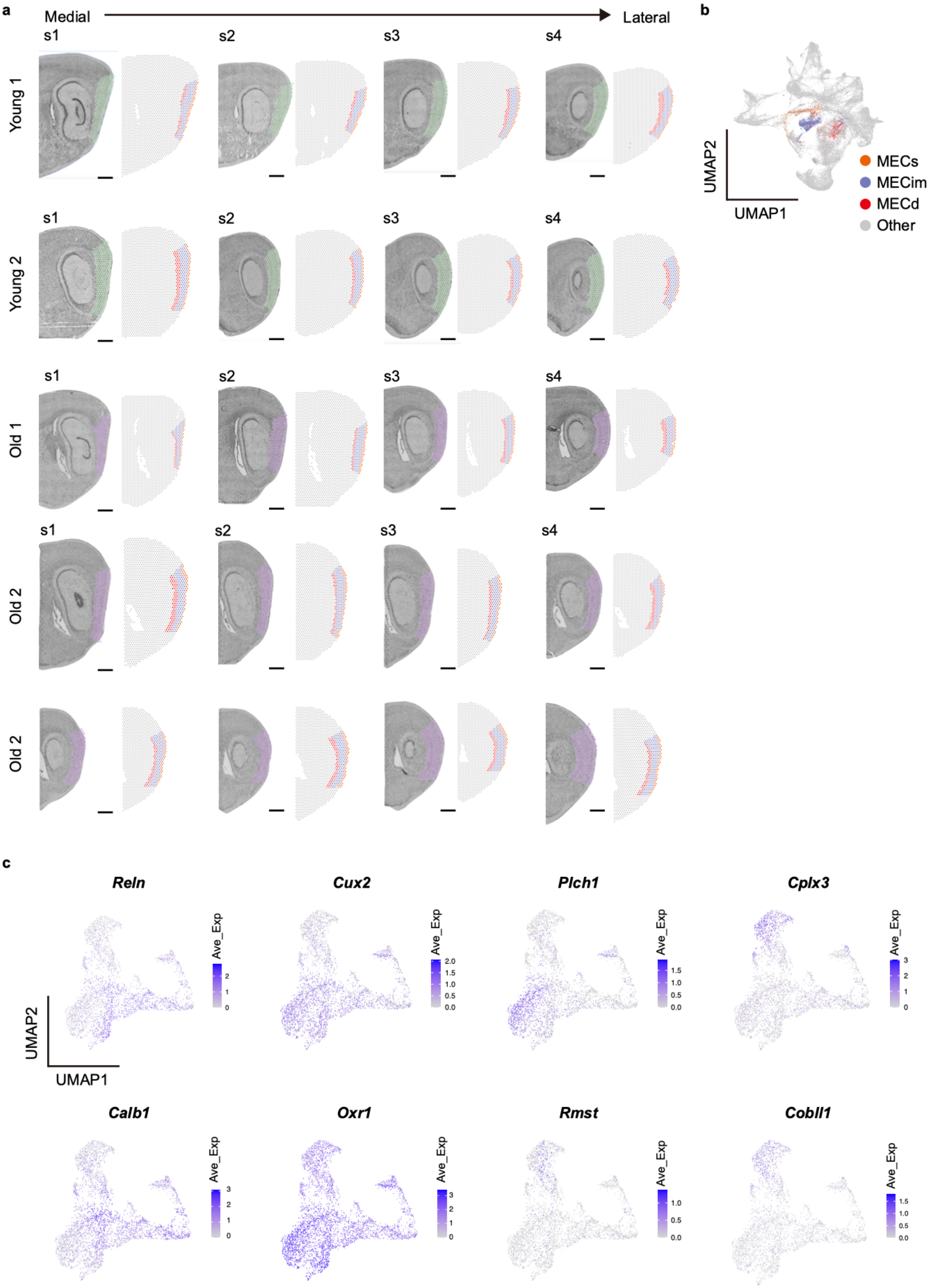

a
Medial
Lateral
s1
s2
s3
s4
Young 1
Young 2
Old 1
Old 2
Old 2
b
UMAP2
UMAP1
MECs
MECim
MECd
Other
c
Reln
Cux2
Plch1
Cplx3
Calb1
Oxr1
Rmst
Cobll1
Ave_Exp
UMAP2
UMAP1

**Supplementary Fig.12 MEC sublayer annotation. a** 20 Sagittal sections represent the MEC from young and old mice. The different MEC layers are presented by colored dots. Scale bar: 1 mm. **b** UMAP presentation of all spatial spots from 2 old and 2 young mice brain slices. Spots from three layers of MEC are shown in different colors. **c** Feature plot showing MEC marker genes' expression level in spots derived from MEC sublayers. Related to Fig.3.

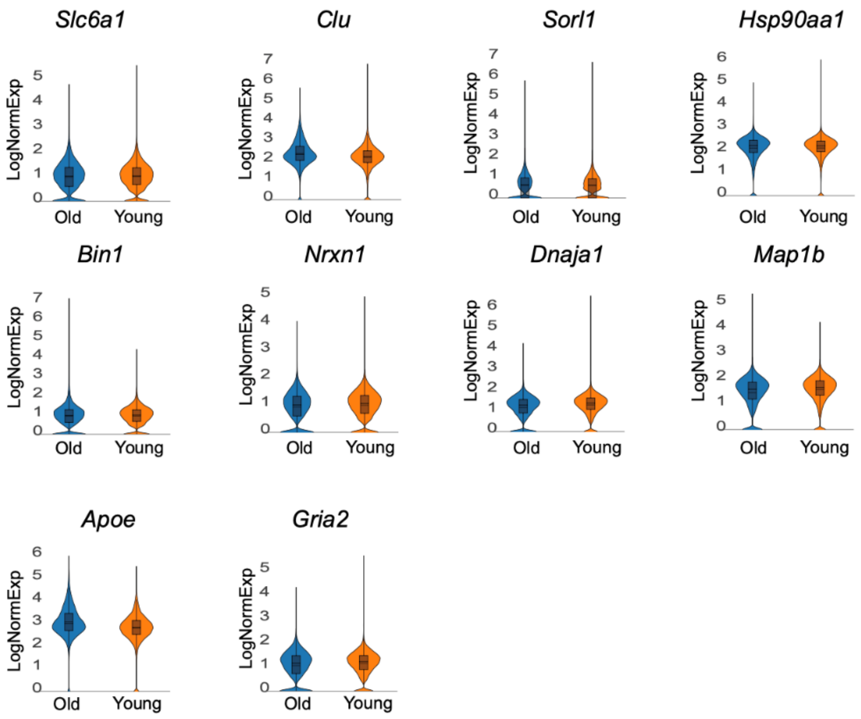

**Supplementary Fig.13 Violin plots show no clear changes in the expression of Alzheimer's Disease (AD)-related genes between old and young mice.** Mean expression of genes in two clusters are calculated in Loupe Browser 7 (see Methods). Related to Fig.3.

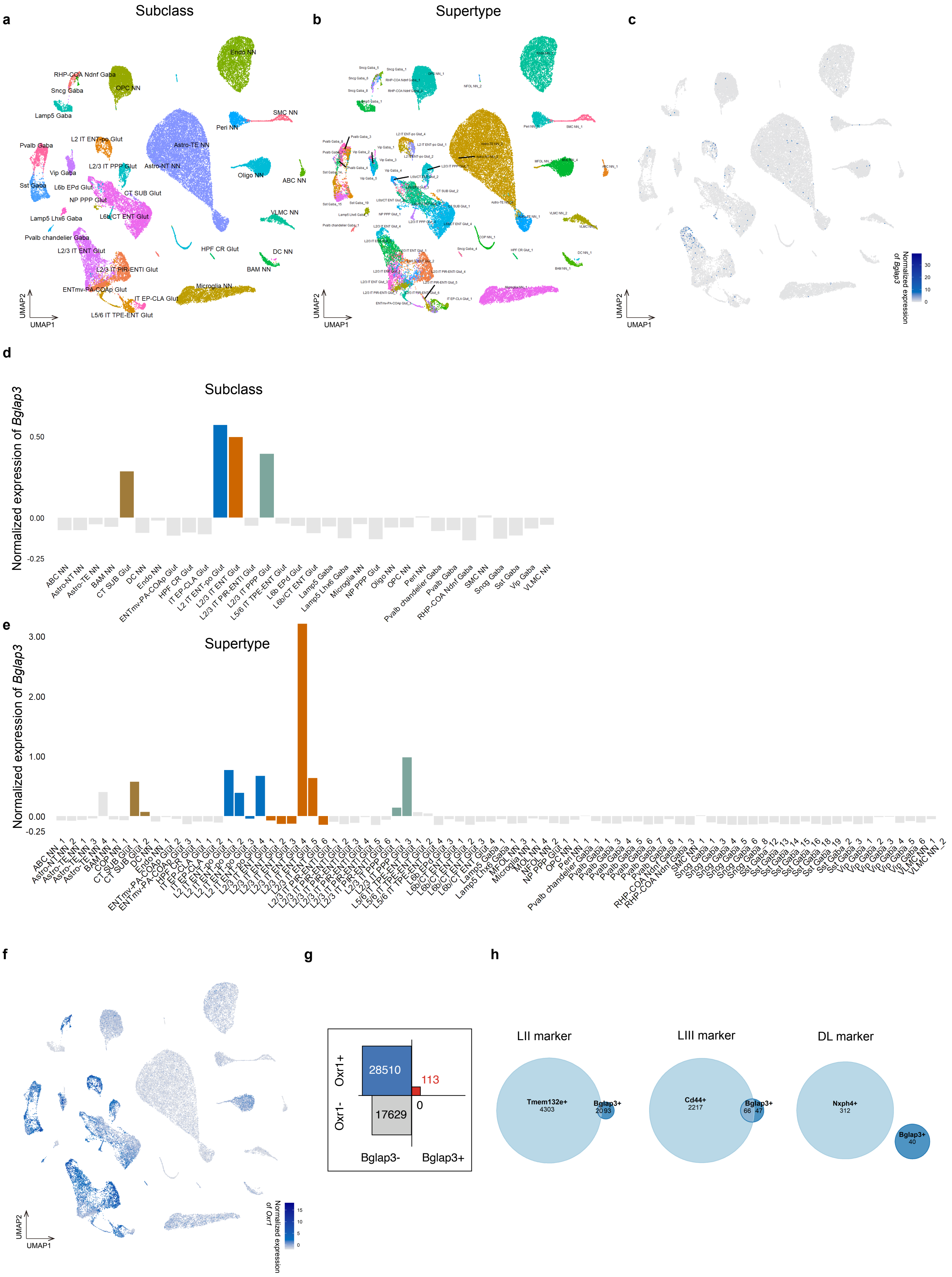
a
Subclass
Endo NN
RHP-COA Ndnf Gaba
OPC NN
Sncg Gaba
Lamp5 Gaba
SMC NN
Peri NN
L2 IT ENT-po Glut
Pvalb Gaba
Astro-TE NN
L2/3 IT PPP Glut
Astro-NT NN
Vip Gaba
Oligo NN
ABC NN
Sst Gaba
L6b EPd Glut
CT SUB Glut
NP PPP Glut
L6b/CT ENT Glut
VLMC NN
Lamp5 Lhx6 Gaba
Pvalb chandelier Gaba
HPF CR Glut
L2/3 IT ENT Glut
DC NN
L2/3 IT PIR-ENTl Glut
BAM NN
ENTmv-PA-COAp Glut
Microglia NN
IT EP-CLA Glut
L5/6 IT TPE-ENT Glut
UMAP2
UMAP1
b
Supertype
UMAP2
UMAP1
c
Normalized expression of Bglap3
30
20
10
0
UMAP2
UMAP1
d
Subclass
Normalized expression of Bglap3
0.50
0.00
-0.25
ABC NN
Astro-NT NN
Astro-TE NN
BAM NN
CT SUB Glut
DC NN
Endo NN
ENTmv-PA-COAp Glut
HPF CR Glut
IT EP-CLA Glut
L2 IT ENT-po Glut
L2/3 IT ENT Glut
L2/3 IT PIR-ENTl Glut
L2/3 IT PPP Glut
L5/6 IT TPE-ENT Glut
L6b EPd Glut
L6b/CT ENT Glut
Lamp5 Gaba
Lamp5 Lhx6 Gaba
Microglia NN
NP PPP Glut
Oligo NN
OPC NN
Peri NN
Pvalb chandelier Gaba
Pvalb Gaba
RHP-COA Ndnf Gaba
SMC NN
Sncg Gaba
Sst Gaba
Vip Gaba
VLMC NN
e
Supertype
Normalized expression of Bglap3
3.00
2.00
1.00
0.00
-0.25
f
Normalized expression of Oxr1
15
10
5
0
UMAP2
UMAP1
g
Oxr1+
Oxr1-
28510
113
17629
0
Bglap3-
Bglap3+
h
LII marker
Tmem132e+
4303
Bglap3+
2093
LIII marker
Cd44+
2217
Bglap3+
66
47
DL marker
Nxph4+
312
Bglap3+
40

**Supplementary Fig.14 scRNA-seq data from Yao et al. showing *Bglap3* expression. a** Uniform Manifold Approximation and Projection (UMAP) visualization of the subclasses of the single-cell RNA-seq data of MEC from Yao et al. (Nature, 2023)[54]. **b** UMAP visualization as in (a) for supertypes. **c** UMAP visualization as in (a) for the expression level of *Bglap3*. **d** Normalized expression of *Bglap3* in different subclasses, highlighting those with higher expression. **e** Normalized expression level of *Bglap3* in different supertypes, highlighting ones that belong to the highlighted subclasses in (d). **f** UMAP visualization as in (a) for the expression level of *Oxr1*.**g** Co-expression of *Bglap3* and Oxr1 in individual cells. **h** Number of cells co-expressing *Bglap3* and marker genes of different MEC layers. Related to Fig.4.

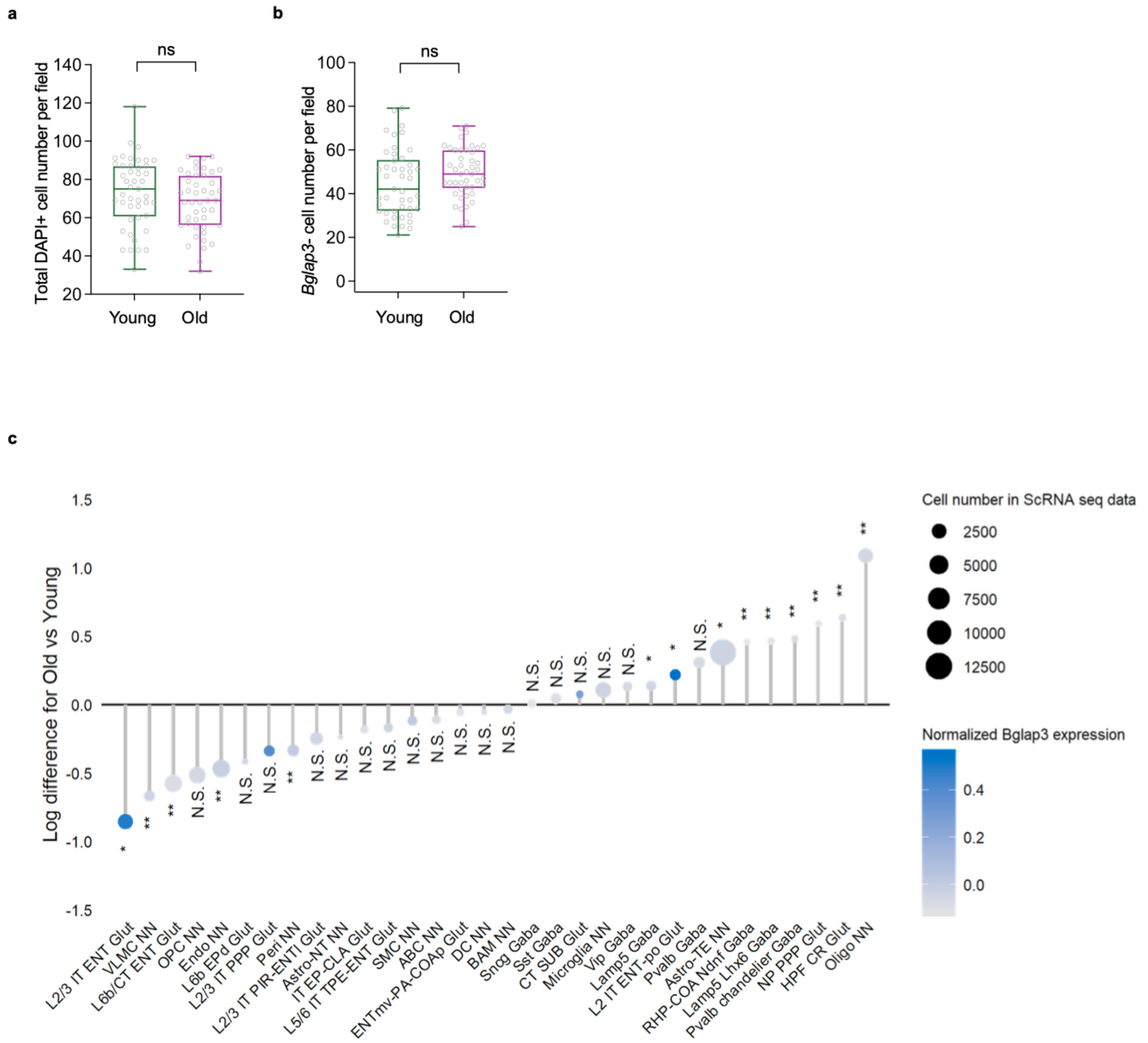

**Supplementary Fig.15 Cell fraction changes in aging MEC. a** Total DAPI+ cell number in each imaging field in young and old mice MEC ISH slices. (young, 73.11 ± 2.706, n = 45 fields of 4 mice; old, 67.93 ± 2.308, n = 45 fields of 4 mice, Linear mixed-effect model with [age] as fixed effect, SE = 3.52, t (88) = -1.47, P = 0.1455). **b** Number of DAPI+ cells that do not express *Bglap3* in each imaging field in young and old mice MEC ISH slices. (young, 44.93 ± 2.29, n = 45 fields of 4 mice; old, 49.60 ± 1.67, n = 45 fields of mice, Linear mixed-effect model with [age] as fixed effect, SE = 2.80, t (88) = 1.66, P = 0.0998). **c** Comparison of the estimated subclass fractions between young and old 10X Visium samples after deconvolution. Two-sided Wilcoxon tests are performed comparing angles. Data in (a) and (b) are presented as median with whiskers from min to max; *P < 0.05; **P < 0.01; ns, non-significant. Source data are provided as a Source Data file. Related to Fig.4.

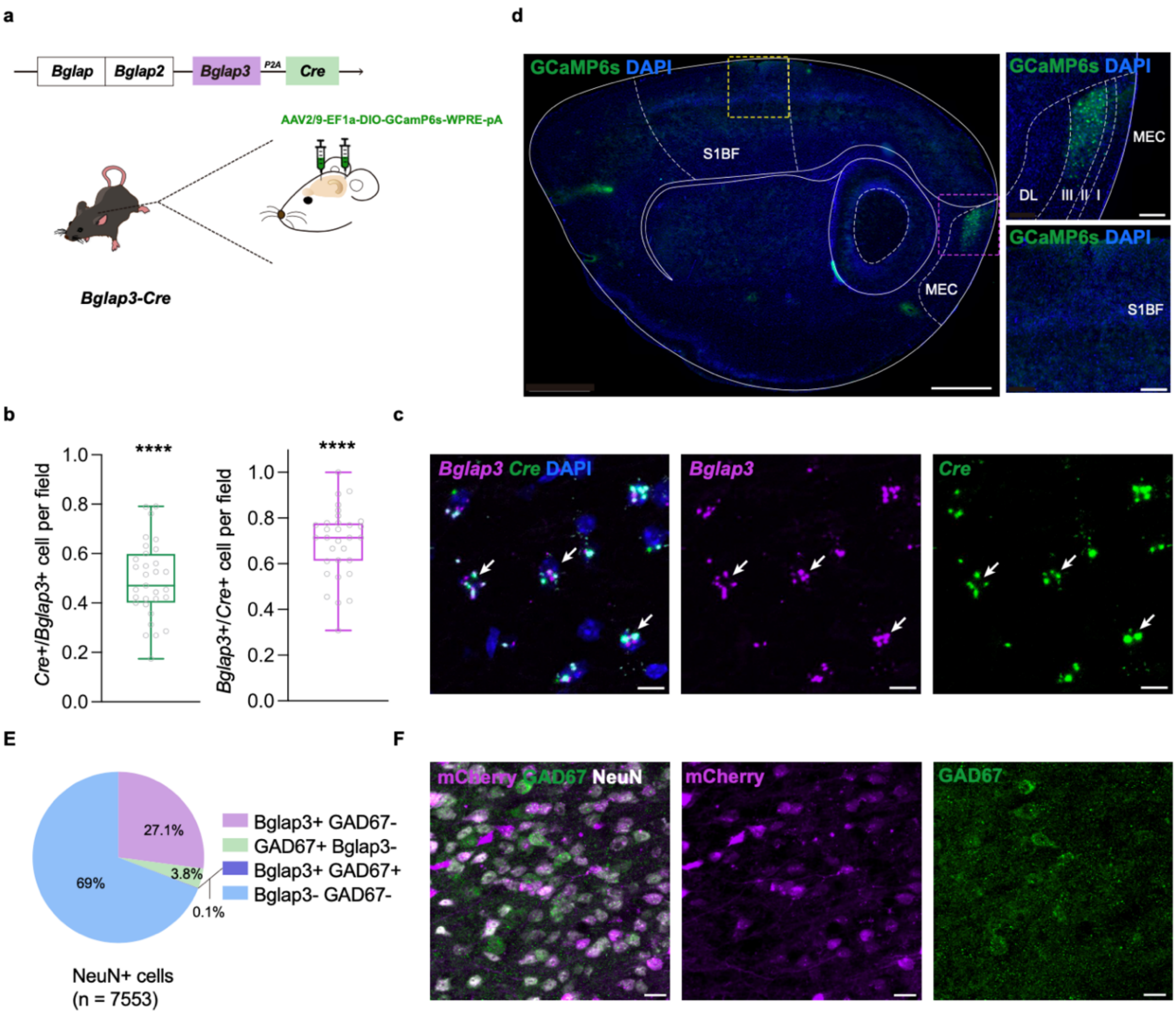

**Supplementary Fig.16 Construction and confirmation of *Bglap3-Cre* knock-in mice. a** Strategy to build the *Bglap3-Cre* transgenic mice. **b** Histogram showing the quantification of the fluorescent signal from *in situ* hybridization (ISH); the percentage of *Cre* in *Bglap3*+ cells (left panel, median = 0.4706, Wilcoxon signed-rank test, n = 31 fields, W = 496, P = 1.23 × $10^{-6}$) and the percentage of *Bglap3* in *Cre3*+ cells (right panel, median = 0.7143, Wilcoxon signed-rank test, n = 31 fields, W = 496, P = 1.222 × $10^{-6}$) are shown. **c** ISH of *Bglap3* (magenta) and *Cre* (green); white arrows indicate cells that co-express both *Bglap3* and *Cre*. **d** Histology showing the specific expression of GCaMP6 in the MEC after AAV-DIO-GCaMP6 injection in *Bglap3-Cre* knock-in mice; please note that the signal is only expressed in the MEC, but not in the primary somatosensory barrel cortex (S1BF), after AAV-DIO-GCaMP6 injection in these brain areas, which is consistent with the endogenous expression pattern of *Bglap3*. **e** Percentages of *Bglap3*+ or *Bglap3*- cells indicated by expression of DIO-hM4D-mCherry co-localizing with GAD67 in Neun-positive neurons in MEC (n = 15 FOVs of 3 mice). **f** Immunofluorescence staining images showing mCherry-expressing *Bglap3*+ cells (magenta) with Neun (white) and GAD67 (green) signals. Scale bars shown in **c**: 10 μm; **d**: 1000 μm (left), 200 μm (top right), and 200 μm (bottom right); **f**: 20 μm. Data are presented as median with whiskers from min to max; ****P < 0.0001. Source data are provided as a Source Data file. Related to Fig.4.

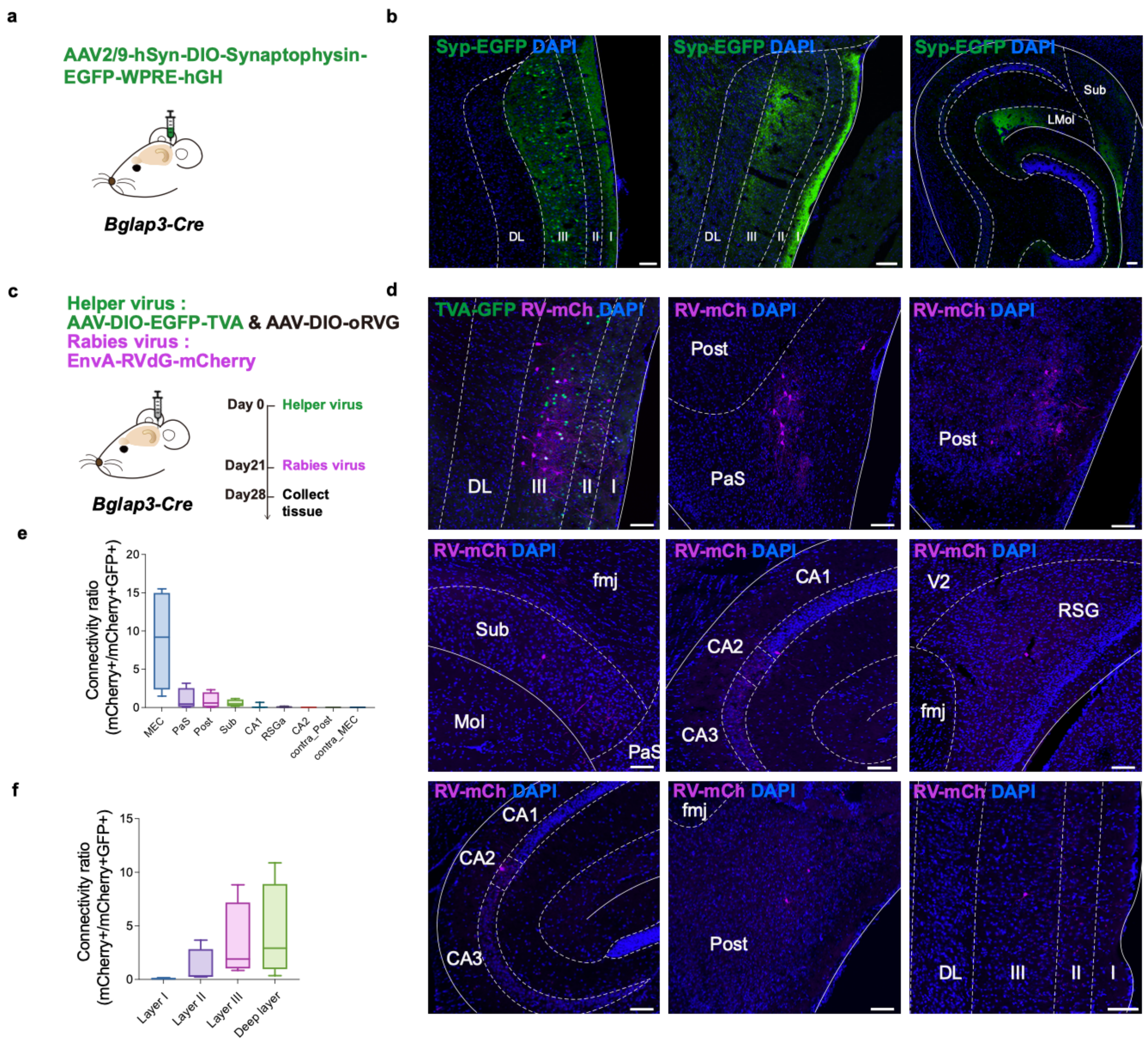

**Supplementary Fig.17 Axonal tagging and retrograde tracing of *Bglap3*+ neurons. a** The axons of *Bglap3*+ cells are labeled by synaptophysin-EGFP. **b** The axons mainly existed in the ipsilateral MEC (middle panel) and the prefrontal pathway in the hippocampus (right panel); the green signal is the soma and axon of the infected *Bglap3*+ cells and the blue signal is DAPI. **c** The experimental diagram shows the rabies virus infection of *Bglap3*+ cells to label their presynaptic cells with mCherry. **d** Representative image shows the presynaptic cells to the *Bglap3*+ cells in different brain areas, including the MEC, parasubiculum (PaS), postsubiculum (Post), subiculum (Sub), hippocampus CA1, CA2, retrosplenial cortex (RSG), contralateral postsubiculum (contra_Post) and contralateral MEC (contra_MEC).**e** Box and whisker plot shows the distribution of the presynaptic cells to the *Bglap3*+ cells in different brain areas (86 starter cells from n = 4 mice). **f** Box and whisker plot showing the distribution of the presynaptic cells to the *Bglap3*+ cells in different layers of the MEC (n = 4 mice). Data are presented as median with min to max. Scale bars shown in **b**: 100μm; **d**: 100 μm. Source data are provided as a Source Data file. Related to Fig.4.

a

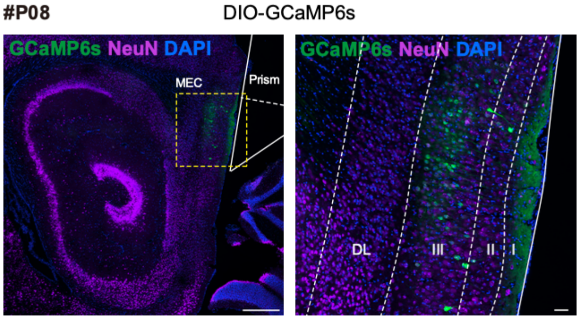
#P08
DIO-GCaMP6s
GCaMP6s NeuN DAPI
MEC
Prism
GCaMP6s NeuN DAPI
DL
III
II
I

b

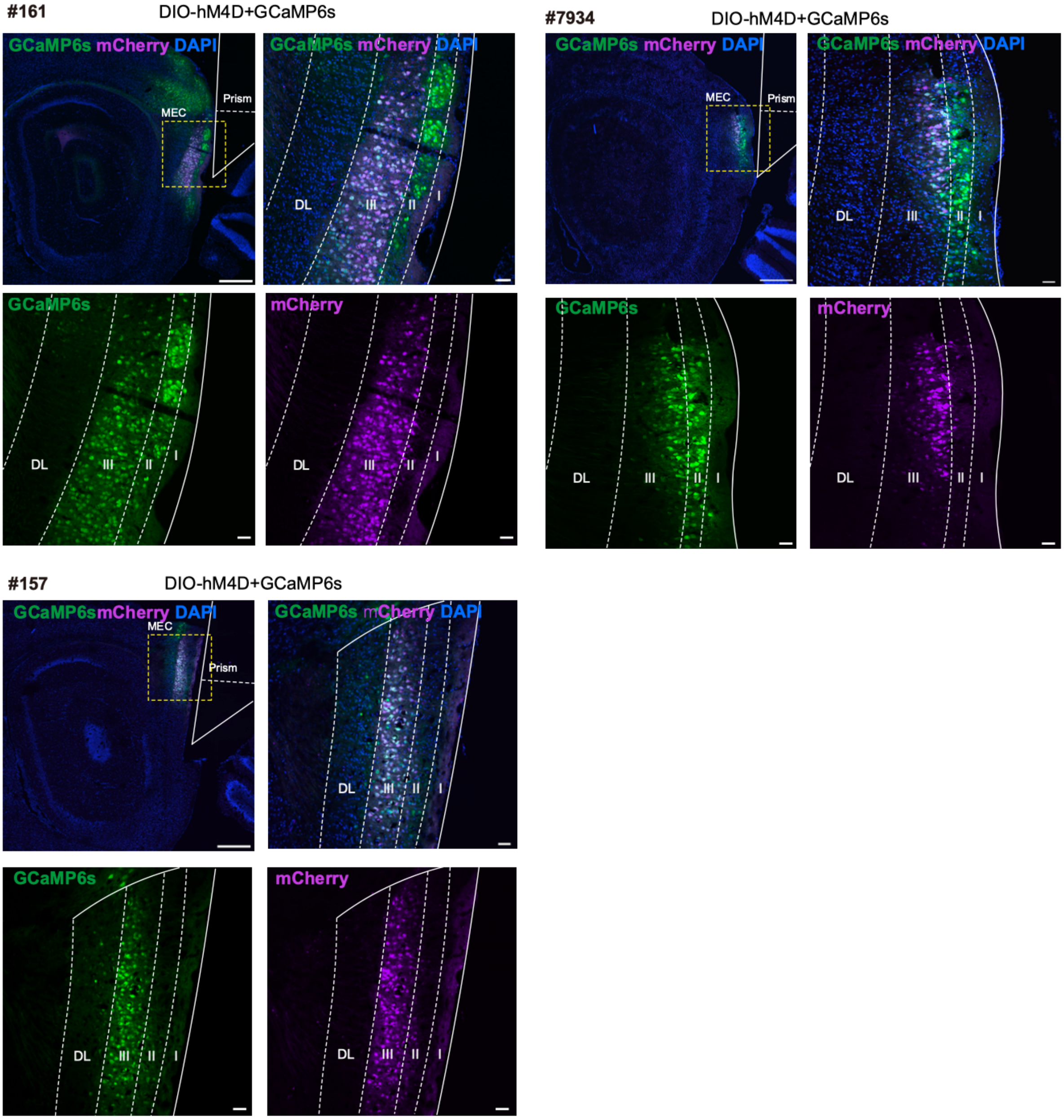
#161
DIO-hM4D+GCaMP6s
GCaMP6s mCherry DAPI
MEC
Prism
GCaMP6s mCherry DAPI
DL
III
II
I
GCaMP6s
mCherry
#7934
DIO-hM4D+GCaMP6s
#157
DIO-hM4D+GCaMP6s
GCaMP6smCherry DAPI

**Supplementary Fig.18 Histology of 2-photon calcium imaging mouse brain sections. a** DIO-GCaMP6s expression in MEC (green) with co-staining of NeuN (magenta). **b** GCaMP6s (green) and DIO - hM4D (magenta) co-expression in MEC of three TPM calcium imaging mice shown in whole sagittal view (top left), enlarged view (top right) and individual fluorescence view (lower left and right). The white outlines indicate the position of the prism. LI: Layer 1; LII: Layer 2; LIII: Layer 3; DL: Deep layer; Scale bars are 500 μm (in whole sagittal section) and 50 μm (in enlarged view). Related to Fig.5.

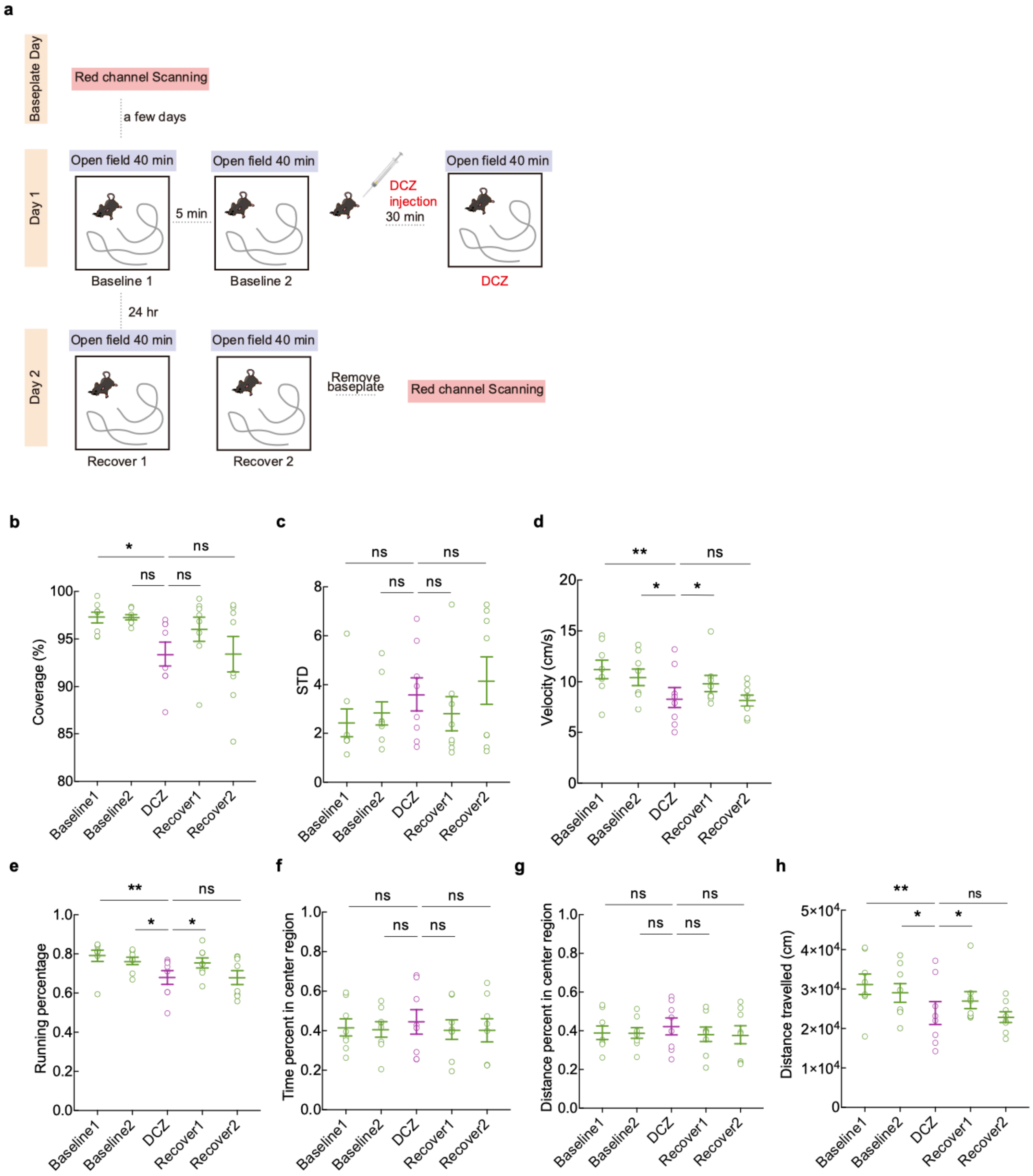
a
Baseplate Day
Red channel Scanning
a few days
Day 1
Open field 40 min
Open field 40 min
Open field 40 min
5 min
DCZ
injection
30 min
Baseline 1
Baseline 2
DCZ
24 hr
Day 2
Open field 40 min
Open field 40 min
Remove
baseplate
Red channel Scanning
Recover 1
Recover 2
b
Coverage (%)
c
STD
d
Velocity (cm/s)
e
Running percentage
f
Time percent in center region
g
Distance percent in center region
h
Distance travelled (cm)
Baseline1
Baseline2
DCZ
Recover1
Recover2
ns
*
**

**Supplementary Fig.19 Two-photon calcium imaging of *Bglap3*-Cre mice. a** Schematic of the open-field/chemogenetic paradigm. Red-channel scans were taken before and after the 2-day protocol: day 1, two 40-min baseline sessions, then intraperitoneal DCZ to suppress *Bglap3*+ MEC neurons, a 30-min rest, and a 40-min experimental session; day 2, two 40-min recovery sessions. **b** Trajectory coverage across five open-field sessions, one point per 40-min session per mouse (q vs. DCZ: BL1 = 0.0325, BL2 = 0.0558, RC1 = 0.0634, RC2 = 0.7273). **c** Uniformity of trajectory coverage across the five sessions (q vs. DCZ: BL1 = 0.1410, BL2 = 0.8338, RC1 = 0.2152, RC2 = 0.9181). **d** Average velocity across the five sessions (q vs. DCZ: BL1 = 0.0019, BL2 = 0.0120, RC1 = 0.0188, RC2 = 0.3672). **e** Percentage running (velocity > 2.5 cm/s) across the five sessions (q vs. DCZ: BL1 = 0.0032, BL2 = 0.0240, RC1 = 0.0279, RC2 = 0.5263). **f** Proportion of time in the central area across the five sessions (q vs. DCZ: BL1 = 0.7045, BL2 = 0.7045, RC1 = 0.7045, RC2 = 0.7045). **g** Proportion of trajectories in the central area across the five sessions (q vs. DCZ: BL1 = 0.4062, BL2 = 0.3985, RC1 = 0.3985, RC2 = 0.3985). **h** Total distance traveled across the five sessions (q vs. DCZ: BL1 = 0.0019, BL2 = 0.0186, RC1 = 0.0188, RC2 = 0.4200). Statistics (n = 8 sessions) are from Friedman tests with two-stage linear step-up correction (Benjamini, Krieger, Yekutieli; Q = 0.05). Data are represented as mean ± SEM. **P < 0.01; *P < 0.05; ns, not significant. Source data are provided as a Source Data file. Related to Fig.5.

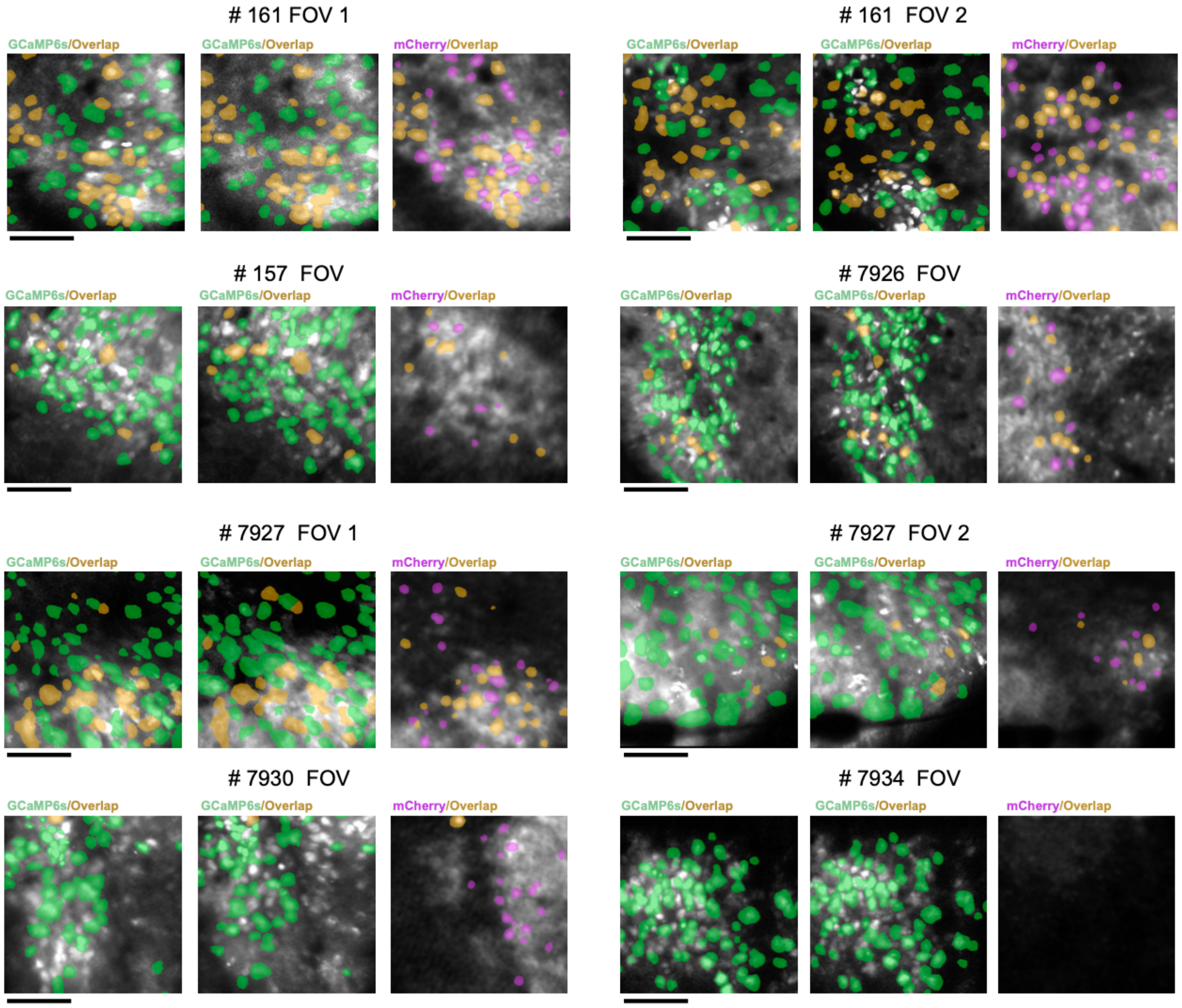

**Supplementary Fig.20 8 FOVs of 6 imaging mice.** Selected from Baseline 1, Baseline 2, and DCZ sessions on day 1 (left panel, GCaMP6s), Recovery sessions 1&2 on day2 (middle panel, GCaMP6s). mCherry signal on the right panels shows *Bglap3*+ neurons in each FOV, with yellow shaded ROIs indicating *Bglap3*+ neurons in both channels. Scale bar: 100 μm. Related to Fig.5.

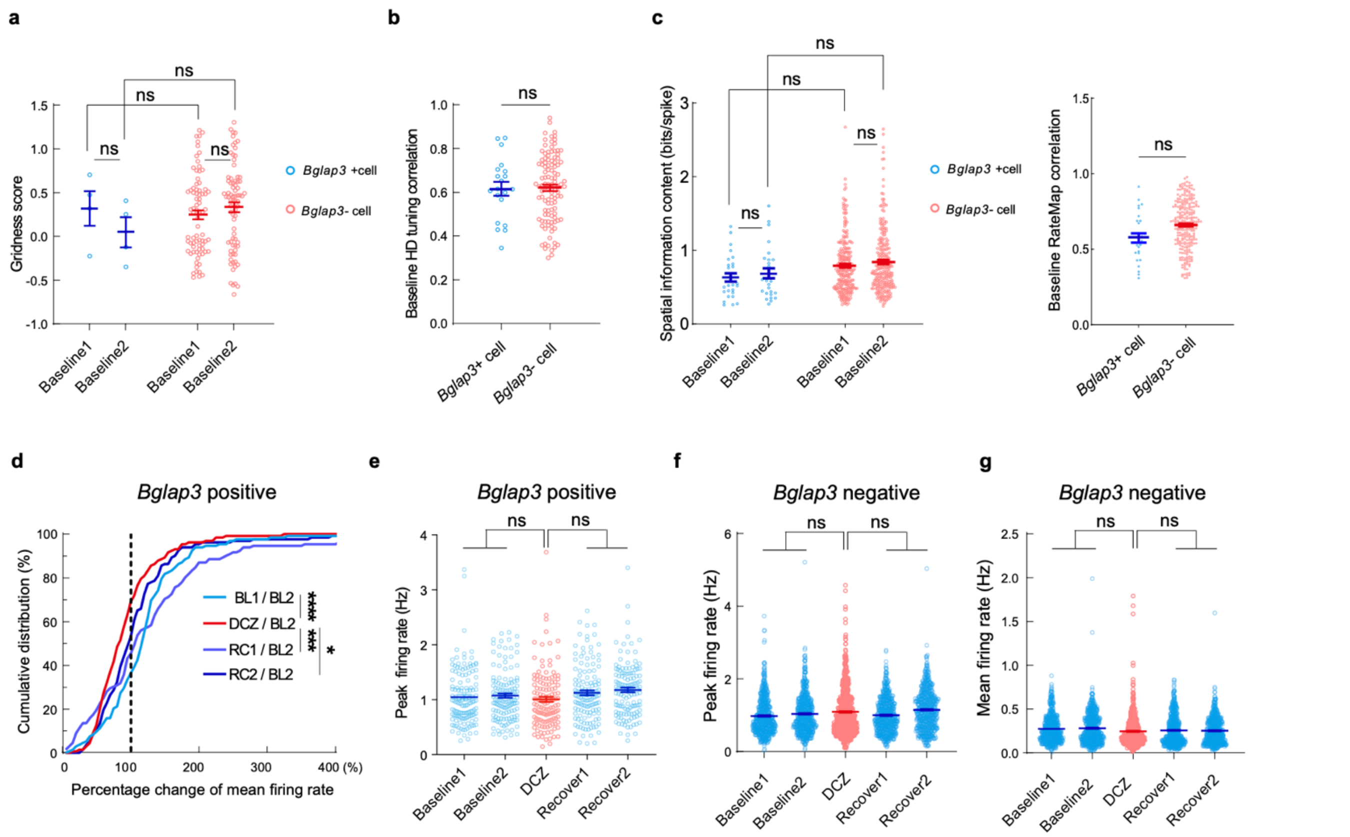

**Supplementary Fig.21 Analysis of *Bglap3*+/- cells. a** Grid scores, BL1/BL2 (*Bglap3*+, blue, n = 4: 0.32 ± 0.20/0.05 ± 0.17; *Bglap3*−, red, n=77: 0.25 ± 0.05/0.33 ± 0.06). LME (Value ~ Group × Phase + (1|MouseID)): Group, t(158) = -0.82, P = 0.413; Phase, t(158) = -0.85, P = 0.394; Group×Phase, t(158) = 1.10, P=0.274. Post hoc [F(1,158), P]: BL1–BL2(+), 0.73, 0.39; BL1–BL2(−), 1.42, 0.23; −vs+@BL1, 0.67, 0.41; @BL2, 0.45, 0.50. **b** HD-cell tuning-curve correlation, BL1/BL2 (*Bglap3*+, blue, n = 20: 0.62 ± 0.03; *Bglap3*−, red, n = 110: 0.62 ± 0.01). Model as in **a**: Group, t(128) = 0.32, P = 0.75. **c** Left: Spatial information, spatially tuned cells, BL1/BL2 (*Bglap3*+, blue, n = 27: 0.63 ± 0.06/0.69 ± 0.07; *Bglap3*−, red, n = 254: 0.80 ± 0.02/0.84 ± 0.03). Model as in **a**: Group, t(558) = -0.09, P = 0.927; Phase, t(558) = 0.63, P=0.532; Group×Phase, t(558) = -0.07, P = 0.944. Post hoc [F(1,558), P]: BL1–BL2(+), 0.39, 0.53; BL1–BL2(−), 2.86, 0.09; −vs+@BL1, 0.008, 0.92; @BL2, 0.036, 0.84. Right: Ratemap correlation, same cells, BL1/BL2 (*Bglap3*+, 0.58 ± 0.03; *Bglap3*−, 0.66 ± 0.01); linear model, t(279) = 0.92, P = 0.36. **d** Cumulative change in *Bglap3*+ firing rate (n=132 cells, 6 mice) at BL1, DCZ (+30 min), RC1 (+24 h), RC2, vs. BL2 (dashed line = no change); Kolmogorov-Smirnov P: BL1 < 0.001, RC1 = 0.001, RC2 = 0.0364. **e-g** Peak **e, f** and mean **g** firing rates (Hz), order BL1/BL2/DCZ/RC1/RC2; LME ANOVA (Condition, random intercept MouseID). **e** *Bglap3*+, n = 132 cells/6 mice: 1.05 ± 0.04/1.07 ± 0.04/1.01 ± 0.05/1.13± 0.04/1.18 ± 0.04; Condition, F(2,657) = 0.53, P = 0.59. **f** *Bglap3*−, n = 669 cells/6 mice: 0.98 ± 0.02/1.04 ± 0.02/1.09 ± 0.02/1.00 ± 0.02/1.15 ± 0.02; Condition, F(1,3342) = 0.073, P = 0.93. **g** *Bglap3*−, same n: 0.27 ± 0.01/0.28 ± 0.01/0.24 ± 0.01/0.26 ± 0.01/0.25 ± 0.01; Condition, F(1,3342) = 0.65, P = 0.52. Data are mean ± SEM. *P < 0.05; ***P < 0.001; ****P < 0.0001; ns, not significant. Source data are provided as a Source Data file. Related to Fig.5.

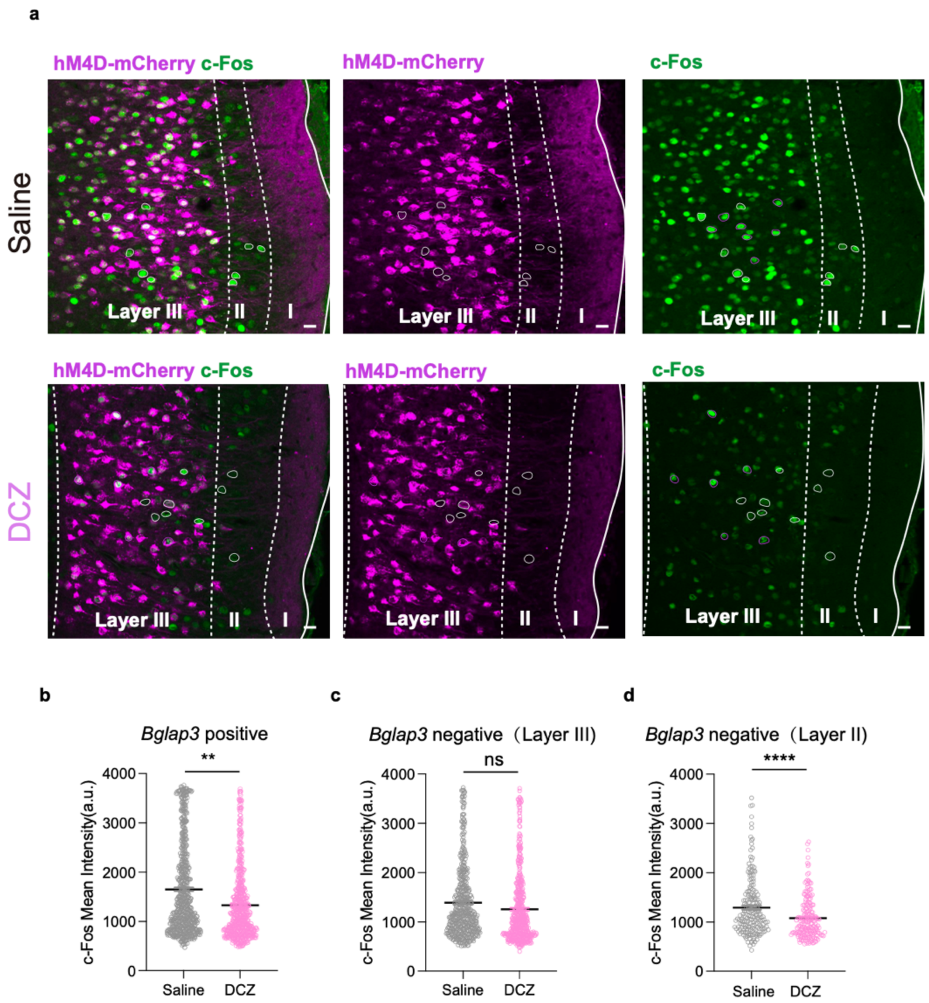

**Supplementary Fig.22 c-Fos intensity decreased in DCZ injection group. a** c-Fos staining of MEC brain slices from *Bglap3-Cre* mice injected with AAV-DIO-hM4Di-mCherry and perfused after DCZ or saline i.p. injection. *Bglap3*+ cells expressing mCherry are in magenta and c-Fos signals are in green. Selected *Bglap3*+/- cells are outlined in either magenta or white. (Saline: n = 8 FOVs of 2 mice; DCZ: n = 8 FOVs of 3 mice). **b** Mean intensities of c-Fos signals in mCherry-expressing *Bglap3+ cells* (Saline: n = 676 cells, DCZ: n = 612 cells, Linear mixed-effect model with condition as fixed effect, FOV as random effect, DCZ as reference, SE = 137.8, t (1286) = 2.66, P = 0.0080). **c** Mean intensities of c-Fos signals in *Bglap3*- cells in MEC layer III (Saline: n = 506 cells, DCZ: n = 546 cells, Linear mixed-effect model with condition as fixed effect, FOV as random effect, DCZ as reference, SE = 116.74, t (1050) = 0.76, P = 0.4475). **d** Mean intensities of c-Fos signals in *Bglap3*- cells in MEC layer II (Saline: n = 197 cells, DCZ: n = 205 cells, Linear mixed-effect model with condition as fixed effect, FOV as random effect, DCZ as reference, SE = 50.53, t (400) = 4.25, P = 2.67 × $10^{-5}$). Data are presented as mean ± SEM. Scale bars in **a** are 20μm. **P < 0.01; ****P < 0.0001; ns, non-significant. Source data are provided as a Source Data file.Related to Fig.5.

**a**

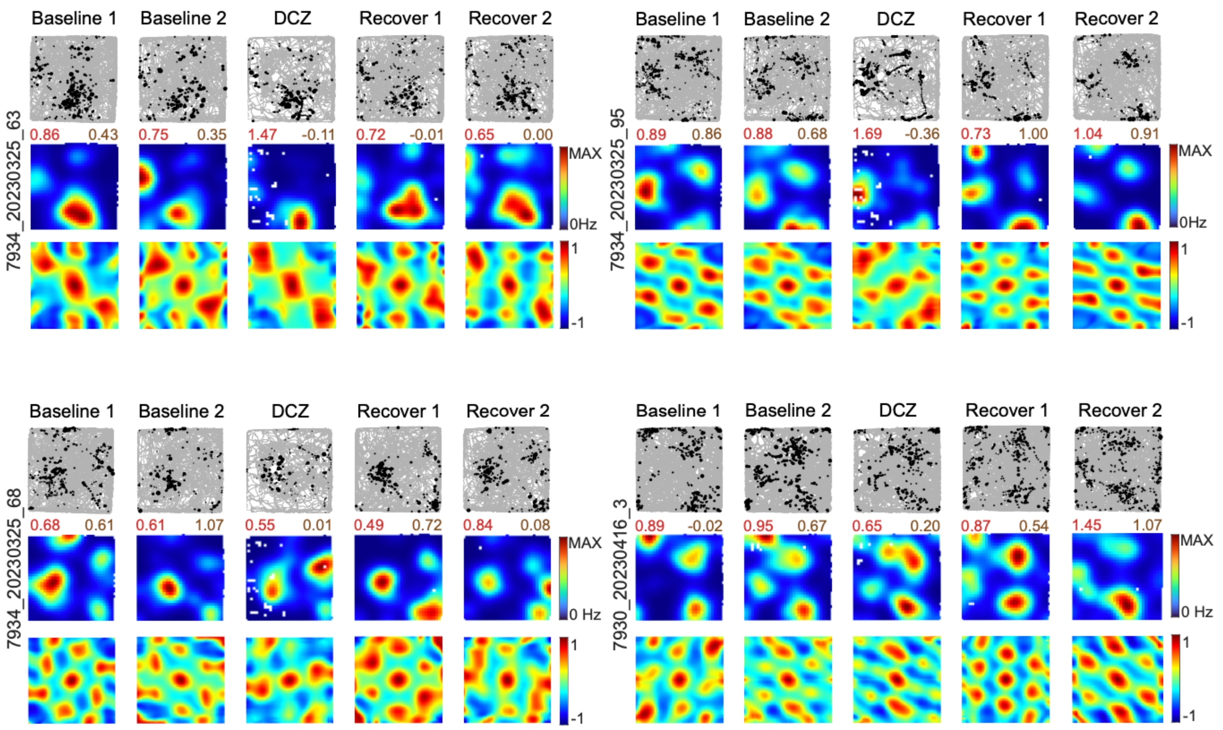

Baseline 1
Baseline 2
DCZ
Recover 1
Recover 2
7934_20230325_63
0.86 0.43
0.75 0.35
1.47 -0.11
0.72 -0.01
0.65 0.00
MAX
0Hz
1
-1
7934_20230325_95
0.89 0.86
0.88 0.68
1.69 -0.36
0.73 1.00
1.04 0.91
7934_20230325_68
0.68 0.61
0.61 1.07
0.55 0.01
0.49 0.72
0.84 0.08
0 Hz
7930_20230416_3
0.89 -0.02
0.95 0.67
0.65 0.20
0.87 0.54
1.45 1.07

**b**

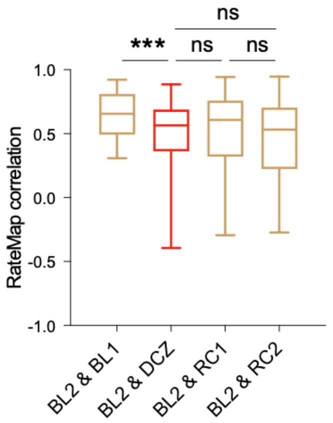

ns
***
ns
ns
RateMap correlation
1.0
0.5
0.0
-0.5
-1.0
BL2 & BL1
BL2 & DCZ
BL2 & RC1
BL2 & RC2

**Supplementary Fig.23 Analysis of *Bglap3*- grid cells. a** 4 example grid cells in total 5 recording sessions: 2 Baseline, 1 after DCZ injection, and 2 sessions after DCZ recovery, were shown with trajectory maps (top), representative color-coded rate maps (middle), and autocorrelograms (bottom); peak rates (red) and grid scores (brown) are indicated above each rate map; animal and cell numbers are indicated to the left of rate maps. **b** Correlation of firing rate maps of *Bglap3*- grid cells in five sessions (n = 77 cells, from 6 mice; Correlation, BL2 & BL1: 0.64 ± 0.02, BL2 & DCZ: 0.53 ± 0.03, BL2 & RC1: 0.52 ± 0.03, BL2 & RC2: 0.48 ± 0.04; A Linear mixed-effects model was used to assess the effect of PairType on correlation values (Value ~ PairType + (1|MouseID)). PairType: $F(3, 303.05) = 9.53$, $P = 4.96 \times 10^{-6}$). Post-hoc Tukey–Kramer contrasts showed that BL2&DCZ correlations were significantly lower than BL2&BL1 (SE = 0.12, $t(304) = -3.60$, $P = 3.70 \times 10^{-4}$), whereas correlations for BL2&RC1 and BL2&RC2 did not differ from BL2&DCZ (difference = 0.005, P = 0.87 and difference = 0.049, P = 0.14, respectively), nor did BL2&RC1 differ from BL2&RC2 (difference = 0.044, P = 0.18). Data are presented as median with whiskers from min to max; ***P < 0.001; ns, non-significant. Source data are provided as a Source Data file. Related to Fig.5.

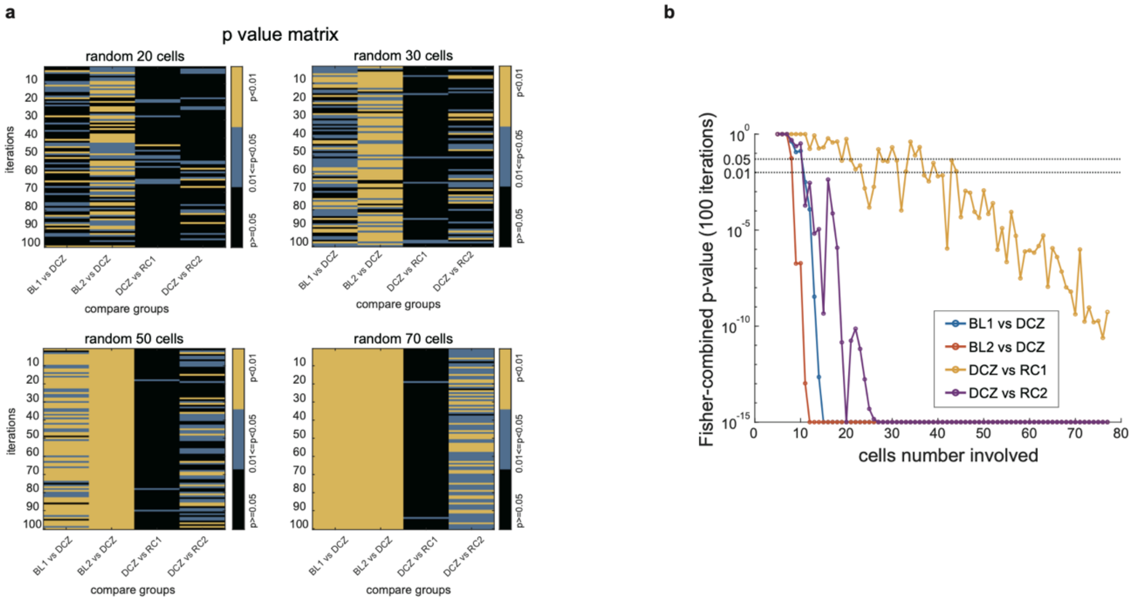

**Supplementary Fig.24 Effect of chemogenetic inhibition on grid score across varying numbers of randomly selected non-*Bglap3* grid cells**. **a** p-value matrix for grid score statistics across different sample sizes of grid cells (n = 20, 30, 50, 70). For each test, n grid cells were randomly sampled and analyzed using paired non-parametric Wilcoxon signed-rank tests, repeated 100 times. The color bar indicates p-values: yellow: $p < 0.01$; blue: $0.01 \leqslant p < 0.05$; black: ns. **b** Fisher-combined p-values are plotted as a function of the number of randomly sampled grid cells (x-axis), based on 100 iterations per sample size. Each curve represents a different comparison: baseline day 1 (BL1) vs DCZ, baseline day 2 (BL2) vs DCZ, and DCZ vs recovery days (RC1, RC2). Dashed horizontal lines indicate standard significance thresholds at p = 0.05 and p = 0.01. Extremely small p-values ($< 1 \times 10^{-15}$) were clipped to $1 \times 10^{-15}$ for stable log-scale visualization. The results show that significant effects of DCZ-induced grid score reduction consistently emerge with as few as 10–20 grid cells in BL vs DCZ conditions. Related to Fig.5.

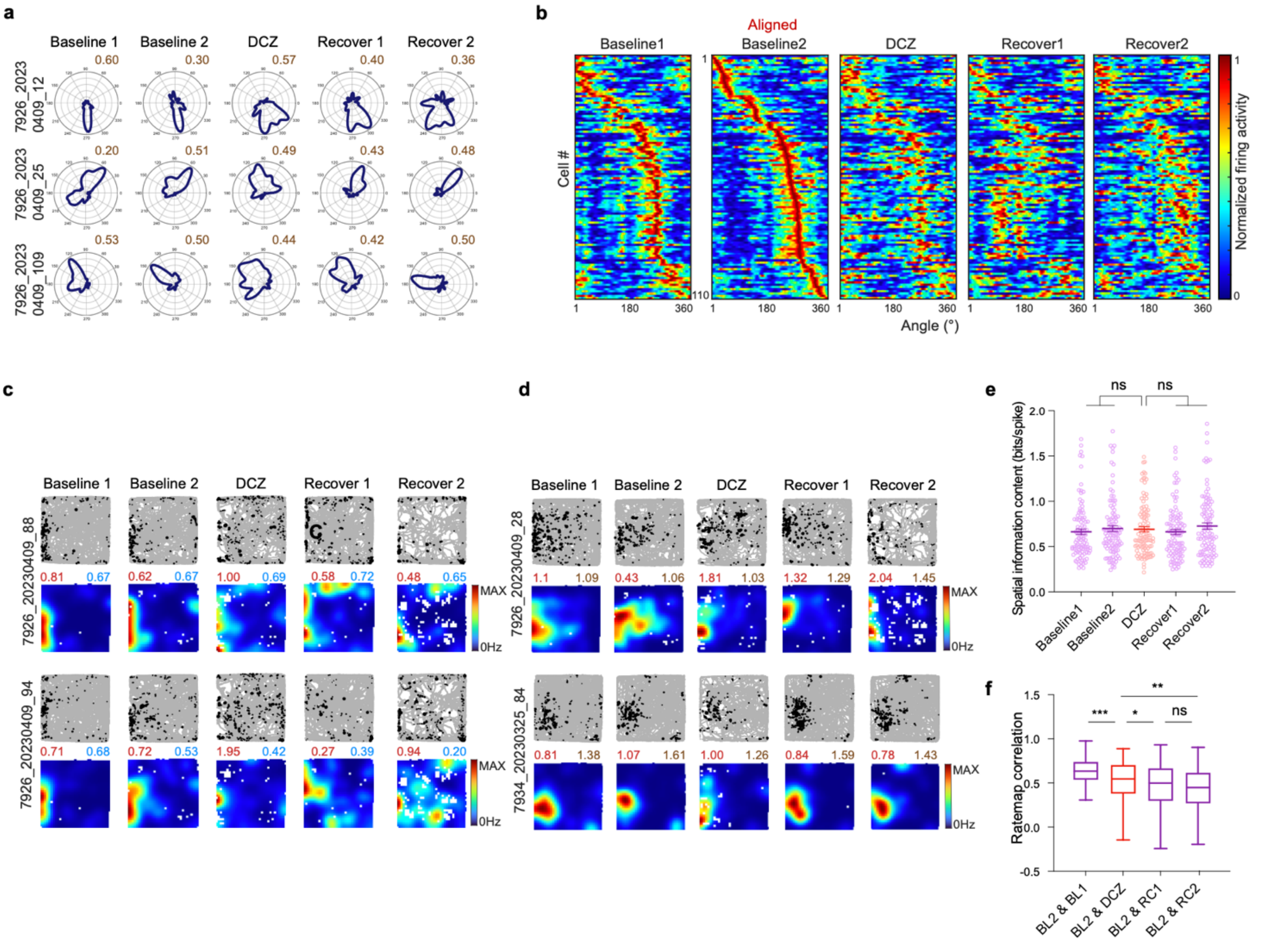

**Supplementary Fig.25 *Bglap3*- HD cell, border cell, and spatial cell changes after *Bglap3*+ neuron inactivation. a** Polar plots of HD cells before, during, and after DCZ injection; firing rate vs. head direction (blue), mean vector length (brown) above each plot; animal/cell numbers at left. **b** Population activity of *Bglap3*− HD cells (n = 110 cells, 6 mice), sorted by preferred angle in BL2; each row shows directional activity across the five sessions, normalized to each session's peak firing rate. **c** Border cells before, during, and after DCZ injection: trajectory maps (top) and representative rate maps (bottom). **d** Two example *Bglap3*− spatial cells before/after silencing; peak rate (red) and spatial information (bits/event, brown) above each map; animal/cell numbers at left. **e** Spatial information (bits/spike) of *Bglap3*− spatial cells across five sessions (n = 103 cells, 6 mice): BL1 0.67 ± 0.03, BL2 0.70 ± 0.03, DCZ 0.70 ± 0.03, RC1 0.66 ± 0.03, RC2 0.72 ± 0.03. LME (Value ~ Condition + (1|MouseID)): Condition, $F(2,512) = 0.32$, $P = 0.73$. **f** Firing-ratemap correlation of *Bglap3*− spatial cells across session pairs: BL2&BL1 0.63 ± 0.01, BL2&DCZ 0.52 ± 0.02, BL2&RC1 0.45 ± 0.03, BL2&RC2 0.43 ± 0.03. LME (Value ~ PairType + (1|MouseID)): PairType, $F(3,406.34) = 17.55$, $P = 9.81 \times 10^{-11}$. Tukey-Kramer post hoc: BL2&DCZ < BL2&BL1 ($P = 0.00018$); BL2&RC1 ($\Delta = 0.069$, $P = 0.026$) and BL2&RC2 ($\Delta = 0.085$, $P = 0.0066$) both < BL2&DCZ; BL2&RC1 vs. BL2&RC2, $\Delta = 0.016$, $P = 0.616$ (ns). Δ represents difference between conditions. Data in **e**, mean ± SEM; in **f**, median with min-to-max whiskers. *P < 0.05; **P < 0.01; ***P < 0.001; ****P < 0.0001; ns, not significant. Source data are provided as a Source Data file. Related to Fig.5.

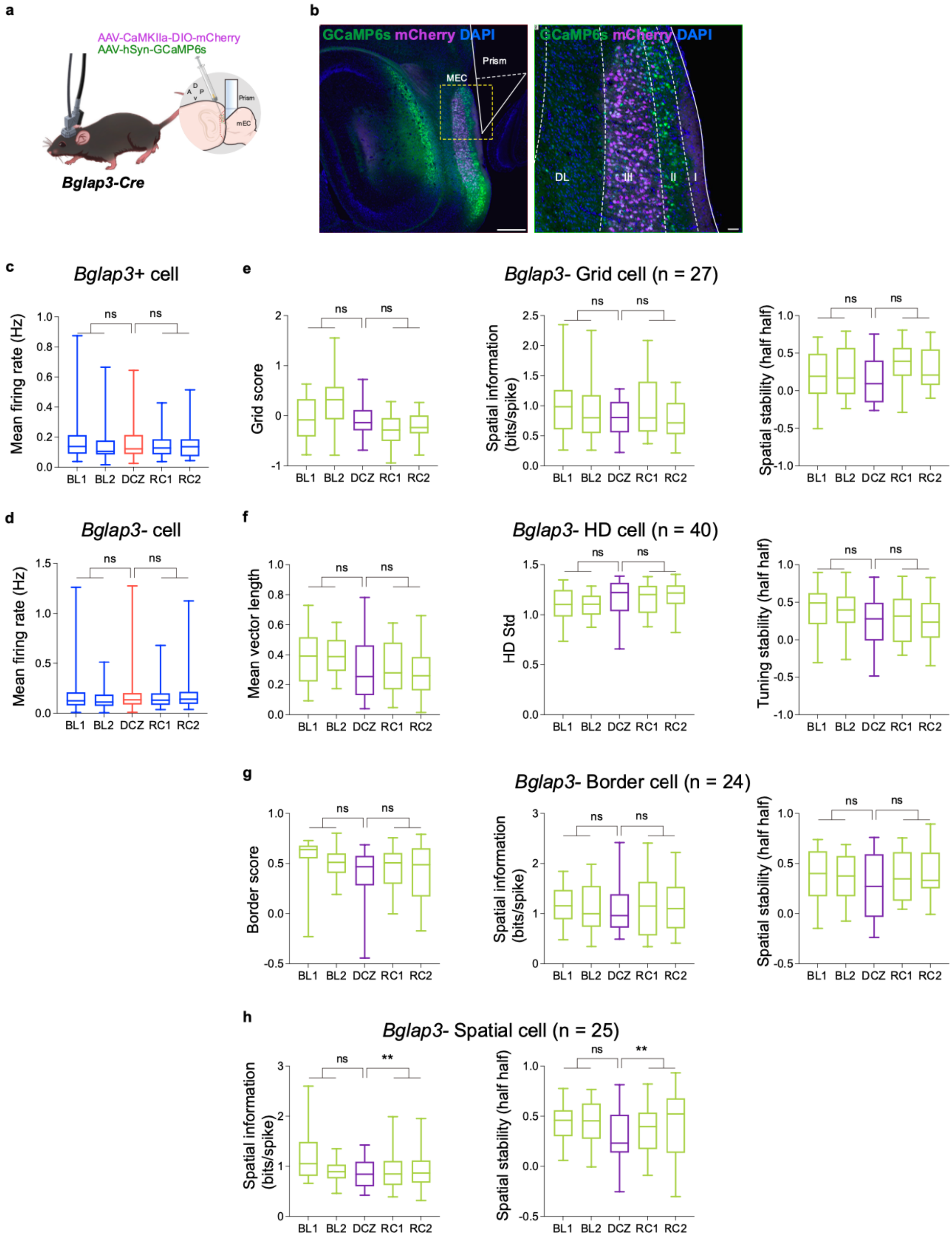

a
AAV-CaMKIIa-DIO-mCherry
AAV-hSyn-GCaMP6s
Prism
mEC
Bglap3-Cre
b
GCaMP6s mCherry DAPI
MEC
Prism
DL
III
II
I
c
Bglap3+ cell
Mean firing rate (Hz)
d
Bglap3- cell
e
Bglap3- Grid cell (n = 27)
Grid score
Spatial information (bits/spike)
Spatial stability (half half)
f
Bglap3- HD cell (n = 40)
Mean vector length
HD Std
Tuning stability (half half)
g
Bglap3- Border cell (n = 24)
Border score
Spatial information (bits/spike)
Spatial stability (half half)
h
Bglap3- Spatial cell (n = 25)
Spatial information (bits/spike)
Spatial stability (half half)
BL1 BL2 DCZ RC1 RC2
ns
**

**Supplementary Fig.26 Two-photon imaging and chemogenetic inhibition of the control group. a** Cartoon of 2-photon imaging in *Bglap3*-Cre mice: miniaturized 2-photon microscope (TPM) in freely behaving mice and prism placement in MEC. **b** GCaMP6s (green) expression in superficial MEC and DIO-mCherry (magenta) in *Bglap3*+ cells; white outline, prism position. Right panel, magnified view of the yellow window (left). LI–LIII, layers 1–3; DL, deep layer. **c-g** Linear mixed-effects ANOVA across five recording sessions; pairwise post-hoc contrasts all P > 0.05 except where noted in **h**. **c** Mean firing rate of *Bglap3*+ neurons (n = 41 cells): P_main = 0.086. **d** Mean firing rate of *Bglap3*- neurons (n = 302 cells): P_main = 0.9296. **e** Grid score, spatial information, and spatial stability of *Bglap3*- grid cells (n = 27 cells): P_main = 0.9050, 0.5104, and 0.1231, respectively. **f** Mean vector length, HD Std, and tuning stability of *Bglap3*- HD cells (n = 40 cells): P_main = 0.2273, 0.2493, and 0.5705, respectively. **g** Border score, spatial information, and spatial stability of *Bglap3*- border cells (n = 24 cells): P_main = 0.4235, 0.3395, and 0.1615, respectively. **h** Spatial information and spatial stability of *Bglap3*- spatial cells (n = 25 cells; P_main = $5.11 \times 10^{-5}$ and 0.0144, respectively). Post hoc: spatial information, BL vs. DCZ P = 0.7035, DCZ vs. RC P = 0.0016; spatial stability, BL vs. DCZ P = 0.1104, DCZ vs. RC P = 0.0041.Data are median with whiskers from min to max; **P < 0.01; ns, non-significant. Source data are provided as a Source Data file. Related to Fig.5.

| **Supplementary Table 1** 1664 Differentially Expressing Genes (DEGs) in MEC during aging. | | | | | | |
|---|---|---|---|---|---|---|
| Up-regulated genes in old MEC | | | | | | |
| | baseMean | log2FoldChange | lfcSE | stat | pvalue | padj |
| Xist | 35.19096693 | 8.412854496 | 1.602102074 | 5.251135138 | 1.51E-07 | 4.17E-06 |
| Smok2b | 17.05226761 | 7.006276082 | 0.872300851 | 8.031949155 | 9.59E-16 | 3.22E-13 |
| Gm16867 | 10.14500837 | 6.61606324 | 0.740392377 | 8.935887847 | 4.04E-19 | 2.63E-16 |
| Gm15446 | 25.55290291 | 5.967707052 | 0.597832575 | 9.982238009 | 1.82E-23 | 2.21E-20 |
| Cst7 | 20.29386418 | 5.834887163 | 0.843104374 | 6.920717466 | 4.49E-12 | 5.60E-10 |
| Defb1 | 52.02601118 | 4.252817363 | 0.574669078 | 7.400463201 | 1.36E-13 | 2.64E-11 |
| H2-Aa | 24.69686355 | 3.942947012 | 0.518545744 | 7.603855698 | 2.87E-14 | 6.58E-12 |
| C4b | 131.9014287 | 3.194774996 | 0.303720292 | 10.51880654 | 7.08E-26 | 1.00E-22 |
| Cd74 | 53.53509275 | 3.13734701 | 0.398629458 | 7.870334084 | 3.54E-15 | 9.83E-13 |
| Cd52 | 58.34399739 | 3.062557238 | 0.274965068 | 11.13798658 | 8.20E-29 | 1.74E-25 |
| Ifit1 | 9.567116844 | 2.90677123 | 0.503328869 | 5.775093398 | 7.69E-09 | 3.23E-07 |
| Lgals3 | 8.69708151 | 2.77784036 | 0.558613479 | 4.972741375 | 6.60E-07 | 1.42E-05 |
| H2-Ab1 | 20.05554282 | 2.745460459 | 0.510007224 | 5.383179548 | 7.32E-08 | 2.24E-06 |
| H2-Eb1 | 14.94595679 | 2.695348135 | 0.346770938 | 7.772704807 | 7.68E-15 | 1.97E-12 |
| Egr2 | 15.40178121 | 2.618480199 | 0.391022366 | 6.696497249 | 2.13E-11 | 2.13E-09 |
| Fosb | 56.52292016 | 2.554809225 | 0.231749196 | 11.02402629 | 2.93E-28 | 5.51E-25 |
| Scnn1a | 44.75162312 | 2.516377644 | 0.366541138 | 6.865198425 | 6.64E-12 | 7.98E-10 |
| Gm11681 | 12.58276192 | 2.430155624 | 0.324103752 | 7.498079267 | 6.48E-14 | 1.39E-11 |
| Sec14l3 | 8.990097085 | 2.408323347 | 0.534705469 | 4.504018542 | 6.67E-06 | 9.58E-05 |
| Eps8l1 | 11.62759592 | 2.403025801 | 0.371313407 | 6.471691456 | 9.69E-11 | 7.75E-09 |
| Lrrc17 | 2010.651858 | 2.363417827 | 0.563548583 | 4.193813803 | 2.74E-05 | 0.000305099 |
| Olfr316 | 10.68289561 | 2.261035211 | 0.402148081 | 5.622394633 | 1.88E-08 | 7.13E-07 |
| Lyz2 | 105.9763087 | 2.231494792 | 0.317865715 | 7.020243738 | 2.21E-12 | 2.98E-10 |
| Fos | 179.3972446 | 2.229781814 | 0.208026081 | 10.71876086 | 8.31E-27 | 1.41E-23 |
| Ifit3b | 13.24985948 | 2.10195621 | 0.345230434 | 6.088559998 | 1.14E-09 | 6.51E-08 |
| Spag6 | 14.99266434 | 2.100366241 | 0.285539826 | 7.355773342 | 1.90E-13 | 3.46E-11 |
| Gstt3 | 10.16714022 | 2.083590415 | 0.355539627 | 5.860360578 | 4.62E-09 | 2.09E-07 |
| Gm34552 | 11.87766212 | 2.06028024 | 0.41512977 | 4.962978777 | 6.94E-07 | 1.47E-05 |
| Serpinb1a | 26.16535999 | 2.056945576 | 0.305186602 | 6.739960293 | 1.58E-11 | 1.64E-09 |
| Gbp2 | 10.56982786 | 2.052626482 | 0.374626556 | 5.479127016 | 4.27E-08 | 1.43E-06 |
| Gm13530 | 18.64604456 | 2.013077432 | 0.269008663 | 7.483318236 | 7.25E-14 | 1.52E-11 |
| Pmaip1 | 10.85099122 | 2.006447199 | 0.315744059 | 6.354663345 | 2.09E-10 | 1.54E-08 |
| Hsf4 | 28.2603832 | 1.981277059 | 0.41044704 | 4.827119863 | 1.39E-06 | 2.62E-05 |
| Gm33651 | 9.921798788 | 1.910272542 | 0.412449874 | 4.63152655 | 3.63E-06 | 5.85E-05 |
| AW112010 | 11.91062461 | 1.907747469 | 0.385385163 | 4.950235904 | 7.41E-07 | 1.55E-05 |
| Wfdc17 | 8.906637594 | 1.898352083 | 0.402967809 | 4.710927376 | 2.47E-06 | 4.26E-05 |
| Psmb9 | 10.34813898 | 1.824886286 | 0.304801128 | 5.987137584 | 2.14E-09 | 1.11E-07 |
| Gpnmb | 14.71038371 | 1.821080494 | 0.351581552 | 5.179681599 | 2.22E-07 | 5.83E-06 |
| Igtp | 12.86579528 | 1.814301006 | 0.322049192 | 5.633614528 | 1.76E-08 | 6.71E-07 |
| Ermn | 89.82518046 | 1.808894678 | 0.427075059 | 4.23554277 | 2.28E-05 | 0.000262557 |
| Trf | 575.661127 | 1.806507865 | 0.3305824 | 5.464622035 | 4.64E-08 | 1.54E-06 |
| Ppp1r14a | 144.6434804 | 1.793325948 | 0.378065254 | 4.743429683 | 2.10E-06 | 3.71E-05 |
| Ifit3 | 25.65220392 | 1.76478593 | 0.29323536 | 6.018325798 | 1.76E-09 | 9.45E-08 |
| Nrp1 | 15.13555844 | 1.750560014 | 0.368815521 | 4.74643803 | 2.07E-06 | 3.68E-05 |
| Npas4 | 57.58041388 | 1.748506382 | 0.410257153 | 4.261976588 | 2.03E-05 | 0.000238523 |
| Mapk13 | 18.74422144 | 1.737866942 | 0.305114137 | 5.695792919 | 1.23E-08 | 4.90E-07 |
| Hapln2 | 40.0565338 | 1.736240416 | 0.554513189 | 3.131107517 | 0.001741484 | 0.00919951 |
| Gjb1 | 30.64724181 | 1.723267743 | 0.314061888 | 5.487032369 | 4.09E-08 | 1.39E-06 |
| Neat1 | 95.33144417 | 1.690004526 | 0.229617901 | 7.360073042 | 1.84E-13 | 3.46E-11 |
| Ass1 | 155.5209485 | 1.678030946 | 0.134860706 | 12.44269732 | 1.53E-35 | 8.66E-32 |
| Bc1 | 129126.854 | 1.675770583 | 0.461317979 | 3.632571586 | 0.000280611 | 0.002061826 |
| Egr1 | 720.7045361 | 1.669964214 | 0.140805505 | 11.86007757 | 1.91E-32 | 6.47E-29 |
| Ccdc36 | 24.44577157 | 1.669174274 | 0.507530087 | 3.288818371 | 0.001006089 | 0.005915442 |
| Pcdhb8 | 12.71343385 | 1.656955224 | 0.328063477 | 5.050715297 | 4.40E-07 | 1.01E-05 |
| Galnt6 | 13.91296204 | 1.639354545 | 0.385909635 | 4.248026994 | 2.16E-05 | 0.000250904 |
| Ankrd22 | 13.85042128 | 1.636509252 | 0.433939365 | 3.771285541 | 0.000162409 | 0.001326742 |
| H2-DMb1 | 9.280714879 | 1.630377996 | 0.330330719 | 4.935593037 | 7.99E-07 | 1.65E-05 |
| Tnnt2 | 53.35243392 | 1.622402358 | 0.189030635 | 8.582748283 | 9.26E-18 | 5.23E-15 |
| Ntn5 | 40.83730131 | 1.61207459 | 0.480558508 | 3.354585473 | 0.00079484 | 0.004857005 |
| H2-K1 | 131.9022446 | 1.611992964 | 0.163140377 | 9.881017779 | 5.03E-23 | 5.69E-20 |
| Arc | 301.3459663 | 1.611652698 | 0.130634298 | 12.33713287 | 5.72E-35 | 2.42E-31 |
| Tmem267 | 69.38845883 | 1.610692755 | 0.235543027 | 6.838210297 | 8.02E-12 | 9.37E-10 |
| Il33 | 152.4496341 | 1.598465984 | 0.21844441 | 7.317495474 | 2.53E-13 | 4.41E-11 |
| Gm40518 | 42.76734459 | 1.597650911 | 0.482887092 | 3.308539276 | 0.00093784 | 0.005574099 |
| Tnfaip6 | 34.93635803 | 1.596440712 | 0.318254692 | 5.016236207 | 5.27E-07 | 1.17E-05 |
| Junb | 273.1790829 | 1.518837283 | 0.151977286 | 9.993843994 | 1.62E-23 | 2.11E-20 |
| Adamtsl4 | 11.70207077 | 1.514628825 | 0.303268917 | 4.99434244 | 5.90E-07 | 1.29E-05 |
| Insc | 10.14356272 | 1.511908253 | 0.384902284 | 3.928031386 | 8.56E-05 | 0.000786431 |
| Pcdhb3 | 27.83199497 | 1.509706105 | 0.217812186 | 6.931228841 | 4.17E-12 | 5.24E-10 |
| Cxcl16 | 10.46281215 | 1.493545653 | 0.336856366 | 4.433775947 | 9.26E-06 | 0.000125569 |

| | | | | | | |
|---|---|---|---|---|---|---|
| Dmp1 | 25.73136093 | 1.480091515 | 0.305113443 | 4.850954786 | 1.23E-06 | 2.38E-05 |
| Mmp14 | 10.579284 | 1.478132086 | 0.43560256 | 3.393304403 | 0.000690549 | 0.004335019 |
| Kazald1 | 30.81965998 | 1.468163659 | 0.223969744 | 6.55518746 | 5.56E-11 | 4.81E-09 |
| Gpr37 | 106.6336583 | 1.450240201 | 0.17070725 | 8.495480997 | 1.97E-17 | 9.83E-15 |
| Plin4 | 12.95765359 | 1.448539102 | 0.410769971 | 3.5263997 | 0.000421251 | 0.002881606 |
| Egr4 | 259.5929926 | 1.446062998 | 0.267149849 | 5.412928369 | 6.20E-08 | 1.98E-06 |
| Aspa | 43.00393553 | 1.444568136 | 0.261613204 | 5.521770743 | 3.36E-08 | 1.18E-06 |
| Gfap | 415.4350741 | 1.427308491 | 0.306702431 | 4.653724094 | 3.26E-06 | 5.34E-05 |
| Pcdhb6 | 11.90847371 | 1.422700851 | 0.267313487 | 5.32221874 | 1.03E-07 | 3.00E-06 |
| S1pr5 | 28.48125302 | 1.418006814 | 0.255728094 | 5.544978621 | 2.94E-08 | 1.05E-06 |
| Oasl2 | 8.194090787 | 1.414091103 | 0.327645403 | 4.315919256 | 1.59E-05 | 0.000195799 |
| Strn3 | 20.62851804 | 1.410727998 | 0.316489363 | 4.457426266 | 8.29E-06 | 0.000114501 |
| Psmb8 | 25.92226367 | 1.40815159 | 0.193466565 | 7.27852686 | 3.37E-13 | 5.72E-11 |
| Cd84 | 11.15692363 | 1.402305166 | 0.283480209 | 4.946748031 | 7.55E-07 | 1.58E-05 |
| Zfp296 | 13.87810934 | 1.400615404 | 0.327419846 | 4.277735203 | 1.89E-05 | 0.000225065 |
| Plekhb1 | 686.2237674 | 1.39713983 | 0.329491897 | 4.240285856 | 2.23E-05 | 0.000258474 |
| Pcdhb9 | 67.92620941 | 1.38274443 | 0.148781829 | 9.293772216 | 1.49E-20 | 1.26E-17 |
| Gstt1 | 80.16678215 | 1.348545709 | 0.28848739 | 4.674539526 | 2.95E-06 | 4.92E-05 |
| Tap2 | 20.63843439 | 1.346983227 | 0.212549977 | 6.337254156 | 2.34E-10 | 1.69E-08 |
| Apol8 | 18.24983212 | 1.327154476 | 0.334423024 | 3.968490145 | 7.23E-05 | 0.000686098 |
| Evi2a | 73.4824001 | 1.318854084 | 0.242908548 | 5.429426405 | 5.65E-08 | 1.82E-06 |
| Opalin | 83.17287677 | 1.318581087 | 0.378702482 | 3.48183904 | 0.000497983 | 0.003306645 |
| Ccn1 | 21.47134078 | 1.314383361 | 0.253638333 | 5.182116382 | 2.19E-07 | 5.77E-06 |
| Tmem88b | 72.56147701 | 1.310258688 | 0.301979842 | 4.338894537 | 1.43E-05 | 0.000178748 |
| Parp14 | 8.547955896 | 1.308287882 | 0.327843094 | 3.990591551 | 6.59E-05 | 0.000635505 |
| Gpat3 | 12.08127071 | 1.29740044 | 0.283549484 | 4.575569743 | 4.75E-06 | 7.23E-05 |
| Exoc3l4 | 9.518689987 | 1.292008942 | 0.300726149 | 4.296297297 | 1.74E-05 | 0.000209983 |
| Lag3 | 8.51437158 | 1.291158179 | 0.341368445 | 3.782300908 | 0.000155385 | 0.001279231 |
| Gm11837 | 13.35996851 | 1.288169706 | 0.256761118 | 5.016996786 | 5.25E-07 | 1.17E-05 |
| Pls1 | 30.78697349 | 1.287715003 | 0.191122097 | 6.737656323 | 1.61E-11 | 1.65E-09 |
| Glt8d2 | 40.99658155 | 1.282746484 | 0.284355388 | 4.511067972 | 6.45E-06 | 9.32E-05 |
| Zfp367 | 52.06515322 | 1.28162135 | 0.145958875 | 8.780701763 | 1.62E-18 | 9.84E-16 |
| Espn | 18.81473867 | 1.280425997 | 0.320030298 | 4.000952423 | 6.31E-05 | 0.000614052 |
| Cd82 | 36.49453362 | 1.267373488 | 0.21466041 | 5.904085847 | 3.55E-09 | 1.70E-07 |
| Ccno | 9.395321794 | 1.266376908 | 0.310131783 | 4.083350937 | 4.44E-05 | 0.000457151 |
| Efhd1 | 51.23290621 | 1.262096822 | 0.213085267 | 5.922966152 | 3.16E-09 | 1.54E-07 |
| Prr5l | 15.13208936 | 1.261464181 | 0.28616999 | 4.408093885 | 1.04E-05 | 0.000138103 |
| Tmem254a | 17.73130791 | 1.260047087 | 0.274997587 | 4.58202961 | 4.60E-06 | 7.04E-05 |
| Gstp1 | 186.1374358 | 1.250829646 | 0.324859493 | 3.850371227 | 0.000117939 | 0.001025222 |
| Ifi27 | 200.0263659 | 1.249999938 | 0.183871103 | 6.798240295 | 1.06E-11 | 1.18E-09 |
| Evpl | 12.59937842 | 1.248058041 | 0.253892616 | 4.915692551 | 8.85E-07 | 1.81E-05 |
| Ccl6 | 19.44171359 | 1.247220363 | 0.308274994 | 4.045804517 | 5.21E-05 | 0.000522888 |
| Depp1 | 18.51398107 | 1.246377484 | 0.265956249 | 4.686400455 | 2.78E-06 | 4.69E-05 |
| Hist1h2be | 8.112898359 | 1.238515658 | 0.324836479 | 3.812735754 | 0.000137437 | 0.001158476 |
| Lgals3bp | 68.47207509 | 1.236977579 | 0.167229296 | 7.396895217 | 1.39E-13 | 2.69E-11 |
| Irgm2 | 8.49406956 | 1.233167379 | 0.309281602 | 3.987199268 | 6.69E-05 | 0.000642829 |
| Smim1 | 45.0028512 | 1.23035469 | 0.164413332 | 7.483302427 | 7.25E-14 | 1.52E-11 |
| Arsg | 35.53301062 | 1.228496081 | 0.242556119 | 5.064791127 | 4.09E-07 | 9.55E-06 |
| Itgb2 | 21.51659315 | 1.226340762 | 0.257775139 | 4.757405097 | 1.96E-06 | 3.51E-05 |
| Abca8a | 23.44181913 | 1.225612556 | 0.299626196 | 4.090471974 | 4.30E-05 | 0.000446321 |
| Sorbs1 | 11.06454243 | 1.225530735 | 0.317692004 | 3.857606482 | 0.000114503 | 0.000998938 |
| Bhlhe40 | 270.68024 | 1.224430286 | 0.107033958 | 11.43964321 | 2.65E-30 | 6.42E-27 |
| H2-T22 | 65.45676294 | 1.223784827 | 0.186315857 | 6.568334271 | 5.09E-11 | 4.47E-09 |
| Adssl1 | 67.85928185 | 1.220555901 | 0.215115778 | 5.673948762 | 1.40E-08 | 5.44E-07 |
| Litaf | 67.46436564 | 1.2105337 | 0.198720037 | 6.091653949 | 1.12E-09 | 6.42E-08 |
| B2m | 701.2554914 | 1.208285359 | 0.143071002 | 8.445354703 | 3.03E-17 | 1.43E-14 |
| Zfp791 | 11.020655 | 1.204808287 | 0.278298999 | 4.329186562 | 1.50E-05 | 0.000186127 |
| Bcl2a1b | 9.80330071 | 1.204183384 | 0.382583469 | 3.147505003 | 0.001646703 | 0.00878881 |
| H2-M3 | 10.01643281 | 1.203679615 | 0.286187739 | 4.205909097 | 2.60E-05 | 0.000291911 |
| Endou | 8.162506227 | 1.201731451 | 0.323780428 | 3.711562986 | 0.000205983 | 0.001605345 |
| Pde8a | 27.2028666 | 1.200418511 | 0.197199426 | 6.087332686 | 1.15E-09 | 6.51E-08 |
| Plekhh1 | 35.87074561 | 1.192732962 | 0.36998081 | 3.223769801 | 0.001265151 | 0.007108242 |
| Herc6 | 9.667425681 | 1.184667335 | 0.309282345 | 3.830374917 | 0.000127948 | 0.001096486 |
| Nr4a1 | 346.5300196 | 1.18292938 | 0.146965143 | 8.049047258 | 8.34E-16 | 2.89E-13 |
| Erbb3 | 13.05198256 | 1.182707743 | 0.322027674 | 3.672689763 | 0.000240011 | 0.00181626 |
| Serpina3n | 65.28329128 | 1.179680145 | 0.306972874 | 3.842945893 | 0.000121566 | 0.001049459 |
| Utp14b | 24.01037314 | 1.176469577 | 0.196837195 | 5.976866187 | 2.27E-09 | 1.17E-07 |
| Asb16 | 13.09168336 | 1.174591281 | 0.271203138 | 4.331038677 | 1.48E-05 | 0.000184839 |
| C1ql3 | 111.1700887 | 1.173431137 | 0.40769164 | 2.878232031 | 0.003999109 | 0.017985909 |
| Zc3hav1 | 16.29347054 | 1.171394465 | 0.229665005 | 5.100448209 | 3.39E-07 | 8.25E-06 |
| Itgb4 | 16.61432699 | 1.171239591 | 0.279588008 | 4.189162466 | 2.80E-05 | 0.000310605 |
| Zfp503 | 39.20110381 | 1.169616245 | 0.198727665 | 5.885522999 | 3.97E-09 | 1.84E-07 |
| Prelp | 67.09700616 | 1.161827134 | 0.194349517 | 5.978029434 | 2.26E-09 | 1.17E-07 |

| | | | | | | |
|---|---|---|---|---|---|---|
| Prr18 | 83.1787052 | 1.156206485 | 0.18534109 | 6.238263106 | 4.42E-10 | 2.92E-08 |
| Acat3 | 12.8580035 | 1.154869684 | 0.264755082 | 4.36203028 | 1.29E-05 | 0.000163497 |
| Adm | 11.05095052 | 1.148982547 | 0.293596366 | 3.913476729 | 9.10E-05 | 0.000826103 |
| Arhgap15 | 32.91972487 | 1.148464536 | 0.174550933 | 6.579538207 | 4.72E-11 | 4.19E-09 |
| Mobp | 1132.770024 | 1.144084255 | 0.334871276 | 3.416489665 | 0.000634341 | 0.004053038 |
| Cox8b | 12.15357786 | 1.13723577 | 0.339687073 | 3.347892401 | 0.000814286 | 0.004952625 |
| Ccl12 | 8.057099546 | 1.135533391 | 0.372565957 | 3.047872116 | 0.00230468 | 0.011517284 |
| Nkx6-2 | 46.98117753 | 1.131570877 | 0.33401384 | 3.387796375 | 0.000704566 | 0.004405419 |
| Tagln | 22.91399964 | 1.131305523 | 0.326237585 | 3.467735098 | 0.000524864 | 0.003447104 |
| Sfrp5 | 8.549703934 | 1.131058246 | 0.38471249 | 2.940009167 | 0.003282025 | 0.015296566 |
| Ighm | 84.91973576 | 1.12783102 | 0.157932339 | 7.141229144 | 9.25E-13 | 1.40E-10 |
| Pcdhb13 | 19.57047912 | 1.125617706 | 0.205189391 | 5.485750019 | 4.12E-08 | 1.39E-06 |
| Hba-a1 | 1117.318277 | 1.125391932 | 0.347741261 | 3.236291048 | 0.001210938 | 0.006849055 |
| Plp1 | 2869.851743 | 1.123217776 | 0.30246658 | 3.713526888 | 0.000204391 | 0.001598075 |
| Klhdc7a | 12.21006682 | 1.121781458 | 0.31365343 | 3.576499887 | 0.000348225 | 0.002471847 |
| Ifi27l2a | 34.89158532 | 1.119058628 | 0.254985969 | 4.388706686 | 1.14E-05 | 0.000148112 |
| AA467197 | 8.31674038 | 1.117030875 | 0.29218109 | 3.823077243 | 0.000131796 | 0.001120964 |
| A930004D18Rik | 8.073498914 | 1.115211715 | 0.321802963 | 3.465511017 | 0.000529225 | 0.003469022 |
| Lrrc38 | 11.0152353 | 1.112431139 | 0.282302974 | 3.94055763 | 8.13E-05 | 0.000750947 |
| Mreg | 8.474259481 | 1.11165439 | 0.358602237 | 3.099965013 | 0.001935435 | 0.009990122 |
| Rapgef3 | 57.48960904 | 1.107158188 | 0.151104989 | 7.327078997 | 2.35E-13 | 4.20E-11 |
| Crb3 | 18.06899181 | 1.106736452 | 0.285673867 | 3.874125631 | 0.000107008 | 0.00094523 |
| Macrod1 | 49.46452314 | 1.102374235 | 0.239127132 | 4.6099923 | 4.03E-06 | 6.36E-05 |
| Mafa | 12.22653206 | 1.098067373 | 0.275583498 | 3.984517877 | 6.76E-05 | 0.000648659 |
| Aif1l | 18.96857063 | 1.084665048 | 0.223684808 | 4.84907785 | 1.24E-06 | 2.40E-05 |
| Anxa4 | 21.44889483 | 1.084151351 | 0.243017447 | 4.461207886 | 8.15E-06 | 0.000113051 |
| Batf3 | 15.68006564 | 1.07360231 | 0.434755818 | 2.469437476 | 0.013532566 | 0.047571658 |
| Scx | 11.86884753 | 1.072785711 | 0.264000564 | 4.063573562 | 4.83E-05 | 0.000493192 |
| AW495222 | 26.67140006 | 1.071281206 | 0.185102407 | 5.787505544 | 7.14E-09 | 3.04E-07 |
| Kank4 | 31.98171127 | 1.070336107 | 0.23818082 | 4.493796385 | 7.00E-06 | 9.94E-05 |
| mt-Nd6 | 144.3040523 | 1.068188077 | 0.385263577 | 2.772616312 | 0.005560764 | 0.023291454 |
| Tmem63a | 49.98772312 | 1.066319842 | 0.34225865 | 3.115538035 | 0.001836097 | 0.009582412 |
| Trim59 | 34.9796185 | 1.06541448 | 0.220054365 | 4.841596677 | 1.29E-06 | 2.47E-05 |
| Hapln4 | 252.0165228 | 1.065106919 | 0.35432465 | 3.006019813 | 0.002646918 | 0.012893074 |
| Tatdn1 | 150.1392345 | 1.061770684 | 0.261118931 | 4.06623404 | 4.78E-05 | 0.00048793 |
| Thbs4 | 9.186173989 | 1.061437362 | 0.339907726 | 3.122722085 | 0.001791869 | 0.009406617 |
| Hspa1b | 25.38243466 | 1.057684308 | 0.252637673 | 4.186566057 | 2.83E-05 | 0.000313972 |
| Pcdhb14 | 30.82757614 | 1.05759625 | 0.156473176 | 6.758961991 | 1.39E-11 | 1.50E-09 |
| Hipk2 | 9.283879184 | 1.056351287 | 0.370475392 | 2.85133995 | 0.00435354 | 0.019258051 |
| 2900041M22Rik | 19.7363301 | 1.054768338 | 0.240797507 | 4.380312525 | 1.19E-05 | 0.000152186 |
| Prss16 | 8.706650035 | 1.050040368 | 0.356590354 | 2.944668463 | 0.003233009 | 0.015134696 |
| Apod | 742.5755866 | 1.0451512 | 0.254774381 | 4.102261752 | 4.09E-05 | 0.000427615 |
| Chil1 | 28.42911158 | 1.038772417 | 0.200847739 | 5.171939796 | 2.32E-07 | 6.03E-06 |
| Pdlim2 | 82.37661372 | 1.033932393 | 0.323744466 | 3.193668159 | 0.001404775 | 0.007708754 |
| Dkkl1 | 86.15387249 | 1.033617588 | 0.250385994 | 4.128096665 | 3.66E-05 | 0.000389222 |
| Gm44053 | 8.508717752 | 1.033122936 | 0.287359027 | 3.59523397 | 0.0003241 | 0.002320026 |
| Deptor | 74.12844294 | 1.028241146 | 0.213452895 | 4.817180596 | 1.46E-06 | 2.72E-05 |
| Irgm1 | 38.44333055 | 1.028036181 | 0.190198651 | 5.405065564 | 6.48E-08 | 2.03E-06 |
| Mafb | 87.10818194 | 1.021840798 | 0.126190608 | 8.097597889 | 5.61E-16 | 2.02E-13 |
| Mpeg1 | 64.85891098 | 1.020328195 | 0.174099639 | 5.86059914 | 4.61E-09 | 2.09E-07 |
| C3ar1 | 10.02488619 | 1.015821335 | 0.406927533 | 2.496319989 | 0.012548934 | 0.044763675 |
| Rgs16 | 52.53908854 | 1.007279773 | 0.267661691 | 3.763257157 | 0.000167715 | 0.001361556 |
| Glra3 | 41.47786057 | 1.005059787 | 0.36322485 | 2.767045779 | 0.005656681 | 0.023611525 |
| Jade2 | 131.6760929 | 1.002404171 | 0.204671772 | 4.897618072 | 9.70E-07 | 1.95E-05 |
| Pdk4 | 9.904526318 | 1.002345256 | 0.287692007 | 3.484091425 | 0.000493811 | 0.00328516 |
| Ctss | 342.8403048 | 1.001557981 | 0.148351708 | 6.751239963 | 1.47E-11 | 1.54E-09 |
| Pcdhb11 | 16.11291197 | 0.999732305 | 0.264039325 | 3.786300793 | 0.000152907 | 0.001263119 |
| Rnf213 | 12.17127188 | 0.99907949 | 0.246017395 | 4.061011584 | 4.89E-05 | 0.000497439 |
| Olfr287 | 9.81345511 | 0.998858654 | 0.330571372 | 3.021612693 | 0.00251432 | 0.012396813 |
| Cfap58 | 26.19889854 | 0.992608638 | 0.236134224 | 4.203578035 | 2.63E-05 | 0.000294544 |
| Plekhg3 | 20.8859692 | 0.990851613 | 0.208813781 | 4.745144727 | 2.08E-06 | 3.70E-05 |
| Ninj2 | 11.55509262 | 0.984076756 | 0.30440701 | 3.232766412 | 0.001225978 | 0.006920262 |
| Plaat1 | 55.77757624 | 0.981838844 | 0.130906229 | 7.500321827 | 6.37E-14 | 1.38E-11 |
| Tent5a | 13.32723881 | 0.978738992 | 0.250978295 | 3.899695759 | 9.63E-05 | 0.000865189 |
| Hba-a2 | 1081.462374 | 0.978085479 | 0.320158912 | 3.055000008 | 0.002250605 | 0.011300886 |
| Anxa11 | 47.26875217 | 0.977761299 | 0.175346045 | 5.576181078 | 2.46E-08 | 8.96E-07 |
| Tapbpl | 20.53493232 | 0.974588409 | 0.202579975 | 4.810882257 | 1.50E-06 | 2.79E-05 |
| Gm14419 | 31.6311546 | 0.971763156 | 0.173281131 | 5.608014848 | 2.05E-08 | 7.59E-07 |
| Tap1 | 10.64226284 | 0.970601425 | 0.270267269 | 3.591265147 | 0.000329077 | 0.002352669 |
| Nr4a3 | 152.2970305 | 0.962537735 | 0.159592152 | 6.031234779 | 1.63E-09 | 8.81E-08 |
| Pold1 | 37.70779439 | 0.959503918 | 0.201519307 | 4.761349841 | 1.92E-06 | 3.45E-05 |
| Ube2l6 | 10.77309478 | 0.958184073 | 0.292196638 | 3.279244001 | 0.001040856 | 0.00607979 |
| Ctxn3 | 24.91981411 | 0.954705176 | 0.31932691 | 2.989742315 | 0.002792129 | 0.013495688 |

| | | | | | | |
|---|---|---|---|---|---|---|
| Tuft1 | 12.78530358 | 0.954334943 | 0.240983701 | 3.960163862 | 7.49E-05 | 0.000706512 |
| Hspa1a | 57.43574077 | 0.952707611 | 0.135937956 | 7.008400283 | 2.41E-12 | 3.17E-10 |
| Pkib | 70.25272801 | 0.951833109 | 0.13408284 | 7.098843589 | 1.26E-12 | 1.79E-10 |
| B230206L02Rik | 15.93057672 | 0.948762649 | 0.208311514 | 4.554537724 | 5.25E-06 | 7.86E-05 |
| Dgkg | 13.84023704 | 0.946074072 | 0.370593956 | 2.552858885 | 0.010684277 | 0.039558095 |
| Sncg | 30.81280067 | 0.944963229 | 0.231185568 | 4.087466354 | 4.36E-05 | 0.000450487 |
| Gm13031 | 9.217472385 | 0.944654257 | 0.31787141 | 2.971812584 | 0.002960473 | 0.014127193 |
| Cyp27a1 | 8.343614481 | 0.94313655 | 0.351187084 | 2.6855673 | 0.007240679 | 0.028750703 |
| Fcgr2b | 15.69396847 | 0.942288779 | 0.287120467 | 3.281858624 | 0.001031253 | 0.006027851 |
| 3110070M22Rik | 9.775279533 | 0.934141724 | 0.343609311 | 2.718615867 | 0.006555569 | 0.026590919 |
| Tmem98 | 28.76723934 | 0.933212954 | 0.197598365 | 4.722776709 | 2.33E-06 | 4.05E-05 |
| Plaat3 | 204.4109947 | 0.927374908 | 0.17627561 | 5.260937179 | 1.43E-07 | 3.98E-06 |
| Hmox1 | 13.54104032 | 0.926122882 | 0.24589167 | 3.766385756 | 0.000165628 | 0.001347843 |
| Fam122a | 8.040331163 | 0.923925331 | 0.314416576 | 2.938538875 | 0.003297633 | 0.015343912 |
| Slc5a5 | 202.8259044 | 0.923544755 | 0.16907121 | 5.462460212 | 4.70E-08 | 1.55E-06 |
| 2900052N01Rik | 278.6055712 | 0.921215972 | 0.243450086 | 3.784003473 | 0.000154326 | 0.001271743 |
| Dusp1 | 199.6575552 | 0.918751216 | 0.110252488 | 8.333156345 | 7.87E-17 | 3.25E-14 |
| Crhbp | 113.1963583 | 0.916571219 | 0.114488109 | 8.005820224 | 1.19E-15 | 3.87E-13 |
| Efcab10 | 72.7371449 | 0.909810572 | 0.118931002 | 7.649902534 | 2.01E-14 | 4.74E-12 |
| Tspo | 45.52107782 | 0.909123691 | 0.173751357 | 5.232325708 | 1.67E-07 | 4.58E-06 |
| Smco4 | 14.53832896 | 0.905055699 | 0.224786475 | 4.026290713 | 5.67E-05 | 0.000563016 |
| Mansc1 | 24.52458131 | 0.904804797 | 0.179532036 | 5.03979579 | 4.66E-07 | 1.06E-05 |
| Zdhhc1 | 47.55881314 | 0.903359383 | 0.173874057 | 5.195481148 | 2.04E-07 | 5.43E-06 |
| Alox8 | 19.39980558 | 0.901833524 | 0.201677727 | 4.47165652 | 7.76E-06 | 0.000108375 |
| Smco3 | 21.99592852 | 0.901701173 | 0.235996748 | 3.820820327 | 0.000133009 | 0.001127314 |
| Kcnk13 | 13.60775437 | 0.898120928 | 0.267001009 | 3.363736084 | 0.00076895 | 0.004722637 |
| Rps27 | 5130.411175 | 0.897047507 | 0.163884727 | 5.473649206 | 4.41E-08 | 1.47E-06 |
| B230209E15Rik | 17.68179614 | 0.891632137 | 0.295889088 | 3.013399866 | 0.002583383 | 0.012656336 |
| Ly86 | 84.06663837 | 0.890568106 | 0.184098473 | 4.837455151 | 1.32E-06 | 2.50E-05 |
| Srek1ip1 | 93.64993672 | 0.89014774 | 0.161528854 | 5.510766134 | 3.57E-08 | 1.24E-06 |
| Tmem243 | 40.86464739 | 0.886881839 | 0.191695381 | 4.62651649 | 3.72E-06 | 5.98E-05 |
| Qdpr | 840.6019405 | 0.880970456 | 0.187271221 | 4.704248998 | 2.55E-06 | 4.38E-05 |
| Efemp1 | 44.55721456 | 0.880107866 | 0.201586731 | 4.365901765 | 1.27E-05 | 0.000161211 |
| Plin3 | 40.65257073 | 0.879456224 | 0.220511456 | 3.988256396 | 6.66E-05 | 0.000640699 |
| Cmip | 55.56935917 | 0.878749474 | 0.339956905 | 2.584884911 | 0.009741151 | 0.03671171 |
| Ak8 | 11.11555062 | 0.876895765 | 0.270886656 | 3.237131639 | 0.001207377 | 0.006833472 |
| Ankrd35 | 209.8222927 | 0.876019306 | 0.266972428 | 3.281310025 | 0.001033261 | 0.006037507 |
| Nectin4 | 9.640267144 | 0.875604542 | 0.315954448 | 2.771299934 | 0.005583297 | 0.023356977 |
| Pigz | 42.75137354 | 0.87438969 | 0.184865107 | 4.729879559 | 2.25E-06 | 3.93E-05 |
| Gstm7 | 42.23506074 | 0.871638108 | 0.174795746 | 4.986609382 | 6.14E-07 | 1.33E-05 |
| Lzts3 | 132.6540902 | 0.870919343 | 0.197228332 | 4.415792273 | 1.01E-05 | 0.000133906 |
| Anln | 41.05084575 | 0.86893701 | 0.197825361 | 4.39244497 | 1.12E-05 | 0.000146148 |
| Bche | 24.78848334 | 0.867912983 | 0.288679233 | 3.006496087 | 0.002642775 | 0.012885206 |
| Sowahb | 60.76872515 | 0.866660881 | 0.145786662 | 5.944719956 | 2.77E-09 | 1.38E-07 |
| Wwox | 161.8022955 | 0.866485753 | 0.191730362 | 4.519293369 | 6.20E-06 | 9.04E-05 |
| Myo1d | 15.80172496 | 0.86212401 | 0.232379574 | 3.709981887 | 0.000207274 | 0.001611699 |
| Parp4 | 10.03198198 | 0.861464355 | 0.286759379 | 3.004136629 | 0.002663357 | 0.012965699 |
| Ggact | 84.38400187 | 0.861330436 | 0.166743111 | 5.165613335 | 2.40E-07 | 6.19E-06 |
| Itih3 | 111.7096986 | 0.859716065 | 0.16638912 | 5.166900724 | 2.38E-07 | 6.17E-06 |
| Cryab | 814.1949876 | 0.857289789 | 0.253092588 | 3.387257595 | 0.000705951 | 0.004410826 |
| Cbx7 | 119.2587477 | 0.856461381 | 0.286722131 | 2.987078044 | 0.002816578 | 0.013582878 |
| Gm973 | 10.35855529 | 0.855922527 | 0.267640464 | 3.198031093 | 0.001383694 | 0.00761773 |
| Ppl | 14.60463244 | 0.855606897 | 0.246380258 | 3.472708828 | 0.000515234 | 0.00339174 |
| Rnf207 | 22.5216259 | 0.854477627 | 0.240662174 | 3.550527326 | 0.00038446 | 0.002676918 |
| Gadd45b | 115.1686262 | 0.85355317 | 0.129751843 | 6.578351055 | 4.76E-11 | 4.20E-09 |
| 9330179D12Rik | 18.7870269 | 0.851475275 | 0.24209933 | 3.517049282 | 0.000436373 | 0.00296828 |
| H2-D1 | 205.3213251 | 0.85105515 | 0.125964474 | 6.756310907 | 1.42E-11 | 1.52E-09 |
| Sgk1 | 254.684038 | 0.850823863 | 0.140995044 | 6.034423898 | 1.60E-09 | 8.67E-08 |
| Arsj | 14.26202633 | 0.84612976 | 0.320167795 | 2.642769737 | 0.008223094 | 0.031962775 |
| Spg20 | 66.35468329 | 0.840384571 | 0.179441919 | 4.683323579 | 2.82E-06 | 4.76E-05 |
| Rbmx | 87.42004772 | 0.839814911 | 0.107328611 | 7.824706759 | 5.09E-15 | 1.37E-12 |
| Prtn3 | 26.87236956 | 0.837804426 | 0.25366425 | 3.302808446 | 0.000957218 | 0.005671372 |
| Ar | 39.23264323 | 0.834461949 | 0.21083587 | 3.957874675 | 7.56E-05 | 0.000712127 |
| Mcam | 19.92774069 | 0.832895122 | 0.226517251 | 3.676961108 | 0.000236029 | 0.001790926 |
| mt-Nd2 | 51841.70991 | 0.832799391 | 0.21345016 | 3.901610527 | 9.56E-05 | 0.00085974 |
| Gm36251 | 18.37084411 | 0.830228825 | 0.314628672 | 2.638757679 | 0.008321044 | 0.032284278 |
| Pdlim1 | 28.40062681 | 0.828516021 | 0.24100986 | 3.437685163 | 0.000586709 | 0.003787247 |
| Pir | 29.63258042 | 0.828086448 | 0.169459339 | 4.886638013 | 1.03E-06 | 2.03E-05 |
| Chrm2 | 10.18754464 | 0.827790478 | 0.268937633 | 3.078001653 | 0.002083938 | 0.010604871 |
| Dipk1c | 11.00855742 | 0.8277353 | 0.268171301 | 3.086591655 | 0.002024655 | 0.010362298 |
| Atf5 | 59.27257421 | 0.823620138 | 0.160115382 | 5.143916385 | 2.69E-07 | 6.81E-06 |
| Fads6 | 27.60146306 | 0.823334675 | 0.221175568 | 3.722538978 | 0.000197229 | 0.001551386 |
| Akain1 | 45.0735502 | 0.819398958 | 0.275585828 | 2.973298609 | 0.002946177 | 0.014075716 |

| | | | | | | |
|---|---|---|---|---|---|---|
| Nfil3 | 52.00438919 | 0.816814972 | 0.136413345 | 5.987793743 | 2.13E-09 | 1.11E-07 |
| Ier5 | 371.3118518 | 0.816740245 | 0.139253454 | 5.865134543 | 4.49E-09 | 2.04E-07 |
| Medag | 19.2404075 | 0.816114013 | 0.221586731 | 3.683045501 | 0.000230464 | 0.001758144 |
| Prune2 | 222.0066628 | 0.813146358 | 0.095220659 | 8.539600196 | 1.35E-17 | 7.13E-15 |
| Gpld1 | 101.5333508 | 0.812372849 | 0.129626778 | 6.26701413 | 3.68E-10 | 2.52E-08 |
| Rtp4 | 12.93146583 | 0.811978672 | 0.25093861 | 3.235766194 | 0.001213167 | 0.006859371 |
| Pi4kb | 40.09439695 | 0.81047434 | 0.199885387 | 4.054695293 | 5.02E-05 | 0.000507718 |
| Lactb2 | 26.31940077 | 0.809176552 | 0.167080541 | 4.843032853 | 1.28E-06 | 2.46E-05 |
| Rapgef4 | 630.8421653 | 0.808128177 | 0.130409076 | 6.196870637 | 5.76E-10 | 3.64E-08 |
| Arhgef10 | 54.74686931 | 0.805900098 | 0.147160575 | 5.476331526 | 4.34E-08 | 1.45E-06 |
| Capg | 10.7837581 | 0.805255533 | 0.283041517 | 2.845008536 | 0.004441023 | 0.019572612 |
| Apold1 | 18.03123469 | 0.803413295 | 0.225738186 | 3.559049135 | 0.0003722 | 0.002612489 |
| Ddo | 14.93085329 | 0.802132315 | 0.27769997 | 2.888485421 | 0.00387102 | 0.017488714 |
| Gpc3 | 15.02482509 | 0.800937795 | 0.285881783 | 2.801639851 | 0.00508436 | 0.021752899 |
| 2010001A14Rik | 36.68328578 | 0.798280479 | 0.193077915 | 4.134499167 | 3.56E-05 | 0.00037996 |
| Aatf | 29.07525734 | 0.79608538 | 0.202682059 | 3.927754547 | 8.57E-05 | 0.00078691 |
| 1700047M11Rik | 20.71500717 | 0.794055345 | 0.267253095 | 2.971173618 | 0.00296664 | 0.014141597 |
| AC160336.1 | 16.94813466 | 0.794008922 | 0.265559399 | 2.989948478 | 0.002790245 | 0.013494278 |
| Slc25a34 | 10.75338812 | 0.793078651 | 0.306203754 | 2.590035691 | 0.009596598 | 0.036237901 |
| Mettl7a1 | 98.76741822 | 0.79264414 | 0.117220871 | 6.761971088 | 1.36E-11 | 1.48E-09 |
| Mthfd2 | 30.77638531 | 0.791213085 | 0.148591845 | 5.324740978 | 1.01E-07 | 2.96E-06 |
| Syt2 | 104.3839299 | 0.790935552 | 0.235216641 | 3.36258332 | 0.000772168 | 0.004740682 |
| Tmem88 | 28.74587666 | 0.789568566 | 0.182954454 | 4.315656443 | 1.59E-05 | 0.00019589 |
| Ankrd6 | 75.78874908 | 0.788943146 | 0.14461522 | 5.455464121 | 4.88E-08 | 1.60E-06 |
| Gngt2 | 26.74668455 | 0.787734023 | 0.224418457 | 3.510112464 | 0.000447917 | 0.003030996 |
| A230056P14Rik | 15.30107436 | 0.787515428 | 0.213172384 | 3.694265708 | 0.000220523 | 0.001691442 |
| 1700112D23Rik | 12.58060899 | 0.787188612 | 0.260014575 | 3.02747879 | 0.00246603 | 0.012197747 |
| Nap1l2 | 363.1451915 | 0.786707558 | 0.096180163 | 8.179519872 | 2.85E-16 | 1.05E-13 |
| Maob | 56.53125616 | 0.785657062 | 0.114590869 | 6.856192534 | 7.07E-12 | 8.32E-10 |
| Rdm1 | 34.54254744 | 0.785117801 | 0.211488565 | 3.712341607 | 0.000205351 | 0.001602921 |
| Ggta1 | 10.05778601 | 0.783376955 | 0.266512669 | 2.939361036 | 0.003288897 | 0.015320169 |
| Ctsz | 222.6344112 | 0.780221513 | 0.125596433 | 6.21213116 | 5.23E-10 | 3.37E-08 |
| Myrf | 61.82142116 | 0.777677459 | 0.193853713 | 4.011671727 | 6.03E-05 | 0.000593485 |
| mt-Atp8 | 1533.482387 | 0.776410681 | 0.260249035 | 2.983337401 | 0.002851235 | 0.013714893 |
| Zar1l | 14.58188232 | 0.774005921 | 0.23787482 | 3.253837124 | 0.001138575 | 0.006535724 |
| H2-T23 | 51.14898354 | 0.773380192 | 0.197690743 | 3.912070855 | 9.15E-05 | 0.000828883 |
| Lap3 | 158.87413 | 0.772590116 | 0.109208238 | 7.074467343 | 1.50E-12 | 2.05E-10 |
| Zfp84 | 9.753512955 | 0.771217566 | 0.278482535 | 2.769357024 | 0.005616705 | 0.023461993 |
| Ehd4 | 34.28286968 | 0.769678956 | 0.153724197 | 5.006882268 | 5.53E-07 | 1.22E-05 |
| Tmc6 | 35.46743943 | 0.768429326 | 0.198680448 | 3.867664551 | 0.000109883 | 0.00096609 |
| Scn1a | 127.3140416 | 0.768128496 | 0.105555948 | 7.276979761 | 3.41E-13 | 5.73E-11 |
| Padi2 | 54.39848086 | 0.767162981 | 0.19386338 | 3.957235138 | 7.58E-05 | 0.0007128 |
| Anxa5 | 269.5995376 | 0.766373406 | 0.091034921 | 8.418455241 | 3.81E-17 | 1.70E-14 |
| Rffl | 22.03521527 | 0.766346196 | 0.184144606 | 4.161654319 | 3.16E-05 | 0.000345305 |
| Haus7 | 34.01773758 | 0.76491301 | 0.146260251 | 5.229807839 | 1.70E-07 | 4.63E-06 |
| Spty2d1 | 43.69114093 | 0.762962362 | 0.1552839 | 4.913338482 | 8.95E-07 | 1.82E-05 |
| Mbp | 5364.241569 | 0.761973815 | 0.242823172 | 3.137978184 | 0.001701175 | 0.00903403 |
| Gatm | 262.6133862 | 0.761019211 | 0.122625403 | 6.206048581 | 5.43E-10 | 3.46E-08 |
| Csrnp1 | 22.25761398 | 0.760340548 | 0.171028983 | 4.445682439 | 8.76E-06 | 0.000120111 |
| Plvap | 86.83172105 | 0.760103092 | 0.161592257 | 4.703833632 | 2.55E-06 | 4.38E-05 |
| Mtrf1 | 29.10175197 | 0.759631699 | 0.161421386 | 4.705892571 | 2.53E-06 | 4.35E-05 |
| Nat8f5 | 10.72415841 | 0.758464167 | 0.273406199 | 2.774129373 | 0.005534966 | 0.023233799 |
| Dusp14 | 260.6633637 | 0.758306035 | 0.127067668 | 5.967733948 | 2.41E-09 | 1.22E-07 |
| Ccdc190 | 16.172407 | 0.758012896 | 0.228440494 | 3.318207211 | 0.000905972 | 0.005414076 |
| Irf8 | 15.58224746 | 0.757807426 | 0.248683588 | 3.047275589 | 0.002309259 | 0.011526574 |
| Fa2h | 70.79962321 | 0.756685529 | 0.226183213 | 3.34545397 | 0.00082148 | 0.004985721 |
| Ndrg1 | 194.4689877 | 0.756467 | 0.175925555 | 4.29992675 | 1.71E-05 | 0.000207239 |
| Gas5 | 2318.41529 | 0.755051572 | 0.150303951 | 5.02349783 | 5.07E-07 | 1.13E-05 |
| Zfhx3 | 13.88092842 | 0.753097679 | 0.249184333 | 3.022251316 | 0.002509021 | 0.012374287 |
| Sf3b6 | 723.5581644 | 0.752281608 | 0.136475962 | 5.512191261 | 3.54E-08 | 1.23E-06 |
| Rhog | 105.9716819 | 0.750702643 | 0.172524411 | 4.351283625 | 1.35E-05 | 0.000170363 |
| Rps29 | 11906.81807 | 0.74965671 | 0.169399654 | 4.425373318 | 9.63E-06 | 0.000129214 |
| Frmd6 | 65.74132079 | 0.7464528 | 0.155441788 | 4.802137238 | 1.57E-06 | 2.90E-05 |
| Smad9 | 20.70030685 | 0.746337781 | 0.18763033 | 3.977703289 | 6.96E-05 | 0.000665268 |
| Bhlhe41 | 136.731813 | 0.745996067 | 0.146860224 | 5.079633192 | 3.78E-07 | 9.08E-06 |
| Tmem106c | 108.0486617 | 0.745283429 | 0.124776181 | 5.972962356 | 2.33E-09 | 1.20E-07 |
| Metrnl | 89.15753298 | 0.74497044 | 0.159858286 | 4.660192828 | 3.16E-06 | 5.21E-05 |
| Crip1 | 114.1956945 | 0.744798053 | 0.176381416 | 4.222656037 | 2.41E-05 | 0.000274823 |
| Fundc1 | 156.059073 | 0.743155693 | 0.08077652 | 9.200144993 | 3.57E-20 | 2.75E-17 |
| Bcas1 | 310.6086417 | 0.742271643 | 0.19477751 | 3.810869342 | 0.000138479 | 0.001166099 |
| Dph7 | 49.36884721 | 0.741222388 | 0.167669376 | 4.420738037 | 9.84E-06 | 0.0001316 |
| Cbr3 | 59.76745778 | 0.740510155 | 0.270580567 | 2.736745524 | 0.006205027 | 0.025473823 |
| Taf9b | 90.58814069 | 0.740213501 | 0.171976191 | 4.304162655 | 1.68E-05 | 0.00020441 |

| | | | | | | |
|---|---|---|---|---|---|---|
| Vamp1 | 303.8702111 | 0.739265556 | 0.242889717 | 3.04362641 | 0.002337452 | 0.011646723 |
| Gm30082 | 24.30896411 | 0.738596194 | 0.195105264 | 3.785629251 | 0.00015332 | 0.001265302 |
| Gng11 | 133.6531839 | 0.737963354 | 0.145588505 | 5.068829807 | 4.00E-07 | 9.40E-06 |
| Zfas1 | 432.795408 | 0.736153328 | 0.076242556 | 9.655412549 | 4.66E-22 | 4.39E-19 |
| Efhd2 | 834.5328719 | 0.735996451 | 0.147454007 | 4.991362841 | 6.00E-07 | 1.30E-05 |
| C030037D09Rik | 15.9144257 | 0.734898473 | 0.250716108 | 2.931197681 | 0.003376578 | 0.015641472 |
| Sox9 | 102.2174844 | 0.734611642 | 0.186106631 | 3.947262057 | 7.91E-05 | 0.000733832 |
| Tceal1 | 182.9845057 | 0.734063735 | 0.101303544 | 7.246180234 | 4.29E-13 | 7.06E-11 |
| Gpr62 | 47.0869692 | 0.73282767 | 0.184262573 | 3.977083671 | 6.98E-05 | 0.000666627 |
| Srcap | 23.52564662 | 0.73237817 | 0.27700819 | 2.643886344 | 0.008196017 | 0.031864836 |
| Tomm34 | 288.7116883 | 0.731055332 | 0.135552215 | 5.393164064 | 6.92E-08 | 2.15E-06 |
| Spin2c | 35.91757832 | 0.730589721 | 0.197647888 | 3.69642059 | 0.000218661 | 0.001679436 |
| Gm14418 | 37.51127494 | 0.730344003 | 0.174927483 | 4.175124407 | 2.98E-05 | 0.000326969 |
| Fbxo32 | 24.58214259 | 0.730020107 | 0.178008987 | 4.101029509 | 4.11E-05 | 0.000429324 |
| Nek7 | 110.8880223 | 0.729115828 | 0.112096923 | 6.504334016 | 7.80E-11 | 6.45E-09 |
| Nol3 | 115.1073906 | 0.726775242 | 0.133674286 | 5.436911326 | 5.42E-08 | 1.75E-06 |
| Lars2 | 162.9566722 | 0.726763263 | 0.251063096 | 2.89474349 | 0.003794685 | 0.017203451 |
| Fstl4 | 13.84055412 | 0.724530827 | 0.245131197 | 2.955685925 | 0.003119746 | 0.014689671 |
| Pabpn1 | 123.9205263 | 0.723723519 | 0.171100217 | 4.229822326 | 2.34E-05 | 0.000268152 |
| Wdyhv1 | 64.63426293 | 0.722534064 | 0.164124809 | 4.402345196 | 1.07E-05 | 0.000141153 |
| Zxdb | 75.53150047 | 0.722053505 | 0.115746754 | 6.238218179 | 4.43E-10 | 2.92E-08 |
| Aass | 15.78224109 | 0.721706437 | 0.218903467 | 3.296916428 | 0.000977526 | 0.005769513 |
| Ifih1 | 11.7715016 | 0.7213315 | 0.272759115 | 2.644573402 | 0.008179396 | 0.031825513 |
| Ankrd33b | 124.5050182 | 0.720414577 | 0.167281675 | 4.306595901 | 1.66E-05 | 0.000202646 |
| Gm20878 | 25.71449704 | 0.71912327 | 0.27013367 | 2.662101578 | 0.007765444 | 0.030477435 |
| Ier3 | 58.92204433 | 0.718873752 | 0.152341843 | 4.718820123 | 2.37E-06 | 4.12E-05 |
| Ppie | 49.42538614 | 0.718425853 | 0.162565495 | 4.419300995 | 9.90E-06 | 0.000132269 |
| Prdm16os | 18.47032099 | 0.717662703 | 0.210993455 | 3.401350537 | 0.000670538 | 0.004234089 |
| Tmbim1 | 93.86311316 | 0.717187873 | 0.156176095 | 4.592174445 | 4.39E-06 | 6.75E-05 |
| Relt | 44.75968836 | 0.715864012 | 0.232476379 | 3.079297838 | 0.002074891 | 0.010568354 |
| Pcgf5 | 53.26085437 | 0.715458079 | 0.125956865 | 5.68018327 | 1.35E-08 | 5.29E-07 |
| Lipo3 | 20.71304287 | 0.714717315 | 0.207676792 | 3.44148862 | 0.000578523 | 0.003747245 |
| Lgi4 | 70.68372775 | 0.714154884 | 0.151080194 | 4.726992106 | 2.28E-06 | 3.97E-05 |
| Eif5a2 | 131.4556901 | 0.713250565 | 0.121950818 | 5.848673879 | 4.96E-09 | 2.22E-07 |
| Cast | 26.10083691 | 0.713138076 | 0.1803904 | 3.953303913 | 7.71E-05 | 0.000721067 |
| Tpd52l1 | 160.3240601 | 0.711582667 | 0.144601314 | 4.920997249 | 8.61E-07 | 1.76E-05 |
| Lrrc6 | 15.64956824 | 0.711014724 | 0.242173047 | 2.935977937 | 0.00332498 | 0.015437341 |
| Hist2h2be | 15.75329731 | 0.710646651 | 0.2098342 | 3.386705558 | 0.000707373 | 0.004418082 |
| Fosl2 | 119.2142112 | 0.71016273 | 0.106905055 | 6.642929379 | 3.08E-11 | 2.91E-09 |
| Sstr2 | 95.48884693 | 0.709337831 | 0.228516761 | 3.104095413 | 0.001908618 | 0.009905995 |
| Zfp119b | 11.56435189 | 0.709238412 | 0.254245781 | 2.789577902 | 0.00527768 | 0.022399086 |
| Resp18 | 687.7708296 | 0.708068501 | 0.169862369 | 4.168483609 | 3.07E-05 | 0.000335988 |
| Hhatl | 30.15608055 | 0.708023488 | 0.220524177 | 3.210638849 | 0.001324403 | 0.007348592 |
| Trem2 | 125.3509078 | 0.705352456 | 0.133230591 | 5.294222978 | 1.20E-07 | 3.40E-06 |
| mt-Nd4l | 2057.004056 | 0.704896773 | 0.231132928 | 3.049746217 | 0.002290348 | 0.011465945 |
| Lama3 | 15.07141988 | 0.704562524 | 0.249637132 | 2.822346649 | 0.004767362 | 0.020694378 |
| Cox17 | 1630.787929 | 0.704418667 | 0.1294362 | 5.442207558 | 5.26E-08 | 1.71E-06 |
| Tyrobp | 443.7043349 | 0.702560128 | 0.13400902 | 5.242633138 | 1.58E-07 | 4.35E-06 |
| Skap2 | 51.31681309 | 0.700734143 | 0.136813604 | 5.12181629 | 3.03E-07 | 7.52E-06 |
| Filip1l | 10.83289587 | 0.699948326 | 0.257085703 | 2.72262642 | 0.006476525 | 0.026313651 |
| Ppp1r3c | 119.4949581 | 0.69824051 | 0.097938624 | 7.129368174 | 1.01E-12 | 1.50E-10 |
| Gm9989 | 94.4341261 | 0.697075824 | 0.211866383 | 3.290167195 | 0.001001279 | 0.0058892 |
| Rbm4b | 72.57131773 | 0.696553174 | 0.115157746 | 6.048687101 | 1.46E-09 | 8.06E-08 |
| Ccdc88b | 14.47868568 | 0.696208455 | 0.269820057 | 2.580269466 | 0.009872325 | 0.037072613 |
| Rhpn2 | 19.43619811 | 0.693677841 | 0.189607684 | 3.658490126 | 0.000253706 | 0.001900381 |
| Ntn4 | 11.72625722 | 0.692960671 | 0.249756242 | 2.77454796 | 0.005527848 | 0.023210938 |
| Rps28 | 4973.146326 | 0.692822887 | 0.16607161 | 4.171832193 | 3.02E-05 | 0.000331323 |
| Sdf2l1 | 59.55311534 | 0.691812339 | 0.174680301 | 3.960448516 | 7.48E-05 | 0.000706063 |
| Lonrf1 | 84.84642071 | 0.68943054 | 0.161087051 | 4.279863196 | 1.87E-05 | 0.000223081 |
| Dusp5 | 123.2653795 | 0.688165544 | 0.128078623 | 5.372992995 | 7.74E-08 | 2.35E-06 |
| Eif2s3x | 339.3217753 | 0.687576348 | 0.147500824 | 4.661508522 | 3.14E-06 | 5.19E-05 |
| Prrg1 | 15.5417094 | 0.687461538 | 0.252259408 | 2.725216645 | 0.006425931 | 0.026165254 |
| Entpd3 | 26.03067747 | 0.687280807 | 0.169510012 | 4.054514536 | 5.02E-05 | 0.000507808 |
| Rps6 | 1871.403423 | 0.686568776 | 0.097887609 | 7.013847619 | 2.32E-12 | 3.07E-10 |
| A830031A19Rik | 9.197556976 | 0.686185361 | 0.268330467 | 2.557239839 | 0.010550645 | 0.039185797 |
| Dnajc12 | 83.21977764 | 0.685888208 | 0.132678723 | 5.169541831 | 2.35E-07 | 6.09E-06 |
| Sqor | 16.61168845 | 0.684115397 | 0.220115899 | 3.107978111 | 0.00188372 | 0.009797773 |
| Synpr | 443.5264372 | 0.684041807 | 0.193135935 | 3.541763518 | 0.000397462 | 0.002748826 |
| Fam122b | 9.988782847 | 0.682220365 | 0.271158072 | 2.515950789 | 0.011871176 | 0.042860128 |
| 2010204K13Rik | 194.821804 | 0.680851596 | 0.214825551 | 3.169323171 | 0.001527944 | 0.008256353 |
| Syde1 | 24.1024649 | 0.680548654 | 0.245234813 | 2.77508991 | 0.005518645 | 0.023178035 |
| Aktip | 364.1907725 | 0.680423613 | 0.110661199 | 6.148709908 | 7.81E-10 | 4.78E-08 |
| Samd4 | 60.09324102 | 0.679806192 | 0.155493698 | 4.371921178 | 1.23E-05 | 0.000157203 |

| | | | | | | |
|---|---|---|---|---|---|---|
| B3galt5 | 40.53183875 | 0.679787316 | 0.217730887 | 3.122144615 | 0.001795387 | 0.00942217 |
| Aldh3b1 | 13.38392177 | 0.679554833 | 0.230587 | 2.947064806 | 0.00320806 | 0.015053054 |
| Btg2 | 62.20437609 | 0.679182592 | 0.119201353 | 5.697775851 | 1.21E-08 | 4.85E-07 |
| Cox6a2 | 327.9152218 | 0.677750175 | 0.110129458 | 6.154122483 | 7.55E-10 | 4.65E-08 |
| Utp3 | 154.8605887 | 0.676570331 | 0.079674265 | 8.491704705 | 2.04E-17 | 9.86E-15 |
| Bbx | 39.37145698 | 0.676379601 | 0.209979332 | 3.221172261 | 0.001276674 | 0.007161247 |
| Sox2ot | 48.1109192 | 0.67603587 | 0.194150005 | 3.4820286 | 0.000497631 | 0.003306645 |
| Mpp5 | 70.82801037 | 0.674011579 | 0.132335415 | 5.09320639 | 3.52E-07 | 8.55E-06 |
| Stard10 | 168.2008687 | 0.67292747 | 0.20574777 | 3.270642839 | 0.001073033 | 0.006231127 |
| Bcap29 | 181.9633363 | 0.67253997 | 0.107117869 | 6.278504003 | 3.42E-10 | 2.36E-08 |
| mt-Nd1 | 47414.18688 | 0.672449371 | 0.169349168 | 3.970786382 | 7.16E-05 | 0.000680279 |
| mt-Co2 | 88555.27371 | 0.670454168 | 0.181333318 | 3.697357851 | 0.000217855 | 0.001674008 |
| Rpl10 | 1848.910823 | 0.668404634 | 0.113083743 | 5.910704903 | 3.41E-09 | 1.65E-07 |
| Slc35g2 | 51.74020049 | 0.668348706 | 0.136021666 | 4.913545952 | 8.94E-07 | 1.82E-05 |
| Wfdc18 | 13.73433786 | 0.668291099 | 0.255547951 | 2.615129948 | 0.008919351 | 0.03417539 |
| Iqcb1 | 111.7932359 | 0.667391837 | 0.101745688 | 6.559411504 | 5.40E-11 | 4.72E-09 |
| Cchcr1 | 12.79905573 | 0.666933121 | 0.240417675 | 2.774060265 | 0.005536142 | 0.023233799 |
| Erbin | 101.5687932 | 0.666126953 | 0.148560599 | 4.483873638 | 7.33E-06 | 0.000103526 |
| Edil3 | 386.5792344 | 0.665949687 | 0.106300136 | 6.264805576 | 3.73E-10 | 2.54E-08 |
| Cdkn2aip | 45.06241057 | 0.665888802 | 0.130871412 | 5.088115052 | 3.62E-07 | 8.74E-06 |
| Nfe2l2 | 52.5644763 | 0.665479475 | 0.154819029 | 4.298434616 | 1.72E-05 | 0.000208414 |
| Dnajb1 | 236.2508753 | 0.665403161 | 0.134366122 | 4.952164656 | 7.34E-07 | 1.54E-05 |
| Lefty1 | 20.59708161 | 0.664757933 | 0.252587626 | 2.631791361 | 0.0084936 | 0.032863506 |
| Fgfr2 | 85.32868662 | 0.664444908 | 0.117119953 | 5.673199926 | 1.40E-08 | 5.45E-07 |
| Rps19 | 3637.331373 | 0.663314846 | 0.134947752 | 4.915345644 | 8.86E-07 | 1.81E-05 |
| Pnrc1 | 209.8439727 | 0.663048513 | 0.109993223 | 6.028085154 | 1.66E-09 | 8.96E-08 |
| Ralbp1 | 223.2463405 | 0.662557533 | 0.101704498 | 6.514535225 | 7.29E-11 | 6.15E-09 |
| Cdc42ep1 | 56.96776812 | 0.660258185 | 0.182892324 | 3.610092378 | 0.000306088 | 0.002221103 |
| Tspyl4 | 1247.57071 | 0.660037379 | 0.047047834 | 14.02907038 | 1.03E-44 | 8.77E-41 |
| Nap1l5 | 2253.757853 | 0.659998245 | 0.073121977 | 9.025990186 | 1.78E-19 | 1.21E-16 |
| Fbxo33 | 156.4985029 | 0.658875314 | 0.128671449 | 5.120602254 | 3.05E-07 | 7.54E-06 |
| Rpl5 | 2585.035244 | 0.65766913 | 0.166933387 | 3.939709967 | 8.16E-05 | 0.000752846 |
| Med30 | 169.0665374 | 0.653656866 | 0.078885166 | 8.286182322 | 1.17E-16 | 4.64E-14 |
| Flywch1 | 952.8211292 | 0.651680219 | 0.136276136 | 4.782056791 | 1.74E-06 | 3.16E-05 |
| Zfp825 | 29.27188714 | 0.65164789 | 0.151551419 | 4.299846845 | 1.71E-05 | 0.000207239 |
| Ldb3 | 21.41661818 | 0.651449094 | 0.204333484 | 3.188166141 | 0.001431782 | 0.007836663 |
| Kif5b | 453.9785084 | 0.650406485 | 0.105561331 | 6.161408543 | 7.21E-10 | 4.48E-08 |
| Wdr78 | 15.63844825 | 0.648908021 | 0.218568727 | 2.96889693 | 0.002988708 | 0.014222794 |
| Stat1 | 61.95158113 | 0.648861389 | 0.149513743 | 4.339811022 | 1.43E-05 | 0.000178136 |
| Senp8 | 26.6757944 | 0.648803078 | 0.17854324 | 3.633870863 | 0.000279201 | 0.002054136 |
| Rpl37 | 9340.449608 | 0.648600652 | 0.132299958 | 4.902500799 | 9.46E-07 | 1.92E-05 |
| Rpl26 | 4546.688683 | 0.647701445 | 0.146422687 | 4.423504718 | 9.71E-06 | 0.000130131 |
| Ddx23 | 26.49308904 | 0.647671228 | 0.203932336 | 3.175912365 | 0.001493661 | 0.008102093 |
| Lgals1 | 159.2147606 | 0.647229294 | 0.097823538 | 6.616294098 | 3.68E-11 | 3.41E-09 |
| Rerg | 70.24411174 | 0.646488246 | 0.18760472 | 3.44601269 | 0.000568924 | 0.003692123 |
| Cyba | 75.74625741 | 0.645296494 | 0.193616181 | 3.332864493 | 0.000859568 | 0.005179714 |
| Gjc2 | 34.96560292 | 0.645220625 | 0.229564737 | 2.810626033 | 0.004944522 | 0.021299769 |
| Gm49492 | 14.83982013 | 0.644761902 | 0.237724458 | 2.712223668 | 0.006683348 | 0.026960836 |
| Stat6 | 30.54212327 | 0.643518016 | 0.173836519 | 3.701857466 | 0.000214027 | 0.001650577 |
| Hspa2 | 159.2243017 | 0.643122817 | 0.097376899 | 6.604470104 | 3.99E-11 | 3.64E-09 |
| Agmo | 14.39009344 | 0.642860851 | 0.252396727 | 2.547025311 | 0.010864555 | 0.039992416 |
| Gm15728 | 23.13246535 | 0.642733682 | 0.233725268 | 2.749953773 | 0.005960367 | 0.024612469 |
| Rsl1 | 13.58303578 | 0.641565643 | 0.226358334 | 2.834292123 | 0.004592733 | 0.020095353 |
| Crebrf | 128.4904489 | 0.640103429 | 0.156587981 | 4.087819674 | 4.35E-05 | 0.00045028 |
| Bag3 | 30.78415323 | 0.639451967 | 0.188973522 | 3.383817801 | 0.000714854 | 0.004453324 |
| Slc5a3 | 61.66030784 | 0.638439091 | 0.195290974 | 3.269168447 | 0.001078641 | 0.006253091 |
| Eif3e | 377.3917574 | 0.638319226 | 0.089079658 | 7.165712545 | 7.74E-13 | 1.21E-10 |
| Amd1 | 59.57560619 | 0.638316655 | 0.148062784 | 4.311121531 | 1.62E-05 | 0.000199307 |
| Grb7 | 26.51110068 | 0.638293337 | 0.218552497 | 2.92054928 | 0.003494149 | 0.016064368 |
| mt-Nd5 | 6821.690135 | 0.636772857 | 0.177922664 | 3.578930545 | 0.000345003 | 0.002457205 |
| Unc5b | 28.7745898 | 0.635632033 | 0.244729752 | 2.597281402 | 0.009396491 | 0.035665006 |
| Hvcn1 | 12.11021646 | 0.633631308 | 0.250479625 | 2.529672051 | 0.011416918 | 0.041643337 |
| Khdrbs1 | 385.5811256 | 0.63356758 | 0.089320226 | 7.09321517 | 1.31E-12 | 1.85E-10 |
| Ints12 | 35.73367717 | 0.633076729 | 0.137796832 | 4.594276355 | 4.34E-06 | 6.69E-05 |
| Tceal9 | 542.4276963 | 0.63254014 | 0.085074311 | 7.435148526 | 1.04E-13 | 2.06E-11 |
| Rpl36a | 2345.114804 | 0.630308983 | 0.110204959 | 5.719424871 | 1.07E-08 | 4.30E-07 |
| mt-Nd3 | 18163.93423 | 0.629834426 | 0.183975422 | 3.423470478 | 0.00061827 | 0.003966801 |
| Dkc1 | 85.70858974 | 0.629064 | 0.13687042 | 4.596055149 | 4.31E-06 | 6.67E-05 |
| Abt1 | 60.33824993 | 0.62841319 | 0.134770462 | 4.662840654 | 3.12E-06 | 5.17E-05 |
| Fcer1g | 151.6976958 | 0.628363878 | 0.135863882 | 4.624951602 | 3.75E-06 | 6.01E-05 |
| Srebf1 | 120.0612729 | 0.627801557 | 0.118202758 | 5.311225962 | 1.09E-07 | 3.16E-06 |
| Hebp1 | 81.53804861 | 0.627676397 | 0.169358614 | 3.706197063 | 0.000210395 | 0.001629967 |
| Klhl15 | 37.19116896 | 0.627670061 | 0.153649705 | 4.085071701 | 4.41E-05 | 0.000454328 |

| | | | | | | |
|---|---|---|---|---|---|---|
| Gask1a | 12.50329293 | 0.627471038 | 0.239169706 | 2.623538946 | 0.00870215 | 0.033479377 |
| Car2 | 623.5782189 | 0.627457777 | 0.218336107 | 2.873815906 | 0.004055453 | 0.018171816 |
| 04.Sep | 443.5985153 | 0.627235833 | 0.17098882 | 3.668285627 | 0.000244182 | 0.001841029 |
| Twistnb | 43.04389645 | 0.6271062 | 0.127977851 | 4.900115092 | 9.58E-07 | 1.93E-05 |
| Rpl30 | 4250.95379 | 0.625262353 | 0.114744361 | 5.449177188 | 5.06E-08 | 1.66E-06 |
| Asah2 | 23.51533406 | 0.624050062 | 0.171926963 | 3.629739341 | 0.000283708 | 0.002082775 |
| Plek | 27.26554262 | 0.620618278 | 0.176355767 | 3.519126645 | 0.00043297 | 0.002952242 |
| Spp1 | 82.43842535 | 0.619748473 | 0.236329513 | 2.622391361 | 0.00873151 | 0.033561866 |
| Rnf7 | 544.3520581 | 0.619074095 | 0.101523196 | 6.097858623 | 1.07E-09 | 6.24E-08 |
| Pip4k2a | 201.8769013 | 0.618111146 | 0.116519507 | 5.304786803 | 1.13E-07 | 3.24E-06 |
| Impact | 969.6042371 | 0.617850478 | 0.076933988 | 8.030917052 | 9.67E-16 | 3.22E-13 |
| D16Ertd472e | 45.55046735 | 0.617265539 | 0.148145315 | 4.166622069 | 3.09E-05 | 0.000338523 |
| Aif1 | 70.7874839 | 0.616522588 | 0.139497204 | 4.419605345 | 9.89E-06 | 0.000132187 |
| Adat2 | 10.47824715 | 0.616449605 | 0.245502493 | 2.510970852 | 0.012039964 | 0.043349496 |
| Cmtm5 | 250.5012639 | 0.61582372 | 0.147372168 | 4.178697564 | 2.93E-05 | 0.000323131 |
| C1qa | 424.1115422 | 0.61522724 | 0.159913677 | 3.847245913 | 0.000119453 | 0.001034141 |
| Cd37 | 18.50947984 | 0.615103462 | 0.195568515 | 3.145206995 | 0.001659694 | 0.008849785 |
| Nrsn2 | 548.5486599 | 0.614017949 | 0.155751277 | 3.942298006 | 8.07E-05 | 0.000746459 |
| Klhl4 | 29.88461527 | 0.612227693 | 0.156465875 | 3.912851246 | 9.12E-05 | 0.000826928 |
| Psmd1 | 180.0877848 | 0.611070218 | 0.196625358 | 3.107789474 | 0.001884923 | 0.009801021 |
| Apoc1 | 21.0667923 | 0.610444667 | 0.182890062 | 3.337768389 | 0.000844541 | 0.005101859 |
| Pex5l | 325.5644308 | 0.610214954 | 0.077983455 | 7.824928449 | 5.08E-15 | 1.37E-12 |
| Epdr1 | 412.2972614 | 0.610187905 | 0.127892381 | 4.77110442 | 1.83E-06 | 3.31E-05 |
| Nudt17 | 42.32986401 | 0.608810295 | 0.170438887 | 3.5720152 | 0.000354245 | 0.002508273 |
| Fth1 | 20730.9311 | 0.608102355 | 0.12953598 | 4.694466792 | 2.67E-06 | 4.56E-05 |
| 1700048O20Rik | 24.03991708 | 0.607516831 | 0.168780263 | 3.59945423 | 0.000318886 | 0.00229393 |
| Zrsr1 | 239.8595239 | 0.607119018 | 0.117920291 | 5.148554257 | 2.63E-07 | 6.68E-06 |
| Chac1 | 50.79635901 | 0.605642536 | 0.127694649 | 4.742896759 | 2.11E-06 | 3.72E-05 |
| Pcsk6 | 49.06533463 | 0.605513724 | 0.169418568 | 3.574069427 | 0.000351476 | 0.002493874 |
| Fgfbp3 | 37.11965844 | 0.605204114 | 0.177478285 | 3.410017824 | 0.000649586 | 0.004130209 |
| Arhgef6 | 31.5787271 | 0.604946129 | 0.185832726 | 3.255326138 | 0.001132622 | 0.006505957 |
| Wls | 115.405721 | 0.604503681 | 0.111764626 | 5.408721 | 6.35E-08 | 2.00E-06 |
| Crebl2 | 89.68707405 | 0.604411721 | 0.168040258 | 3.596826895 | 0.000322123 | 0.002309772 |
| Krt12 | 127.9579805 | 0.604331579 | 0.095264033 | 6.34375386 | 2.24E-10 | 1.63E-08 |
| Tlr3 | 27.2315836 | 0.603849191 | 0.187029959 | 3.228622798 | 0.001243878 | 0.007009635 |
| Gsn | 154.1214985 | 0.602056661 | 0.172929893 | 3.481507164 | 0.0004986 | 0.003309231 |
| Fbxl14 | 57.41742256 | 0.601189202 | 0.111150032 | 5.408808154 | 6.34E-08 | 2.00E-06 |
| Efr3a | 591.2864674 | 0.600574139 | 0.170719548 | 3.517899059 | 0.000434978 | 0.002961168 |
| Amotl1 | 36.05828352 | 0.600509711 | 0.151755915 | 3.957076135 | 7.59E-05 | 0.0007128 |
| Mcm8 | 18.40843989 | 0.59977575 | 0.197156388 | 3.042131965 | 0.002349089 | 0.011690959 |
| Cyp4v3 | 24.66989602 | 0.599268478 | 0.17713017 | 3.383209531 | 0.000716439 | 0.004459921 |
| Gm4631 | 50.3843985 | 0.598662763 | 0.124603718 | 4.804533688 | 1.55E-06 | 2.87E-05 |
| Oxr1 | 1753.661968 | 0.598269308 | 0.121010388 | 4.943950002 | 7.66E-07 | 1.59E-05 |
| Cpm | 14.2235023 | 0.597853415 | 0.220033507 | 2.717101698 | 0.006585637 | 0.026674583 |
| Klf10 | 93.67422742 | 0.597074275 | 0.133980422 | 4.456429263 | 8.33E-06 | 0.000114941 |
| Eif4ebp1 | 27.18064756 | 0.596996384 | 0.16581935 | 3.600281778 | 0.000317873 | 0.002288979 |
| Paip1 | 169.106056 | 0.596925563 | 0.134185278 | 4.448517554 | 8.65E-06 | 0.000118678 |
| Rpl22l1 | 2523.245791 | 0.596476232 | 0.127099412 | 4.692989709 | 2.69E-06 | 4.57E-05 |
| Rpl35 | 2583.981956 | 0.595650627 | 0.154813076 | 3.847547266 | 0.000119306 | 0.001033904 |
| Sertad1 | 69.80482112 | 0.594693732 | 0.133489772 | 4.454976013 | 8.39E-06 | 0.000115441 |
| Mylk | 34.81846453 | 0.594311705 | 0.183386304 | 3.240763845 | 0.001192099 | 0.006765072 |
| Zfp850 | 14.73575305 | 0.594106031 | 0.222708902 | 2.667634865 | 0.007638722 | 0.030077581 |
| Kcna1 | 14.26606606 | 0.593803179 | 0.226111307 | 2.62615429 | 0.008635566 | 0.033298791 |
| Umad1 | 187.5614386 | 0.593519436 | 0.099067187 | 5.991079905 | 2.08E-09 | 1.09E-07 |
| Mir99ahg | 18.80046321 | 0.59265425 | 0.226120611 | 2.620965195 | 0.008768121 | 0.033672049 |
| Polr2k | 567.0506564 | 0.592198073 | 0.084393774 | 7.017082455 | 2.27E-12 | 3.02E-10 |
| Prkg2 | 34.83202281 | 0.591485301 | 0.214888206 | 2.752525656 | 0.005913751 | 0.024449755 |
| Cyb5r1 | 96.66519572 | 0.591437722 | 0.092871877 | 6.368318826 | 1.91E-10 | 1.41E-08 |
| Vwa5a | 25.93950758 | 0.591068377 | 0.173641118 | 3.403965506 | 0.000664151 | 0.004207014 |
| Rnaset2b | 82.98390525 | 0.589268981 | 0.137398624 | 4.2887546 | 1.80E-05 | 0.000216469 |
| Xrcc3 | 13.82457144 | 0.587474146 | 0.235981154 | 2.489496033 | 0.012792435 | 0.045469608 |
| Gm42418 | 1420.613404 | 0.587092236 | 0.211516394 | 2.775634665 | 0.005509408 | 0.023144974 |
| Amd2 | 19.58024523 | 0.586512936 | 0.235374677 | 2.491826828 | 0.012708798 | 0.045229234 |
| Atp6v1h | 609.8308229 | 0.586193439 | 0.083265163 | 7.040080331 | 1.92E-12 | 2.61E-10 |
| Atf4 | 733.1888577 | 0.586082594 | 0.087305197 | 6.713032151 | 1.91E-11 | 1.92E-09 |
| Acot13 | 814.0767494 | 0.585454202 | 0.0936853 | 6.249157546 | 4.13E-10 | 2.75E-08 |
| 2310010J17Rik | 48.32299317 | 0.585085421 | 0.139077525 | 4.206901289 | 2.59E-05 | 0.000291018 |
| Rps21 | 11350.67167 | 0.584646677 | 0.128105077 | 4.56380567 | 5.02E-06 | 7.59E-05 |
| Fam133b | 84.12027778 | 0.584535325 | 0.116941111 | 4.998544308 | 5.78E-07 | 1.26E-05 |
| Il17ra | 39.86287303 | 0.584463726 | 0.174704083 | 3.345449727 | 0.000821493 | 0.004985721 |
| Scn1b | 3480.310119 | 0.584025173 | 0.130653763 | 4.470021838 | 7.82E-06 | 0.000109027 |
| Tmem144 | 27.06429778 | 0.58331141 | 0.163436535 | 3.569039256 | 0.000358293 | 0.002530592 |
| Usf1 | 68.70165033 | 0.582219062 | 0.194081295 | 2.999872106 | 0.00270093 | 0.013114713 |

| | | | | | | |
|---|---|---|---|---|---|---|
| Zhx1 | 135.6124527 | 0.581021094 | 0.075821231 | 7.663039634 | 1.82E-14 | 4.34E-12 |
| Srgn | 82.12555806 | 0.580920612 | 0.173783945 | 3.342774919 | 0.000829451 | 0.005026826 |
| Gcat | 81.13319281 | 0.580467325 | 0.123860728 | 4.686451756 | 2.78E-06 | 4.69E-05 |
| Kiz | 46.43045107 | 0.579938368 | 0.134556522 | 4.309998216 | 1.63E-05 | 0.000199809 |
| Zfp729b | 39.95171727 | 0.579788559 | 0.151935456 | 3.816018822 | 0.000135622 | 0.001146028 |
| Gpr150 | 38.39106283 | 0.579463944 | 0.206261906 | 2.809359977 | 0.004964011 | 0.021356585 |
| Bex1 | 673.0427031 | 0.579275205 | 0.077708119 | 7.4545004 | 9.02E-14 | 1.80E-11 |
| Tob1 | 114.727816 | 0.578992202 | 0.100940943 | 5.735950013 | 9.70E-09 | 3.99E-07 |
| Celf4 | 1224.176734 | 0.578827153 | 0.13264718 | 4.363659693 | 1.28E-05 | 0.000162481 |
| Nfkbiz | 14.77279397 | 0.577781898 | 0.230137605 | 2.510593164 | 0.012052851 | 0.04337747 |
| Kctd4 | 96.56220991 | 0.577316388 | 0.170121984 | 3.393543699 | 0.000689945 | 0.004334791 |
| Grm3 | 18.11202313 | 0.57655548 | 0.222586306 | 2.590255845 | 0.009590463 | 0.036232863 |
| Gnai1 | 377.426836 | 0.57645481 | 0.106565908 | 5.40937359 | 6.32E-08 | 2.00E-06 |
| Csf1 | 32.86199423 | 0.576447485 | 0.160209022 | 3.598096283 | 0.000320555 | 0.002302427 |
| BC005561 | 19.89501126 | 0.575953605 | 0.209942341 | 2.743389453 | 0.006080852 | 0.025036804 |
| Sik1 | 16.70687996 | 0.57578809 | 0.203915557 | 2.82365945 | 0.00474788 | 0.020625658 |
| Fn3k | 198.2823183 | 0.575361919 | 0.115780024 | 4.969440297 | 6.71E-07 | 1.44E-05 |
| Gm2415 | 46.46905863 | 0.575324288 | 0.183326967 | 3.138241458 | 0.001699648 | 0.009028748 |
| Tceal7 | 13.13244668 | 0.574191034 | 0.233516994 | 2.458883289 | 0.013936992 | 0.04872055 |
| Ogfrl1 | 444.2018209 | 0.574170656 | 0.154530203 | 3.715588579 | 0.000202731 | 0.001587297 |
| Rpl37a | 5492.64902 | 0.57416442 | 0.150885152 | 3.805307637 | 0.000141628 | 0.001185549 |
| Lrif1 | 68.00175308 | 0.572842151 | 0.127229458 | 4.502433323 | 6.72E-06 | 9.63E-05 |
| Stx4a | 237.2190479 | 0.572200846 | 0.094448045 | 6.058366217 | 1.38E-09 | 7.67E-08 |
| Dnajb4 | 190.1387377 | 0.571246952 | 0.121707293 | 4.693613151 | 2.68E-06 | 4.57E-05 |
| Rnf6 | 173.1846067 | 0.570905317 | 0.081750422 | 6.983515229 | 2.88E-12 | 3.75E-10 |
| Gad1 | 1009.562827 | 0.570831798 | 0.116364872 | 4.905533659 | 9.32E-07 | 1.89E-05 |
| Tmcc3 | 59.20724988 | 0.569709417 | 0.136929145 | 4.160614736 | 3.17E-05 | 0.000346434 |
| Rbm12b1 | 26.0631684 | 0.569323084 | 0.158147633 | 3.599946921 | 0.000318282 | 0.002290956 |
| Dusp28 | 146.5075467 | 0.568401 | 0.18502721 | 3.071986012 | 0.002126397 | 0.010756357 |
| Arrdc3 | 79.9084919 | 0.568083203 | 0.122867109 | 4.623557969 | 3.77E-06 | 6.03E-05 |
| Trmt2b | 32.47776219 | 0.567745424 | 0.15507845 | 3.661020757 | 0.000251212 | 0.001883371 |
| Morf4l1 | 2058.321725 | 0.566606605 | 0.071107207 | 7.968342891 | 1.61E-15 | 5.14E-13 |
| Grhpr | 192.933954 | 0.56610213 | 0.157365198 | 3.597378182 | 0.000321441 | 0.002306836 |
| Htr5a | 50.80173703 | 0.565875067 | 0.12313034 | 4.595740294 | 4.31E-06 | 6.67E-05 |
| Nop58 | 96.83227925 | 0.56565388 | 0.120100995 | 4.709818446 | 2.48E-06 | 4.28E-05 |
| Efna1 | 31.40475844 | 0.565123485 | 0.221566232 | 2.550584897 | 0.010754233 | 0.039696957 |
| Tifa | 20.51096028 | 0.564847238 | 0.205219136 | 2.752410174 | 0.005915837 | 0.024452416 |
| Pttg1 | 280.1115558 | 0.562782224 | 0.160083546 | 3.515553198 | 0.000438839 | 0.002980271 |
| Bst2 | 32.06940746 | 0.562474848 | 0.173697723 | 3.23823962 | 0.001202697 | 0.006811535 |
| Tspyl2 | 188.3556177 | 0.561672058 | 0.116952 | 4.802586172 | 1.57E-06 | 2.90E-05 |
| Zmat4 | 96.91300055 | 0.561097509 | 0.191898021 | 2.923935881 | 0.003456359 | 0.015925183 |
| Fmn1 | 26.24644927 | 0.560861735 | 0.224007148 | 2.50376713 | 0.012287888 | 0.044036362 |
| Rpl17 | 4668.754691 | 0.557700982 | 0.121134789 | 4.603970407 | 4.15E-06 | 6.49E-05 |
| Gm14295 | 81.93943722 | 0.557144125 | 0.132246384 | 4.212925203 | 2.52E-05 | 0.000284683 |
| Peli3 | 55.67582883 | 0.555600244 | 0.117115348 | 4.744042996 | 2.09E-06 | 3.71E-05 |
| Gskip | 172.7690015 | 0.554950377 | 0.132045511 | 4.202720518 | 2.64E-05 | 0.000295467 |
| Plekhb2 | 739.2452875 | 0.554584007 | 0.057022212 | 9.725754033 | 2.34E-22 | 2.48E-19 |
| 9930021J03Rik | 105.5284266 | 0.554582707 | 0.136068711 | 4.075754825 | 4.59E-05 | 0.00047005 |
| Eepd1 | 60.64923955 | 0.554133852 | 0.147998656 | 3.744181643 | 0.000180983 | 0.00144846 |
| Tmem192 | 120.6860997 | 0.553756351 | 0.10490795 | 5.278497474 | 1.30E-07 | 3.67E-06 |
| Eif2ak3 | 50.33644724 | 0.553741297 | 0.136334941 | 4.061624207 | 4.87E-05 | 0.000496433 |
| Bmi1 | 117.2240206 | 0.553594007 | 0.104082682 | 5.318790782 | 1.04E-07 | 3.04E-06 |
| Ahcy | 20.4136345 | 0.552766816 | 0.178099005 | 3.103705242 | 0.001911136 | 0.009911542 |
| Klf5 | 109.9126272 | 0.552171817 | 0.131080936 | 4.212449433 | 2.53E-05 | 0.000285093 |
| M6pr | 290.0965911 | 0.551727736 | 0.069481219 | 7.94067438 | 2.01E-15 | 6.09E-13 |
| Arhgap10 | 24.84037955 | 0.551476685 | 0.169053576 | 3.262141496 | 0.001105739 | 0.006386163 |
| Zfp941 | 57.29023481 | 0.551361126 | 0.145102813 | 3.799796244 | 0.000144815 | 0.001205678 |
| Ints9 | 47.70920586 | 0.55013977 | 0.168034411 | 3.27397089 | 0.001060475 | 0.006173117 |
| Set | 650.3105067 | 0.548971875 | 0.079102035 | 6.940047455 | 3.92E-12 | 4.96E-10 |
| Ciart | 70.77824892 | 0.548596558 | 0.131849137 | 4.160789893 | 3.17E-05 | 0.000346391 |
| Fnbp1 | 141.3100137 | 0.548241491 | 0.101342749 | 5.409775201 | 6.31E-08 | 2.00E-06 |
| Aldh9a1 | 73.82626668 | 0.548125776 | 0.120438726 | 4.551075831 | 5.34E-06 | 7.96E-05 |
| Rpl35a | 6003.084354 | 0.547729813 | 0.132822122 | 4.123784544 | 3.73E-05 | 0.000394111 |
| Plcl1 | 79.79221035 | 0.547207743 | 0.111140767 | 4.923555575 | 8.50E-07 | 1.74E-05 |
| Rps3a1 | 3744.85882 | 0.547173962 | 0.099904435 | 5.476973693 | 4.33E-08 | 1.45E-06 |
| Pfdn2 | 808.1164259 | 0.547146921 | 0.118743282 | 4.60781368 | 4.07E-06 | 6.41E-05 |
| Rsrc1 | 113.6025634 | 0.54712937 | 0.089362281 | 6.122598531 | 9.21E-10 | 5.44E-08 |
| Lats2 | 60.05765262 | 0.545883842 | 0.127737667 | 4.273475899 | 1.92E-05 | 0.000228606 |
| Gadd45a | 46.33408732 | 0.544890773 | 0.162765936 | 3.347695393 | 0.000814865 | 0.004954369 |
| Flrt1 | 37.16944385 | 0.541121276 | 0.155525479 | 3.479309486 | 0.000502708 | 0.003327371 |
| Necab1 | 784.6442918 | 0.540074418 | 0.086067436 | 6.275014595 | 3.50E-10 | 2.40E-08 |
| Ccdc92 | 391.1322757 | 0.539312587 | 0.119143723 | 4.526571543 | 5.99E-06 | 8.76E-05 |
| Pacsin2 | 192.5337734 | 0.538505548 | 0.113544514 | 4.742682226 | 2.11E-06 | 3.72E-05 |

| | | | | | | |
|---|---|---|---|---|---|---|
| Znfx1 | 97.82889968 | 0.538016332 | 0.118962357 | 4.52257627 | 6.11E-06 | 8.91E-05 |
| Arl4d | 89.89382519 | 0.537878884 | 0.165776221 | 3.244608194 | 0.001176123 | 0.006687845 |
| Rps23 | 4063.603794 | 0.537329094 | 0.120141261 | 4.472477574 | 7.73E-06 | 0.000108137 |
| Mcub | 29.01193055 | 0.536845127 | 0.175723641 | 3.055053508 | 0.002250204 | 0.011300886 |
| Peg13 | 696.1808014 | 0.536119229 | 0.113807867 | 4.710739632 | 2.47E-06 | 4.26E-05 |
| Svip | 173.9230116 | 0.53601368 | 0.086085348 | 6.226537882 | 4.77E-10 | 3.11E-08 |
| Hscb | 53.73631308 | 0.535756426 | 0.122209076 | 4.383933221 | 1.17E-05 | 0.00015036 |
| Gnpda2 | 64.06970313 | 0.535654687 | 0.120658341 | 4.439433557 | 9.02E-06 | 0.000123002 |
| Socs4 | 33.30690666 | 0.535578682 | 0.150186783 | 3.566083985 | 0.000362355 | 0.002551842 |
| Ephx2 | 73.92328303 | 0.535395923 | 0.124098818 | 4.314270939 | 1.60E-05 | 0.000196836 |
| Kif3a | 150.0923949 | 0.534830394 | 0.213795916 | 2.501593128 | 0.012363592 | 0.044251531 |
| Zhx2 | 18.52981091 | 0.534701845 | 0.216013251 | 2.475319651 | 0.013311697 | 0.046921727 |
| Ide | 197.1324532 | 0.534229633 | 0.183195375 | 2.91617424 | 0.003543525 | 0.016264191 |
| Isca1 | 428.2988675 | 0.534214907 | 0.100611186 | 5.309696951 | 1.10E-07 | 3.17E-06 |
| S100b | 659.987427 | 0.534179502 | 0.139176179 | 3.838153252 | 0.000123963 | 0.001065027 |
| Dpp9 | 96.39334149 | 0.533914951 | 0.187671738 | 2.844940619 | 0.00444197 | 0.019572612 |
| Nupr1 | 59.52481511 | 0.533339588 | 0.176588758 | 3.020235222 | 0.002525784 | 0.012434248 |
| Zbtb4 | 291.2532452 | 0.532429268 | 0.119977568 | 4.437740123 | 9.09E-06 | 0.000123675 |
| Sh3bp1 | 34.00963551 | 0.532222464 | 0.213677712 | 2.490772006 | 0.012746589 | 0.045353717 |
| Cd83 | 46.34763201 | 0.531824494 | 0.18816074 | 2.826437095 | 0.004706899 | 0.020484375 |
| Hlf | 440.533512 | 0.531703636 | 0.114778335 | 4.632438999 | 3.61E-06 | 5.84E-05 |
| Them6 | 99.4169463 | 0.531653904 | 0.124461664 | 4.271627798 | 1.94E-05 | 0.000229865 |
| Gm38393 | 18.68934645 | 0.531598292 | 0.189565595 | 2.804297325 | 0.005042638 | 0.021610478 |
| Papss2 | 30.34307348 | 0.531410611 | 0.196510736 | 2.704231953 | 0.006846248 | 0.027519742 |
| Pcnt | 156.9482032 | 0.53126778 | 0.1298657 | 4.090901462 | 4.30E-05 | 0.000445845 |
| Cpeb1 | 157.6362564 | 0.530299001 | 0.18238826 | 2.907528152 | 0.003642976 | 0.016631317 |
| Carhsp1 | 93.45864709 | 0.530211293 | 0.179355206 | 2.956207991 | 0.00311447 | 0.014672979 |
| Clp1 | 26.90434028 | 0.530195778 | 0.161050085 | 3.292117344 | 0.000994361 | 0.005854608 |
| Mapt | 461.2868973 | 0.530104916 | 0.118354581 | 4.478955623 | 7.50E-06 | 0.000105429 |
| Hes1 | 61.98805526 | 0.529940496 | 0.129765123 | 4.083843817 | 4.43E-05 | 0.000456459 |
| Ndufb1-ps | 3394.281914 | 0.52985391 | 0.105943544 | 5.001285482 | 5.69E-07 | 1.25E-05 |
| Usp5 | 563.8125932 | 0.52948715 | 0.072498966 | 7.303375239 | 2.81E-13 | 4.81E-11 |
| Elmsan1 | 23.57174132 | 0.5291931 | 0.184321668 | 2.871030338 | 0.004091363 | 0.018303691 |
| Rpl38 | 11035.15552 | 0.52789962 | 0.110242388 | 4.788535774 | 1.68E-06 | 3.08E-05 |
| Uap1l1 | 74.58444104 | 0.527562206 | 0.13781863 | 3.827945501 | 0.000129217 | 0.001105107 |
| Zfp934 | 17.56585858 | 0.52702416 | 0.203638063 | 2.588043472 | 0.00965228 | 0.03643193 |
| Xrcc5 | 57.69125347 | 0.526951196 | 0.113065102 | 4.660599846 | 3.15E-06 | 5.21E-05 |
| Scly | 42.89264812 | 0.526657939 | 0.154919954 | 3.39954877 | 0.000674971 | 0.004258072 |
| Psme2 | 148.1104952 | 0.526534628 | 0.104757129 | 5.026241492 | 5.00E-07 | 1.12E-05 |
| Cib2 | 201.1275521 | 0.526127644 | 0.10807634 | 4.868111234 | 1.13E-06 | 2.20E-05 |
| Cry1 | 69.49382618 | 0.526024471 | 0.141895365 | 3.707129344 | 0.000209622 | 0.001626969 |
| Ttyh2 | 72.81848221 | 0.525634679 | 0.205241901 | 2.561049549 | 0.010435647 | 0.038852329 |
| Cpeb3 | 27.43505213 | 0.52488034 | 0.159088805 | 3.29929149 | 0.000969292 | 0.005734893 |
| Id4 | 236.9502202 | 0.524074528 | 0.12926983 | 4.054113223 | 5.03E-05 | 0.000508377 |
| Lcp1 | 23.36378285 | 0.523922079 | 0.188088881 | 2.785502667 | 0.00534448 | 0.022640872 |
| B230118H07Rik | 194.8426427 | 0.522751898 | 0.098235562 | 5.321412023 | 1.03E-07 | 3.00E-06 |
| Lrrk2 | 53.55001155 | 0.521613329 | 0.13744173 | 3.795159792 | 0.000147548 | 0.001223627 |
| Cd300c2 | 28.81973931 | 0.521372623 | 0.177138684 | 2.943301892 | 0.003247316 | 0.015180709 |
| Eif5b | 177.8215226 | 0.521222104 | 0.130428006 | 3.996243747 | 6.44E-05 | 0.000624079 |
| Hspa4l | 554.8826606 | 0.520863106 | 0.092162864 | 5.651550782 | 1.59E-08 | 6.08E-07 |
| Lrp1b | 36.28950314 | 0.520321977 | 0.162791524 | 3.196247343 | 0.001392277 | 0.007650077 |
| Ctxn2 | 224.5785578 | 0.519107933 | 0.125305108 | 4.142751577 | 3.43E-05 | 0.000369097 |
| Naga | 62.35754547 | 0.518781912 | 0.165391763 | 3.1366853 | 0.001708694 | 0.009065439 |
| Cog7 | 117.2414389 | 0.517262157 | 0.111875089 | 4.62356866 | 3.77E-06 | 6.03E-05 |
| Prpsap2 | 118.9729462 | 0.517234214 | 0.084285063 | 6.136724534 | 8.42E-10 | 5.06E-08 |
| Clic4 | 134.0125894 | 0.516721676 | 0.125344355 | 4.122416801 | 3.75E-05 | 0.000395718 |
| Mest | 75.51144992 | 0.516345534 | 0.124903997 | 4.133939227 | 3.57E-05 | 0.000380408 |
| Adar | 432.0353718 | 0.514516994 | 0.072079366 | 7.138200865 | 9.46E-13 | 1.42E-10 |
| Serpinb9 | 26.19110312 | 0.514252161 | 0.198786196 | 2.586961123 | 0.009682652 | 0.03653843 |
| Tmem184c | 165.1739953 | 0.513950605 | 0.087074916 | 5.90239563 | 3.58E-09 | 1.71E-07 |
| Fam204a | 55.62419448 | 0.513010126 | 0.115861978 | 4.427769454 | 9.52E-06 | 0.000128193 |
| Rpl14 | 3014.50797 | 0.512354487 | 0.112528473 | 4.55310975 | 5.29E-06 | 7.89E-05 |
| Soat1 | 62.16469231 | 0.511973034 | 0.120154618 | 4.260951775 | 2.04E-05 | 0.000239121 |
| Rida | 244.3213114 | 0.511642964 | 0.140308771 | 3.646550101 | 0.000265785 | 0.001972555 |
| P4ha1 | 96.33463235 | 0.51120296 | 0.11925854 | 4.286510287 | 1.82E-05 | 0.000217892 |
| Supt16 | 77.13056347 | 0.510966189 | 0.120607363 | 4.23660857 | 2.27E-05 | 0.000261848 |
| Arpc3 | 986.1444228 | 0.509932107 | 0.103358666 | 4.933617353 | 8.07E-07 | 1.67E-05 |
| P3h4 | 42.21064836 | 0.50991511 | 0.176657789 | 2.886456993 | 0.00389606 | 0.017587779 |
| Ccdc82 | 30.06456663 | 0.509869375 | 0.177605495 | 2.870797299 | 0.00409438 | 0.018306176 |
| Prepl | 100.7777408 | 0.509701104 | 0.095231273 | 5.352245002 | 8.69E-08 | 2.59E-06 |
| Ak4 | 158.6073873 | 0.509560668 | 0.133896566 | 3.805629107 | 0.000141444 | 0.001184595 |
| 2310040G24Rik | 22.86017183 | 0.509526092 | 0.191261142 | 2.664033518 | 0.007720987 | 0.030338074 |
| Etv3 | 52.42539849 | 0.50842844 | 0.123543211 | 4.115389561 | 3.87E-05 | 0.000406704 |

| | | | | | | |
|---|---|---|---|---|---|---|
| Jade1 | 92.91028845 | 0.507840933 | 0.121502321 | 4.179680916 | 2.92E-05 | 0.000322156 |
| Zbtb38 | 157.3005868 | 0.507661382 | 0.086206499 | 5.888899195 | 3.89E-09 | 1.82E-07 |
| Meis2 | 253.4764423 | 0.507235447 | 0.132952758 | 3.815155518 | 0.000136097 | 0.001149469 |
| Tspyl1 | 695.6664201 | 0.50676751 | 0.094758599 | 5.347984421 | 8.89E-08 | 2.64E-06 |
| Asb3 | 44.40751903 | 0.506705226 | 0.13563478 | 3.735806009 | 0.000187115 | 0.001484265 |
| Cenps | 27.04633912 | 0.506030271 | 0.161574922 | 3.131861448 | 0.001737018 | 0.009188073 |
| Rrm2b | 77.60878962 | 0.505214906 | 0.103251375 | 4.893057395 | 9.93E-07 | 1.98E-05 |
| Zwint | 4491.234316 | 0.505151377 | 0.063724147 | 7.927157979 | 2.24E-15 | 6.55E-13 |
| Smad7 | 61.07335157 | 0.504853314 | 0.117250252 | 4.305775929 | 1.66E-05 | 0.000203218 |
| Hprt | 790.1845763 | 0.504173808 | 0.065039354 | 7.751826853 | 9.06E-15 | 2.29E-12 |
| Map7d2 | 438.6879168 | 0.50390295 | 0.063655489 | 7.916095834 | 2.45E-15 | 6.92E-13 |
| Cmc1 | 111.3923996 | 0.503797716 | 0.106321667 | 4.73842945 | 2.15E-06 | 3.79E-05 |
| Scrt1 | 334.2053551 | 0.503447402 | 0.151149579 | 3.330789306 | 0.000866001 | 0.005211071 |
| Sbds | 278.2963586 | 0.503174863 | 0.092418242 | 5.444540489 | 5.19E-08 | 1.69E-06 |
| Siglech | 36.21883109 | 0.50314665 | 0.143696632 | 3.501450533 | 0.000462733 | 0.003115085 |
| Ptk2b | 574.8988015 | 0.502945037 | 0.16315648 | 3.082593088 | 0.002052055 | 0.010484172 |
| Scrn3 | 70.61470831 | 0.502312734 | 0.12478925 | 4.025288521 | 5.69E-05 | 0.000564757 |
| Mettl5 | 63.23682051 | 0.502096755 | 0.111175464 | 4.5162551 | 6.29E-06 | 9.14E-05 |
| Dusp23 | 35.95588058 | 0.501610599 | 0.164608198 | 3.047300219 | 0.00230907 | 0.011526574 |
| Nagpa | 43.70611974 | 0.501251415 | 0.126918584 | 3.949393391 | 7.83E-05 | 0.000729556 |
| Gstm1 | 1101.760962 | 0.501186488 | 0.167061515 | 3.000011634 | 0.002699693 | 0.013112463 |
| Ramac | 380.1437112 | 0.500727709 | 0.069483844 | 7.206390428 | 5.75E-13 | 9.28E-11 |
| Fgfrl1 | 43.7590368 | 0.50063157 | 0.154768996 | 3.234701932 | 0.001217698 | 0.006878106 |
| Rps10 | 4066.531744 | 0.500500522 | 0.116923641 | 4.280575927 | 1.86E-05 | 0.000222524 |
| Phyh | 332.8469909 | 0.50036768 | 0.113015177 | 4.427437922 | 9.54E-06 | 0.000128288 |
| Rpl9 | 5449.088111 | 0.500321863 | 0.111997324 | 4.467266234 | 7.92E-06 | 0.000110078 |
| Mvp | 24.20545289 | 0.500060244 | 0.18597961 | 2.688790689 | 0.007171137 | 0.028548131 |
| Down-regulated genes in old MEC | | | | | | |
| | baseMean | log2FoldChange | lfcSE | stat | pvalue | padj |
| Chrna3 | 11.53362042 | -4.690055067 | 0.916872174 | -5.115276919 | 3.13E-07 | 7.71E-06 |
| Mkrn3 | 15.87954593 | -4.133440997 | 0.755621415 | -5.470253909 | 4.49E-08 | 1.49E-06 |
| 9630013A20Rik | 12.70380968 | -3.884178658 | 0.596902303 | -6.507226787 | 7.66E-11 | 6.36E-09 |
| Eln | 36.99955295 | -3.380604883 | 0.506687204 | -6.671976032 | 2.52E-11 | 2.47E-09 |
| Col3a1 | 17.77648454 | -2.743010997 | 0.655220193 | -4.1863957 | 2.83E-05 | 0.000314002 |
| Gm29521 | 8.943345093 | -2.728758959 | 0.485520496 | -5.620275521 | 1.91E-08 | 7.18E-07 |
| C1qtnf6 | 16.42107373 | -2.577139915 | 0.390984822 | -6.591406542 | 4.36E-11 | 3.89E-09 |
| Cd24a | 38.01706439 | -2.576008926 | 0.378752941 | -6.801290884 | 1.04E-11 | 1.17E-09 |
| Emid1 | 20.3316492 | -2.518195987 | 0.354693124 | -7.09964704 | 1.25E-12 | 1.79E-10 |
| Atp6v1c2 | 9.119994553 | -2.500851535 | 0.471014672 | -5.309498157 | 1.10E-07 | 3.17E-06 |
| Ppic | 13.95618146 | -2.265124802 | 0.417872031 | -5.420618355 | 5.94E-08 | 1.91E-06 |
| Gm43305 | 45.72712575 | -2.222024458 | 0.369358388 | -6.0159036 | 1.79E-09 | 9.54E-08 |
| Robo3 | 23.40804783 | -2.208917124 | 0.349635809 | -6.317765711 | 2.65E-10 | 1.90E-08 |
| Slc29a4 | 30.08396681 | -2.022985342 | 0.275255257 | -7.349488484 | 1.99E-13 | 3.59E-11 |
| Col4a1 | 86.83240045 | -2.004839656 | 0.336889014 | -5.951038978 | 2.66E-09 | 1.34E-07 |
| Gpr17 | 107.79416 | -1.995591575 | 0.315686454 | -6.321435557 | 2.59E-10 | 1.86E-08 |
| Slc38a5 | 8.039008282 | -1.977232547 | 0.443792854 | -4.455305062 | 8.38E-06 | 0.000115358 |
| Nid1 | 15.39046975 | -1.977070734 | 0.385865494 | -5.123730328 | 3.00E-07 | 7.47E-06 |
| Apcdd1 | 54.61659373 | -1.96869676 | 0.263458286 | -7.472517914 | 7.87E-14 | 1.61E-11 |
| Marcksl1 | 837.5591319 | -1.937967343 | 0.385083896 | -5.03258475 | 4.84E-07 | 1.09E-05 |
| Gm47283 | 389.5626559 | -1.910164719 | 0.558154254 | -3.422288205 | 0.000620965 | 0.003979573 |
| Col11a1 | 8.086702884 | -1.860667267 | 0.321675085 | -5.784306447 | 7.28E-09 | 3.07E-07 |
| C030013G03Rik | 10.60591165 | -1.851272583 | 0.364802833 | -5.074720964 | 3.88E-07 | 9.25E-06 |
| 1500035N22Rik | 13.51541705 | -1.772118048 | 0.348294259 | -5.087990972 | 3.62E-07 | 8.74E-06 |
| Dnmt3a | 241.8234805 | -1.763014565 | 0.303495741 | -5.809025723 | 6.28E-09 | 2.72E-07 |
| Wdfy1 | 74.63867611 | -1.749012993 | 0.115714577 | -15.114889 | 1.29E-51 | 2.19E-47 |
| Tet1 | 27.40046757 | -1.746362666 | 0.308237243 | -5.665644581 | 1.46E-08 | 5.67E-07 |
| Gm15690 | 8.732677995 | -1.737491775 | 0.347131512 | -5.005283915 | 5.58E-07 | 1.23E-05 |
| Met | 18.17846764 | -1.735463466 | 0.341491629 | -5.082008808 | 3.73E-07 | 8.99E-06 |
| Dpysl5 | 143.4056448 | -1.731551194 | 0.355341713 | -4.872918469 | 1.10E-06 | 2.16E-05 |
| Lsm11 | 66.24368577 | -1.715781948 | 0.325367247 | -5.273370206 | 1.34E-07 | 3.77E-06 |
| Trim67 | 14.40015885 | -1.69808662 | 0.329687982 | -5.150586952 | 2.60E-07 | 6.62E-06 |
| Col4a2 | 81.19970288 | -1.679464821 | 0.234287396 | -7.168395966 | 7.59E-13 | 1.20E-10 |
| Cxadr | 108.3209442 | -1.637974005 | 0.30080258 | -5.445345594 | 5.17E-08 | 1.69E-06 |
| Col9a3 | 46.42155772 | -1.615122631 | 0.294308025 | -5.487864739 | 4.07E-08 | 1.38E-06 |
| Pla2g3 | 12.52009199 | -1.604132489 | 0.292272828 | -5.488476304 | 4.05E-08 | 1.38E-06 |
| E130006D01Rik | 12.68657731 | -1.596606052 | 0.334532315 | -4.772651195 | 1.82E-06 | 3.29E-05 |
| Nrep | 785.5554182 | -1.594712918 | 0.250362343 | -6.36961972 | 1.89E-10 | 1.41E-08 |
| Cnbd2 | 20.66880177 | -1.588125985 | 0.179865398 | -8.829524763 | 1.05E-18 | 6.60E-16 |
| Enpp6 | 21.95189529 | -1.587597494 | 0.346931339 | -4.576114401 | 4.74E-06 | 7.22E-05 |
| Gp1bb | 43.44393365 | -1.561728561 | 0.250840931 | -6.225971783 | 4.79E-10 | 3.11E-08 |
| Gm3435 | 12.01387833 | -1.547876962 | 0.267383669 | -5.788973455 | 7.08E-09 | 3.02E-07 |
| Adcyap1 | 15.11057168 | -1.547871526 | 0.304638936 | -5.081003582 | 3.75E-07 | 9.03E-06 |
| Nts | 67.93900217 | -1.547222642 | 0.494545607 | -3.128574233 | 0.001756566 | 0.00925569 |

| | | | | | | |
|---|---|---|---|---|---|---|
| Gpx8 | 23.04158289 | -1.535496806 | 0.214712084 | -7.151422385 | 8.59E-13 | 1.31E-10 |
| Gm21860 | 11.91129234 | -1.525817308 | 0.294487472 | -5.181263906 | 2.20E-07 | 5.79E-06 |
| Selenov | 10.08980936 | -1.512524955 | 0.356342999 | -4.244576044 | 2.19E-05 | 0.000253925 |
| Prom1 | 36.09073992 | -1.508029191 | 0.14223432 | -10.60242841 | 2.90E-26 | 4.47E-23 |
| Gdf10 | 67.63205965 | -1.492216957 | 0.231547953 | -6.444526651 | 1.16E-10 | 9.06E-09 |
| Mia | 18.13319385 | -1.488586574 | 0.293234877 | -5.07643084 | 3.85E-07 | 9.20E-06 |
| Hcrtr1 | 26.80109176 | -1.488271173 | 0.326054269 | -4.564489157 | 5.01E-06 | 7.58E-05 |
| Cd248 | 12.69936466 | -1.485409565 | 0.284102344 | -5.22843123 | 1.71E-07 | 4.66E-06 |
| Igsf1 | 17.59495615 | -1.483717892 | 0.236118139 | -6.283794634 | 3.30E-10 | 2.29E-08 |
| AW551984 | 23.58631756 | -1.479204402 | 0.297721197 | -4.968421522 | 6.75E-07 | 1.44E-05 |
| Bglap3 | 94.95281509 | -1.476486945 | 0.299571056 | -4.928670221 | 8.28E-07 | 1.71E-05 |
| Col5a2 | 9.033239228 | -1.444562727 | 0.312589976 | -4.621270154 | 3.81E-06 | 6.08E-05 |
| Gm20033 | 13.75796659 | -1.437346152 | 0.316621327 | -4.53963782 | 5.64E-06 | 8.31E-05 |
| Mpzl1 | 99.85984212 | -1.42837054 | 0.212434165 | -6.723826836 | 1.77E-11 | 1.80E-09 |
| Fam184b | 20.17236307 | -1.424925229 | 0.248621407 | -5.731305477 | 9.97E-09 | 4.08E-07 |
| Cemip | 147.979018 | -1.413846227 | 0.271127742 | -5.214686688 | 1.84E-07 | 4.95E-06 |
| Ncaph | 12.21750223 | -1.410970599 | 0.335826215 | -4.201490341 | 2.65E-05 | 0.000296882 |
| Kcp | 9.678126369 | -1.407153518 | 0.365674991 | -3.848098861 | 0.000119038 | 0.001033187 |
| Plekhg2 | 10.82226274 | -1.403506757 | 0.278752326 | -5.034959812 | 4.78E-07 | 1.08E-05 |
| Dpysl3 | 138.2590564 | -1.400462696 | 0.278058082 | -5.036583317 | 4.74E-07 | 1.07E-05 |
| Tmem229b | 107.7411812 | -1.399511635 | 0.204069535 | -6.858013541 | 6.98E-12 | 8.32E-10 |
| Tnc | 11.03640249 | -1.393240288 | 0.33668346 | -4.138131075 | 3.50E-05 | 0.000375892 |
| Gnb4 | 63.46329783 | -1.392974887 | 0.218238468 | -6.382810954 | 1.74E-10 | 1.30E-08 |
| Gabra2 | 223.930885 | -1.392907102 | 0.120345113 | -11.57427224 | 5.56E-31 | 1.57E-27 |
| Lhx1os | 23.99716567 | -1.378418585 | 0.344130079 | -4.00551614 | 6.19E-05 | 0.000606338 |
| Col22a1 | 12.19934365 | -1.372234857 | 0.315495608 | -4.349457872 | 1.36E-05 | 0.000170981 |
| Ism1 | 13.09067954 | -1.369277364 | 0.300484344 | -4.556900853 | 5.19E-06 | 7.80E-05 |
| Mfsd2a | 113.0219066 | -1.358883425 | 0.149136584 | -9.111670574 | 8.11E-20 | 5.98E-17 |
| Aspdh | 9.232091923 | -1.356875979 | 0.268797342 | -5.047951621 | 4.47E-07 | 1.02E-05 |
| Gm15627 | 8.774549131 | -1.353555416 | 0.267138462 | -5.066868337 | 4.04E-07 | 9.48E-06 |
| Col26a1 | 23.49065135 | -1.352745079 | 0.270896084 | -4.993594072 | 5.93E-07 | 1.29E-05 |
| Lpl | 43.09655668 | -1.350327942 | 0.382096507 | -3.533997084 | 0.000409326 | 0.002813657 |
| Megf6 | 9.844989606 | -1.348453784 | 0.374670566 | -3.599038479 | 0.000319396 | 0.002295074 |
| Col18a1 | 22.82306105 | -1.330295692 | 0.281354255 | -4.728187563 | 2.27E-06 | 3.95E-05 |
| Rras2 | 62.95352397 | -1.305067968 | 0.220177015 | -5.927357883 | 3.08E-09 | 1.51E-07 |
| Nid2 | 9.460519112 | -1.303514008 | 0.38610693 | -3.376044062 | 0.000735361 | 0.00454843 |
| AI506816 | 15.86954562 | -1.297884531 | 0.270590378 | -4.796491805 | 1.61E-06 | 2.97E-05 |
| Col5a1 | 36.9291506 | -1.297314795 | 0.293030433 | -4.427235704 | 9.54E-06 | 0.000128307 |
| Kif21b | 197.9955635 | -1.286161796 | 0.276294022 | -4.655047507 | 3.24E-06 | 5.32E-05 |
| Ctsk | 18.97176416 | -1.283576873 | 0.313590702 | -4.093159862 | 4.26E-05 | 0.000442529 |
| Cd38 | 15.0384224 | -1.280014612 | 0.282582016 | -4.529710104 | 5.91E-06 | 8.67E-05 |
| Col1a2 | 59.45274699 | -1.272339717 | 0.217564726 | -5.848097437 | 4.97E-09 | 2.22E-07 |
| Dcx | 55.10365419 | -1.267339076 | 0.234061167 | -5.414563613 | 6.14E-08 | 1.96E-06 |
| Pcp2 | 385.4694087 | -1.265509328 | 0.36454855 | -3.471442498 | 0.00051767 | 0.003405133 |
| Gjc1 | 13.81445075 | -1.264133682 | 0.330824611 | -3.821159738 | 0.000132826 | 0.001126326 |
| Ap1g2 | 20.13651837 | -1.257485602 | 0.303831335 | -4.138762052 | 3.49E-05 | 0.000375097 |
| Slc26a6 | 26.77812451 | -1.257469739 | 0.168551819 | -7.460434123 | 8.62E-14 | 1.74E-11 |
| Lama4 | 15.00092845 | -1.25610148 | 0.272624832 | -4.607436058 | 4.08E-06 | 6.42E-05 |
| Bmp1 | 70.86391157 | -1.246289007 | 0.169418281 | -7.356284111 | 1.89E-13 | 3.46E-11 |
| Col1a1 | 32.56335448 | -1.244347678 | 0.258739414 | -4.809269917 | 1.51E-06 | 2.81E-05 |
| Rnf122 | 10.16043215 | -1.241912429 | 0.289048898 | -4.29654787 | 1.73E-05 | 0.000209983 |
| Grin2d | 51.31498593 | -1.237719116 | 0.192961623 | -6.414327857 | 1.41E-10 | 1.08E-08 |
| Hmgcs2 | 36.54573951 | -1.230375204 | 0.2889746 | -4.257727847 | 2.07E-05 | 0.000241757 |
| Hrk | 100.6356743 | -1.225169862 | 0.21096405 | -5.8074817 | 6.34E-09 | 2.74E-07 |
| Cnpy1 | 15.83161897 | -1.223687638 | 0.357913105 | -3.418951756 | 0.000628629 | 0.004021089 |
| Traf4 | 41.23177584 | -1.219414472 | 0.216518388 | -5.631921077 | 1.78E-08 | 6.76E-07 |
| Ncan | 421.8171209 | -1.214761938 | 0.186785148 | -6.50352532 | 7.85E-11 | 6.46E-09 |
| Ankrd23 | 9.551139973 | -1.213159795 | 0.264086701 | -4.5937936 | 4.35E-06 | 6.70E-05 |
| Dhrs13 | 25.70335346 | -1.210536451 | 0.230965047 | -5.241210591 | 1.60E-07 | 4.38E-06 |
| Cenpa | 21.9127765 | -1.210176777 | 0.263267981 | -4.596748811 | 4.29E-06 | 6.66E-05 |
| Kmt5c | 30.63028931 | -1.208545117 | 0.192064525 | -6.292391154 | 3.13E-10 | 2.21E-08 |
| Ccl17 | 29.03245339 | -1.189324855 | 0.271172207 | -4.385865606 | 1.16E-05 | 0.000149486 |
| Hspg2 | 13.99337785 | -1.182376999 | 0.367235666 | -3.219668212 | 0.00128339 | 0.007183559 |
| Elovl2 | 41.71938056 | -1.180210766 | 0.275577436 | -4.282682885 | 1.85E-05 | 0.000221049 |
| Penk | 133.9838136 | -1.180165007 | 0.151738105 | -7.777644277 | 7.39E-15 | 1.93E-12 |
| AU021092 | 11.6241543 | -1.176371819 | 0.296242529 | -3.970975486 | 7.16E-05 | 0.00068012 |
| Nptx2 | 63.05468958 | -1.171355088 | 0.204292163 | -5.733725026 | 9.82E-09 | 4.03E-07 |
| Lama1 | 8.103792634 | -1.168050765 | 0.310028124 | -3.76756389 | 0.000164848 | 0.001342789 |
| Dll4 | 8.19463865 | -1.167589946 | 0.3525761 | -3.311596971 | 0.000927651 | 0.005519343 |
| Clmp | 35.0957874 | -1.166029269 | 0.213624726 | -5.45830667 | 4.81E-08 | 1.58E-06 |
| Muc3a | 15.89369383 | -1.162617102 | 0.242696786 | -4.790409967 | 1.66E-06 | 3.06E-05 |
| Crybb1 | 36.50124042 | -1.162289777 | 0.247131074 | -4.703130853 | 2.56E-06 | 4.39E-05 |
| Egflam | 9.698278966 | -1.161523459 | 0.283931282 | -4.090861179 | 4.30E-05 | 0.000445845 |

| | | | | | | |
|---|---|---|---|---|---|---|
| Vav2 | 22.62335405 | -1.155187769 | 0.249087392 | -4.637680619 | 3.52E-06 | 5.70E-05 |
| Zfp41 | 22.6387277 | -1.148665767 | 0.243409664 | -4.71906393 | 2.37E-06 | 4.12E-05 |
| Rgs11 | 76.65428293 | -1.141843687 | 0.312833994 | -3.649998752 | 0.000262242 | 0.00195053 |
| 2410018L13Rik | 8.527894684 | -1.134483461 | 0.314851783 | -3.603230221 | 0.000314287 | 0.002267012 |
| Gm27004 | 27.80398682 | -1.133566102 | 0.238150759 | -4.759867686 | 1.94E-06 | 3.47E-05 |
| Gm20501 | 22.94636485 | -1.131563985 | 0.257574677 | -4.393149191 | 1.12E-05 | 0.000145899 |
| Adam19 | 37.32295537 | -1.131417122 | 0.188720403 | -5.995202991 | 2.03E-09 | 1.07E-07 |
| Dera | 8.861054158 | -1.130443928 | 0.335027557 | -3.374181934 | 0.000740354 | 0.004571856 |
| Tmem163 | 69.00020862 | -1.130308615 | 0.153627903 | -7.357443497 | 1.87E-13 | 3.46E-11 |
| Gpr4 | 12.98213354 | -1.120131665 | 0.248763539 | -4.502796781 | 6.71E-06 | 9.62E-05 |
| Arhgef33 | 14.10128022 | -1.118766261 | 0.232259587 | -4.81687871 | 1.46E-06 | 2.72E-05 |
| Pde3a | 8.692018426 | -1.116981608 | 0.31897875 | -3.50174301 | 0.000462225 | 0.003112904 |
| Cyp26b1 | 20.30537911 | -1.108741992 | 0.219917042 | -5.041637437 | 4.62E-07 | 1.05E-05 |
| Ptpro | 60.42413014 | -1.105080892 | 0.16721007 | -6.608937424 | 3.87E-11 | 3.57E-09 |
| Cspg4 | 49.96616196 | -1.098635166 | 0.177033774 | -6.205794198 | 5.44E-10 | 3.46E-08 |
| Dynlt1f | 19.72022707 | -1.092836437 | 0.276131823 | -3.957662044 | 7.57E-05 | 0.000712365 |
| Nav1 | 185.3745695 | -1.085047557 | 0.201270168 | -5.391000416 | 7.01E-08 | 2.17E-06 |
| Gm34466 | 71.38580131 | -1.083206563 | 0.178971465 | -6.052398164 | 1.43E-09 | 7.91E-08 |
| Fbxw9 | 88.99986811 | -1.080314246 | 0.195208287 | -5.53416182 | 3.13E-08 | 1.10E-06 |
| Tie1 | 16.43859832 | -1.077555076 | 0.265286627 | -4.061852232 | 4.87E-05 | 0.000496246 |
| Zic4 | 9.629984941 | -1.073509748 | 0.281907814 | -3.808017004 | 0.000140086 | 0.001177289 |
| Insyn2b | 10.1849998 | -1.067585482 | 0.354020126 | -3.015606753 | 0.002564656 | 0.012590273 |
| Pip5kl1 | 21.97951071 | -1.066590936 | 0.203044544 | -5.252989892 | 1.50E-07 | 4.13E-06 |
| 04.Mar | 49.45509418 | -1.062058082 | 0.200907559 | -5.286302264 | 1.25E-07 | 3.53E-06 |
| Eva1b | 23.31030942 | -1.062012495 | 0.252789263 | -4.201177229 | 2.66E-05 | 0.000297096 |
| Gja4 | 14.93546241 | -1.061245098 | 0.233326774 | -4.548321138 | 5.41E-06 | 8.03E-05 |
| Gm17750 | 70.93135172 | -1.05229849 | 0.194588297 | -5.407820034 | 6.38E-08 | 2.01E-06 |
| Notch4 | 18.49121566 | -1.049421047 | 0.207535241 | -5.05659204 | 4.27E-07 | 9.88E-06 |
| Zfp282 | 70.24953774 | -1.043731601 | 0.185120513 | -5.638119658 | 1.72E-08 | 6.55E-07 |
| Inha | 127.1962679 | -1.042111644 | 0.145508325 | -7.161869541 | 7.96E-13 | 1.23E-10 |
| Fam71e1 | 44.94842655 | -1.039794806 | 0.189925646 | -5.47474671 | 4.38E-08 | 1.46E-06 |
| Ppp1r14c | 112.7223146 | -1.035238489 | 0.225092185 | -4.599175619 | 4.24E-06 | 6.61E-05 |
| Sppl2b | 147.1772336 | -1.033662087 | 0.153371226 | -6.739608958 | 1.59E-11 | 1.64E-09 |
| Thsd1 | 9.646745652 | -1.028345791 | 0.275281779 | -3.735611536 | 0.00018726 | 0.001484676 |
| Erc1 | 133.0830645 | -1.027694509 | 0.207615173 | -4.949997117 | 7.42E-07 | 1.55E-05 |
| Neu4 | 25.14695425 | -1.027660187 | 0.185116922 | -5.551411375 | 2.83E-08 | 1.01E-06 |
| Grap | 13.06794587 | -1.024255317 | 0.241819178 | -4.235624841 | 2.28E-05 | 0.000262557 |
| Npnt | 34.58941645 | -1.023061602 | 0.233070433 | -4.389495441 | 1.14E-05 | 0.000147689 |
| Brwd3 | 15.28911546 | -1.018401372 | 0.220472529 | -4.6191758 | 3.85E-06 | 6.13E-05 |
| Tbc1d16 | 92.975041 | -1.017309863 | 0.168405935 | -6.040819546 | 1.53E-09 | 8.38E-08 |
| Flna | 114.0164288 | -1.01657561 | 0.131821748 | -7.711744276 | 1.24E-14 | 3.09E-12 |
| Cd93 | 14.43915846 | -1.016328803 | 0.3132333 | -3.244638432 | 0.001175998 | 0.006687845 |
| Tcap | 34.20304643 | -1.014556426 | 0.218153668 | -4.650650325 | 3.31E-06 | 5.39E-05 |
| D330050I16Rik | 11.12727819 | -1.013413258 | 0.231004133 | -4.38699188 | 1.15E-05 | 0.000148828 |
| Rem2 | 36.15289237 | -1.012265935 | 0.236924917 | -4.272517849 | 1.93E-05 | 0.000229295 |
| Mmp15 | 40.44360396 | -1.005413779 | 0.170415018 | -5.899795619 | 3.64E-09 | 1.72E-07 |
| Col16a1 | 41.53142961 | -1.005292292 | 0.200461249 | -5.014895882 | 5.31E-07 | 1.17E-05 |
| Serpinh1 | 70.65521997 | -1.003569818 | 0.215715797 | -4.652277816 | 3.28E-06 | 5.36E-05 |
| Mrc1 | 10.97244542 | -1.002693328 | 0.256225873 | -3.913318024 | 9.10E-05 | 0.000826103 |
| Ccdc28b | 151.6609014 | -1.002437457 | 0.122364619 | -8.192216548 | 2.56E-16 | 9.66E-14 |
| Etv4 | 25.22434339 | -1.000303935 | 0.198595242 | -5.036897796 | 4.73E-07 | 1.07E-05 |
| Vcan | 22.70572375 | -1.000111522 | 0.213108471 | -4.692969357 | 2.69E-06 | 4.57E-05 |
| Nav3 | 226.7386043 | -0.998688284 | 0.196831394 | -5.073826191 | 3.90E-07 | 9.26E-06 |
| Slc39a8 | 11.16857499 | -0.998355985 | 0.337021424 | -2.962292343 | 0.003053577 | 0.014442295 |
| Ctla2a | 53.8959707 | -0.997862364 | 0.185976273 | -5.365535875 | 8.07E-08 | 2.44E-06 |
| Ushbp1 | 8.190348221 | -0.996115577 | 0.296717483 | -3.357117915 | 0.000787595 | 0.004821425 |
| Dynlt1a | 49.95406534 | -0.995094001 | 0.350701624 | -2.837437678 | 0.004547722 | 0.019940104 |
| Ndst4 | 10.85231367 | -0.99478916 | 0.235137152 | -4.230676235 | 2.33E-05 | 0.000267575 |
| Igf2 | 141.831784 | -0.992797988 | 0.315687508 | -3.14487575 | 0.001661574 | 0.00885424 |
| Afap1 | 63.48336512 | -0.992345008 | 0.194876858 | -5.092164458 | 3.54E-07 | 8.58E-06 |
| Robo4 | 13.11870419 | -0.9899634 | 0.235922385 | -4.196140186 | 2.72E-05 | 0.000302182 |
| AU022754 | 8.416201944 | -0.989263712 | 0.278732907 | -3.549145746 | 0.000386483 | 0.002686152 |
| Proscos | 10.65671658 | -0.988780536 | 0.246270058 | -4.015025396 | 5.94E-05 | 0.000585789 |
| Hif3a | 12.6252882 | -0.988269783 | 0.31358751 | -3.151495998 | 0.001624364 | 0.008680515 |
| Npff | 8.446801557 | -0.987102595 | 0.31086135 | -3.175378969 | 0.001496409 | 0.008111811 |
| Faap100 | 68.157883 | -0.986663207 | 0.194032948 | -5.0850292 | 3.68E-07 | 8.86E-06 |
| Zic1 | 105.1038203 | -0.985828778 | 0.17990287 | -5.479783488 | 4.26E-08 | 1.43E-06 |
| Sema5a | 55.83054781 | -0.982210628 | 0.189464852 | -5.184131088 | 2.17E-07 | 5.72E-06 |
| Traf3 | 124.4592134 | -0.980725969 | 0.181273076 | -5.410213085 | 6.29E-08 | 2.00E-06 |
| Spef1 | 51.42781639 | -0.979645527 | 0.155698919 | -6.291922463 | 3.14E-10 | 2.21E-08 |
| Sema4g | 115.8563783 | -0.978130569 | 0.154731057 | -6.321488326 | 2.59E-10 | 1.86E-08 |
| Jak3 | 11.39213209 | -0.976252741 | 0.279019888 | -3.49886436 | 0.000467244 | 0.003139222 |
| Dpysl4 | 290.1438184 | -0.975603548 | 0.146361036 | -6.665732716 | 2.63E-11 | 2.57E-09 |

| | | | | | | |
|---|---|---|---|---|---|---|
| Calb2 | 187.4371111 | -0.971449073 | 0.249337008 | -3.896128694 | 9.77E-05 | 0.00087478 |
| Meg3 | 3526.170776 | -0.969380016 | 0.23538093 | -4.118345591 | 3.82E-05 | 0.000402272 |
| Gpx7 | 32.57533878 | -0.966294337 | 0.187194926 | -5.16196863 | 2.44E-07 | 6.28E-06 |
| Sap25 | 24.02598784 | -0.966232496 | 0.206384977 | -4.681699755 | 2.85E-06 | 4.79E-05 |
| Fmo1 | 19.49532084 | -0.963161257 | 0.224247637 | -4.295078733 | 1.75E-05 | 0.000210989 |
| Iglon5 | 49.34793642 | -0.960677663 | 0.166089856 | -5.784083897 | 7.29E-09 | 3.07E-07 |
| Grm4 | 129.8731413 | -0.960034114 | 0.14802863 | -6.48546239 | 8.85E-11 | 7.14E-09 |
| Miat | 357.549306 | -0.957307367 | 0.193893215 | -4.93729172 | 7.92E-07 | 1.64E-05 |
| Angptl4 | 22.85412459 | -0.956642518 | 0.238040674 | -4.018819564 | 5.85E-05 | 0.000577651 |
| Slco1c1 | 122.8137691 | -0.956556896 | 0.17805786 | -5.372168903 | 7.78E-08 | 2.36E-06 |
| Myt1 | 21.66498397 | -0.956200282 | 0.186020434 | -5.140297006 | 2.74E-07 | 6.93E-06 |
| Nanos3 | 35.54530796 | -0.954428926 | 0.181362377 | -5.262551917 | 1.42E-07 | 3.96E-06 |
| Csrp2 | 35.97159445 | -0.951947148 | 0.214132905 | -4.445590215 | 8.77E-06 | 0.000120111 |
| Esam | 54.78621649 | -0.951063085 | 0.165966979 | -5.730435608 | 1.00E-08 | 4.08E-07 |
| Nnmt | 11.12574407 | -0.949568857 | 0.28616029 | -3.318311067 | 0.000905636 | 0.005414076 |
| Rflnb | 31.10380384 | -0.946418462 | 0.210261268 | -4.501154548 | 6.76E-06 | 9.67E-05 |
| Rhbdl1 | 418.3439327 | -0.945715842 | 0.133094222 | -7.105611565 | 1.20E-12 | 1.75E-10 |
| Dot1l | 107.1042506 | -0.945505733 | 0.209821767 | -4.506232819 | 6.60E-06 | 9.50E-05 |
| Morn1 | 11.9918661 | -0.944261264 | 0.234651639 | -4.024098305 | 5.72E-05 | 0.000566775 |
| Syndig1l | 57.77223334 | -0.943403409 | 0.258200546 | -3.65376226 | 0.000258426 | 0.001928918 |
| Pdgfra | 96.16288132 | -0.943246281 | 0.152528241 | -6.184076289 | 6.25E-10 | 3.94E-08 |
| Igsf3 | 28.28513157 | -0.942860508 | 0.261345894 | -3.607711192 | 0.00030891 | 0.002238707 |
| Zfp46 | 78.95288613 | -0.942084885 | 0.178721418 | -5.271247819 | 1.35E-07 | 3.80E-06 |
| Car8 | 158.8277777 | -0.941181133 | 0.261348584 | -3.601248257 | 0.000316693 | 0.002281455 |
| Emilin1 | 9.627842636 | -0.939416428 | 0.270135744 | -3.477571742 | 0.000505978 | 0.003342433 |
| Tet3 | 69.00246288 | -0.937666348 | 0.197413216 | -4.749764806 | 2.04E-06 | 3.62E-05 |
| Ccdc3 | 39.82041889 | -0.935307875 | 0.212618198 | -4.399001977 | 1.09E-05 | 0.00014279 |
| Smim3 | 37.76921918 | -0.931430684 | 0.190276879 | -4.895133301 | 9.82E-07 | 1.96E-05 |
| Gm48045 | 9.741397076 | -0.930801095 | 0.285281666 | -3.262744178 | 0.001103391 | 0.006378059 |
| Cwh43 | 19.84206661 | -0.930546297 | 0.219484858 | -4.239683351 | 2.24E-05 | 0.000258992 |
| Slc17a8 | 9.280540416 | -0.929699749 | 0.364376697 | -2.55147971 | 0.010726657 | 0.039639756 |
| Adamts10 | 49.20130162 | -0.928509558 | 0.197392644 | -4.703871116 | 2.55E-06 | 4.38E-05 |
| Hacd1 | 99.17070228 | -0.928132113 | 0.169989348 | -5.459942787 | 4.76E-08 | 1.57E-06 |
| Ppil6 | 15.90233113 | -0.920963632 | 0.197284377 | -4.668203555 | 3.04E-06 | 5.05E-05 |
| Dcakd | 175.2242984 | -0.918242634 | 0.129165317 | -7.109049539 | 1.17E-12 | 1.72E-10 |
| Itga1 | 17.50290831 | -0.916964855 | 0.268930284 | -3.409674963 | 0.000650403 | 0.004133854 |
| Ebf4 | 22.91125917 | -0.91540793 | 0.18498004 | -4.948684891 | 7.47E-07 | 1.56E-05 |
| Notum | 9.246269147 | -0.906387614 | 0.334983207 | -2.705770299 | 0.006814616 | 0.0274186 |
| Tmem252 | 9.230669416 | -0.906250109 | 0.334620225 | -2.708294478 | 0.006762998 | 0.027223839 |
| Fabp7 | 208.4794683 | -0.902619696 | 0.332081415 | -2.718067487 | 0.006566444 | 0.026618363 |
| Dok4 | 82.30595084 | -0.90229537 | 0.194803557 | -4.631821834 | 3.62E-06 | 5.85E-05 |
| Ifitm1 | 37.46301589 | -0.901028551 | 0.282512191 | -3.189343963 | 0.001425961 | 0.007807321 |
| Kcnk10 | 22.3079904 | -0.897756703 | 0.213829674 | -4.198466408 | 2.69E-05 | 0.000299883 |
| Shisa6 | 51.79688574 | -0.897632767 | 0.138661567 | -6.473551289 | 9.57E-11 | 7.69E-09 |
| Srarp | 12.85717454 | -0.89761144 | 0.239094752 | -3.754208036 | 0.00017389 | 0.001403659 |
| Parvb | 45.92917801 | -0.895420202 | 0.228893977 | -3.91194305 | 9.16E-05 | 0.000828883 |
| Rab34 | 73.08125164 | -0.893762789 | 0.149505285 | -5.978135061 | 2.26E-09 | 1.17E-07 |
| Zic5 | 11.17565214 | -0.889715204 | 0.24739601 | -3.596319942 | 0.000322751 | 0.002313298 |
| Ezh2 | 11.84021355 | -0.889566327 | 0.247554637 | -3.593414121 | 0.000326373 | 0.002334326 |
| C1ql1 | 63.28792715 | -0.888885056 | 0.150633687 | -5.900971257 | 3.61E-09 | 1.72E-07 |
| Thsd4 | 19.61548524 | -0.88613395 | 0.193691871 | -4.574967174 | 4.76E-06 | 7.23E-05 |
| Zbtb5 | 28.17877014 | -0.885304207 | 0.164654734 | -5.376730973 | 7.59E-08 | 2.31E-06 |
| Stk36 | 9.787749025 | -0.884540256 | 0.250711028 | -3.52812663 | 0.000418512 | 0.002864027 |
| Tubb2b | 613.5537071 | -0.884054634 | 0.231080433 | -3.825744236 | 0.000130378 | 0.001112805 |
| N4bp2 | 13.00120549 | -0.882903539 | 0.225898466 | -3.908408735 | 9.29E-05 | 0.000839025 |
| Cacna1h | 69.06099845 | -0.882122313 | 0.141129184 | -6.250459939 | 4.09E-10 | 2.74E-08 |
| Cryga | 11.73491682 | -0.880723494 | 0.243591791 | -3.615571328 | 0.000299686 | 0.002183058 |
| Fras1 | 14.71856036 | -0.879954408 | 0.278096381 | -3.164206608 | 0.001555064 | 0.008373533 |
| Itm2a | 207.1598944 | -0.877672259 | 0.096450881 | -9.099681067 | 9.06E-20 | 6.40E-17 |
| Arap3 | 11.07955512 | -0.875177062 | 0.296706865 | -2.949635367 | 0.003181492 | 0.014955481 |
| 1700016K19Rik | 12.30301738 | -0.872876338 | 0.290298533 | -3.006823115 | 0.002639933 | 0.012877557 |
| Eme2 | 31.51791821 | -0.872808317 | 0.155450629 | -5.614697872 | 1.97E-08 | 7.37E-07 |
| Gm35339 | 8.576799658 | -0.872065518 | 0.267169878 | -3.264086218 | 0.001098177 | 0.006357652 |
| Lefty2 | 27.78953012 | -0.871666714 | 0.235198884 | -3.706083542 | 0.000210489 | 0.001629967 |
| Atp13a5 | 21.06039184 | -0.870563629 | 0.195883609 | -4.444290337 | 8.82E-06 | 0.000120742 |
| Cntn6 | 36.49798386 | -0.870314329 | 0.215119721 | -4.045720791 | 5.22E-05 | 0.000522888 |
| Neto1 | 260.4512246 | -0.868342521 | 0.148875468 | -5.832677016 | 5.45E-09 | 2.41E-07 |
| Fn1 | 192.0067322 | -0.867716022 | 0.183230307 | -4.73565774 | 2.18E-06 | 3.83E-05 |
| Slc27a3 | 15.61153541 | -0.867229123 | 0.234218756 | -3.702645933 | 0.000213363 | 0.001647703 |
| Deaf1 | 248.3996597 | -0.865657553 | 0.185538848 | -4.665640455 | 3.08E-06 | 5.10E-05 |
| Kif26a | 9.016532609 | -0.863490422 | 0.316058878 | -2.732055582 | 0.006294052 | 0.025746809 |
| Hapln1 | 38.27714878 | -0.863122185 | 0.258473718 | -3.33930348 | 0.000839888 | 0.005079177 |
| Sdk1 | 12.38468891 | -0.862549894 | 0.272241595 | -3.168325155 | 0.001533199 | 0.008282111 |

| | | | | | | |
|---|---|---|---|---|---|---|
| Pkp2 | 19.59542998 | -0.862217028 | 0.246504024 | -3.497780737 | 0.000469147 | 0.003148434 |
| Nkiras2 | 49.26739221 | -0.86220013 | 0.205072446 | -4.20436848 | 2.62E-05 | 0.000293711 |
| Gm5577 | 13.10136793 | -0.861624693 | 0.273575168 | -3.149498904 | 0.001635507 | 0.008731806 |
| Shisal1 | 58.82519852 | -0.8615302 | 0.132564828 | -6.498935003 | 8.09E-11 | 6.62E-09 |
| Gm3764 | 207.7505715 | -0.861007638 | 0.178396215 | -4.82637839 | 1.39E-06 | 2.62E-05 |
| Clec2l | 391.7921631 | -0.86069091 | 0.132634658 | -6.489185559 | 8.63E-11 | 7.00E-09 |
| Slc16a1 | 148.1834668 | -0.860024811 | 0.306010581 | -2.810441415 | 0.00494736 | 0.021306578 |
| Tmem132a | 938.861961 | -0.858956021 | 0.126171006 | -6.807871676 | 9.91E-12 | 1.13E-09 |
| Mirt1 | 21.49729225 | -0.857485605 | 0.195657847 | -4.382577128 | 1.17E-05 | 0.000150955 |
| Car15 | 29.19171297 | -0.857107073 | 0.138943124 | -6.168762073 | 6.88E-10 | 4.29E-08 |
| Ccm2l | 10.41151216 | -0.856674249 | 0.249626015 | -3.431830807 | 0.000599521 | 0.003853807 |
| Ccnb1 | 16.32765381 | -0.85621347 | 0.263293855 | -3.251931079 | 0.001146238 | 0.00657196 |
| Adamtsl2 | 29.19817652 | -0.855146546 | 0.237685066 | -3.597813529 | 0.000320904 | 0.002303954 |
| Sema6a | 72.261126 | -0.85325227 | 0.194333869 | -4.390651371 | 1.13E-05 | 0.000147132 |
| Spata2l | 206.1595025 | -0.852978746 | 0.157381659 | -5.419810361 | 5.97E-08 | 1.91E-06 |
| Pus7l | 9.85403213 | -0.852783701 | 0.247651342 | -3.443485081 | 0.000574268 | 0.003722532 |
| Snhg9 | 296.2617045 | -0.85048877 | 0.130077831 | -6.538306841 | 6.22E-11 | 5.35E-09 |
| Gm3488 | 8.908084136 | -0.85021128 | 0.294175594 | -2.890148936 | 0.003850593 | 0.017405709 |
| Gm6569 | 12.1672686 | -0.850019186 | 0.244564435 | -3.475645123 | 0.000509626 | 0.003361351 |
| Igsf9b | 173.8564905 | -0.849657637 | 0.182600002 | -4.653108583 | 3.27E-06 | 5.35E-05 |
| Cldn5 | 188.0697595 | -0.845901496 | 0.168702325 | -5.014166196 | 5.33E-07 | 1.18E-05 |
| A230057D06Rik | 29.62117281 | -0.845461259 | 0.205858792 | -4.106996117 | 4.01E-05 | 0.000419679 |
| A430027H14Rik | 11.07139868 | -0.845215581 | 0.257303874 | -3.284892555 | 0.001020213 | 0.005986025 |
| 2310002F09Rik | 10.2882307 | -0.844100531 | 0.274978523 | -3.069696217 | 0.002142766 | 0.010823011 |
| Bdh1 | 255.1009171 | -0.840342308 | 0.179531634 | -4.680747849 | 2.86E-06 | 4.80E-05 |
| Fam83h | 8.62135854 | -0.838521119 | 0.272188195 | -3.080666744 | 0.002065377 | 0.010535721 |
| Sox12 | 53.74398957 | -0.837659205 | 0.128602613 | -6.51354731 | 7.34E-11 | 6.16E-09 |
| Gm27032 | 97.2684022 | -0.836744268 | 0.197463241 | -4.237468528 | 2.26E-05 | 0.000261025 |
| Rimklb | 97.47311951 | -0.836556678 | 0.145632154 | -5.744313031 | 9.23E-09 | 3.82E-07 |
| Spink10 | 56.0919219 | -0.834012684 | 0.12771723 | -6.530150093 | 6.57E-11 | 5.62E-09 |
| Catsperz | 39.14605742 | -0.833508488 | 0.147414631 | -5.6541775 | 1.57E-08 | 6.02E-07 |
| Ttyh3 | 471.6411013 | -0.833187419 | 0.157061414 | -5.304851129 | 1.13E-07 | 3.24E-06 |
| Tmem169 | 56.88777909 | -0.831011018 | 0.181748909 | -4.572302634 | 4.82E-06 | 7.31E-05 |
| Fam57b | 334.41199 | -0.828263678 | 0.12263946 | -6.753647479 | 1.44E-11 | 1.53E-09 |
| Arhgef1 | 96.34329887 | -0.825473002 | 0.168535495 | -4.8979178 | 9.69E-07 | 1.95E-05 |
| Leng8 | 120.2187684 | -0.823783808 | 0.215089914 | -3.829950899 | 0.000128169 | 0.001097268 |
| Kdr | 28.92623403 | -0.823031686 | 0.292272251 | -2.815976147 | 0.004862928 | 0.021007005 |
| Ddah2 | 70.5919402 | -0.822465172 | 0.159310499 | -5.162655169 | 2.43E-07 | 6.26E-06 |
| Fbxl12os | 9.556209768 | -0.820905786 | 0.264772459 | -3.100419845 | 0.001932465 | 0.009980632 |
| Pcdh18 | 14.90337711 | -0.819039438 | 0.211935339 | -3.864572293 | 0.000111284 | 0.000975375 |
| Ackr1 | 168.0062598 | -0.818813399 | 0.153771115 | -5.324884325 | 1.01E-07 | 2.96E-06 |
| A330102I10Rik | 23.81595798 | -0.817890371 | 0.188017061 | -4.350085922 | 1.36E-05 | 0.000170727 |
| Gm11627 | 33.45920432 | -0.81760937 | 0.151314921 | -5.403362503 | 6.54E-08 | 2.05E-06 |
| Abcc9 | 12.0673745 | -0.816978256 | 0.240563417 | -3.396103478 | 0.000683525 | 0.004299234 |
| Gm10244 | 10.07862901 | -0.816091135 | 0.240729225 | -3.390079191 | 0.000698724 | 0.004380206 |
| Plppr1 | 35.27695271 | -0.815371184 | 0.22307253 | -3.655184181 | 0.000256997 | 0.001919948 |
| 2810029C07Rik | 25.82199015 | -0.814958779 | 0.186657619 | -4.366062216 | 1.27E-05 | 0.000161211 |
| Dab2 | 24.30508887 | -0.814916547 | 0.193349483 | -4.214733514 | 2.50E-05 | 0.000282788 |
| Catsper2 | 8.307659932 | -0.81481438 | 0.254843585 | -3.19731172 | 0.001387149 | 0.007626847 |
| Ryr1 | 12.78564125 | -0.812460422 | 0.249805537 | -3.25237155 | 0.001144463 | 0.006565074 |
| Mycn | 31.39065839 | -0.810774032 | 0.192081002 | -4.221000632 | 2.43E-05 | 0.000276329 |
| Col11a2 | 25.0156901 | -0.808327635 | 0.23887386 | -3.38390996 | 0.000714614 | 0.004453324 |
| Slc16a7 | 84.84593567 | -0.807540053 | 0.154520264 | -5.226111019 | 1.73E-07 | 4.71E-06 |
| Pald1 | 21.96362113 | -0.805522558 | 0.214886112 | -3.748602226 | 0.000177823 | 0.00142789 |
| Neurod1 | 94.7216568 | -0.805357968 | 0.135451663 | -5.945722268 | 2.75E-09 | 1.38E-07 |
| 4933439C10Rik | 67.28394811 | -0.803262003 | 0.196879256 | -4.079972765 | 4.50E-05 | 0.000462721 |
| Slc16a2 | 95.172482 | -0.803186851 | 0.181503682 | -4.425182135 | 9.64E-06 | 0.000129226 |
| Hist3h2ba | 288.0221245 | -0.801388162 | 0.11652901 | -6.877155838 | 6.11E-12 | 7.45E-10 |
| Mapkbp1 | 101.5324169 | -0.80055806 | 0.115330019 | -6.941454329 | 3.88E-12 | 4.95E-10 |
| Tmem181a | 225.199988 | -0.799498779 | 0.109151964 | -7.324639417 | 2.40E-13 | 4.23E-11 |
| Meak7 | 30.58401584 | -0.799210711 | 0.167558129 | -4.769751948 | 1.84E-06 | 3.33E-05 |
| Nasp | 70.15136112 | -0.799127582 | 0.120559715 | -6.628479371 | 3.39E-11 | 3.18E-09 |
| Tubd1 | 18.05909548 | -0.79763025 | 0.23654684 | -3.371975928 | 0.00074631 | 0.004601926 |
| 2310039H08Rik | 69.40799082 | -0.797521459 | 0.129130729 | -6.176078044 | 6.57E-10 | 4.11E-08 |
| Sh3rf3 | 39.28963348 | -0.797195559 | 0.179936177 | -4.430435115 | 9.40E-06 | 0.000127022 |
| Ppfibp1 | 36.29760376 | -0.795526236 | 0.16287297 | -4.884335546 | 1.04E-06 | 2.05E-05 |
| Slc44a5 | 41.60872865 | -0.794881418 | 0.129398473 | -6.142896422 | 8.10E-10 | 4.92E-08 |
| E230016M11Rik | 9.579759597 | -0.79326005 | 0.283736118 | -2.795766914 | 0.005177673 | 0.022074127 |
| Zfp512b | 113.1404979 | -0.791702867 | 0.147000145 | -5.385728469 | 7.22E-08 | 2.22E-06 |
| Amer1 | 15.9578421 | -0.791091401 | 0.21273864 | -3.718607021 | 0.000200324 | 0.001570629 |
| Tdg | 110.6862315 | -0.790566997 | 0.139322876 | -5.674351674 | 1.39E-08 | 5.44E-07 |
| Zdhhc12 | 36.70732227 | -0.789701468 | 0.192582533 | -4.100587186 | 4.12E-05 | 0.000429616 |
| Stx2 | 44.8106624 | -0.789544015 | 0.135728352 | -5.81708984 | 5.99E-09 | 2.62E-07 |

| | | | | | | |
|---|---|---|---|---|---|---|
| Lmnb1 | 13.6916822 | -0.786896091 | 0.235249463 | -3.344943191 | 0.000822994 | 0.004991261 |
| Pf4 | 17.46480003 | -0.786072001 | 0.277772184 | -2.829916193 | 0.00465602 | 0.020288996 |
| Ecm1 | 24.46981439 | -0.783948531 | 0.16891684 | -4.641032427 | 3.47E-06 | 5.63E-05 |
| Tek | 24.61679949 | -0.782784068 | 0.17483177 | -4.477355972 | 7.56E-06 | 0.000106134 |
| Nckap5l | 38.07396974 | -0.782748922 | 0.190949417 | -4.099247515 | 4.14E-05 | 0.00043158 |
| AI480526 | 51.1911992 | -0.780855694 | 0.179480014 | -4.350655407 | 1.36E-05 | 0.000170428 |
| Sema5b | 31.86756253 | -0.780640378 | 0.156282467 | -4.995060505 | 5.88E-07 | 1.28E-05 |
| Snhg20 | 314.9666056 | -0.778913391 | 0.159064476 | -4.896840658 | 9.74E-07 | 1.95E-05 |
| Thpo | 20.93235637 | -0.777343667 | 0.225199036 | -3.451807256 | 0.000556845 | 0.00362067 |
| Fzd6 | 17.87740918 | -0.776889344 | 0.253140816 | -3.069000707 | 0.002147761 | 0.010845009 |
| Lpar6 | 10.12274098 | -0.776351493 | 0.23890992 | -3.249557384 | 0.001155848 | 0.006608018 |
| Myo19 | 15.51907279 | -0.775298565 | 0.238415396 | -3.251881284 | 0.001146439 | 0.00657196 |
| Pglyrp1 | 73.7600905 | -0.773732092 | 0.19089072 | -4.053272436 | 5.05E-05 | 0.000509904 |
| Trpc3 | 34.48645498 | -0.773153386 | 0.197515473 | -3.914394018 | 9.06E-05 | 0.000824193 |
| Sertad4 | 42.2673509 | -0.772978512 | 0.171319667 | -4.511907634 | 6.42E-06 | 9.30E-05 |
| Adgra2 | 18.21088924 | -0.771781198 | 0.210641756 | -3.663951603 | 0.000248354 | 0.001866066 |
| Hspa12b | 10.45431681 | -0.770249 | 0.248703031 | -3.097063177 | 0.001954482 | 0.010072485 |
| Itga9 | 10.49743927 | -0.768821087 | 0.275585975 | -2.789768553 | 0.005274573 | 0.022391508 |
| Anks6 | 16.97860818 | -0.768366752 | 0.207471931 | -3.703473273 | 0.000212668 | 0.001643834 |
| 3010001F23Rik | 12.21410226 | -0.767895707 | 0.234580443 | -3.273485621 | 0.001062298 | 0.006179483 |
| Dgkd | 231.5813677 | -0.767093247 | 0.09680964 | -7.923727925 | 2.30E-15 | 6.62E-13 |
| Sox4 | 66.00097381 | -0.765421101 | 0.146770242 | -5.215097355 | 1.84E-07 | 4.95E-06 |
| Slc24a3 | 288.3754365 | -0.764916427 | 0.121256022 | -6.308275767 | 2.82E-10 | 2.01E-08 |
| Lratd2 | 12.75698095 | -0.764198117 | 0.27651236 | -2.763703283 | 0.005714948 | 0.023772782 |
| Cacng5 | 23.78182443 | -0.762603657 | 0.214553825 | -3.554369908 | 0.000378886 | 0.002648006 |
| Pigc | 40.26495497 | -0.76131018 | 0.151139535 | -5.037134598 | 4.73E-07 | 1.07E-05 |
| Rmst | 21.40967273 | -0.760041507 | 0.252878898 | -3.005555278 | 0.002650964 | 0.012909075 |
| Ky | 8.490017048 | -0.759862689 | 0.295647653 | -2.570163106 | 0.010165064 | 0.037986773 |
| Pear1 | 12.7750998 | -0.759626557 | 0.252166427 | -3.012401633 | 0.002591894 | 0.012687143 |
| Cdh5 | 23.75301945 | -0.759377252 | 0.20312817 | -3.738414274 | 0.000185185 | 0.001472678 |
| Stard9 | 14.52947161 | -0.759321867 | 0.224015134 | -3.389600761 | 0.000699945 | 0.004386235 |
| Cyyr1 | 12.44485587 | -0.759294103 | 0.242959882 | -3.125183042 | 0.001776945 | 0.009345637 |
| Ryr3 | 54.99297318 | -0.758162233 | 0.166735169 | -4.54710446 | 5.44E-06 | 8.07E-05 |
| Cartpt | 90.34454989 | -0.75751857 | 0.229921121 | -3.294688923 | 0.000985307 | 0.005805382 |
| Rhcg | 10.20535971 | -0.757242114 | 0.304916546 | -2.483440548 | 0.013012005 | 0.046105038 |
| Pdzd2 | 42.63114585 | -0.756729295 | 0.177594968 | -4.260983869 | 2.04E-05 | 0.000239121 |
| Dnase1l2 | 16.9267475 | -0.755941258 | 0.291563458 | -2.59271605 | 0.009522135 | 0.036045045 |
| Ecscr | 19.51779065 | -0.755766933 | 0.221704833 | -3.408887944 | 0.000652283 | 0.004144244 |
| Gm43597 | 42.18880876 | -0.75275868 | 0.146737845 | -5.129955953 | 2.90E-07 | 7.27E-06 |
| Ntf3 | 36.24344074 | -0.75267513 | 0.210966403 | -3.567748796 | 0.000360061 | 0.002540966 |
| Dnhd1 | 10.70995603 | -0.751794952 | 0.257261295 | -2.922301047 | 0.003474555 | 0.015991633 |
| Ankrd13b | 231.2870331 | -0.751077492 | 0.128290485 | -5.854506604 | 4.78E-09 | 2.16E-07 |
| Actn2 | 23.87060209 | -0.750964687 | 0.177742315 | -4.225019161 | 2.39E-05 | 0.000272723 |
| Ebi3 | 16.81470845 | -0.750697345 | 0.201478367 | -3.725945153 | 0.000194585 | 0.001532716 |
| Srpk3 | 11.59229529 | -0.749776013 | 0.25413255 | -2.950334433 | 0.003174301 | 0.01492582 |
| Pcsk5 | 53.6717507 | -0.748352002 | 0.247350007 | -3.025477994 | 0.002482405 | 0.012260851 |
| Slc22a8 | 46.50081308 | -0.747014101 | 0.180582753 | -4.136685754 | 3.52E-05 | 0.000377549 |
| Uck2 | 89.77960229 | -0.746678104 | 0.178052741 | -4.193578262 | 2.75E-05 | 0.000305216 |
| Sema6c | 61.59362589 | -0.745342596 | 0.187932831 | -3.966005262 | 7.31E-05 | 0.000692124 |
| Ccdc57 | 26.17823221 | -0.745193234 | 0.191153469 | -3.898402882 | 9.68E-05 | 0.000867522 |
| Usp43 | 19.32571675 | -0.744119286 | 0.228509583 | -3.25640298 | 0.001128335 | 0.006488488 |
| Cd59a | 46.00912772 | -0.742210467 | 0.149321281 | -4.970560531 | 6.68E-07 | 1.43E-05 |
| Cyren | 43.99368565 | -0.740850956 | 0.12115198 | -6.115054469 | 9.65E-10 | 5.64E-08 |
| Cemip2 | 18.75608445 | -0.740409125 | 0.210440827 | -3.518372059 | 0.000434203 | 0.002958271 |
| Nectin3 | 25.70626651 | -0.740041887 | 0.183409177 | -4.034922899 | 5.46E-05 | 0.000544308 |
| 9330160F10Rik | 9.312977815 | -0.739836689 | 0.261782367 | -2.826151734 | 0.004711094 | 0.020497371 |
| Ret | 10.47048184 | -0.739118844 | 0.29974694 | -2.465809474 | 0.013670403 | 0.047966673 |
| Nectin2 | 39.04872811 | -0.738958922 | 0.186420018 | -3.963946212 | 7.37E-05 | 0.000697345 |
| Ccdc39 | 29.8998726 | -0.73531532 | 0.176927408 | -4.156028335 | 3.24E-05 | 0.000352914 |
| Sulf2 | 221.4133512 | -0.734292567 | 0.163750666 | -4.484211191 | 7.32E-06 | 0.000103465 |
| Rab3d | 29.97602734 | -0.732689102 | 0.170704998 | -4.292136191 | 1.77E-05 | 0.000213501 |
| Carmil3 | 177.6441774 | -0.732155986 | 0.151144703 | -4.844073079 | 1.27E-06 | 2.45E-05 |
| Rgl2 | 110.4171606 | -0.731947748 | 0.139227054 | -5.257223541 | 1.46E-07 | 4.05E-06 |
| Piezo1 | 15.41279122 | -0.731527901 | 0.266488402 | -2.745064683 | 0.006049897 | 0.024927518 |
| Zfp760 | 17.571873 | -0.73144887 | 0.185290828 | -3.94757191 | 7.89E-05 | 0.000733285 |
| Prss36 | 46.23318603 | -0.731187357 | 0.127711042 | -5.725326073 | 1.03E-08 | 4.19E-07 |
| Wdr90 | 29.27364364 | -0.729979633 | 0.18821198 | -3.878497169 | 0.000105104 | 0.000930833 |
| Gm48957 | 11.16448865 | -0.729189273 | 0.244240254 | -2.985540928 | 0.002830773 | 0.013631939 |
| Rep15 | 35.8417772 | -0.728568224 | 0.171528904 | -4.247495366 | 2.16E-05 | 0.000251162 |
| Slc26a8 | 9.267902446 | -0.728390028 | 0.294169403 | -2.476090374 | 0.013282995 | 0.046830293 |
| Arhgdig | 964.4418072 | -0.724345111 | 0.105635015 | -6.857055047 | 7.03E-12 | 8.32E-10 |
| Sbk1 | 212.7208369 | -0.723383767 | 0.145499805 | -4.971716396 | 6.64E-07 | 1.42E-05 |
| 8430429K09Rik | 58.37242236 | -0.723190207 | 0.138849959 | -5.208429399 | 1.90E-07 | 5.10E-06 |

| | | | | | | |
|---|---|---|---|---|---|---|
| Igflr1 | 25.49376075 | -0.721769095 | 0.226483401 | -3.186852073 | 0.001438303 | 0.007862198 |
| Rps6kb2 | 79.16199157 | -0.721558234 | 0.175399067 | -4.113808841 | 3.89E-05 | 0.000409247 |
| Adamts17 | 12.35600048 | -0.720639861 | 0.239424946 | -3.009877926 | 0.002613527 | 0.01277448 |
| Chaf1a | 9.772813945 | -0.720451522 | 0.245887973 | -2.929999029 | 0.003389631 | 0.015685948 |
| Mmp24 | 134.6058243 | -0.720123524 | 0.110419984 | -6.521677481 | 6.95E-11 | 5.92E-09 |
| Pdgfrb | 39.00877376 | -0.719288036 | 0.162400305 | -4.429105208 | 9.46E-06 | 0.000127503 |
| A230051N06Rik | 24.14822512 | -0.718586896 | 0.198885604 | -3.613066415 | 0.000302597 | 0.002200482 |
| Fam131c | 20.98150182 | -0.717767565 | 0.213345199 | -3.364348339 | 0.000767246 | 0.00471388 |
| Nrm | 8.299823364 | -0.717325226 | 0.2912337 | -2.463057081 | 0.013775798 | 0.048256573 |
| Rhobtb1 | 9.900649858 | -0.716334498 | 0.259107715 | -2.764620494 | 0.005698906 | 0.023717689 |
| Zfp580 | 210.1449628 | -0.716197979 | 0.147002391 | -4.872015844 | 1.10E-06 | 2.17E-05 |
| Pak6 | 166.384471 | -0.716027174 | 0.147879669 | -4.841958193 | 1.29E-06 | 2.47E-05 |
| Srgap1 | 38.25831974 | -0.715406561 | 0.182836618 | -3.91281883 | 9.12E-05 | 0.000826928 |
| Mmp16 | 27.76045428 | -0.715058034 | 0.170393362 | -4.196513443 | 2.71E-05 | 0.000301883 |
| Chfr | 81.25807675 | -0.714670409 | 0.1294863 | -5.519274302 | 3.40E-08 | 1.19E-06 |
| Trpc4 | 43.5471434 | -0.714312623 | 0.166501506 | -4.290127096 | 1.79E-05 | 0.000215288 |
| Grtp1 | 12.86983407 | -0.713940796 | 0.256452023 | -2.783915631 | 0.0053707 | 0.022716171 |
| Tmem198b | 23.50844829 | -0.713928489 | 0.191377675 | -3.730469028 | 0.000191124 | 0.001510367 |
| Opn3 | 18.29030576 | -0.713263021 | 0.28514603 | -2.501395589 | 0.012370491 | 0.044266877 |
| Adamts15 | 16.44196487 | -0.712007041 | 0.250050552 | -2.847452388 | 0.004407069 | 0.019454224 |
| Cdkl2 | 131.7045087 | -0.711808193 | 0.162000696 | -4.393858855 | 1.11E-05 | 0.000145577 |
| Gpr101 | 34.78359024 | -0.711405734 | 0.28427623 | -2.502515715 | 0.012331415 | 0.04415501 |
| 2810049E08Rik | 16.00862631 | -0.711396419 | 0.231151021 | -3.077626108 | 0.002086565 | 0.010611872 |
| Cfap65 | 8.485866911 | -0.709512414 | 0.260278557 | -2.725973369 | 0.006411217 | 0.026112585 |
| Fbn1 | 11.40057104 | -0.709341348 | 0.261086865 | -2.716878727 | 0.006590075 | 0.026679809 |
| Afap1l2 | 23.95474381 | -0.708346642 | 0.192167043 | -3.686098461 | 0.000227718 | 0.00174033 |
| Gm42067 | 99.06936138 | -0.708222222 | 0.131276473 | -5.394890695 | 6.86E-08 | 2.13E-06 |
| Miip | 54.06928382 | -0.708039583 | 0.192403833 | -3.679966104 | 0.000233265 | 0.001773923 |
| Gpam | 156.2257556 | -0.707676231 | 0.166610477 | -4.247489382 | 2.16E-05 | 0.000251162 |
| Epha8 | 30.46696253 | -0.706856394 | 0.220858618 | -3.200492697 | 0.001371928 | 0.007580039 |
| Aldh1b1 | 39.27060599 | -0.706723995 | 0.211688231 | -3.338513402 | 0.00084228 | 0.005090012 |
| Sfxn2 | 12.72460553 | -0.706664516 | 0.238806076 | -2.959156352 | 0.003084825 | 0.014557594 |
| Ndst3 | 29.09313622 | -0.705890868 | 0.173494873 | -4.068655495 | 4.73E-05 | 0.000483433 |
| Ccdc141 | 27.05214813 | -0.704475011 | 0.225656977 | -3.121884462 | 0.001796974 | 0.009427581 |
| Atp11b | 105.3693286 | -0.704214124 | 0.152376456 | -4.621541556 | 3.81E-06 | 6.08E-05 |
| Gm3294 | 11.01120162 | -0.703742728 | 0.267057112 | -2.635176888 | 0.008409345 | 0.032604483 |
| D430019H16Rik | 259.0788795 | -0.703032662 | 0.139892821 | -5.025509214 | 5.02E-07 | 1.13E-05 |
| Agrn | 261.59344 | -0.703021569 | 0.118757264 | -5.919819492 | 3.22E-09 | 1.57E-07 |
| Cdca8 | 9.044048604 | -0.698830297 | 0.2508867 | -2.785441785 | 0.005345484 | 0.022640872 |
| Homer3 | 63.15018622 | -0.698767096 | 0.226049556 | -3.09121198 | 0.001993413 | 0.010220913 |
| Bbc3 | 52.83186382 | -0.69805202 | 0.136081087 | -5.129676977 | 2.90E-07 | 7.27E-06 |
| Il16 | 26.16505106 | -0.697021127 | 0.195580602 | -3.563856122 | 0.000365446 | 0.002568276 |
| Ache | 242.1541418 | -0.696608899 | 0.114435539 | -6.087347572 | 1.15E-09 | 6.51E-08 |
| Srl | 10.1335546 | -0.69621745 | 0.27582113 | -2.524162854 | 0.011597419 | 0.042115025 |
| Tubb2a | 4889.919136 | -0.696009824 | 0.149476419 | -4.656318548 | 3.22E-06 | 5.29E-05 |
| Megf11 | 49.74665084 | -0.694809247 | 0.157345204 | -4.415827312 | 1.01E-05 | 0.000133906 |
| Nit2 | 36.49477871 | -0.694604061 | 0.137460569 | -5.053114995 | 4.35E-07 | 1.00E-05 |
| Parp6 | 458.5916796 | -0.694065201 | 0.09634528 | -7.203935666 | 5.85E-13 | 9.35E-11 |
| Id3 | 364.315994 | -0.691897715 | 0.120911664 | -5.722340542 | 1.05E-08 | 4.25E-07 |
| B4galt2 | 342.0018164 | -0.69161391 | 0.090693766 | -7.625815326 | 2.42E-14 | 5.63E-12 |
| Gm16099 | 8.586470762 | -0.691489723 | 0.277907721 | -2.488199034 | 0.012839186 | 0.045574758 |
| Phtf2 | 62.83056402 | -0.69019157 | 0.252225005 | -2.73641216 | 0.006211318 | 0.025493473 |
| Ap1s3 | 8.119626623 | -0.68925913 | 0.274781879 | -2.50838641 | 0.012128396 | 0.043603061 |
| Syce2 | 26.01240301 | -0.688675276 | 0.161600662 | -4.261586965 | 2.03E-05 | 0.000238773 |
| Atp11c | 21.51613118 | -0.687774417 | 0.173301003 | -3.968669581 | 7.23E-05 | 0.000685965 |
| Casp6 | 16.47254886 | -0.686289937 | 0.194721744 | -3.524464829 | 0.000424339 | 0.002899588 |
| Cntnap5a | 128.9195336 | -0.685616169 | 0.173370137 | -3.954638219 | 7.67E-05 | 0.000718244 |
| Casp3 | 27.79295397 | -0.685565245 | 0.270678897 | -2.532762072 | 0.011316774 | 0.041369557 |
| Epha3 | 40.9911174 | -0.685056599 | 0.14476201 | -4.732295433 | 2.22E-06 | 3.88E-05 |
| Ndc1 | 63.32129887 | -0.683500567 | 0.111716861 | -6.118150484 | 9.47E-10 | 5.55E-08 |
| Cdk5r2 | 635.789404 | -0.682836328 | 0.14178572 | -4.815973913 | 1.46E-06 | 2.73E-05 |
| Ocel1 | 59.82618119 | -0.681466965 | 0.13330304 | -5.112163722 | 3.18E-07 | 7.81E-06 |
| Bak1 | 44.46693696 | -0.67910101 | 0.180386005 | -3.764710074 | 0.000166742 | 0.001354962 |
| Heg1 | 38.86936496 | -0.678461833 | 0.157637802 | -4.303928531 | 1.68E-05 | 0.000204479 |
| Smim40 | 10.95330516 | -0.678412833 | 0.248046624 | -2.735021432 | 0.006237621 | 0.025582851 |
| Slc2a6 | 130.8891782 | -0.678186595 | 0.130654215 | -5.190698184 | 2.10E-07 | 5.57E-06 |
| Palmd | 36.75906604 | -0.677462366 | 0.184280877 | -3.67624887 | 0.000236689 | 0.001794276 |
| Rad52 | 41.68552819 | -0.677118456 | 0.143511041 | -4.718232471 | 2.38E-06 | 4.13E-05 |
| Rgs9 | 29.04913033 | -0.676687723 | 0.220437265 | -3.069751949 | 0.002142366 | 0.010823011 |
| Trim46 | 266.6377697 | -0.67599278 | 0.080643435 | -8.382489901 | 5.18E-17 | 2.20E-14 |
| Scmh1 | 136.06662 | -0.675404641 | 0.132590047 | -5.093931667 | 3.51E-07 | 8.53E-06 |
| Kctd5 | 85.86847604 | -0.674963971 | 0.135596008 | -4.977756946 | 6.43E-07 | 1.39E-05 |
| Ranbp3l | 12.19861954 | -0.67484021 | 0.247713823 | -2.724273524 | 0.006444311 | 0.026214225 |

| | | | | | | |
|---|---|---|---|---|---|---|
| Mirg | 110.8126199 | -0.674256333 | 0.127042302 | -5.307337179 | 1.11E-07 | 3.21E-06 |
| Sidt2 | 236.9245943 | -0.67372379 | 0.10414418 | -6.469144913 | 9.86E-11 | 7.84E-09 |
| Adap1 | 849.3113498 | -0.673459465 | 0.099178892 | -6.790350743 | 1.12E-11 | 1.24E-09 |
| B930059L03Rik | 8.207920114 | -0.673018207 | 0.271555363 | -2.478383045 | 0.013197937 | 0.046598257 |
| Vat1l | 61.64930977 | -0.673002096 | 0.132864581 | -5.065323572 | 4.08E-07 | 9.53E-06 |
| Rgs5 | 154.5444842 | -0.672034761 | 0.168115988 | -3.997447067 | 6.40E-05 | 0.000621627 |
| Hr | 26.83766412 | -0.671662063 | 0.2370198 | -2.833780396 | 0.004600093 | 0.020112504 |
| Hsd11b1 | 144.9847518 | -0.671548853 | 0.182586984 | -3.67796673 | 0.000235101 | 0.00178548 |
| Cask | 160.7709463 | -0.671506822 | 0.098922462 | -6.788213817 | 1.14E-11 | 1.25E-09 |
| Myorg | 79.6100729 | -0.671103905 | 0.140323978 | -4.782531943 | 1.73E-06 | 3.15E-05 |
| Slc7a6 | 56.86235432 | -0.670970869 | 0.132356458 | -5.069422966 | 3.99E-07 | 9.39E-06 |
| 9430021M05Rik | 12.66394656 | -0.670853582 | 0.250661881 | -2.67632868 | 0.007443361 | 0.029417676 |
| Pkia | 433.1145732 | -0.670276711 | 0.07952123 | -8.428902742 | 3.49E-17 | 1.60E-14 |
| Gm21781 | 16.21994713 | -0.669698135 | 0.217554361 | -3.078302514 | 0.002081835 | 0.010597351 |
| Rtel1 | 28.2032196 | -0.668670441 | 0.202954197 | -3.294686444 | 0.000985316 | 0.005805382 |
| Cbln2 | 30.97581074 | -0.668350493 | 0.185265163 | -3.607534635 | 0.00030912 | 0.002239273 |
| Stmn2 | 3217.322333 | -0.668088678 | 0.110201025 | -6.062454305 | 1.34E-09 | 7.50E-08 |
| Gmip | 15.99636788 | -0.667726566 | 0.19708949 | -3.387935934 | 0.000704207 | 0.004405419 |
| Mdm1 | 16.08988851 | -0.667626693 | 0.207659291 | -3.215009978 | 0.0013044 | 0.007270925 |
| Cnksr3 | 20.53569954 | -0.66637845 | 0.224029018 | -2.974518465 | 0.002934488 | 0.014031978 |
| Ptprb | 47.60021621 | -0.665714687 | 0.189741279 | -3.508539044 | 0.000450575 | 0.003046549 |
| Tgfb2 | 23.43098636 | -0.665517973 | 0.202626468 | -3.284457263 | 0.00102179 | 0.005991132 |
| Cd276 | 27.09023151 | -0.665410665 | 0.220570918 | -3.016765178 | 0.002554876 | 0.01255296 |
| Glis2 | 29.32708671 | -0.661440436 | 0.156630575 | -4.22293308 | 2.41E-05 | 0.000274782 |
| Cckbr | 181.0498139 | -0.661351312 | 0.162995874 | -4.05747271 | 4.96E-05 | 0.000502319 |
| Fam193b | 161.6288139 | -0.66134748 | 0.150277793 | -4.400833075 | 1.08E-05 | 0.000141725 |
| Ypel1 | 146.4665303 | -0.658296615 | 0.181664647 | -3.62369137 | 0.000290428 | 0.002124147 |
| Lrrc45 | 119.9808166 | -0.657404697 | 0.15065794 | -4.363558264 | 1.28E-05 | 0.000162481 |
| Ccnl2 | 222.1737238 | -0.656319938 | 0.129437998 | -5.070535288 | 3.97E-07 | 9.38E-06 |
| Focad | 97.17516642 | -0.655158647 | 0.106038427 | -6.17850215 | 6.47E-10 | 4.06E-08 |
| Frmd4a | 86.2833396 | -0.654900966 | 0.174999686 | -3.742297942 | 0.000182345 | 0.001455241 |
| Fchsd1 | 15.29117755 | -0.654177878 | 0.223443874 | -2.927705584 | 0.003414732 | 0.015763379 |
| Igsf21 | 318.8371541 | -0.653288931 | 0.167546271 | -3.899155308 | 9.65E-05 | 0.000866615 |
| Dlg2 | 497.8796111 | -0.653186352 | 0.117313652 | -5.567863078 | 2.58E-08 | 9.34E-07 |
| Samd14 | 166.1934979 | -0.652869542 | 0.120652621 | -5.411150917 | 6.26E-08 | 1.99E-06 |
| 9330020H09Rik | 33.36223722 | -0.652613366 | 0.141456821 | -4.613516415 | 3.96E-06 | 6.27E-05 |
| Purg | 112.0931123 | -0.651261335 | 0.149880513 | -4.345203551 | 1.39E-05 | 0.000173943 |
| B230307C23Rik | 23.92517247 | -0.651164828 | 0.175278236 | -3.715035254 | 0.000203175 | 0.00159004 |
| Snhg11 | 2171.034065 | -0.650535518 | 0.233773928 | -2.782754793 | 0.005389952 | 0.022767278 |
| Siah3 | 17.74369326 | -0.650405119 | 0.251599148 | -2.585084746 | 0.009735507 | 0.036705198 |
| Cdc7 | 30.15745594 | -0.650373569 | 0.152499813 | -4.264749934 | 2.00E-05 | 0.000235743 |
| Zkscan2 | 38.07245522 | -0.649347879 | 0.167301181 | -3.881310786 | 0.000103895 | 0.000923505 |
| Adam17 | 35.52648711 | -0.648725691 | 0.157231627 | -4.125923675 | 3.69E-05 | 0.000391796 |
| Sirt4 | 41.24576818 | -0.648513962 | 0.131849228 | -4.918602664 | 8.72E-07 | 1.78E-05 |
| 5430416N02Rik | 75.83340005 | -0.6482195 | 0.116852591 | -5.54732673 | 2.90E-08 | 1.04E-06 |
| Myo7a | 37.02547463 | -0.648087405 | 0.146693682 | -4.41796399 | 9.96E-06 | 0.000132985 |
| Amy1 | 37.38362037 | -0.647381943 | 0.232036909 | -2.78999555 | 0.005270877 | 0.022387028 |
| Mycl | 55.61809866 | -0.646496104 | 0.177076352 | -3.650945464 | 0.000261277 | 0.001945915 |
| Sh2b2 | 30.3349551 | -0.645071713 | 0.146369869 | -4.407134598 | 1.05E-05 | 0.0001385 |
| Grip1 | 39.35692575 | -0.644801019 | 0.159392362 | -4.045369617 | 5.22E-05 | 0.000523363 |
| Cep89 | 43.00123409 | -0.644658677 | 0.180719698 | -3.567174383 | 0.000360851 | 0.002545481 |
| Nup210 | 67.35939221 | -0.644005212 | 0.122313803 | -5.265188359 | 1.40E-07 | 3.91E-06 |
| Sh3bp5l | 131.2063954 | -0.643934108 | 0.085110735 | -7.565838844 | 3.85E-14 | 8.60E-12 |
| Eogt | 38.67558472 | -0.643489184 | 0.17574289 | -3.661537516 | 0.000250706 | 0.001882906 |
| Zcchc7 | 76.51546276 | -0.643385163 | 0.158226999 | -4.066216066 | 4.78E-05 | 0.00048793 |
| Tbata | 42.91467097 | -0.643038525 | 0.172828156 | -3.720681522 | 0.000198686 | 0.001560669 |
| Grik1 | 48.8104111 | -0.642817839 | 0.156148613 | -4.116705412 | 3.84E-05 | 0.000404641 |
| Fkbp10 | 25.08459157 | -0.641938779 | 0.213363951 | -3.008656218 | 0.002624059 | 0.012818564 |
| Gm42595 | 10.01090574 | -0.641476698 | 0.234445659 | -2.736142362 | 0.006216413 | 0.025508209 |
| Apc | 434.2416905 | -0.641398703 | 0.169869516 | -3.7758317 | 0.000159475 | 0.001307183 |
| Adam11 | 343.4674225 | -0.640973515 | 0.107362198 | -5.970197404 | 2.37E-09 | 1.21E-07 |
| Rhbdl3 | 29.85237837 | -0.639304002 | 0.166230525 | -3.845888123 | 0.000120117 | 0.001038295 |
| Gm12371 | 14.27742367 | -0.638816493 | 0.227803203 | -2.804247192 | 0.005043422 | 0.021610478 |
| Sipa1 | 43.98666541 | -0.638309622 | 0.155282639 | -4.110630958 | 3.95E-05 | 0.000413893 |
| Ust | 97.85020977 | -0.638110128 | 0.116243322 | -5.489434709 | 4.03E-08 | 1.38E-06 |
| Zkscan17 | 30.30938119 | -0.638037483 | 0.167638842 | -3.80602417 | 0.000141219 | 0.00118329 |
| Tmem121b | 170.6346058 | -0.638006984 | 0.169418939 | -3.765853964 | 0.000165981 | 0.00134942 |
| Gm19531 | 62.63363669 | -0.637454515 | 0.17608116 | -3.620231226 | 0.00029434 | 0.002149657 |
| Trmu | 53.37911839 | -0.636822125 | 0.126970639 | -5.015506968 | 5.29E-07 | 1.17E-05 |
| Csdc2 | 293.0054497 | -0.636329207 | 0.158937196 | -4.003651898 | 6.24E-05 | 0.000610039 |
| Unc13b | 118.3393522 | -0.636163455 | 0.110630666 | -5.750335591 | 8.91E-09 | 3.69E-07 |
| Inhbb | 27.43008128 | -0.636142345 | 0.215813507 | -2.947648427 | 0.00320201 | 0.015039421 |
| F2r | 12.41262359 | -0.63608868 | 0.232611082 | -2.734558801 | 0.006246394 | 0.025612632 |

| | | | | | | |
|---|---|---|---|---|---|---|
| Sfi1 | 38.1205773 | -0.635636396 | 0.197893699 | -3.212009268 | 0.001318101 | 0.007325618 |
| Map4k4 | 159.8254736 | -0.635339153 | 0.145204937 | -4.375465218 | 1.21E-05 | 0.000154903 |
| Obsl1 | 38.03695656 | -0.63522714 | 0.139124497 | -4.565889922 | 4.97E-06 | 7.53E-05 |
| C78859 | 38.37425667 | -0.635189567 | 0.202775998 | -3.132469194 | 0.001733426 | 0.009179415 |
| Plppr2 | 555.2712805 | -0.633796858 | 0.112968626 | -5.610379449 | 2.02E-08 | 7.52E-07 |
| Nfya | 51.17293305 | -0.6326559 | 0.163139394 | -3.878008149 | 0.000105315 | 0.000931732 |
| Hip1 | 73.78995345 | -0.63236645 | 0.152599526 | -4.143960761 | 3.41E-05 | 0.000368186 |
| Eif4e2 | 152.8783757 | -0.632106628 | 0.105203804 | -6.008400878 | 1.87E-09 | 9.92E-08 |
| Ogfr | 138.5402638 | -0.632005388 | 0.127435012 | -4.959432863 | 7.07E-07 | 1.50E-05 |
| Efcab6 | 19.18530091 | -0.63169376 | 0.221526613 | -2.851547953 | 0.004350692 | 0.019253226 |
| Nmb | 18.35165735 | -0.631589962 | 0.232467872 | -2.716891403 | 0.006589823 | 0.026679809 |
| Gdpd5 | 129.5988309 | -0.63041259 | 0.113770553 | -5.541087501 | 3.01E-08 | 1.07E-06 |
| Arxes1 | 174.6936517 | -0.629267346 | 0.130188617 | -4.833505093 | 1.34E-06 | 2.55E-05 |
| Plpp5 | 95.79264515 | -0.62919854 | 0.143201551 | -4.393796972 | 1.11E-05 | 0.000145577 |
| Fam189b | 167.7328058 | -0.628544617 | 0.130386433 | -4.820628991 | 1.43E-06 | 2.68E-05 |
| Gjd2 | 19.72299815 | -0.627881018 | 0.235702166 | -2.663874621 | 0.007724635 | 0.030345373 |
| Brinp3 | 128.9222799 | -0.627826059 | 0.222005756 | -2.827971984 | 0.004684391 | 0.020396893 |
| Cacna1g | 149.3760242 | -0.627479149 | 0.123607831 | -5.076370524 | 3.85E-07 | 9.20E-06 |
| Lrrc56 | 15.68018994 | -0.626131157 | 0.226736952 | -2.761487054 | 0.00575388 | 0.023893683 |
| Raver2 | 67.07644615 | -0.626123063 | 0.128581928 | -4.869448378 | 1.12E-06 | 2.19E-05 |
| Slco1a4 | 77.46074092 | -0.626029406 | 0.135732743 | -4.612220994 | 3.98E-06 | 6.30E-05 |
| Lrfn4 | 163.9767989 | -0.625400933 | 0.123164473 | -5.077770537 | 3.82E-07 | 9.16E-06 |
| R74862 | 52.33392914 | -0.625384821 | 0.152095567 | -4.111788602 | 3.93E-05 | 0.000412333 |
| Gdf11 | 40.86736306 | -0.625013212 | 0.133201457 | -4.692240062 | 2.70E-06 | 4.59E-05 |
| Abcd1 | 46.15217091 | -0.624126033 | 0.122156121 | -5.109248968 | 3.23E-07 | 7.92E-06 |
| Kank3 | 46.82714069 | -0.62412544 | 0.135674293 | -4.600174634 | 4.22E-06 | 6.58E-05 |
| Abca3 | 336.7677528 | -0.623707854 | 0.124275013 | -5.018771174 | 5.20E-07 | 1.16E-05 |
| Hps4 | 55.34427622 | -0.621977303 | 0.12720959 | -4.889390042 | 1.01E-06 | 2.01E-05 |
| Slc45a4 | 144.847162 | -0.621886522 | 0.109658228 | -5.671134152 | 1.42E-08 | 5.50E-07 |
| Fnbp1l | 209.2110649 | -0.62152578 | 0.122844312 | -5.059459149 | 4.20E-07 | 9.78E-06 |
| Sestd1 | 110.6053966 | -0.621466015 | 0.13538943 | -4.590210724 | 4.43E-06 | 6.79E-05 |
| Tspan4 | 124.6139774 | -0.621179508 | 0.125307175 | -4.957254116 | 7.15E-07 | 1.51E-05 |
| Irgq | 248.6234257 | -0.621055042 | 0.072181811 | -8.604037912 | 7.70E-18 | 4.50E-15 |
| Dchs1 | 33.61425739 | -0.61955046 | 0.222959259 | -2.778760857 | 0.005456668 | 0.022968955 |
| Bcl6 | 84.99686225 | -0.619165789 | 0.108044201 | -5.730671187 | 1.00E-08 | 4.08E-07 |
| Mapk7 | 37.17309519 | -0.618707486 | 0.161654168 | -3.827352518 | 0.000129529 | 0.001106676 |
| C230057M02Rik | 42.74770545 | -0.618308164 | 0.149214472 | -4.14375466 | 3.42E-05 | 0.000368186 |
| Kctd15 | 39.32120512 | -0.618227839 | 0.160674283 | -3.847708723 | 0.000119228 | 0.001033774 |
| Igfbp7 | 187.2800332 | -0.616308671 | 0.163432485 | -3.771029175 | 0.000162576 | 0.001327314 |
| Flywch2 | 196.5761962 | -0.614955805 | 0.132488378 | -4.641583004 | 3.46E-06 | 5.62E-05 |
| Mast1 | 405.2192322 | -0.614258126 | 0.089298426 | -6.878711679 | 6.04E-12 | 7.42E-10 |
| Mllt11 | 1865.616462 | -0.612208374 | 0.092130244 | -6.645031512 | 3.03E-11 | 2.89E-09 |
| Snhg5 | 38.65335932 | -0.612045118 | 0.141995827 | -4.310303571 | 1.63E-05 | 0.000199677 |
| Guf1 | 65.92937949 | -0.611805386 | 0.163551772 | -3.74074447 | 0.000183476 | 0.001462889 |
| Prep | 90.83848103 | -0.611797109 | 0.172287609 | -3.551022113 | 0.000383738 | 0.00267436 |
| Syngap1 | 172.438917 | -0.610445964 | 0.169355125 | -3.604531983 | 0.000312716 | 0.002259223 |
| Epb41l2 | 148.2656893 | -0.609842303 | 0.117493719 | -5.190424713 | 2.10E-07 | 5.57E-06 |
| Gm28050 | 52.52143774 | -0.6095395 | 0.152271555 | -4.002976786 | 6.26E-05 | 0.000610349 |
| Sh3d19 | 37.66916592 | -0.609416834 | 0.152037789 | -4.008324749 | 6.12E-05 | 0.000600562 |
| Myo5b | 71.82379996 | -0.608777235 | 0.220231739 | -2.764257487 | 0.00570525 | 0.023738265 |
| Map4k1 | 15.13026777 | -0.608624601 | 0.238707589 | -2.54966591 | 0.010782619 | 0.039767724 |
| Slc31a1 | 71.75708162 | -0.608469144 | 0.154692402 | -3.933413257 | 8.37E-05 | 0.000770273 |
| Hcn4 | 15.13284473 | -0.607740502 | 0.238205452 | -2.551329099 | 0.010731294 | 0.039644224 |
| Glra2 | 44.94745834 | -0.607179469 | 0.230487905 | -2.63432248 | 0.008430538 | 0.032664238 |
| Lncppara | 31.3981214 | -0.606001564 | 0.221244156 | -2.739062463 | 0.006161467 | 0.025328163 |
| Ints6l | 56.10407094 | -0.605609705 | 0.188581751 | -3.211390822 | 0.001320942 | 0.007333749 |
| Impdh1 | 226.3411631 | -0.60545152 | 0.120316731 | -5.032147368 | 4.85E-07 | 1.09E-05 |
| Fam163b | 184.9575743 | -0.605228706 | 0.129305608 | -4.68060678 | 2.86E-06 | 4.80E-05 |
| Foxq1 | 11.79963741 | -0.605022554 | 0.233561014 | -2.590426132 | 0.009585719 | 0.036232863 |
| Gm5914 | 90.48356268 | -0.60380287 | 0.172960329 | -3.490990534 | 0.000481233 | 0.003218156 |
| Fscn1 | 416.6525087 | -0.603107287 | 0.125043125 | -4.823194285 | 1.41E-06 | 2.66E-05 |
| Bcan | 767.0437611 | -0.603026426 | 0.147207709 | -4.096432379 | 4.20E-05 | 0.000436591 |
| Slc16a13 | 14.70906733 | -0.602835524 | 0.211804716 | -2.846185564 | 0.00442464 | 0.019511466 |
| Sirt2 | 579.4041784 | -0.60185198 | 0.118811603 | -5.065599371 | 4.07E-07 | 9.53E-06 |
| Mir124a-1hg | 268.1134073 | -0.598624186 | 0.166167975 | -3.602524417 | 0.000315142 | 0.002272211 |
| Car4 | 104.6865482 | -0.598567309 | 0.135321578 | -4.423295378 | 9.72E-06 | 0.000130154 |
| Id1 | 130.2526162 | -0.59850261 | 0.107679038 | -5.558209104 | 2.73E-08 | 9.81E-07 |
| 5031434O11Rik | 23.34214572 | -0.598451989 | 0.189607934 | -3.156260261 | 0.001598062 | 0.008558846 |
| Asxl3 | 21.309071 | -0.597432215 | 0.185924328 | -3.213308454 | 0.001312153 | 0.007299739 |
| Lnx2 | 12.37996361 | -0.596385429 | 0.220012872 | -2.710684261 | 0.006714453 | 0.027060555 |
| Grik5 | 856.947278 | -0.595748659 | 0.064296849 | -9.265596515 | 1.94E-20 | 1.57E-17 |
| Nphp3 | 15.06083451 | -0.595392995 | 0.198569843 | -2.998405938 | 0.002713959 | 0.013170433 |
| Shisa7 | 259.6115843 | -0.595244666 | 0.103269923 | -5.76396932 | 8.22E-09 | 3.42E-07 |

| | | | | | | |
|---|---|---|---|---|---|---|
| Pmm2 | 46.35142158 | -0.595088585 | 0.128304872 | -4.638082541 | 3.52E-06 | 5.70E-05 |
| Pantr1 | 504.1927808 | -0.595061425 | 0.127245926 | -4.676467406 | 2.92E-06 | 4.88E-05 |
| Atp10a | 31.94160911 | -0.594828583 | 0.153356129 | -3.878740204 | 0.000104999 | 0.000930389 |
| 1810058I24Rik | 239.2466106 | -0.593830436 | 0.092895187 | -6.392478003 | 1.63E-10 | 1.24E-08 |
| Ptprf | 218.7980415 | -0.593068826 | 0.13952692 | -4.250569169 | 2.13E-05 | 0.000248585 |
| Agt | 182.6798657 | -0.591680014 | 0.179051481 | -3.304524553 | 0.000951376 | 0.005642681 |
| Pomc | 31.28408668 | -0.591659322 | 0.150028002 | -3.943659273 | 8.02E-05 | 0.000742915 |
| Myo16 | 57.44763189 | -0.591595804 | 0.216064909 | -2.738046671 | 0.00618053 | 0.025385552 |
| Lsr | 24.40564673 | -0.591463803 | 0.226769272 | -2.608218464 | 0.009101485 | 0.03475541 |
| Gpr173 | 41.95981023 | -0.591103418 | 0.124199353 | -4.759311567 | 1.94E-06 | 3.48E-05 |
| C230037L18Rik | 12.23681962 | -0.590596717 | 0.235793551 | -2.504719552 | 0.012254852 | 0.043964444 |
| Pheta1 | 58.3501493 | -0.590466383 | 0.126148064 | -4.680740751 | 2.86E-06 | 4.80E-05 |
| Mchr1 | 68.70621476 | -0.590115153 | 0.149308941 | -3.952309559 | 7.74E-05 | 0.000722608 |
| Narf | 161.0781666 | -0.58978109 | 0.114954792 | -5.130548091 | 2.89E-07 | 7.26E-06 |
| Slc1a6 | 66.19364526 | -0.589121492 | 0.15109396 | -3.899040663 | 9.66E-05 | 0.000866615 |
| Csmd2 | 54.7198971 | -0.589040689 | 0.149077167 | -3.951246853 | 7.77E-05 | 0.000724894 |
| Nfasc | 364.056305 | -0.588889826 | 0.106816992 | -5.513072562 | 3.53E-08 | 1.22E-06 |
| 2610008E11Rik | 50.73046798 | -0.58872306 | 0.118605836 | -4.96369387 | 6.92E-07 | 1.47E-05 |
| Usp40 | 25.16194156 | -0.588655934 | 0.165865851 | -3.54898812 | 0.000386715 | 0.002686557 |
| Plod1 | 47.91972306 | -0.587295883 | 0.14154529 | -4.149172919 | 3.34E-05 | 0.000361418 |
| 5830448L01Rik | 18.75123212 | -0.587111936 | 0.193211731 | -3.038697144 | 0.002376036 | 0.011811198 |
| Mgat4b | 248.3621921 | -0.586554241 | 0.100874522 | -5.814691648 | 6.07E-09 | 2.65E-07 |
| Kdm4b | 147.8139103 | -0.585881267 | 0.090635352 | -6.464158319 | 1.02E-10 | 8.03E-09 |
| Tekt5 | 39.72485744 | -0.585526793 | 0.239006223 | -2.449839107 | 0.014292006 | 0.049735946 |
| Cacna1i | 155.4841598 | -0.58506271 | 0.12273992 | -4.766686412 | 1.87E-06 | 3.37E-05 |
| Mir670hg | 40.91579768 | -0.584398603 | 0.175704424 | -3.326032373 | 0.000880917 | 0.005287685 |
| Btbd17 | 101.234321 | -0.584036803 | 0.115521538 | -5.055652925 | 4.29E-07 | 9.91E-06 |
| Slc38a6 | 18.59970308 | -0.583866018 | 0.22777229 | -2.563375985 | 0.010365973 | 0.038609888 |
| Usp6nl | 43.31992556 | -0.583440848 | 0.134021961 | -4.353322721 | 1.34E-05 | 0.000169244 |
| Fam102b | 92.8560136 | -0.581749298 | 0.195783397 | -2.971392405 | 0.002964527 | 0.014139477 |
| Serpinf1 | 31.63630662 | -0.581687033 | 0.194797649 | -2.986109108 | 0.002825518 | 0.013614371 |
| Ccdc134 | 22.73436834 | -0.581403363 | 0.200001266 | -2.906998416 | 0.00364915 | 0.016655021 |
| Nectin1 | 104.4828283 | -0.581311503 | 0.129618155 | -4.484800018 | 7.30E-06 | 0.000103266 |
| Slc29a3 | 35.44754659 | -0.58076837 | 0.157598072 | -3.685123572 | 0.000228592 | 0.001745627 |
| Rnf165 | 137.4114591 | -0.580616872 | 0.137681227 | -4.217109938 | 2.47E-05 | 0.000280574 |
| Adgrf5 | 39.12607556 | -0.580560752 | 0.14519912 | -3.998376528 | 6.38E-05 | 0.000620257 |
| Plxna1 | 111.3638158 | -0.580510862 | 0.136962302 | -4.238471859 | 2.25E-05 | 0.000260216 |
| Dcbld2 | 72.09459463 | -0.580510019 | 0.102028729 | -5.689672156 | 1.27E-08 | 5.06E-07 |
| Klhdc10 | 213.0003891 | -0.580463032 | 0.102537375 | -5.660989759 | 1.51E-08 | 5.81E-07 |
| Ptprr | 76.82014725 | -0.580055535 | 0.132207358 | -4.38746787 | 1.15E-05 | 0.00014873 |
| Sephs1 | 80.67418745 | -0.580046128 | 0.138828353 | -4.178153194 | 2.94E-05 | 0.000323694 |
| Cyp11a1 | 21.34891062 | -0.579332367 | 0.194068026 | -2.98520256 | 0.002833906 | 0.013643152 |
| Agbl3 | 19.13938499 | -0.578794621 | 0.20802227 | -2.782368549 | 0.005396372 | 0.02277914 |
| Hs6st1 | 249.0881111 | -0.57873714 | 0.12422944 | -4.658615047 | 3.18E-06 | 5.24E-05 |
| Trp53inp2 | 432.27968 | -0.578375427 | 0.119911663 | -4.823345881 | 1.41E-06 | 2.66E-05 |
| Nsmf | 1385.765987 | -0.577527425 | 0.090004013 | -6.416685274 | 1.39E-10 | 1.08E-08 |
| Gm15663 | 13.33764747 | -0.576512353 | 0.211451867 | -2.726447215 | 0.006402019 | 0.026086689 |
| Grin3a | 172.9948455 | -0.575790914 | 0.140872444 | -4.0873211 | 4.36E-05 | 0.000450495 |
| Gabrg3 | 49.66486452 | -0.575679959 | 0.188225327 | -3.058461731 | 0.002224765 | 0.011190502 |
| Gm39469 | 54.73150494 | -0.575547877 | 0.212357439 | -2.710278855 | 0.006722666 | 0.027080778 |
| 1110017D15Rik | 21.54786704 | -0.575392591 | 0.177056822 | -3.249762329 | 0.001155015 | 0.006606039 |
| Slc35f2 | 13.60606733 | -0.574179213 | 0.225903171 | -2.54170497 | 0.011031324 | 0.040535873 |
| Rfc5 | 40.96772995 | -0.573963733 | 0.137330963 | -4.1794197 | 2.92E-05 | 0.000322316 |
| Ephb2 | 55.82872428 | -0.57346874 | 0.170796877 | -3.357606706 | 0.000786204 | 0.004816387 |
| Nhsl1 | 53.34811989 | -0.572940885 | 0.173698451 | -3.298480097 | 0.000972098 | 0.0057458 |
| Ttc13 | 95.58776523 | -0.571873596 | 0.115439301 | -4.953889981 | 7.27E-07 | 1.53E-05 |
| Gm2a | 231.1190741 | -0.571763787 | 0.164930361 | -3.466698216 | 0.000526893 | 0.003456411 |
| Pigo | 53.66977272 | -0.571393238 | 0.116654125 | -4.8981829 | 9.67E-07 | 1.94E-05 |
| Parp8 | 63.48853569 | -0.5712255 | 0.155723529 | -3.668202891 | 0.000244261 | 0.001841029 |
| Ick | 49.97052168 | -0.570525095 | 0.147015587 | -3.880711611 | 0.000104151 | 0.000924813 |
| Abcb6 | 79.05910141 | -0.570077603 | 0.107662415 | -5.295047496 | 1.19E-07 | 3.39E-06 |
| Cdk16 | 658.4124783 | -0.569543694 | 0.107963657 | -5.275327898 | 1.33E-07 | 3.73E-06 |
| Abhd5 | 58.33350168 | -0.568987736 | 0.161925652 | -3.513882635 | 0.000441608 | 0.0029948 |
| Sfswap | 158.9216836 | -0.568636516 | 0.100095532 | -5.680938038 | 1.34E-08 | 5.29E-07 |
| Sh3bp4 | 76.29231674 | -0.568602387 | 0.151890909 | -3.743491897 | 0.00018148 | 0.001449706 |
| Pcdh15 | 31.69614386 | -0.568270966 | 0.19487133 | -2.916134282 | 0.003543979 | 0.016264191 |
| Sdc3 | 534.1951601 | -0.568117638 | 0.147282243 | -3.857339666 | 0.000114628 | 0.000999515 |
| Slc30a10 | 44.88523582 | -0.56774377 | 0.174401848 | -3.255377027 | 0.001132419 | 0.006505957 |
| Atp13a2 | 562.9152214 | -0.567625058 | 0.093893937 | -6.04538563 | 1.49E-09 | 8.20E-08 |
| Gpsm2 | 22.1776462 | -0.566172059 | 0.176549914 | -3.20686681 | 0.001341891 | 0.007438324 |
| A730063M14Rik | 21.15995075 | -0.566103829 | 0.174315481 | -3.247582071 | 0.001163901 | 0.006638386 |
| Eng | 51.12246478 | -0.565676951 | 0.167049584 | -3.386281707 | 0.000708466 | 0.004421653 |
| Fbxo41 | 250.0278313 | -0.56547997 | 0.088732122 | -6.372889038 | 1.86E-10 | 1.39E-08 |

| | | | | | | |
|---|---|---|---|---|---|---|
| Ncam1 | 707.490578 | -0.565117493 | 0.124111558 | -4.553302717 | 5.28E-06 | 7.89E-05 |
| Trit1 | 47.39238816 | -0.564772835 | 0.152119734 | -3.712686175 | 0.000205071 | 0.001602238 |
| Tmem223 | 488.159775 | -0.564651614 | 0.0659868 | -8.557038923 | 1.16E-17 | 6.33E-15 |
| Firre | 47.0891786 | -0.564576141 | 0.224357328 | -2.516414977 | 0.01185555 | 0.042812831 |
| Lamc1 | 48.91686871 | -0.564442725 | 0.211960271 | -2.662964723 | 0.007745553 | 0.030406409 |
| Vcam1 | 66.48577391 | -0.564247777 | 0.141997501 | -3.973645834 | 7.08E-05 | 0.000674048 |
| Sarm1 | 71.72277509 | -0.563990602 | 0.123617087 | -4.562400037 | 5.06E-06 | 7.62E-05 |
| Tspoap1 | 284.0856671 | -0.563663574 | 0.14556553 | -3.872232483 | 0.000107843 | 0.000950622 |
| Cgref1 | 90.42114035 | -0.563639487 | 0.09725602 | -5.795420059 | 6.82E-09 | 2.92E-07 |
| Shroom2 | 272.8575588 | -0.563210641 | 0.122469043 | -4.598800063 | 4.25E-06 | 6.61E-05 |
| Lims2 | 51.88342348 | -0.56272378 | 0.144346435 | -3.898425217 | 9.68E-05 | 0.000867522 |
| Zfp579 | 70.62663813 | -0.562712805 | 0.105976372 | -5.309794953 | 1.10E-07 | 3.17E-06 |
| Shmt2 | 67.18222472 | -0.562645485 | 0.115085643 | -4.888928527 | 1.01E-06 | 2.01E-05 |
| Slc6a7 | 281.5527295 | -0.562509911 | 0.119445517 | -4.70934302 | 2.49E-06 | 4.29E-05 |
| Fam189a2 | 47.97909897 | -0.562239499 | 0.182222773 | -3.085451342 | 0.002032435 | 0.010395991 |
| Dvl2 | 45.12227816 | -0.561582795 | 0.188929495 | -2.972446385 | 0.002954368 | 0.014110874 |
| Tmem131l | 25.34501769 | -0.560964086 | 0.170802524 | -3.284284524 | 0.001022416 | 0.005992732 |
| Tbc1d4 | 21.53378867 | -0.56094769 | 0.194178924 | -2.888818616 | 0.003866921 | 0.017474852 |
| Slc8a2 | 349.4939923 | -0.560547319 | 0.149081505 | -3.760005773 | 0.000169909 | 0.001377396 |
| Tac1 | 145.0368258 | -0.560371938 | 0.136601545 | -4.102237181 | 4.09E-05 | 0.000427615 |
| D630045J12Rik | 88.18233122 | -0.560247018 | 0.114337685 | -4.899933186 | 9.59E-07 | 1.93E-05 |
| Pomt2 | 60.43411213 | -0.560157563 | 0.132809179 | -4.217762429 | 2.47E-05 | 0.000279951 |
| Rimkla | 43.85573187 | -0.560154187 | 0.14277397 | -3.923363533 | 8.73E-05 | 0.000797942 |
| Higd1b | 26.44967892 | -0.559976642 | 0.173727296 | -3.223308342 | 0.001267191 | 0.007117346 |
| Ankrd16 | 63.01038996 | -0.559185066 | 0.16432044 | -3.403015864 | 0.000666464 | 0.004215385 |
| Nat8f1 | 126.108822 | -0.55907679 | 0.099496083 | -5.619083418 | 1.92E-08 | 7.22E-07 |
| Ankrd10 | 103.9702183 | -0.558269692 | 0.129632411 | -4.306559521 | 1.66E-05 | 0.000202646 |
| Gm45051 | 21.20027398 | -0.558256439 | 0.186846624 | -2.987779099 | 0.002810126 | 0.013559477 |
| Aco1 | 70.70774653 | -0.558085347 | 0.129177315 | -4.320304587 | 1.56E-05 | 0.000192227 |
| Ing4 | 146.0540004 | -0.55804061 | 0.0918137 | -6.077966697 | 1.22E-09 | 6.83E-08 |
| P2ry1 | 11.20701326 | -0.557784476 | 0.225162573 | -2.477252184 | 0.013239832 | 0.046713171 |
| Tbcd | 123.0415091 | -0.556851044 | 0.103557886 | -5.37719595 | 7.57E-08 | 2.31E-06 |
| 09.Mar | 150.9644432 | -0.556595551 | 0.127700462 | -4.35860249 | 1.31E-05 | 0.000165583 |
| Carmil1 | 45.09554742 | -0.556471494 | 0.160067769 | -3.476474363 | 0.000508053 | 0.003353584 |
| Tax1bp3 | 48.97380271 | -0.556244398 | 0.127100268 | -4.376421912 | 1.21E-05 | 0.000154458 |
| Oxtr | 27.35650158 | -0.555462583 | 0.157058451 | -3.536661548 | 0.000405219 | 0.002795629 |
| Fbxl19 | 101.831572 | -0.55518542 | 0.105359505 | -5.269438377 | 1.37E-07 | 3.83E-06 |
| Iqce | 57.49818009 | -0.55493263 | 0.131263747 | -4.227615325 | 2.36E-05 | 0.000270325 |
| Cidea | 22.64010949 | -0.554822354 | 0.221538486 | -2.504406182 | 0.012265713 | 0.043975487 |
| Tfrc | 152.9173001 | -0.554674542 | 0.136935423 | -4.050628614 | 5.11E-05 | 0.000514781 |
| Wsb1 | 131.6926338 | -0.554208043 | 0.149769205 | -3.700413866 | 0.000215248 | 0.001656726 |
| Pik3r3 | 116.9817156 | -0.554195312 | 0.085050134 | -6.516101569 | 7.22E-11 | 6.12E-09 |
| Zfc3h1 | 73.19331185 | -0.554150513 | 0.116397358 | -4.760851287 | 1.93E-06 | 3.46E-05 |
| Snhg16 | 19.31997824 | -0.554069104 | 0.188585229 | -2.938030232 | 0.003303048 | 0.015356547 |
| Wbp1 | 163.4836481 | -0.553814524 | 0.082065304 | -6.748461271 | 1.49E-11 | 1.56E-09 |
| Wnt7b | 218.9634332 | -0.553649883 | 0.123723776 | -4.474886724 | 7.65E-06 | 0.000107191 |
| Thbs3 | 38.88785026 | -0.553550379 | 0.151035785 | -3.665027987 | 0.000247312 | 0.001859895 |
| Zfp618 | 13.66106033 | -0.553422072 | 0.225841315 | -2.450490837 | 0.014266159 | 0.049686802 |
| Pcyt1b | 79.77467634 | -0.552663834 | 0.149329337 | -3.700972927 | 0.000214774 | 0.001654837 |
| Atoh7 | 12.63251321 | -0.552229749 | 0.22042861 | -2.505254419 | 0.012236334 | 0.043907303 |
| Rac3 | 330.4466929 | -0.551638475 | 0.138788779 | -3.974661909 | 7.05E-05 | 0.000671932 |
| Kmt2b | 105.5257474 | -0.551576292 | 0.132973092 | -4.148029377 | 3.35E-05 | 0.000362995 |
| 2700081O15Rik | 112.1327605 | -0.551489207 | 0.10404319 | -5.300579573 | 1.15E-07 | 3.30E-06 |
| Zfyve27 | 164.6317468 | -0.550786249 | 0.11805245 | -4.665606255 | 3.08E-06 | 5.10E-05 |
| Echdc2 | 29.12136031 | -0.550653202 | 0.142443615 | -3.865762616 | 0.000110743 | 0.000972641 |
| Ahr | 29.41819811 | -0.550431873 | 0.186863376 | -2.945638064 | 0.003222893 | 0.015100611 |
| Slc17a7 | 3995.046065 | -0.550034104 | 0.083025367 | -6.624892207 | 3.48E-11 | 3.24E-09 |
| Tysnd1 | 80.74718417 | -0.54970085 | 0.093043417 | -5.908003653 | 3.46E-09 | 1.66E-07 |
| Phka1 | 56.24675987 | -0.549614652 | 0.125084441 | -4.393948996 | 1.11E-05 | 0.000145577 |
| D630023F18Rik | 96.51623385 | -0.548340556 | 0.132841824 | -4.127770464 | 3.66E-05 | 0.00038953 |
| Eml4 | 79.16024438 | -0.54771384 | 0.113390707 | -4.830323872 | 1.36E-06 | 2.58E-05 |
| Slc4a7 | 47.29357388 | -0.547264233 | 0.154689537 | -3.537823203 | 0.00040344 | 0.002784492 |
| Nsd2 | 192.5720143 | -0.547060633 | 0.111905049 | -4.888614395 | 1.02E-06 | 2.01E-05 |
| Mbd6 | 80.97931234 | -0.546874337 | 0.161856183 | -3.378767036 | 0.000728117 | 0.004514377 |
| Wdr83 | 111.3290289 | -0.546499485 | 0.116422925 | -4.694088263 | 2.68E-06 | 4.57E-05 |
| Kif22 | 114.173608 | -0.546336984 | 0.165422376 | -3.302678858 | 0.00095766 | 0.00567201 |
| Mindy1 | 77.20674537 | -0.544800134 | 0.1179744 | -4.617952144 | 3.88E-06 | 6.16E-05 |
| Ttll12 | 70.57700758 | -0.544272453 | 0.121037092 | -4.496741003 | 6.90E-06 | 9.82E-05 |
| Tyk2 | 30.8756712 | -0.544095375 | 0.201868251 | -2.695299395 | 0.007032541 | 0.028141785 |
| Tab1 | 78.75365108 | -0.544043851 | 0.118309726 | -4.59847105 | 4.26E-06 | 6.61E-05 |
| Lysmd1 | 92.60192682 | -0.543807854 | 0.122396642 | -4.442996527 | 8.87E-06 | 0.000121317 |
| Jarid2 | 80.78469382 | -0.543001107 | 0.122513474 | -4.432174591 | 9.33E-06 | 0.000126303 |
| Mterf1a | 24.6907531 | -0.542992882 | 0.155542141 | -3.490969572 | 0.000481271 | 0.003218156 |

| | | | | | | |
|---|---|---|---|---|---|---|
| Nrros | 20.04724171 | -0.542926746 | 0.189451152 | -2.865787513 | 0.004159734 | 0.018531312 |
| Pgap1 | 126.5381585 | -0.542925367 | 0.140089893 | -3.875549884 | 0.000106384 | 0.000940208 |
| Elfn1 | 59.75759046 | -0.542320967 | 0.125496047 | -4.321418723 | 1.55E-05 | 0.00019158 |
| Spon1 | 160.8317744 | -0.541891241 | 0.138525156 | -3.911861613 | 9.16E-05 | 0.000828883 |
| Bhlhe22 | 121.8688811 | -0.541399901 | 0.186110086 | -2.909030424 | 0.003625516 | 0.01658287 |
| Hexdc | 120.4991249 | -0.540767671 | 0.13029678 | -4.150276555 | 3.32E-05 | 0.000360579 |
| Lrrc3 | 40.57925819 | -0.5401737 | 0.19324815 | -2.795233487 | 0.005186224 | 0.022105026 |
| Clic1 | 51.95209641 | -0.540149209 | 0.122485849 | -4.409890734 | 1.03E-05 | 0.000137177 |
| 4931428F04Rik | 79.2738767 | -0.539902331 | 0.103533738 | -5.214747781 | 1.84E-07 | 4.95E-06 |
| Vps37c | 68.11684609 | -0.539681046 | 0.120377519 | -4.483237837 | 7.35E-06 | 0.000103679 |
| Sox6 | 36.8774757 | -0.539677135 | 0.161599834 | -3.339589672 | 0.000839023 | 0.005075811 |
| Klhl23 | 178.6456388 | -0.53907049 | 0.121667908 | -4.430671155 | 9.39E-06 | 0.000126984 |
| H2afy2 | 81.05891199 | -0.538732284 | 0.133297703 | -4.041572151 | 5.31E-05 | 0.000531285 |
| Sox17 | 18.6961612 | -0.538709889 | 0.182025782 | -2.959525204 | 0.003081135 | 0.014552332 |
| Zfp821 | 113.2353591 | -0.538378029 | 0.092629002 | -5.812197217 | 6.17E-09 | 2.69E-07 |
| 1810055G02Rik | 103.6579776 | -0.538081665 | 0.118173743 | -4.553309837 | 5.28E-06 | 7.89E-05 |
| Ttc9b | 2330.871453 | -0.537963159 | 0.080414823 | -6.68985067 | 2.23E-11 | 2.21E-09 |
| Gucd1 | 35.23390466 | -0.53788063 | 0.205958857 | -2.611592619 | 0.009012157 | 0.034484216 |
| Mief2 | 73.71131736 | -0.537735064 | 0.102938275 | -5.223859301 | 1.75E-07 | 4.75E-06 |
| Spaca6 | 50.18827029 | -0.537632363 | 0.212941935 | -2.524783872 | 0.011576946 | 0.042071677 |
| Chpf2 | 180.2795036 | -0.536764409 | 0.12966453 | -4.139639492 | 3.48E-05 | 0.000373902 |
| Fam89b | 151.0235609 | -0.536007308 | 0.09212854 | -5.818037587 | 5.95E-09 | 2.61E-07 |
| Crmp1 | 706.8610323 | -0.535883793 | 0.15867101 | -3.377326411 | 0.000731941 | 0.00453146 |
| Cmtm3 | 40.48056608 | -0.535428661 | 0.188478682 | -2.840791623 | 0.004500171 | 0.01978791 |
| Zfp174 | 15.64856286 | -0.535394871 | 0.216211816 | -2.47625168 | 0.013276995 | 0.046828618 |
| Atf6b | 205.0699221 | -0.535360331 | 0.0854805 | -6.262952707 | 3.78E-10 | 2.55E-08 |
| Celsr3 | 112.6683695 | -0.535222843 | 0.161279522 | -3.318603857 | 0.000904687 | 0.005413111 |
| Jpt1 | 487.4001622 | -0.535065326 | 0.067213784 | -7.960648715 | 1.71E-15 | 5.37E-13 |
| Trim47 | 14.62678721 | -0.534945614 | 0.21120885 | -2.532780303 | 0.011316185 | 0.041369557 |
| Cldn1 | 39.61507343 | -0.534942178 | 0.184907833 | -2.893020639 | 0.003815562 | 0.017279614 |
| Scn3b | 375.3411715 | -0.534837301 | 0.118800142 | -4.501992099 | 6.73E-06 | 9.64E-05 |
| Stk17b | 30.53462902 | -0.53483556 | 0.190322494 | -2.81015422 | 0.004951777 | 0.021320184 |
| Neto2 | 140.6380997 | -0.534626712 | 0.142596706 | -3.749222037 | 0.000177384 | 0.001425716 |
| Mzf1 | 30.19892103 | -0.534505836 | 0.158209082 | -3.378477571 | 0.000728884 | 0.004515851 |
| Cdhr1 | 19.27920106 | -0.533579543 | 0.190239925 | -2.804771617 | 0.005035224 | 0.021597188 |
| Pcdhb22 | 25.61632148 | -0.533572381 | 0.151696594 | -3.517365598 | 0.000435853 | 0.002965935 |
| Col6a1 | 30.95398669 | -0.532905325 | 0.187024195 | -2.849392434 | 0.004380281 | 0.019361187 |
| Mdm4 | 103.9363023 | -0.532689162 | 0.123406433 | -4.316542881 | 1.58E-05 | 0.000195389 |
| 2700049A03Rik | 21.6836724 | -0.532059059 | 0.187418635 | -2.83888023 | 0.004527215 | 0.019868832 |
| Apbb1ip | 17.92086879 | -0.532027171 | 0.197471577 | -2.694196192 | 0.007055862 | 0.028195171 |
| Slc19a1 | 67.46541159 | -0.531643926 | 0.150063379 | -3.542795917 | 0.000395909 | 0.002741443 |
| Rfx3 | 65.75602659 | -0.531574587 | 0.158708388 | -3.34937928 | 0.000809928 | 0.0049332 |
| Acvr2a | 165.1255205 | -0.530845673 | 0.136497549 | -3.889049117 | 0.000100638 | 0.000896904 |
| Trim9 | 843.7500877 | -0.52964593 | 0.089996832 | -5.885161917 | 3.98E-09 | 1.84E-07 |
| Wfs1 | 777.7889091 | -0.529556172 | 0.130464471 | -4.059006767 | 4.93E-05 | 0.000500826 |
| Col23a1 | 57.28119897 | -0.529537076 | 0.183682163 | -2.882898738 | 0.003940341 | 0.017764021 |
| Slc6a20a | 66.30282856 | -0.529522014 | 0.207379615 | -2.553394727 | 0.010667852 | 0.03952585 |
| Triobp | 20.23740385 | -0.529359492 | 0.171107951 | -3.093716504 | 0.001976663 | 0.010149534 |
| Tunar | 35.89909645 | -0.529315073 | 0.1423241 | -3.719082524 | 0.000199948 | 0.001569127 |
| St6gal2 | 44.34011819 | -0.52929546 | 0.17982475 | -2.943396062 | 0.003246328 | 0.015180278 |
| Marveld1 | 20.07370492 | -0.52921749 | 0.183641889 | -2.881790709 | 0.003954223 | 0.017821864 |
| Dact1 | 21.99435415 | -0.529154357 | 0.193712336 | -2.731650284 | 0.006301799 | 0.025758814 |
| Eml5 | 89.71925728 | -0.529076393 | 0.119185138 | -4.439113834 | 9.03E-06 | 0.000123029 |
| Numbl | 333.3327956 | -0.528620511 | 0.079468996 | -6.651908757 | 2.89E-11 | 2.77E-09 |
| Znrf1 | 394.1477015 | -0.52830119 | 0.087743092 | -6.021000345 | 1.73E-09 | 9.33E-08 |
| Abcb1a | 59.86345197 | -0.528161889 | 0.134495568 | -3.926983584 | 8.60E-05 | 0.000787305 |
| Fkbp5 | 68.04934876 | -0.528102378 | 0.13355519 | -3.954188371 | 7.68E-05 | 0.000719199 |
| Cdh9 | 48.2223366 | -0.52804194 | 0.181731098 | -2.905622344 | 0.003665235 | 0.01671943 |
| Snapc4 | 83.90587838 | -0.527955849 | 0.164404931 | -3.211313961 | 0.001321295 | 0.007333749 |
| Mov10 | 28.9910201 | -0.527340189 | 0.196500581 | -2.683657146 | 0.007282174 | 0.028888401 |
| Tmem265 | 23.1093605 | -0.527327311 | 0.215347368 | -2.448728841 | 0.014336133 | 0.049869031 |
| Mir9-3hg | 192.704393 | -0.526726408 | 0.17960984 | -2.932614423 | 0.00336121 | 0.015584212 |
| 9430038I01Rik | 17.98349196 | -0.526720771 | 0.187515315 | -2.808948016 | 0.004970367 | 0.021378506 |
| Zfp629 | 52.10608094 | -0.526428453 | 0.112151587 | -4.693901052 | 2.68E-06 | 4.57E-05 |
| Vstm2l | 351.1824445 | -0.523619302 | 0.107901608 | -4.852747913 | 1.22E-06 | 2.36E-05 |
| Cfap69 | 40.04419763 | -0.523331628 | 0.167081998 | -3.132184401 | 0.001735109 | 0.009183906 |
| Abcc10 | 25.34335119 | -0.52311196 | 0.156507205 | -3.342414552 | 0.000830529 | 0.005031559 |
| Lhfpl4 | 238.3292639 | -0.52262515 | 0.068746713 | -7.602183776 | 2.91E-14 | 6.58E-12 |
| C130026L21Rik | 34.8300393 | -0.522261338 | 0.206292927 | -2.531649264 | 0.011352749 | 0.041483174 |
| Cfp | 33.44940571 | -0.521868965 | 0.205958568 | -2.533854113 | 0.011281569 | 0.041278381 |
| Slc41a2 | 53.28390697 | -0.521311323 | 0.167659448 | -3.109346525 | 0.001875017 | 0.009761489 |
| Bach2 | 22.0777157 | -0.521234999 | 0.198316364 | -2.628300508 | 0.008581267 | 0.03313225 |
| Arhgef7 | 308.8827937 | -0.521225793 | 0.090622938 | -5.751587858 | 8.84E-09 | 3.67E-07 |

| | | | | | | |
|---|---|---|---|---|---|---|
| Pkdcc | 44.6458481 | -0.520861457 | 0.176855947 | -2.945117004 | 0.003228326 | 0.015116947 |
| Fmnl3 | 19.97910718 | -0.520671 | 0.175979713 | -2.958698996 | 0.003089407 | 0.014575157 |
| Sdk2 | 58.82512884 | -0.520020246 | 0.20257825 | -2.567009276 | 0.010257987 | 0.038241288 |
| Gng2 | 545.6840301 | -0.51999018 | 0.084594283 | -6.14687138 | 7.90E-10 | 4.82E-08 |
| Spred3 | 76.57305517 | -0.519466517 | 0.141558437 | -3.669625966 | 0.000242906 | 0.001835315 |
| Klhl36 | 15.31873586 | -0.519303983 | 0.205530239 | -2.52665489 | 0.01151546 | 0.041897094 |
| 4930570G19Rik | 27.95869205 | -0.518721347 | 0.166064226 | -3.123618847 | 0.001786417 | 0.00938381 |
| Sugp2 | 178.7749339 | -0.518255627 | 0.080337366 | -6.450991038 | 1.11E-10 | 8.72E-09 |
| Tctn3 | 59.17224981 | -0.517973398 | 0.132496308 | -3.909342133 | 9.25E-05 | 0.000836682 |
| Akap8l | 404.6011167 | -0.517350948 | 0.088101624 | -5.872206732 | 4.30E-09 | 1.98E-07 |
| Dlk2 | 143.2480378 | -0.517216909 | 0.131283642 | -3.93969045 | 8.16E-05 | 0.000752846 |
| B3gat1 | 506.1794442 | -0.516400873 | 0.106405 | -4.853163603 | 1.22E-06 | 2.36E-05 |
| Caskin2 | 53.96980792 | -0.516372214 | 0.145189553 | -3.556538346 | 0.000375774 | 0.00262995 |
| Tgfb1i1 | 56.06706307 | -0.516270053 | 0.143569945 | -3.595947978 | 0.000323212 | 0.002315627 |
| P3h3 | 49.38659207 | -0.516024325 | 0.159429157 | -3.236699821 | 0.001209205 | 0.006841535 |
| Slc7a10 | 121.9802143 | -0.515512165 | 0.141026872 | -3.655417983 | 0.000256763 | 0.001919045 |
| Gfm2 | 63.72896225 | -0.515071215 | 0.141651335 | -3.636190334 | 0.0002767 | 0.002040165 |
| Hgh1 | 37.69285958 | -0.514967136 | 0.125538127 | -4.102077575 | 4.09E-05 | 0.000427647 |
| Arhgap33 | 386.7509935 | -0.514674664 | 0.128996521 | -3.989833688 | 6.61E-05 | 0.000637177 |
| Spata33 | 20.24867953 | -0.513971901 | 0.181329084 | -2.834470286 | 0.004590173 | 0.020095353 |
| Lzic | 29.94639415 | -0.513779162 | 0.173155058 | -2.967162316 | 0.003005622 | 0.01429125 |
| P2ry12 | 111.9399901 | -0.513473445 | 0.131569446 | -3.90268 | 9.51E-05 | 0.000856858 |
| Dtx2 | 34.83336082 | -0.512664976 | 0.143548032 | -3.571382823 | 0.000355101 | 0.002512239 |
| Lhpp | 67.97340069 | -0.512614807 | 0.12695507 | -4.037765547 | 5.40E-05 | 0.000538389 |
| Cd2ap | 40.13861227 | -0.511468741 | 0.132029667 | -3.87389253 | 0.000107111 | 0.000945522 |
| Dock6 | 29.23181383 | -0.511432185 | 0.194594437 | -2.628195302 | 0.008583922 | 0.03313225 |
| Mlxip | 43.30647762 | -0.511426459 | 0.152333022 | -3.357292141 | 0.000787099 | 0.004820128 |
| Tmem63b | 419.0405091 | -0.51080823 | 0.098984328 | -5.160496033 | 2.46E-07 | 6.32E-06 |
| Dcaf15 | 69.16349745 | -0.509853584 | 0.129081241 | -3.94986583 | 7.82E-05 | 0.000728688 |
| Ednrb | 132.9144261 | -0.509808426 | 0.158080503 | -3.22499243 | 0.00125976 | 0.007082651 |
| Eml6 | 57.34018096 | -0.509699499 | 0.167085584 | -3.050529467 | 0.002284383 | 0.011446222 |
| Gpcpd1 | 264.6281807 | -0.508914462 | 0.130429962 | -3.901821758 | 9.55E-05 | 0.000859446 |
| Hars2 | 88.92038896 | -0.508736167 | 0.098287788 | -5.1759855 | 2.27E-07 | 5.92E-06 |
| Sprn | 592.5653175 | -0.508385589 | 0.121750592 | -4.175631352 | 2.97E-05 | 0.000326477 |
| Limd2 | 392.7606856 | -0.507891788 | 0.094979997 | -5.347355255 | 8.92E-08 | 2.65E-06 |
| Doc2b | 38.13513645 | -0.507515252 | 0.174182979 | -2.913690273 | 0.003571841 | 0.016386002 |
| Ajm1 | 229.1205198 | -0.507215959 | 0.074374703 | -6.819737587 | 9.12E-12 | 1.05E-09 |
| Myo9b | 77.30987007 | -0.507106091 | 0.164382937 | -3.084907113 | 0.002036157 | 0.010411735 |
| Psme4 | 68.00812354 | -0.50706215 | 0.146707189 | -3.456286991 | 0.000547672 | 0.003571984 |
| Ago4 | 21.90307309 | -0.506945302 | 0.169506368 | -2.990715387 | 0.002783248 | 0.013464278 |
| Efna2 | 44.2530418 | -0.506704705 | 0.1495036 | -3.389247516 | 0.000700847 | 0.004390266 |
| 1500026H17Rik | 49.01579567 | -0.506609931 | 0.192367965 | -2.633546235 | 0.008449833 | 0.03272485 |
| Lcorl | 48.47637188 | -0.506456156 | 0.155974765 | -3.247039065 | 0.001166124 | 0.006644404 |
| Oxct1 | 1203.201875 | -0.506015941 | 0.114332785 | -4.425816616 | 9.61E-06 | 0.000129051 |
| Zfp341 | 40.02245835 | -0.506004256 | 0.133161645 | -3.799924938 | 0.00014474 | 0.001205644 |
| Gng3 | 3415.347042 | -0.505519276 | 0.059499433 | -8.49620326 | 1.96E-17 | 9.83E-15 |
| Tmem117 | 25.35330588 | -0.505179664 | 0.194118683 | -2.602426799 | 0.009256657 | 0.03522374 |
| Dancr | 55.5248015 | -0.504871954 | 0.189142688 | -2.66926499 | 0.007601745 | 0.029966786 |
| Smpd2 | 59.34740333 | -0.504326863 | 0.116985148 | -4.311033235 | 1.62E-05 | 0.000199307 |
| Usp37 | 126.5537653 | -0.504037763 | 0.103340177 | -4.87746178 | 1.07E-06 | 2.11E-05 |
| Srd5a1 | 51.3218746 | -0.503769317 | 0.159622702 | -3.156000443 | 0.001599486 | 0.008563768 |
| Cdh13 | 270.1707131 | -0.503496943 | 0.141950159 | -3.546998083 | 0.000389647 | 0.002704715 |
| Slc9a5 | 44.55618414 | -0.503082669 | 0.137095928 | -3.669566812 | 0.000242962 | 0.001835315 |
| 3110039M20Rik | 84.84310442 | -0.502776924 | 0.113262555 | -4.43903923 | 9.04E-06 | 0.000123029 |
| Sh3pxd2b | 32.42692052 | -0.502563883 | 0.175215353 | -2.868263963 | 0.00412731 | 0.018415909 |
| Rgma | 126.7056671 | -0.502504946 | 0.180810065 | -2.779186794 | 0.005449518 | 0.02295506 |
| Setd4 | 22.19021883 | -0.50189546 | 0.190900667 | -2.629092232 | 0.008561314 | 0.033065124 |
| Ramp2 | 153.2560328 | -0.501894175 | 0.131316213 | -3.822027467 | 0.000132359 | 0.001123413 |
| Plxnb3 | 27.97218923 | -0.501781133 | 0.165857836 | -3.025368854 | 0.002483301 | 0.012260871 |
| Ireb2 | 126.0379534 | -0.501533565 | 0.120814769 | -4.151260377 | 3.31E-05 | 0.000359284 |
| Matk | 646.7698742 | -0.501003691 | 0.117012927 | -4.281609753 | 1.86E-05 | 0.000221961 |
| Lama2 | 23.03129154 | -0.500907885 | 0.172054112 | -2.911339226 | 0.003598831 | 0.016483053 |
| Pfkfb4 | 33.24096091 | -0.500515045 | 0.17000892 | -2.944051674 | 0.003239459 | 0.015156727 |

**Supplementary Table 2** 573 overlapping DEGs between 10x visium and bulk RNA-seq analysis.

| 573 overlapping DEGs between 10x visium and bulk RNA-seq analysis (bulk data) | | | | | | |
|---|---|---|---|---|---|---|
| gene | Young1 | Young2 | Young3 | Old1 | Old2 | Old3 |
| Aass | -1.633524021 | 0.105332221 | -0.4854104 | 1.300191316 | 0.092219019 | 0.621191866 |
| Sik1 | -1.629724512 | 0.212034501 | -0.536266762 | 0.905731372 | 1.090557658 | -0.042332256 |
| Efemp1 | -1.578744978 | -0.069112841 | -0.568141088 | 1.340122027 | 0.453689819 | 0.422187061 |
| Dmp1 | -1.558573589 | -0.61510223 | -0.237155688 | 1.067962631 | 0.454567885 | 0.888300991 |
| Serpinb9 | -1.519079534 | -0.38576656 | -0.567394971 | 1.048505645 | 0.945072682 | 0.478662737 |
| Dusp1 | -1.511383178 | -0.724682292 | -0.244542681 | 0.937925754 | 0.946149854 | 0.596532544 |
| Itgb4 | -1.50182013 | -0.505563653 | -0.497434936 | 1.151153102 | 0.668424139 | 0.685241478 |
| Arc | -1.473279814 | -0.561913438 | -0.517039477 | 1.044941965 | 0.849831722 | 0.657459042 |
| Fos | -1.470979853 | -0.376036382 | -0.664857649 | 0.89759217 | 1.075613192 | 0.538668522 |
| Rtp4 | -1.466365464 | -0.39386024 | -0.657496772 | 0.608870938 | 1.141192356 | 0.767659182 |
| Gpc3 | -1.441824993 | -0.334794143 | -0.763011537 | 1.062247113 | 0.798553887 | 0.678829674 |
| Csrnp1 | -1.434421371 | -0.732880683 | -0.09502656 | 1.276920532 | 0.867037181 | 0.118370901 |
| Nr4a1 | -1.426423294 | -0.628320064 | -0.334608156 | 1.019400689 | 1.175387213 | 0.194563613 |
| Sgk1 | -1.416380298 | -0.582950934 | -0.581363176 | 1.049633103 | 0.620189833 | 0.910871471 |
| Gatm | -1.403994976 | -0.725583551 | -0.263076268 | 1.388821373 | 0.486884732 | 0.51694869 |
| Rdm1 | -1.400367648 | -0.895000105 | -0.151584133 | 1.191828345 | 0.686064576 | 0.569058966 |
| Ifit1 | -1.38687125 | -0.167280891 | -0.543868084 | -0.226217481 | 1.179409607 | 1.144828099 |
| Dusp5 | -1.381803805 | -0.706432428 | -0.310225652 | 1.29942776 | 0.86801432 | 0.231019804 |
| Egr1 | -1.380134019 | -0.526813492 | -0.697484478 | 0.881429433 | 1.048485617 | 0.674516939 |
| Nfe2l2 | -1.378918167 | -0.300057224 | -0.647251683 | 1.45070396 | 0.212147654 | 0.663375461 |
| Parp14 | -1.375356923 | -0.488203875 | -0.715537333 | 0.904524916 | 1.099768593 | 0.574804622 |
| Sdf2l1 | -1.370746755 | -0.846898992 | -0.294416411 | 1.213241993 | 0.620158055 | 0.678662111 |
| Arpc3 | -1.36820697 | -0.890554051 | -0.315928988 | 0.732261874 | 0.949796205 | 0.89263193 |
| Pdk4 | -1.355654732 | -0.894420636 | 0.034211538 | 1.421497887 | 0.508710263 | 0.285655679 |
| Usf1 | -1.355068077 | -0.978453304 | -0.212094132 | 0.875510745 | 0.847128889 | 0.822975879 |
| Plin4 | -1.349843943 | -0.866650703 | -0.141879988 | 1.18266289 | 0.189818567 | 0.985893177 |
| Irgm1 | -1.340017787 | -0.47229772 | -0.79337989 | 0.95190384 | 0.62915685 | 1.024634707 |
| Atf4 | -1.339820617 | -0.919910773 | -0.327470947 | 0.897501795 | 0.940647535 | 0.749053006 |
| Tnfaip6 | -1.335898607 | -0.774827652 | -0.414904542 | 1.294060686 | 0.545444073 | 0.686126043 |
| Btg2 | -1.323988514 | -0.923656854 | -0.309709321 | 1.098831332 | 0.845272629 | 0.613250728 |
| Stx4a | -1.31877342 | -1.03811205 | -0.178037334 | 0.806812548 | 0.80350048 | 0.924609775 |
| H2-DMb1 | -1.312796608 | -0.512867287 | -0.748817419 | 1.227497725 | 0.53407941 | 0.81290418 |
| Efhd2 | -1.311557313 | -1.059999628 | -0.067932826 | 1.116357813 | 0.539072132 | 0.784059823 |
| Pdlim1 | -1.305722606 | -0.454911485 | -0.622300892 | 0.90845271 | 1.365154468 | 0.109327804 |
| Krt12 | -1.29456078 | -0.965972284 | -0.187405975 | 1.173734705 | 0.925578711 | 0.348625622 |
| Gadd45b | -1.285373218 | -0.448169259 | -0.776942703 | 1.29611534 | 0.858240706 | 0.356129134 |
| Sertad1 | -1.279910365 | -0.811451727 | -0.52830952 | 0.598972036 | 0.901070984 | 1.119628593 |
| Nupr1 | -1.278362174 | -0.397024603 | -0.603482318 | 1.620298551 | 0.358829595 | 0.299740949 |
| Gng11 | -1.275821239 | -0.845470496 | -0.488835778 | 0.896484148 | 1.127495629 | 0.586147736 |
| Clic4 | -1.274949284 | -0.527573651 | -0.590887424 | 1.501319371 | 0.664161746 | 0.227929243 |
| Sqor | -1.271668499 | -1.074899877 | 0.157130695 | 1.239881652 | 0.802363568 | 0.147192462 |
| Ndrg1 | -1.264733806 | -0.706015477 | -0.653817525 | 1.141708676 | 0.930612006 | 0.552246126 |
| Cyp27a1 | -1.264574826 | -0.580657477 | -0.792664403 | 0.663683447 | 1.138987894 | 0.835225365 |
| Ggact | -1.256219157 | -0.806838337 | -0.550166634 | 1.19572967 | 0.578694765 | 0.838799695 |
| Hapln2 | -1.250229611 | -0.648238985 | -0.558739742 | 1.479339274 | 0.626708618 | 0.351160446 |
| Gpld1 | -1.248709438 | -0.846240779 | -0.486895229 | 1.251026018 | 0.529527329 | 0.801292099 |
| Cryab | -1.237527212 | -0.9969903 | -0.376502236 | 1.064539177 | 0.730007963 | 0.816472608 |
| Tuft1 | -1.234154974 | -0.676788614 | -0.676788614 | 1.325951097 | 0.573515681 | 0.688265425 |
| Nol3 | -1.233125751 | -0.742456578 | -0.64926734 | 1.247358611 | 0.720104122 | 0.657386936 |
| Apod | -1.226207022 | -0.634687807 | -0.764361838 | 1.239760139 | 0.613586433 | 0.771910095 |
| Anxa5 | -1.224977869 | -0.871986138 | -0.506923531 | 1.245310949 | 0.744224409 | 0.614352179 |
| Pcdhb6 | -1.224338698 | -0.113608821 | -1.046780935 | 1.190181668 | 0.959443275 | 0.235103511 |
| Bst2 | -1.223859971 | -0.649469186 | -0.808636868 | 0.895905104 | 1.01305171 | 0.773009211 |
| Prelp | -1.222936617 | -0.529035733 | -0.59457453 | 1.604839319 | 0.47198256 | 0.269725 |
| H2-Eb1 | -1.21488096 | -0.603417931 | -0.68273454 | 0.942768777 | 1.322693395 | 0.235571259 |
| Egr4 | -1.214363507 | -0.886820839 | -0.33580829 | 1.317625637 | 0.922639631 | 0.196727368 |
| Acot13 | -1.21332432 | -0.936660128 | -0.485123017 | 1.06693511 | 0.630406706 | 0.937765649 |
| Itih3 | -1.211558715 | -0.907389349 | -0.512867428 | 1.148444555 | 0.626321949 | 0.85704899 |
| Stat1 | -1.209691225 | -0.958009193 | -0.486280863 | 0.92558343 | 0.990714132 | 0.73768372 |
| Abt1 | -1.200241316 | -0.911678581 | -0.53878899 | 0.847404267 | 0.684947902 | 1.118356719 |
| Adamtsl4 | -1.198361144 | -0.334524462 | -0.881791257 | 1.131172904 | 1.176179143 | 0.107324816 |
| Ass1 | -1.197154079 | -0.981480053 | -0.416960296 | 1.139150816 | 0.539297812 | 0.917145799 |
| Prr5l | -1.196578543 | -0.522226256 | -0.762317004 | 1.490533053 | 0.463431406 | 0.527157343 |
| Crip1 | -1.189092693 | -0.7952498 | -0.458490156 | 0.818636648 | 1.426203372 | 0.197992629 |
| Kcnk13 | -1.186312798 | -0.517146256 | -0.967552904 | 0.975235748 | 0.839232702 | 0.856543508 |
| Mvp | -1.182742155 | -0.772892609 | -0.653420207 | 1.309802459 | 0.74233282 | 0.556919693 |
| Gcat | -1.180162941 | -0.603132483 | -0.73084409 | 1.25892432 | 0.217339073 | 1.03787612 |
| Igtp | -1.176959728 | -0.828392789 | -0.688639483 | 0.902875833 | 1.027984509 | 0.763131659 |
| Them6 | -1.175088494 | -0.756011157 | -0.555429493 | 1.486270084 | 0.622306418 | 0.377952643 |
| Rps10 | -1.17435765 | -1.019099538 | -0.439129797 | 1.103453762 | 0.72747986 | 0.801653363 |

| | | | | | | |
|---|---|---|---|---|---|---|
| Pdlim2 | -1.174021626 | -0.944011991 | -0.522256504 | 1.151914643 | 0.851948121 | 0.636427357 |
| Ccn1 | -1.171424817 | -0.665283108 | -0.752151025 | 1.336789549 | 0.781896707 | 0.470172694 |
| Gsn | -1.170560577 | -0.798409941 | -0.615097809 | 1.337506144 | 0.778366336 | 0.468195847 |
| Sf3b6 | -1.170514663 | -0.920253649 | -0.580488463 | 0.724959131 | 1.088167921 | 0.858129723 |
| Ccdc92 | -1.168006695 | -0.927643853 | -0.44045814 | 1.383748336 | 0.540773781 | 0.61158657 |
| Gm973 | -1.163140684 | -0.572286662 | -0.181232818 | 1.811087549 | 0.028958884 | 0.076613732 |
| Bag3 | -1.161507475 | -0.612913726 | -0.742924417 | 1.451719466 | 0.68778324 | 0.377842913 |
| Cd82 | -1.158903148 | -0.894068782 | -0.40358505 | 1.310245276 | 0.97374646 | 0.172565244 |
| Aldh9a1 | -1.158718645 | -0.749567397 | -0.512387388 | 1.492787612 | 0.758966706 | 0.168919114 |
| Mcam | -1.158500815 | -1.011243799 | -0.246412787 | 1.300094423 | 0.919624342 | 0.196438636 |
| Junb | -1.157745095 | -0.722767843 | -0.784842124 | 1.035468902 | 1.060702602 | 0.569183558 |
| Mobp | -1.157581548 | -0.844478265 | -0.438480618 | 1.439747906 | 0.801169678 | 0.199622847 |
| Rpl10 | -1.156689146 | -1.041433704 | -0.443507466 | 0.992897062 | 0.6906095 | 0.958123754 |
| Adm | -1.156534414 | -1.006045592 | -0.360824228 | 0.967382305 | 1.210258374 | 0.345763555 |
| Slc5a5 | -1.155595024 | -0.972162008 | -0.439117347 | 1.191418203 | 0.403773914 | 0.971682262 |
| Vwa5a | -1.153494647 | -0.984294038 | -0.536637715 | 0.904370496 | 1.003772136 | 0.766283768 |
| Ttyh2 | -1.149823421 | -1.042522787 | -0.43087225 | 1.059975976 | 0.954810222 | 0.608432259 |
| Rpl14 | -1.149049179 | -1.065162417 | -0.430847972 | 0.913430179 | 0.752104151 | 0.979525237 |
| Anxa11 | -1.14791483 | -0.881961634 | -0.544757967 | 1.291557054 | 0.882796409 | 0.400280968 |
| Srebf1 | -1.146296949 | -0.973807182 | -0.283494569 | 1.472430413 | 0.29863326 | 0.632535028 |
| Plekhb1 | -1.144574215 | -0.975006825 | -0.512932646 | 1.210259142 | 0.717470572 | 0.704783972 |
| Psme2 | -1.143059703 | -0.939704716 | -0.565590445 | 1.191109986 | 0.658079217 | 0.799165662 |
| Aif1l | -1.142414411 | -1.261361607 | -0.030008211 | 0.964456847 | 0.518657642 | 0.950669741 |
| Chac1 | -1.141013989 | -0.74731038 | -0.809356835 | 1.093622301 | 0.770064083 | 0.833994818 |
| Bex1 | -1.139093371 | -1.097385566 | -0.404951438 | 0.885379406 | 0.943107543 | 0.812943426 |
| Rpl35a | -1.131705497 | -0.953574716 | -0.609028328 | 0.786975925 | 0.972803488 | 0.934529128 |
| Gpr37 | -1.13054065 | -0.830080727 | -0.611190769 | 1.387739294 | 0.719423749 | 0.464649103 |
| Gstm1 | -1.127725612 | -0.895255645 | -0.524041498 | 1.418501163 | 0.524218671 | 0.604302921 |
| Ifi27 | -1.127132117 | -0.748077788 | -0.824555343 | 1.028632247 | 0.969047306 | 0.702085694 |
| Dkkl1 | -1.121216341 | -1.079497807 | -0.269528293 | 0.713516747 | 1.354159112 | 0.402566582 |
| Plin3 | -1.118344249 | -1.02139428 | -0.434270934 | 1.311329331 | 0.650094423 | 0.612585707 |
| Tspo | -1.118171258 | -0.689797657 | -0.801576208 | 1.35325146 | 0.554250226 | 0.702043437 |
| 2010001A14Rik | -1.117457784 | -0.941229949 | -0.594692549 | 0.865526637 | 1.180604665 | 0.607248978 |
| Atf5 | -1.116424182 | -0.826765001 | -0.741420206 | 1.170962472 | 0.799462362 | 0.714184556 |
| Rps29 | -1.114795714 | -0.893179856 | -0.697851996 | 0.805184075 | 1.044975368 | 0.855668123 |
| H2-T22 | -1.113629292 | -0.980094119 | -0.531526981 | 0.940560972 | 0.516016482 | 1.168672937 |
| Thbs4 | -1.111274392 | -0.946259678 | -0.506459592 | 1.329219556 | 0.82232894 | 0.412445166 |
| Oasl2 | -1.102527225 | -0.728714344 | -0.809307436 | 0.739628312 | 1.300196543 | 0.600724152 |
| Anxa4 | -1.100485731 | -0.205176694 | -1.164349424 | 1.286118581 | 0.726522746 | 0.457370523 |
| Srgn | -1.099244498 | -0.72009158 | -0.668550896 | 1.442076738 | 0.838915013 | 0.206895222 |
| Plekhb2 | -1.098743869 | -1.111502625 | -0.376151472 | 1.157904299 | 0.548725764 | 0.879767903 |
| Rpl17 | -1.09823521 | -1.24535471 | -0.206954421 | 0.995208335 | 0.768550451 | 0.786785554 |
| Rpl36a | -1.097690627 | -1.105789589 | -0.43889106 | 0.694078854 | 0.971679292 | 0.97661313 |
| Carhsp1 | -1.097282796 | -0.861301303 | -0.661854561 | 1.328306288 | 0.554200946 | 0.737931426 |
| Rps21 | -1.093904267 | -0.992267047 | -0.613961934 | 0.814533165 | 0.96043793 | 0.925162154 |
| Psmb8 | -1.092100101 | -0.864440453 | -0.742390648 | 1.024254665 | 1.006967017 | 0.667709521 |
| Metrnl | -1.088607354 | -1.120084551 | -0.14146338 | 1.464758847 | 0.480452913 | 0.404943526 |
| Rhpn2 | -1.086428873 | -0.421366568 | -1.086428873 | 0.510487108 | 0.918951227 | 1.164785979 |
| Pcdhb14 | -1.086314112 | -0.445380441 | -0.836234705 | 1.428410108 | 0.000416482 | 0.939102668 |
| Rpl9 | -1.086281586 | -1.124592415 | -0.425274912 | 0.901739857 | 0.698084292 | 1.036324764 |
| Qdpr | -1.084841343 | -1.041331561 | -0.535095908 | 1.112594363 | 0.861966322 | 0.686708126 |
| Exoc3l4 | -1.083400865 | -0.935169784 | -0.523964053 | 1.344166957 | 0.875167817 | 0.323199928 |
| Ints9 | -1.081379439 | -0.863272742 | -0.558202232 | 1.502863402 | 0.587172149 | 0.412818861 |
| Eif4ebp1 | -1.080384831 | -0.808778536 | -0.685344673 | 1.428308314 | 0.65054994 | 0.495649786 |
| Rps23 | -1.078517241 | -1.096133307 | -0.487797221 | 0.775886927 | 0.854427232 | 1.032133609 |
| Cyp4v3 | -1.076571109 | -0.994979625 | -0.568999499 | 1.24474093 | 0.65473072 | 0.741078583 |
| Cd300c2 | -1.07405026 | -0.715027334 | -0.83146108 | 1.321325761 | 0.485372538 | 0.813840375 |
| Pold1 | -1.071583814 | -1.00984619 | -0.58196146 | 1.175335547 | 0.792353989 | 0.695701928 |
| Cox17 | -1.065372552 | -1.103986454 | -0.21813195 | 0.316664811 | 1.456231694 | 0.61459445 |
| Rpl37a | -1.06379105 | -1.056837419 | -0.5729146 | 0.951113335 | 0.854409106 | 0.888020627 |
| Fth1 | -1.062596526 | -0.970251428 | -0.614248225 | 1.262072113 | 0.681833631 | 0.703190436 |
| S1pr5 | -1.059267531 | -0.807006146 | -0.653255395 | 1.421145403 | 0.846742813 | 0.251640855 |
| Espn | -1.059215696 | -0.825528341 | -0.757027443 | 0.674480897 | 0.642006655 | 1.325283929 |
| Prr18 | -1.057564645 | -0.838685227 | -0.669063107 | 1.445696538 | 0.642602249 | 0.477014191 |
| Litaf | -1.056118336 | -1.096278654 | -0.399862031 | 1.27804728 | 0.83419075 | 0.44002099 |
| Gm15446 | -1.052425703 | -1.052425703 | -0.560597979 | 0.75789256 | 1.149664265 | 0.75789256 |
| H2-Aa | -1.050999658 | -0.66527681 | -0.940832677 | 0.876064128 | 1.217700844 | 0.563344172 |
| Bcas1 | -1.049188693 | -0.974163374 | -0.43336103 | 1.410493977 | 0.858986429 | 0.18723269 |
| Rpl37 | -1.04917284 | -0.985832326 | -0.678757226 | 0.817783158 | 0.931839028 | 0.964140206 |
| Rps27 | -1.048342634 | -1.065641735 | -0.571629839 | 0.769732458 | 1.031545382 | 0.884336367 |
| Pcdhb8 | -1.047823618 | -0.824596913 | -0.352616164 | 1.568609283 | 0.787443431 | -0.131016018 |
| Psmb9 | -1.047102794 | -1.047102794 | -0.526650224 | 1.035426939 | 1.109644806 | 0.475784066 |
| Ctsz | -1.044855641 | -0.925762416 | -0.668066609 | 1.280030469 | 0.82677036 | 0.531883836 |

| | | | | | | |
|---|---|---|---|---|---|---|
| Myo1d | -1.044805297 | -1.026725219 | -0.319341334 | 1.514295299 | 0.631879149 | 0.244697401 |
| Tmem106c | -1.044480523 | -1.119431588 | -0.4287611 | 1.256728648 | 0.681322245 | 0.654622317 |
| Gbp2 | -1.041659462 | -0.734652809 | -0.919593885 | 1.120591042 | 0.916196978 | 0.659118135 |
| Gstp1 | -1.040298088 | -1.150458389 | -0.393746037 | 1.22713216 | 0.756495731 | 0.600874623 |
| Tap2 | -1.038180376 | -0.842623858 | -0.638818096 | 1.517029592 | 0.508476174 | 0.494116564 |
| Rpl38 | -1.037941112 | -1.01231864 | -0.631242382 | 0.632352189 | 1.029330734 | 1.019819211 |
| Resp18 | -1.037848831 | -1.008246482 | -0.616803937 | 0.59452317 | 0.905125783 | 1.163250297 |
| Asb16 | -1.035767198 | -1.055501802 | -0.492024086 | 1.023934721 | 0.388033305 | 1.17132506 |
| Hspa2 | -1.034972204 | -1.079728762 | -0.445568445 | 1.290341409 | 0.850773615 | 0.419154387 |
| H2-T23 | -1.034709904 | -0.817105722 | -0.863419293 | 1.096297968 | 0.765462117 | 0.853474834 |
| H2-M3 | -1.032319343 | -0.700069676 | -0.882180672 | 1.138291666 | 1.112685514 | 0.36359251 |
| Rps28 | -1.030813651 | -0.981301616 | -0.682709073 | 0.702216027 | 1.118834526 | 0.873773788 |
| Slc25a34 | -1.029581922 | -1.029581922 | -0.544055913 | 1.328507871 | 0.580141715 | 0.694570171 |
| Eepd1 | -1.028957846 | -1.268364233 | 0.02092499 | 1.29359113 | 0.787821839 | 0.194984119 |
| Mbp | -1.028895623 | -1.011444314 | -0.55830744 | 1.351650771 | 0.65656584 | 0.590430766 |
| Pttg1 | -1.027163782 | -0.917038604 | -0.749013998 | 1.139307485 | 0.914094549 | 0.63981435 |
| Egr2 | -1.025158606 | -0.596909217 | -0.914671405 | 0.382739618 | 1.457643119 | 0.696356491 |
| Znfx1 | -1.024720039 | -0.783057793 | -0.650947613 | 1.583790851 | 0.332854185 | 0.542080409 |
| Ifit3 | -1.020895774 | -0.811742548 | -0.870394879 | 0.7871707 | 1.165724819 | 0.750137683 |
| Lgals3bp | -1.019472426 | -0.848782079 | -0.825861146 | 1.155851909 | 0.908912087 | 0.629351655 |
| Cmtm5 | -1.016937166 | -1.010466705 | -0.494638629 | 1.447843061 | 0.421068008 | 0.653131433 |
| Uap1l1 | -1.016578592 | -1.2458319 | -0.180605952 | 1.286240413 | 0.749227912 | 0.407548118 |
| Rhog | -1.013953941 | -1.060326486 | -0.531391652 | 1.315639589 | 0.678397543 | 0.611634946 |
| Lap3 | -1.013771377 | -1.251620976 | -0.312632977 | 1.074782628 | 0.858733771 | 0.644508931 |
| Lgals1 | -1.013237202 | -0.85135841 | -0.858737021 | 0.980322334 | 0.996075668 | 0.746934631 |
| Rps19 | -1.01266395 | -1.073746329 | -0.616860309 | 0.954501024 | 0.895529583 | 0.853239981 |
| Efhd1 | -1.00846584 | -1.105662528 | -0.505425422 | 1.230666008 | 0.808480521 | 0.580407261 |
| C1qa | -1.002461593 | -0.94961237 | -0.697144449 | 1.305372904 | 0.68492131 | 0.658924198 |
| Ctss | -1.001157372 | -0.951265442 | -0.731090878 | 1.196332255 | 0.640588467 | 0.846592969 |
| Trf | -0.999746868 | -0.931382524 | -0.698368543 | 1.314751798 | 0.819334759 | 0.495411378 |
| Stat6 | -0.998050331 | -0.686881502 | -0.591301455 | 1.319319821 | 1.179885792 | -0.222972326 |
| Dnajc12 | -0.996953318 | -1.459897808 | 0.331596258 | 0.978509082 | 0.299663722 | 0.847082063 |
| Rpl30 | -0.992090671 | -1.190941658 | -0.478232993 | 0.925522369 | 0.814421405 | 0.921321547 |
| Smim1 | -0.989177228 | -0.786165848 | -0.862619396 | 1.346192517 | 0.565915145 | 0.72585481 |
| Ier5 | -0.986844982 | -1.227556388 | -0.329960847 | 1.110883793 | 0.989571705 | 0.443906719 |
| Adssl1 | -0.983963215 | -1.130446515 | -0.454787172 | 1.332904795 | 0.673964584 | 0.562327523 |
| Pcdhb9 | -0.983897494 | -1.176845242 | -0.469717076 | 0.920073201 | 1.097165401 | 0.61322121 |
| Gjc2 | -0.983400591 | -0.983400591 | -0.595239282 | 1.135282622 | 1.16329622 | 0.263461621 |
| Rps6 | -0.981944774 | -1.032505854 | -0.680493138 | 1.136785565 | 0.764326088 | 0.793832113 |
| Bhlhe40 | -0.97977862 | -0.875538258 | -0.811438308 | 1.245741307 | 0.874096152 | 0.546917728 |
| Cyb5r1 | -0.978023367 | -1.345431065 | -0.232233931 | 0.868612985 | 0.873141435 | 0.813933941 |
| Vamp1 | -0.975892178 | -0.960469887 | -0.59766879 | 1.341707164 | 0.95458064 | 0.237743051 |
| S100b | -0.974978107 | -1.219469672 | -0.263839897 | 1.238920747 | 0.937057446 | 0.282309483 |
| Cdc42ep1 | -0.973736083 | -0.967910125 | -0.641590707 | 1.424534124 | 0.616471783 | 0.542231008 |
| Lyz2 | -0.973450155 | -0.887946095 | -0.774145995 | 1.348934341 | 0.73641867 | 0.550189234 |
| Tmem63a | -0.973198204 | -1.066658727 | -0.522040733 | 1.348429997 | 0.816593956 | 0.396873711 |
| Tmem88b | -0.972848821 | -0.879659594 | -0.738631175 | 1.301726736 | 0.967508928 | 0.321903925 |
| Hebp1 | -0.971755685 | -0.91078063 | -0.781488989 | 1.224001456 | 0.920549438 | 0.519474409 |
| Ephx2 | -0.97151891 | -1.083459125 | -0.549725689 | 1.329036133 | 0.643667197 | 0.632000392 |
| Macrod1 | -0.969600294 | -0.671592135 | -1.043877232 | 1.160523641 | 0.83377194 | 0.69077408 |
| Ube2l6 | -0.965816935 | -0.827173216 | -0.881093326 | 1.072524146 | 1.095612228 | 0.505947103 |
| Rpl26 | -0.963176996 | -1.195356101 | -0.494044305 | 0.92971991 | 0.701998495 | 1.020858998 |
| Lcp1 | -0.962602663 | -1.020382405 | -0.648776676 | 1.207257209 | 0.976097044 | 0.44840749 |
| Fa2h | -0.960321705 | -0.852204866 | -0.632148973 | 1.489415089 | 0.849986366 | 0.105274089 |
| B2m | -0.959314877 | -0.963125902 | -0.745698383 | 1.240819996 | 0.851106159 | 0.576213006 |
| Spag6 | -0.957299762 | -0.921896502 | -0.852272464 | 0.809146706 | 1.010841397 | 0.911480625 |
| Tifa | -0.956556436 | -1.017925525 | -0.26837791 | 0.821928788 | 1.51411696 | -0.093185877 |
| Galnt6 | -0.954903509 | -0.920755001 | -0.752825074 | 1.316225683 | 0.856333193 | 0.455924709 |
| Ppp1r14a | -0.953751632 | -0.798990585 | -0.859105381 | 1.286386447 | 0.963310091 | 0.362151059 |
| Tmbim1 | -0.951045985 | -0.963951694 | -0.764336149 | 1.237932002 | 0.794198076 | 0.64720375 |
| Rps3a1 | -0.946047623 | -1.311195407 | -0.348867876 | 0.860968784 | 0.869110758 | 0.876031363 |
| Rpl22l1 | -0.943169424 | -1.315966883 | -0.211772729 | 0.436092307 | 1.208979309 | 0.825837419 |
| Wfdc17 | -0.941183428 | -0.941183428 | -0.418600288 | 1.050886082 | -0.138964842 | 1.389045904 |
| Csf1 | -0.936554908 | -0.858620345 | -0.792697965 | 1.453996602 | 0.579621659 | 0.554254956 |
| Nudt17 | -0.936374822 | -0.887220838 | -0.862811716 | 0.710389698 | 1.247451385 | 0.728566293 |
| Grhpr | -0.935291751 | -0.957841351 | -0.669613976 | 1.47000833 | 0.519696023 | 0.573042725 |
| Ppp1r3c | -0.933494897 | -1.302146154 | -0.28835404 | 1.114919411 | 0.943349679 | 0.465726001 |
| Gfap | -0.932518391 | -1.00242198 | -0.727607224 | 1.28036164 | 0.71939857 | 0.662787386 |
| Hba-a2 | -0.93110817 | -0.74868866 | -0.707749784 | 0.515035842 | 1.661921595 | 0.210589178 |
| Fn3k | -0.930894297 | -1.075639191 | -0.701642088 | 0.913786876 | 1.037216961 | 0.757171738 |
| Cox6a2 | -0.929315019 | -0.996016509 | -0.679920889 | 1.089438143 | 1.174026332 | 0.341787943 |
| Cd83 | -0.929127884 | -1.096795769 | -0.645617746 | 1.106638222 | 0.598292675 | 0.966610502 |
| Cd52 | -0.92448043 | -0.883099906 | -0.883099906 | 1.117939065 | 1.008923827 | 0.56381735 |

| | | | | | | |
|---|---|---|---|---|---|---|
| Tyrobp | -0.923986231 | -0.951776442 | -0.783430722 | 1.308865819 | 0.705041382 | 0.645286193 |
| Rnf213 | -0.919490693 | -0.23529136 | -1.33048417 | 1.005630141 | 1.073759104 | 0.405876977 |
| Ighm | -0.916967375 | -0.807420169 | -0.715362485 | 1.066999253 | 1.362685327 | 0.010065449 |
| Cd74 | -0.916711428 | -0.906532882 | -0.862174047 | 1.029470871 | 1.113983263 | 0.541964223 |
| Serpina3n | -0.915754294 | -0.910814121 | -0.675501873 | 1.557701313 | 0.511928485 | 0.43244049 |
| Relt | -0.914197157 | -1.367696179 | -0.114468964 | 1.192326329 | 0.343122069 | 0.860913902 |
| P3h4 | -0.914011434 | -1.00277693 | -0.771636914 | 1.178570669 | 0.886720147 | 0.623134463 |
| Fcer1g | -0.912270488 | -1.150464142 | -0.576005028 | 1.208495523 | 0.594390032 | 0.835854103 |
| Ccdc190 | -0.912233638 | -1.094276697 | -0.568694437 | 1.330541775 | 0.848208161 | 0.396454836 |
| Mthfd2 | -0.905784177 | -1.120134512 | -0.641001844 | 1.143507182 | 0.91331067 | 0.610102681 |
| Lgi4 | -0.901275105 | -1.017066804 | -0.652533022 | 1.446805682 | 0.597151849 | 0.526917399 |
| Ier3 | -0.898329507 | -1.039577439 | -0.767637446 | 1.133790366 | 0.751747532 | 0.820006494 |
| Dipk1c | -0.891418061 | -0.837096458 | -0.968957131 | 1.1976849 | 0.692290546 | 0.807496204 |
| C4b | -0.890717862 | -0.785956851 | -0.986043977 | 1.298387557 | 0.689265724 | 0.675065409 |
| Capg | -0.889859304 | -0.889859304 | -0.873085242 | 1.203812868 | 1.002946696 | 0.446044284 |
| Cst7 | -0.889658229 | -0.889658229 | -0.889658229 | 1.279853969 | 0.801027118 | 0.5880936 |
| Arhgef10 | -0.887266909 | -0.847967931 | -0.647401504 | 1.681347955 | 0.380186243 | 0.321102145 |
| Scnn1a | -0.88662116 | -0.911114437 | -0.79867873 | 1.063425006 | 0.297461179 | 1.235528142 |
| Nagpa | -0.886356848 | -1.051021054 | -0.649303359 | 1.342200449 | 0.385319385 | 0.859161427 |
| Myrf | -0.885803102 | -0.858563269 | -0.701186043 | 1.598001687 | 0.615842541 | 0.231708187 |
| Lag3 | -0.885097658 | -0.997594683 | -0.812210049 | 1.20465813 | 0.752423873 | 0.737820389 |
| Irf8 | -0.883168006 | -1.167367537 | -0.356272341 | 1.525418515 | 0.354057309 | 0.527332061 |
| Ankrd22 | -0.882844295 | -0.882844295 | -0.400209573 | 0.632388029 | 1.690135518 | -0.156625384 |
| Mpeg1 | -0.879101572 | -0.96976975 | -0.76372051 | 1.393351988 | 0.706458396 | 0.512781448 |
| H2-K1 | -0.879081827 | -0.993595767 | -0.842401812 | 1.102903821 | 0.8909108 | 0.721264785 |
| Bc1 | -0.87708133 | -0.720299234 | -0.826419913 | 0.112457855 | 1.571079757 | 0.740262865 |
| Cbr3 | -0.876607514 | -1.100776732 | -0.727277729 | 0.936055516 | 1.043571283 | 0.725035176 |
| Gm16867 | -0.874640237 | -0.874640237 | -0.874640237 | 0.39009061 | 0.946850024 | 1.286980077 |
| Stard10 | -0.874470499 | -1.314182091 | -0.286367807 | 1.328864529 | 0.614926967 | 0.531228902 |
| Ccl12 | -0.873471858 | -1.118793559 | -0.455114968 | 0.728976826 | 0.238574821 | 1.479828738 |
| Unc5b | -0.870270482 | -1.010122878 | -0.595461456 | 1.472236603 | 0.815338485 | 0.188279727 |
| C3ar1 | -0.869542363 | -0.484851048 | -1.076895906 | 1.540270185 | 0.64516046 | 0.245858672 |
| H2-D1 | -0.867347837 | -0.927147065 | -0.885422709 | 1.229498116 | 0.867703084 | 0.582716412 |
| Ggta1 | -0.866196343 | -1.108076325 | -0.498903136 | 1.507887635 | 0.611777098 | 0.353511071 |
| Hmox1 | -0.865870952 | -0.865870952 | -0.788208266 | 1.552386183 | 0.508979466 | 0.45858452 |
| Trem2 | -0.861111666 | -1.033208911 | -0.768089599 | 1.28470278 | 0.716279453 | 0.661427944 |
| Ifi27l2a | -0.858136976 | -0.629920485 | -1.139980895 | 0.586012796 | 1.284545966 | 0.757479595 |
| Syde1 | -0.85808835 | -0.784612008 | -0.60324471 | 1.658121907 | 0.719368928 | -0.131545766 |
| Scly | -0.855154786 | -0.715652751 | -0.439689993 | 1.874092848 | 0.212592474 | -0.076187791 |
| Rpl35 | -0.854504197 | -1.280011432 | -0.511775315 | 0.778774825 | 0.83582726 | 1.031688858 |
| Lgals3 | -0.848955156 | -1.084123649 | -0.723222177 | 1.265451047 | 0.773855247 | 0.616994688 |
| Irgm2 | -0.847993873 | -0.97982281 | -0.847993873 | 1.177596106 | 0.533811864 | 0.964402586 |
| Gstt1 | -0.835404652 | -1.03481273 | -0.826202707 | 1.12498859 | 0.941203627 | 0.630227872 |
| Plaat3 | -0.832669965 | -0.905442983 | -0.862279715 | 1.347088056 | 0.3590005 | 0.894304107 |
| Arl4d | -0.832429012 | -1.338318324 | -0.342866373 | 1.088397011 | 1.027393421 | 0.397823277 |
| Nfil3 | -0.832396219 | -0.353270319 | -1.297721769 | 0.921395205 | 1.244329302 | 0.317663801 |
| Cyba | -0.824602372 | -0.878261521 | -0.953613498 | 1.316078588 | 0.736951176 | 0.603447628 |
| Gstt3 | -0.816782967 | -1.271078956 | -0.523917119 | 1.205426795 | 0.722767506 | 0.683584742 |
| Lrrc38 | -0.811631188 | -0.884855636 | -0.884855636 | 1.40079148 | 0.831365455 | 0.349185526 |
| Plekhg3 | -0.809305166 | -0.8019754 | -1.05649657 | 1.110543462 | 1.043078994 | 0.51415468 |
| Depp1 | -0.808689355 | -0.979034242 | -0.730686064 | 0.714790347 | 1.500209524 | 0.30340979 |
| Gjb1 | -0.804350385 | -1.175375136 | -0.574579585 | 1.379629056 | 0.742360291 | 0.432315758 |
| Hba-a1 | -0.79879985 | -0.913846846 | -0.61398843 | 1.106125219 | 1.378852941 | -0.158343034 |
| Ly86 | -0.796981223 | -1.380028101 | -0.385671212 | 1.087392309 | 0.880719929 | 0.594568297 |
| Hhatl | -0.788327525 | -1.201422504 | -0.534091817 | 1.431398328 | 0.592330419 | 0.500113098 |
| Tmem192 | -0.787835688 | -0.848435178 | -1.088075462 | 0.888246576 | 0.896240025 | 0.939859726 |
| Padi2 | -0.783123143 | -1.277067461 | -0.609808891 | 0.909823227 | 0.820668403 | 0.939507865 |
| Zfas1 | -0.780897117 | -1.146383241 | -0.786897598 | 0.917775319 | 0.876354423 | 0.920048214 |
| Il33 | -0.780184944 | -1.260914405 | -0.521360977 | 1.315255198 | 0.728322418 | 0.51888271 |
| Bcl2a1b | -0.778704655 | -1.100894458 | -0.812689296 | 1.160729049 | 0.788775352 | 0.742784008 |
| Dusp28 | -0.775417834 | -1.336390459 | -0.531699092 | 0.896797034 | 0.860309983 | 0.886400368 |
| Evi2a | -0.773286746 | -1.248001095 | -0.654265377 | 1.025880196 | 0.784111985 | 0.865561037 |
| Fgfrl1 | -0.772363102 | -1.031313981 | -0.786715265 | 1.428564866 | 0.529890801 | 0.63193668 |
| Tap1 | -0.766160519 | -0.895494163 | -0.99227704 | 0.924074797 | 1.236652701 | 0.493204224 |
| Sfrp5 | -0.764933494 | -0.901084834 | -1.042100192 | 0.68915226 | 0.965285592 | 1.053680668 |
| Hsf4 | -0.763768557 | -1.053031818 | -0.89573537 | 1.011552712 | 0.717207617 | 0.983775416 |
| Plekhh1 | -0.763641058 | -0.914286216 | -0.781294156 | 1.596175658 | 0.590128935 | 0.272916837 |
| Nkx6-2 | -0.7606365 | -1.074330266 | -0.608663585 | 1.590906885 | 0.459692357 | 0.393031109 |
| Gm11681 | -0.757099834 | -1.064837354 | -0.64385192 | 1.455213981 | 0.858909664 | 0.151665464 |
| Pcsk6 | -0.749987867 | -1.012058957 | -0.76172958 | 1.522066169 | 0.415339635 | 0.5863706 |
| Gngt2 | -0.740704237 | -0.658243503 | -1.105883416 | 1.39283165 | 0.211875403 | 0.900124104 |
| Mansc1 | -0.73929037 | -1.283079883 | -0.279242266 | 1.424124456 | 0.836271642 | 0.041216421 |
| Gstm7 | -0.729594286 | -1.23895625 | -0.515213711 | 1.464978455 | 0.476909819 | 0.541875972 |

| | | | | | | |
|---|---|---|---|---|---|---|
| Tmem88 | -0.726915349 | -0.345555568 | -1.322073096 | 0.980743774 | 1.273924918 | 0.13987532 |
| Naga | -0.725816542 | -1.278923833 | -0.556484667 | 1.330630013 | 0.607857759 | 0.622737269 |
| Cd84 | -0.714295434 | -1.198121633 | -0.658667448 | 1.311937811 | 0.860373861 | 0.398772843 |
| Syt2 | -0.712248855 | -1.137932322 | -0.539841594 | 1.506679354 | 0.792493729 | 0.090849688 |
| Ccdc88b | -0.706801271 | -1.02333389 | -0.299698539 | 1.818849511 | -0.022823515 | 0.233807704 |
| Ehd4 | -0.705491099 | -0.76471308 | -1.064453405 | 1.49957286 | 0.514695683 | 0.520389041 |
| H2-Ab1 | -0.705179952 | -0.819610402 | -1.106521615 | 1.061857731 | 1.13646196 | 0.432992278 |
| Arsg | -0.699191615 | -1.311440698 | -0.472007971 | 1.380625594 | 0.717015833 | 0.384998856 |
| Bhlhe41 | -0.698987096 | -1.144177526 | -0.818510469 | 1.20711696 | 0.820323163 | 0.634234967 |
| Endou | -0.698256119 | -0.281943379 | -1.301150993 | 0.969647873 | 1.341217466 | -0.029514849 |
| Rgs16 | -0.694437297 | -1.051913326 | -0.926741198 | 0.881528346 | 0.603853767 | 1.187709708 |
| Ninj2 | -0.691067124 | -1.560573058 | -0.219205081 | 0.823814153 | 0.779009385 | 0.868021723 |
| Ppie | -0.681923642 | -1.215616879 | -0.786947085 | 0.955878022 | 0.98723098 | 0.741378604 |
| Gpnmb | -0.675268141 | -0.795900068 | -1.029907184 | 1.450951085 | 0.835873695 | 0.214250612 |
| Mafb | -0.674631433 | -1.310827848 | -0.636333016 | 1.114091136 | 0.605690837 | 0.902010324 |
| Ccl6 | -0.674067756 | -1.311235232 | -0.606035276 | 1.177571182 | 0.897898597 | 0.515868485 |
| Cd37 | -0.673629307 | -0.934611847 | -1.001698257 | 1.022886283 | 1.219941673 | 0.367111456 |
| Hvcn1 | -0.67110814 | -0.931820955 | -0.822958702 | 1.425716799 | 0.985483862 | 0.014687137 |
| Tmem98 | -0.668913605 | -0.84236969 | -0.98626837 | 1.453155626 | 0.848694406 | 0.195701634 |
| Erbb3 | -0.667428006 | -1.213370914 | -0.510192066 | 1.527484099 | 0.672528435 | 0.190978452 |
| Opalin | -0.663426485 | -0.995380157 | -0.663426485 | 1.610762339 | 0.730767585 | -0.019296797 |
| Ctxn2 | -0.655915031 | -1.45228754 | -0.438529742 | 0.588168815 | 0.932377616 | 1.026185882 |
| Chil1 | -0.654404171 | -1.068562465 | -0.583138667 | 1.577284419 | -0.045749682 | 0.774570566 |
| Aldh3b1 | -0.652640677 | -1.552694141 | -0.105930849 | 1.267969237 | 0.521648215 | 0.521648215 |
| Cxcl16 | -0.64638984 | -0.812116838 | -0.932824989 | 1.634779854 | 0.594832438 | 0.161719375 |
| Fcgr2b | -0.641585857 | -0.948927645 | -1.109088893 | 1.01139433 | 0.773804614 | 0.91440345 |
| Lefty1 | -0.635525232 | -0.128382598 | -1.335256658 | 1.500587546 | 0.72695954 | -0.128382598 |
| Ifih1 | -0.60834734 | -0.634590689 | -1.332445419 | 0.658073717 | 1.259236015 | 0.658073717 |
| Cyp11a1 | -0.594630594 | 1.023644438 | 1.413469827 | -0.322068589 | -0.347224256 | -1.173190824 |
| Nectin4 | -0.583894476 | -1.553524221 | -0.239790958 | 1.101652883 | 0.354080996 | 0.921475775 |
| Smok2b | -0.578092377 | -1.042577498 | -0.924674849 | 0.451311237 | 1.451768418 | 0.642265069 |
| Aif1 | -0.57518964 | -1.54770718 | -0.336615725 | 0.902374251 | 1.037248576 | 0.519889717 |
| Batf3 | -0.569345644 | -1.041857946 | -0.408961521 | 0.525412696 | 1.754830211 | -0.260077797 |
| Itgb2 | -0.562351238 | -0.945265998 | -0.975732126 | 1.524625081 | 0.643864932 | 0.314859349 |
| Kazald1 | -0.53897735 | -1.430838417 | -0.53897735 | 0.448336307 | 1.185085889 | 0.87537092 |
| Gm30082 | -0.533293249 | -0.608187628 | -1.187116981 | 1.637955557 | 0.432118597 | 0.258523703 |
| Tnnt2 | -0.530615863 | -1.42904082 | -0.530615863 | 0.738374118 | 0.479108214 | 1.272790214 |
| Parp4 | -0.484002475 | -0.603108017 | -1.303481977 | 1.478175727 | 0.681687221 | 0.230729522 |
| Pir | -0.483708522 | -1.671096707 | -0.23513882 | 0.823928579 | 0.702041487 | 0.863973983 |
| Evpl | -0.432716741 | -1.288814972 | -0.814183184 | 1.065392957 | 1.10450666 | 0.365815281 |
| Ifit3b | -0.423388254 | -1.042017626 | -1.161295969 | 0.7916273 | 1.112358017 | 0.722716533 |
| Gpr62 | -0.40758424 | -1.222580428 | -0.432497599 | 1.614565209 | 0.695695491 | -0.247598433 |
| B3galt5 | -0.404157788 | -1.303569618 | -0.585195848 | 1.597889032 | 0.355356376 | 0.339677846 |
| Zc3hav1 | -0.398810942 | -1.118129716 | -1.118129716 | 0.896591113 | 0.982135637 | 0.756343624 |
| Serpinb1a | -0.390352562 | -1.534671236 | -0.412717233 | 0.93575251 | 1.182478887 | 0.219509634 |
| Sncg | -0.375565956 | -1.000762881 | -1.000762881 | 1.586220894 | 0.511759244 | 0.279111581 |
| AW112010 | -0.363825027 | -1.587338932 | -0.485544547 | 1.005304779 | 0.91209044 | 0.519313288 |
| Zic4 | -0.295633302 | 1.663248403 | 0.687179895 | -1.090023755 | -0.583494311 | -0.38127693 |
| Pcdhb13 | -0.276882163 | -1.085192139 | -1.102303458 | 1.276084187 | 0.907363737 | 0.280929836 |
| Pigz | -0.254480757 | -0.993959296 | -0.410574279 | 1.924835981 | 0.005633561 | -0.27145521 |
| Insc | -0.206086327 | -1.066545933 | -0.878116488 | 1.614710747 | 0.653998577 | -0.117960576 |
| Adcyap1 | -0.154111553 | 1.30830626 | 1.169756124 | -0.791473961 | -0.547627754 | -0.984849116 |
| Polr2k | -0.106750229 | -1.023163025 | -1.249605543 | 0.5722886 | 1.361694276 | 0.44553592 |
| Nts | -0.064958333 | 0.81326139 | 1.440367163 | -1.047140509 | -1.076571379 | -0.064958333 |
| Col26a1 | -0.061278181 | 1.296518358 | 0.980536369 | -0.23298069 | -0.582855486 | -1.399940371 |
| Tmc6 | -0.057182252 | -1.434366622 | -0.866014944 | 1.05975293 | 0.98365003 | 0.314160859 |
| P2ry1 | 0.096404565 | 0.843283434 | 1.130052867 | -0.895338915 | -1.45648746 | 0.282085509 |
| Traf3 | 0.105310071 | 1.054835197 | 1.327322712 | -0.874789614 | -0.65009532 | -0.962583047 |
| Ranbp3l | 0.138035114 | 1.122744941 | 1.211550626 | -1.145892324 | -0.490380221 | -0.836058136 |
| Mirt1 | 0.141255759 | 0.808870356 | 1.341129486 | -1.436283522 | -0.205283121 | -0.649688958 |
| Dnhd1 | 0.160849617 | 1.476618397 | 0.674756664 | -0.86377003 | -0.203017261 | -1.245437387 |
| Clmp | 0.179451326 | 1.510767505 | 0.637474904 | -0.901569568 | -0.239109683 | -1.187014485 |
| Efna1 | 0.206005992 | -1.452485324 | -0.935634813 | 0.98075779 | 0.98075779 | 0.220598564 |
| Dnase1l2 | 0.242370819 | 1.26343333 | 1.051336215 | -1.036901651 | -0.69319828 | -0.827040434 |
| Pcdhb22 | 0.243822339 | 0.908560518 | 1.347861014 | -0.704098756 | -0.590245136 | -1.205899979 |
| Gm12371 | 0.33590122 | 0.86132695 | 0.943560178 | -1.620396757 | 0.236740364 | -0.757131955 |
| Arhgef33 | 0.350543892 | 1.577956617 | 0.509288313 | -0.845913058 | -0.526499407 | -1.065376357 |
| Phtf2 | 0.373123147 | 0.8079437 | 1.171620785 | -1.571725392 | -0.563742646 | -0.217219594 |
| Mterf1a | 0.375375404 | 1.214562933 | 0.753829878 | -0.162270173 | -0.638050314 | -1.543447729 |
| B930059L03Rik | 0.40217252 | 0.146373591 | 1.718672888 | -0.822951078 | -0.45493352 | -0.989334401 |
| Grip1 | 0.425850041 | 0.744338783 | 1.408753759 | -0.993388907 | -0.927336511 | -0.658217167 |
| Gm6569 | 0.445245631 | 1.011617071 | 1.106164125 | -1.330126479 | -0.503467712 | -0.729432636 |
| Ppil6 | 0.448301692 | 1.130019851 | 1.055367246 | -1.050650103 | -0.951846083 | -0.631192602 |

| | | | | | | |
|---|---|---|---|---|---|---|
| Sox4 | 0.456876071 | 1.243767382 | 0.91250169 | -0.590324711 | -1.104316362 | -0.91850407 |
| 4933439C10Rik | 0.460644819 | 1.405677763 | 0.62246214 | -0.49940944 | -1.307897208 | -0.681478073 |
| 2810029C07Rik | 0.497018679 | 0.827281468 | 0.984761374 | -1.040117023 | -1.412890309 | 0.14394581 |
| Stx2 | 0.497818167 | 1.443246089 | 0.405179567 | -0.173931394 | -0.846026515 | -1.326285913 |
| Purg | 0.502452719 | 1.003070785 | 1.083451299 | -1.308589422 | -0.77327896 | -0.507106421 |
| Efcab6 | 0.506630818 | 0.487411052 | 1.45308096 | -0.64858982 | -1.321334128 | -0.477198882 |
| Gpr101 | 0.509616492 | 1.112835159 | 0.810090891 | -1.568610206 | -0.509148325 | -0.354784011 |
| Spink10 | 0.510864183 | 1.186811151 | 0.879587894 | -1.29636034 | -0.473245013 | -0.807657874 |
| Col5a2 | 0.51128852 | 1.422401164 | 0.311985439 | 0.016628208 | -0.959775782 | -1.302527548 |
| Col11a1 | 0.513351508 | 1.224402316 | 0.865644357 | -0.707695382 | -0.639394157 | -1.256308641 |
| Plpp5 | 0.526176926 | 0.908694817 | 1.07554805 | -0.252763022 | -0.87563698 | -1.38201979 |
| Zfp618 | 0.540257135 | 0.695594203 | 1.294154425 | -1.29937575 | -0.387463295 | -0.843166717 |
| Dancr | 0.545783384 | 1.316659636 | 0.377416726 | -1.596664764 | -0.508541674 | -0.134653308 |
| 2700049A03Rik | 0.55219135 | 0.894897929 | 1.121771194 | -0.374932961 | -1.307484669 | -0.886442843 |
| Fam184b | 0.56931995 | 1.40963664 | 0.13312769 | -0.088736889 | -0.456590463 | -1.566756928 |
| Cwh43 | 0.574622434 | 1.097966518 | 0.775061111 | -0.057626243 | -1.149563352 | -1.240460469 |
| Chfr | 0.57694664 | 1.239426312 | 0.751483432 | -0.640413747 | -0.577340473 | -1.350102164 |
| Insyn2b | 0.589093015 | 1.108482797 | 0.984889325 | -0.77796817 | -1.085364046 | -0.819132921 |
| C130026L21Rik | 0.591628102 | 0.634159991 | 0.940234465 | -1.797532004 | -0.364193179 | -0.004297376 |
| Ndst4 | 0.613571659 | 1.162406977 | 0.799900191 | -0.940592632 | -0.364484982 | -1.270801213 |
| Myo19 | 0.626102757 | 1.385060994 | 0.44498629 | -0.455986208 | -1.376470997 | -0.623692836 |
| Zfp760 | 0.630853738 | 1.265142958 | 0.759535983 | -0.996176068 | -0.663180544 | -0.996176068 |
| Sirt4 | 0.63255437 | 1.211420332 | 0.295843866 | -0.239186453 | -0.181745703 | -1.718886411 |
| Rgs5 | 0.640580837 | 0.69481298 | 1.1050401 | -1.399550978 | -0.081513283 | -0.959369656 |
| Cfp | 0.65973976 | 1.1699292 | 0.863923431 | -0.888282248 | -1.027209698 | -0.778100446 |
| Sox17 | 0.659966288 | 0.821259361 | 0.568995927 | -0.046554549 | -0.119588338 | -1.884078688 |
| Bglap3 | 0.663675162 | 1.084956093 | 0.889759904 | -0.777866025 | -0.573274869 | -1.287250264 |
| Megf11 | 0.665927221 | 0.715770004 | 1.198154045 | -1.188636924 | -0.357935209 | -1.033279136 |
| Sh3rf3 | 0.702668661 | 0.934426501 | 0.899494727 | -0.869474616 | -1.415962276 | -0.251152997 |
| Car8 | 0.715058942 | 1.031690332 | 0.749283154 | -1.044317769 | -0.126464638 | -1.32525002 |
| Jarid2 | 0.72048488 | 0.813171583 | 1.156277142 | -0.684850033 | -0.960846166 | -1.044237406 |
| Fras1 | 0.726073465 | 1.270120052 | 0.469323663 | -0.929847243 | -0.222128682 | -1.313541255 |
| Igflr1 | 0.726941694 | 1.183150781 | 0.546234598 | -1.513283096 | -0.610529305 | -0.332514672 |
| Slco1a4 | 0.734258463 | 0.587675683 | 1.173313149 | -0.866718926 | -0.23911897 | -1.389409398 |
| P3h3 | 0.741590541 | 0.753440052 | 0.927937268 | -0.208223514 | -0.594829156 | -1.619915192 |
| Sfxn2 | 0.743703436 | 1.013817176 | 0.849717634 | -1.382503491 | -0.478658062 | -0.746076694 |
| 5430416N02Rik | 0.749969643 | 1.165779726 | 0.749969643 | -1.199541265 | -0.699097267 | -0.767080481 |
| Tnc | 0.762465008 | 0.67454684 | 0.696762803 | -1.127313702 | 0.418950979 | -1.425411928 |
| Cacna1g | 0.765254877 | 0.91271077 | 0.999158473 | -0.536815866 | -1.02327457 | -1.117033683 |
| Cd59a | 0.765406473 | 0.51391097 | 1.283399202 | -1.068859292 | -0.397164575 | -1.096692778 |
| Dlg2 | 0.768276406 | 0.845738646 | 1.080510019 | -1.04041451 | -1.022596041 | -0.631514521 |
| Carmil3 | 0.772163915 | 1.103981432 | 0.689635222 | -0.270576143 | -1.114450913 | -1.180753513 |
| Wdfy1 | 0.777801284 | 0.952716936 | 0.989657632 | -1.068790683 | -0.793417512 | -0.857967657 |
| Muc3a | 0.780380921 | 1.190095738 | 0.709592097 | -0.687147208 | -1.077446498 | -0.91547505 |
| Tgfb2 | 0.785808222 | 0.785808222 | 0.977907789 | -0.202997483 | -1.088394765 | -1.258131985 |
| Ahr | 0.793213229 | 0.836280129 | 1.03721564 | -0.690556423 | -1.27632476 | -0.699827815 |
| Eml4 | 0.794836479 | 0.768923685 | 1.063157821 | -1.135743423 | -0.395606656 | -1.095567906 |
| P2ry12 | 0.799600472 | 0.736362661 | 1.159584021 | -0.893577776 | -0.740133898 | -1.06183548 |
| Ankrd16 | 0.807017701 | 0.847801684 | 0.897775639 | -1.432583686 | -0.291263301 | -0.828748037 |
| 2810049E08Rik | 0.814904677 | 0.670729705 | 1.164292532 | -1.22525579 | -0.797127285 | -0.627543839 |
| Myt1 | 0.814917984 | 0.959035089 | 0.909299483 | -0.572108795 | -0.926029679 | -1.185114082 |
| Amer1 | 0.81748476 | 0.693001092 | 1.017225642 | -0.92679044 | -1.381969562 | -0.218951492 |
| Robo3 | 0.819083496 | 0.819083496 | 0.740417929 | -0.321297509 | -0.361358945 | -1.695928467 |
| Fabp7 | 0.823015222 | 0.278219778 | 1.411110883 | -0.946261489 | -0.525860542 | -1.040223852 |
| Cnbd2 | 0.824172488 | 0.663970398 | 0.859966966 | -0.06275997 | -0.630332778 | -1.655017104 |
| Cntn6 | 0.826144653 | 0.675129968 | 1.038499727 | -1.365803633 | -0.248843617 | -0.925127098 |
| Gng2 | 0.826261411 | 0.680066327 | 1.092927691 | -1.372852877 | -0.504678693 | -0.721723859 |
| Usp40 | 0.828768637 | 1.149763585 | 0.63245539 | -0.775149261 | -0.527292228 | -1.308546123 |
| Psme4 | 0.828828397 | 1.087989563 | 0.792385909 | -1.073316681 | -0.742529317 | -0.893357871 |
| AW551984 | 0.83324397 | 0.346757626 | 0.814139616 | -1.863503619 | 0.078662959 | -0.209300551 |
| Mzf1 | 0.834429055 | 0.234062434 | 1.414984987 | -0.609136399 | -1.181237551 | -0.693102526 |
| Nav3 | 0.840778486 | 0.882562555 | 1.007428776 | -0.947471118 | -0.967423651 | -0.815875048 |
| Cdkl2 | 0.843574464 | 0.867324496 | 0.965254427 | -1.257252306 | -0.617009386 | -0.801891695 |
| Ccnl2 | 0.849287444 | 1.170819904 | 0.680438639 | -0.869820414 | -0.836213454 | -0.994512119 |
| Myo9b | 0.849462036 | 1.248514015 | 0.379242572 | -0.231177128 | -1.121019245 | -1.12502225 |
| Glra2 | 0.850650938 | 0.575259737 | 1.229346898 | -0.939169243 | -1.057614925 | -0.658473405 |
| Raver2 | 0.85265113 | 0.710267468 | 1.078270126 | -0.597151198 | -0.738994512 | -1.305043014 |
| Asxl3 | 0.855806723 | 0.978778062 | 0.859879624 | -1.084157754 | -0.58350767 | -1.026798984 |
| Cxadr | 0.855827941 | 1.068539779 | 0.786955341 | -1.093036999 | -0.77502707 | -0.843258992 |
| Neto2 | 0.862076004 | 0.848035694 | 0.96517575 | -1.243958744 | -0.584683606 | -0.846645098 |
| Ireb2 | 0.871113334 | 1.039304932 | 0.758443172 | -1.186015222 | -0.541770641 | -0.941075575 |
| Cask | 0.875967143 | 0.433655986 | 1.317911258 | -0.939396577 | -0.830589217 | -0.857548593 |
| Ebf4 | 0.879469721 | 0.911122756 | 0.51799261 | 0.092756203 | -1.544362572 | -0.856978718 |

| | | | | | | |
|---|---|---|---|---|---|---|
| Slc17a8 | 0.880149322 | 0.846248919 | 0.741880375 | -1.254706361 | -1.176060484 | -0.037511771 |
| Adamts10 | 0.887211851 | 1.313306991 | 0.428593212 | -0.883570275 | -0.883570275 | -0.861971505 |
| Neto1 | 0.891004371 | 0.972348491 | 0.86966318 | -1.007728714 | -0.87861605 | -0.846671278 |
| Apc | 0.891619077 | 0.930132395 | 0.900839993 | -1.102351837 | -0.827275805 | -0.792963824 |
| Dnmt3a | 0.907457536 | 0.873595804 | 0.940957097 | -1.07469105 | -0.733772274 | -0.913547113 |
| Fam193b | 0.909798214 | 1.10414324 | 0.694464775 | -1.025216706 | -0.801464908 | -0.881724616 |
| Trpc4 | 0.911723308 | 1.158515667 | 0.527698104 | -0.710016876 | -1.305824345 | -0.582095857 |
| Pik3r3 | 0.91558271 | 0.714489722 | 1.05632091 | -0.999007849 | -1.082174707 | -0.605210786 |
| Nrep | 0.919984198 | 0.496832771 | 1.215448086 | -0.957128899 | -0.60268947 | -1.072446686 |
| Neurod1 | 0.921415045 | 1.231408647 | 0.411276759 | -1.280165249 | -0.676685191 | -0.607250011 |
| Mir124a-1hg | 0.922196438 | 1.138771651 | 0.640613969 | -0.961275377 | -0.825529482 | -0.914777199 |
| Slc7a6 | 0.92236208 | 1.02340801 | 0.604265117 | -1.40828115 | -0.796366438 | -0.345387619 |
| Gpcpd1 | 0.922739367 | 0.839628606 | 0.92546603 | -1.015465912 | -0.555550869 | -1.116817221 |
| Mirg | 0.924193114 | 1.280609815 | 0.351513605 | -0.922359697 | -0.50337924 | -1.130577598 |
| Tmem169 | 0.944441035 | 0.881787199 | 0.902807675 | -1.05625365 | -0.854401594 | -0.818380665 |
| Tet3 | 0.945996868 | 0.815491614 | 0.883692297 | -0.908188365 | -0.464903685 | -1.272088729 |
| Zfc3h1 | 0.946403213 | 1.083721929 | 0.633082483 | -1.158853596 | -0.917392219 | -0.58696181 |
| Mdm4 | 0.947050926 | 1.013843506 | 0.698578156 | -1.251990966 | -0.582867295 | -0.824614326 |
| Gm5577 | 0.94771355 | 1.021944443 | 0.668581071 | -1.243434956 | -0.911409287 | -0.483394822 |
| Palmd | 0.950782588 | 0.919751044 | 0.490527746 | -1.069319615 | 0.072359648 | -1.364101411 |
| Pkia | 0.952009995 | 0.660858009 | 1.029873899 | -1.303815957 | -0.671198231 | -0.667727715 |
| Ccdc141 | 0.953225057 | 0.700880433 | 0.991941638 | -1.296995146 | -0.784100995 | -0.564950987 |
| D630023F18Rik | 0.956908379 | 0.565791181 | 1.091050285 | -1.236773075 | -0.469198828 | -0.907777941 |
| Rps6kb2 | 0.957767145 | 1.1237646 | 0.562649667 | -0.689913635 | -0.734545272 | -1.219722505 |
| Inha | 0.958149986 | 1.070352608 | 0.654005734 | -0.623481665 | -1.024737789 | -1.034288875 |
| Nav1 | 0.959214019 | 1.170703559 | 0.55181052 | -0.949436229 | -0.8272867 | -0.905005169 |
| Epha3 | 0.959262788 | 0.683666812 | 1.059644063 | -0.739778317 | -1.070880379 | -0.891914968 |
| St6gal2 | 0.960912061 | 0.873261682 | 0.878633328 | -1.147021357 | -0.766083944 | -0.79970177 |
| Fnbp1l | 0.964143408 | 0.9246585 | 0.811826934 | -1.177926345 | -0.692252573 | -0.830449924 |
| Gabra2 | 0.964199687 | 0.91510205 | 0.852314564 | -1.017169067 | -0.824287968 | -0.890159267 |
| N4bp2 | 0.968620021 | 1.173772201 | 0.457227333 | -1.147940593 | -0.49832714 | -0.953351822 |
| Csmd2 | 0.971322474 | 1.026113934 | 0.684229577 | -0.680670167 | -0.813216404 | -1.187779414 |
| Cdc7 | 0.972139964 | 0.816181496 | 0.752327744 | -1.509613192 | -0.594083276 | -0.436952736 |
| Ndst3 | 0.978194573 | 1.06304691 | 0.647400639 | -0.792354505 | -0.763536074 | -1.132751543 |
| Hapln1 | 0.983204341 | 0.68348671 | 1.034998379 | -0.848570027 | -0.756505821 | -1.096613582 |
| 2610008E11Rik | 0.987719035 | 0.843639506 | 0.859081888 | -1.084942723 | -0.569564755 | -1.035932952 |
| Tfrc | 0.988005398 | 0.917327609 | 0.770582671 | -1.061326567 | -0.525607978 | -1.088981134 |
| Agbl3 | 0.989176847 | 0.43528488 | 1.199317115 | -1.128608294 | -0.714641884 | -0.780528663 |
| Brwd3 | 0.989257815 | 0.932284021 | 0.737494543 | -1.046948322 | -1.132236103 | -0.479851954 |
| Zcchc7 | 0.993192769 | 1.155714664 | 0.52309823 | -0.99898961 | -0.778248336 | -0.894767717 |
| Spata33 | 0.993615624 | 0.396294763 | 1.161376387 | -0.543421393 | -0.691031079 | -1.316834302 |
| Cldn1 | 0.995964218 | 0.76930979 | 0.855864177 | -0.37662351 | -1.229369204 | -1.01514547 |
| Cd2ap | 0.998303917 | 0.89900757 | 0.819067885 | -1.030685333 | -0.699306044 | -0.986387996 |
| Ankrd10 | 0.99884828 | 1.066764122 | 0.627268806 | -1.080042199 | -0.763209078 | -0.84962993 |
| Snhg11 | 1.004566264 | 1.043200959 | 0.645445241 | -1.077863371 | -0.708899478 | -0.906449615 |
| C230037L18Rik | 1.004740726 | 1.004740726 | 0.695359224 | -1.099513648 | -0.802663514 | -0.802663514 |
| Miat | 1.005801305 | 1.057291405 | 0.622584135 | -1.046184197 | -0.670719195 | -0.968773453 |
| Rgs11 | 1.007521456 | 1.082969278 | 0.536509008 | -0.50566225 | -1.156859848 | -0.964477643 |
| Unc13b | 1.009085432 | 1.009085432 | 0.702316642 | -0.948558475 | -0.90350154 | -0.86842749 |
| Pgap1 | 1.009394343 | 0.887804922 | 0.757621295 | -1.287786157 | -0.797864786 | -0.569169616 |
| Pcp2 | 1.011991747 | 0.90746176 | 0.566542738 | -0.140135076 | -1.346488977 | -0.999372192 |
| Mmp16 | 1.013485692 | 1.164847743 | 0.451285565 | -0.908818273 | -0.630411665 | -1.090389062 |
| Acvr2a | 1.013775285 | 0.913236164 | 0.774563242 | -1.165662389 | -0.77176779 | -0.764144513 |
| Amy1 | 1.017066589 | 1.095239054 | 0.580136856 | -0.913828454 | -0.809771515 | -0.968842529 |
| Ints6l | 1.019905812 | 1.060100094 | 0.556255538 | -1.032410049 | -0.506528047 | -1.097323348 |
| Sestd1 | 1.02040905 | 0.844070183 | 0.863151265 | -1.02685373 | -0.866688672 | -0.834088097 |
| Cfap65 | 1.021190736 | 1.365868186 | -0.244858045 | -0.548186001 | -0.317075924 | -1.276938951 |
| Akap8l | 1.022492526 | 1.115298742 | 0.516533456 | -0.814165113 | -0.710494012 | -1.1296656 |
| Cfap69 | 1.022762798 | 0.957492015 | 0.730152407 | -1.067834476 | -0.735409474 | -0.90716327 |
| Pip5kl1 | 1.025941463 | 0.918395943 | 0.657454667 | -1.161031049 | -0.342321582 | -1.098439441 |
| Thpo | 1.029684531 | 1.198786276 | 0.387358784 | -0.892420807 | -0.673116763 | -1.050292021 |
| Lrrc45 | 1.031116222 | 1.122427409 | 0.522918831 | -0.981178459 | -0.891270278 | -0.804013724 |
| Lcorl | 1.036256559 | 0.875572909 | 0.707590178 | -1.211895737 | -0.387844631 | -1.019679277 |
| Rad52 | 1.037217182 | 0.214247476 | 1.12532022 | -0.267837676 | -0.656750854 | -1.452196348 |
| Eml5 | 1.039923265 | 1.104196454 | 0.517001462 | -1.034435306 | -0.673862812 | -0.952823063 |
| Eml6 | 1.042967154 | 1.08226824 | 0.542379536 | -1.092299001 | -0.70761996 | -0.86769597 |
| Vcan | 1.043261334 | 0.75062793 | 0.635594224 | -0.142280256 | -0.750988255 | -1.536214976 |
| Gm3764 | 1.044658327 | 1.118155752 | 0.508454436 | -0.847462156 | -0.815560415 | -1.008245944 |
| Meg3 | 1.047467541 | 1.04389871 | 0.54881128 | -1.178161332 | -0.565050869 | -0.89696533 |
| Rmst | 1.054070621 | 0.422500856 | 0.68083193 | -0.231392442 | -0.144899605 | -1.78111136 |
| Cemip2 | 1.054431011 | 0.738303544 | 0.762227017 | -0.75365676 | -0.365981914 | -1.435322898 |
| Nsd2 | 1.057165387 | 0.855741424 | 0.803742164 | -1.068178236 | -0.870999988 | -0.777470752 |
| Spaca6 | 1.064869786 | 1.151423262 | 0.417247583 | -1.054408846 | -0.728024623 | -0.851107162 |

| | | | | | | |
|---|---|---|---|---|---|---|
| Gabrg3 | 1.067274956 | 0.720541253 | 0.713791431 | -0.700159276 | -1.500480444 | -0.30096792 |
| Nectin3 | 1.067719107 | 0.733511946 | 0.841017573 | -1.19510075 | -0.459716698 | -0.987431178 |
| Firre | 1.069463474 | 1.076498765 | 0.507793208 | -0.834855083 | -0.699257332 | -1.119643032 |
| Slc44a5 | 1.069622114 | 1.011111342 | 0.578456462 | -0.775168053 | -1.19030089 | -0.693720974 |
| Kif22 | 1.07075538 | 0.921496283 | 0.652324313 | -0.589206437 | -0.783385454 | -1.271984085 |
| Met | 1.070940301 | 1.172114478 | -0.052933747 | -1.457250218 | -0.571745568 | -0.161125247 |
| Pcdh15 | 1.077323359 | 0.987702924 | 0.630827752 | -1.059346612 | -0.770399036 | -0.866108388 |
| Atp11c | 1.080903289 | 1.130295512 | 0.441225828 | -0.981249037 | -0.843843161 | -0.827332431 |
| Gm39469 | 1.08249087 | 1.130473755 | 0.424814486 | -1.034940871 | -0.88546506 | -0.71737318 |
| Agrn | 1.08361596 | 0.784125786 | 0.718152413 | -0.331709362 | -0.977970526 | -1.276214271 |
| Leng8 | 1.084590154 | 1.138127935 | 0.371037683 | -0.583016571 | -0.884506773 | -1.126232429 |
| Wsb1 | 1.090929196 | 0.955156485 | 0.647358435 | -1.037358757 | -0.703435123 | -0.952650236 |
| Slc4a7 | 1.09143754 | 0.965918272 | 0.634651844 | -0.719357583 | -0.916205375 | -1.056444699 |
| B230307C23Rik | 1.091779558 | 0.573006499 | 1.009356443 | -1.107337737 | -0.723214869 | -0.843589894 |
| Col23a1 | 1.093760311 | 0.965187336 | 0.630658947 | -0.853272189 | -0.744622351 | -1.091712053 |
| Sfi1 | 1.095299496 | 1.127901525 | 0.132936655 | -0.215094874 | -0.777104405 | -1.363938396 |
| Tmem181a | 1.097330016 | 1.091767561 | 0.383612763 | -0.592578057 | -0.720677026 | -1.259455257 |
| Zkscan2 | 1.100429114 | 1.029033082 | 0.553158674 | -0.963815304 | -0.954471004 | -0.764334563 |
| Eogt | 1.10671927 | 0.753879426 | 0.670241887 | -1.10191077 | -0.203345287 | -1.225584526 |
| Dot1l | 1.107403482 | 1.15463295 | 0.278877669 | -0.49234212 | -0.919445205 | -1.129126776 |
| Slc16a7 | 1.114359728 | 0.906836846 | 0.63244124 | -1.227841466 | -0.636004962 | -0.789791386 |
| Cntnap5a | 1.116103629 | 0.843610215 | 0.714204148 | -1.176310065 | -0.865908206 | -0.631699721 |
| Dynlt1a | 1.121440729 | 0.79631767 | 0.555077089 | -0.86058523 | -0.189742173 | -1.422508086 |
| Mdm1 | 1.122381188 | 0.900865813 | 0.60161213 | -0.472217416 | -0.960472609 | -1.192169107 |
| Atp11b | 1.122821205 | 0.917587712 | 0.605845817 | -1.227502354 | -0.801747694 | -0.617004687 |
| Thbs3 | 1.123195248 | 1.199834984 | 0.229933828 | -0.791399101 | -0.696268535 | -1.065296425 |
| A230057D06Rik | 1.125023865 | 0.748706496 | 0.829833706 | -0.973862276 | -1.007669017 | -0.722032774 |
| Ryr3 | 1.128043372 | 1.004418577 | 0.543317 | -1.026963078 | -0.893393666 | -0.755422205 |
| Celsr3 | 1.128991595 | 0.97793664 | 0.54345414 | -0.601170487 | -1.104088233 | -0.945123656 |
| C230057M02Rik | 1.130858268 | 1.076153255 | 0.414084211 | -1.127449434 | -0.69704286 | -0.79660344 |
| 4930570G19Rik | 1.130951439 | 0.744036157 | 0.752623296 | -1.222198796 | -0.947201181 | -0.458210915 |
| Dcx | 1.132462608 | 0.877690598 | 0.672368742 | -0.897503474 | -0.672907358 | -1.112111115 |
| Ccdc39 | 1.133493872 | 0.979682173 | 0.555139876 | -0.716273879 | -0.85570505 | -1.096336991 |
| Sfswap | 1.133986819 | 0.839268022 | 0.645931283 | -0.678085887 | -0.617888575 | -1.323211661 |
| Rimklb | 1.144964432 | 0.726297141 | 0.745574651 | -0.939125399 | -0.44114067 | -1.236570155 |
| Cdh9 | 1.164028631 | 0.761286883 | 0.686393795 | -1.26490024 | -0.4643313 | -0.882477769 |
| Kcnk10 | 1.181671804 | 0.600537179 | 0.474963472 | 0.055646118 | -0.744247193 | -1.568571379 |
| Tet1 | 1.186572064 | 1.033239326 | 0.396124958 | -1.107501266 | -0.794843872 | -0.71359121 |
| Gm43305 | 1.188913657 | -0.179285515 | 1.289565816 | -0.554313235 | -1.048215066 | -0.696665656 |
| Igsf9b | 1.199246111 | 0.944254318 | 0.498479337 | -1.00461997 | -0.628974776 | -1.00838502 |
| Slc26a8 | 1.201547246 | 0.192894973 | 0.950076147 | -1.143025048 | -1.143025048 | -0.05846827 |
| Guf1 | 1.206543808 | 0.89593567 | 0.556763921 | -1.094749842 | -0.714173314 | -0.850320242 |
| Rfx3 | 1.210977178 | 0.816201722 | 0.613063396 | -1.130025576 | -0.5625373 | -0.947679419 |
| Ago4 | 1.214546636 | 0.955162816 | 0.41752914 | -1.20695137 | -0.569973181 | -0.810314041 |
| Fbxl12os | 1.217371768 | 0.972041047 | 0.426425485 | -0.605353259 | -0.961505429 | -1.048979613 |
| Ezh2 | 1.218605187 | 0.000230275 | 1.160869824 | -0.442206458 | -0.750936202 | -1.186562626 |
| Abcc10 | 1.2284587 | 1.090955123 | -0.152728563 | -0.536007845 | -0.240655619 | -1.390021795 |
| Col5a1 | 1.229579497 | 0.924738004 | 0.480507393 | -0.781083051 | -0.734729843 | -1.119012 |
| Ccdc57 | 1.237639056 | 0.955527328 | -0.115522826 | 0.154572804 | -1.231398856 | -1.000817504 |
| Stard9 | 1.247269501 | 0.921617443 | 0.426598931 | -0.842007258 | -0.588092831 | -1.165385786 |
| Slc30a10 | 1.253324284 | 0.176153697 | 0.961836963 | -1.315886815 | -0.252068454 | -0.823359676 |
| Parp8 | 1.253785321 | 0.914411493 | 0.476200922 | -0.775821272 | -0.840963464 | -1.027613 |
| Tmem131l | 1.277037386 | 1.123598314 | -0.046381582 | -0.635190266 | -1.193817846 | -0.525246007 |
| Gm48957 | 1.292566362 | 0.139132061 | 1.071319279 | -0.625406326 | -0.874214057 | -1.003397319 |
| Slc38a6 | 1.307107696 | 0.86175389 | 0.418740484 | -1.133582923 | -0.852329218 | -0.601689929 |
| Rgs9 | 1.324379973 | 0.762561471 | 0.367325994 | -0.512033242 | -0.534496881 | -1.407737315 |
| Adamts17 | 1.356658453 | 0.959602655 | 0.239489937 | -0.896705889 | -0.851751956 | -0.807293199 |
| Nphp3 | 1.360955933 | 0.931808974 | 0.203501216 | -1.135699482 | -0.573666872 | -0.786899769 |
| 9630013A20Rik | 1.389479456 | 0.333048612 | 0.837794075 | -0.96256098 | -0.96256098 | -0.635200184 |
| Gm35339 | 1.393329579 | 0.931582154 | 0.073207776 | -0.618322255 | -0.558789541 | -1.221007712 |
| Bach2 | 1.397168927 | 0.51157816 | 0.689582004 | -0.747182034 | -0.785877906 | -1.065269151 |
| Ap1g2 | 1.398562362 | 0.652814658 | 0.276904208 | -0.593100193 | -0.282788863 | -1.452392172 |
| Tubd1 | 1.40399422 | 1.082793407 | -0.347255017 | -0.828844344 | -0.96343325 | -0.347255017 |
| Ryr1 | 1.458423073 | 0.809704289 | -0.259703661 | -0.391231382 | -0.221562252 | -1.395630068 |
| Catsper2 | 1.589793768 | 0.319881192 | 0.351349742 | -0.698626299 | -0.262468987 | -1.299929415 |

**Supplementary Table 3** DEGs in sublayers of MEC between old and young.

DEGs in sublayers of MEC between old and young

| n = 31 | MEC_s | | MEC_im | | MEC_d | |
|---|---|---|---|---|---|---|
| gene | log2FoldChange | padj | log2FoldChange | padj | log2FoldChange | padj |
| Cd52 | 2.731383891 | 2.15E-09 | 3.175643793 | 3.85E-22 | 2.432382238 | 0.006086541 |
| C4b | 2.45960638 | 2.67E-13 | 3.676540308 | 2.98E-23 | 3.280503619 | 6.23E-06 |
| Lyz2 | 1.66157364 | 1.12E-05 | 2.529575239 | 2.95E-12 | 2.2064467 | 0.021232819 |
| Trf | 1.369620327 | 9.15E-12 | 1.823010681 | 5.02E-06 | 2.043719828 | 0.019098929 |
| Fos | 1.094383162 | 0.004281791 | 2.394190584 | 5.67E-17 | 1.745626246 | 0.034329131 |
| Zic1 | -0.634189128 | 0.013238105 | -1.084557799 | 3.01E-06 | -1.383497043 | 0.013475088 |
| Flna | -0.693023175 | 0.031973741 | -0.933937068 | 4.91E-06 | -1.694855946 | 0.019097425 |
| Hacd1 | -0.750103895 | 0.037808361 | -0.878834278 | 0.00010132 | -1.539139128 | 0.029364672 |
| Gm17750 | -0.870639574 | 0.014047239 | -1.047853308 | 4.89E-05 | -1.540393097 | 0.018127776 |
| Mfsd2a | -0.885948402 | 0.002592636 | -1.367244324 | 4.47E-14 | -2.027856832 | 2.08E-05 |
| Sema4g | -0.92463645 | 0.001790406 | -0.93451967 | 1.82E-07 | -1.57043989 | 0.036381809 |
| Gdf10 | -0.970677031 | 0.009049745 | -1.604062064 | 3.19E-09 | -2.072948651 | 0.004482748 |
| Tbc1d16 | -0.984318753 | 0.008700123 | -1.002121884 | 2.76E-06 | -1.407885331 | 0.021232819 |
| Gnb4 | -1.056136062 | 0.010429466 | -1.318809379 | 1.42E-06 | -2.137299303 | 0.002268553 |
| Cxadr | -1.241527232 | 0.014459121 | -1.831300152 | 3.42E-08 | -1.922474663 | 0.038734655 |
| Neurod1 | -1.254928371 | 1.62E-05 | -0.539484073 | 0.002686344 | -1.254114803 | 0.034329131 |
| Pcp2 | -1.334622049 | 0.020124875 | -1.210590503 | 0.003607781 | -1.548386393 | 0.029304431 |
| Col9a3 | -1.341000331 | 0.001945581 | -1.701526461 | 4.24E-06 | -1.888094554 | 0.039049874 |
| Mpzl1 | -1.345692305 | 5.14E-05 | -1.420294397 | 6.60E-08 | -1.867160833 | 0.00403097 |
| Gabra2 | -1.360125987 | 1.14E-11 | -1.390307772 | 1.67E-25 | -1.847982038 | 0.000801576 |
| Dpysl3 | -1.387678134 | 0.000259774 | -1.291159583 | 5.78E-05 | -2.189254611 | 0.005245142 |
| Tmem229b | -1.518754874 | 4.16E-05 | -1.334984023 | 1.21E-08 | -1.889693589 | 0.004189656 |
| Cemip | -1.55833796 | 3.53E-07 | -1.367462742 | 5.97E-05 | -2.171904986 | 0.008221945 |
| Gpr17 | -1.679804857 | 1.71E-05 | -1.958423519 | 1.06E-06 | -2.672133977 | 3.02E-05 |
| Apcdd1 | -1.748367492 | 2.84E-05 | -2.068964867 | 1.89E-09 | -2.026024456 | 0.005245142 |
| Nrep | -1.752395272 | 2.33E-12 | -1.506741459 | 3.94E-09 | -2.380491426 | 0.015549952 |
| Wdfy1 | -1.863245451 | 9.52E-09 | -1.741695084 | 5.65E-32 | -1.930283884 | 0.000906931 |
| Dnmt3a | -1.986914766 | 2.85E-08 | -1.765851679 | 4.80E-07 | -2.136292032 | 0.013875166 |
| Col4a2 | -2.036653838 | 0.000978651 | -1.640773695 | 1.26E-08 | -1.84723535 | 0.000906931 |
| Col4a1 | -2.138009505 | 0.001232516 | -1.945431932 | 1.55E-06 | -2.28703039 | 0.001088508 |
| Eln | -4.102590067 | 3.30E-06 | -3.436207801 | 4.03E-07 | -2.624712434 | 0.000972744 |
| n = 166 | MEC_s | | MEC_im | | MEC_d | |
| gene | log2FoldChange | padj | log2FoldChange | padj | log2FoldChange | padj |
| H2-Ab1 | 3.219091214 | 3.50E-05 | 2.502133381 | 7.23E-05 | 1.470701677 | 0.440677298 |
| Bche | 1.970596597 | 0.000198954 | 0.216699668 | 0.670643976 | 0.421072299 | 0.993542839 |
| Hba-a1 | 1.899092841 | 0.000869657 | 1.024218017 | 0.076295005 | 0.234978646 | 0.993542839 |
| Hba-a2 | 1.688087202 | 0.002515293 | 0.93790397 | 0.064136989 | 0.13518377 | 0.993542839 |
| Hbb-bt | 1.453807685 | 0.025322435 | 0.619239474 | 0.293474399 | -0.628558483 | 0.993542839 |
| Ptgis | 1.393572927 | 0.01800354 | -0.178060119 | 0.819783088 | -0.302826381 | 0.993542839 |
| Bst2 | 1.379350246 | 0.000764241 | 0.287628786 | 0.337470788 | -0.241114781 | 0.993542839 |
| Gm42418 | 1.365824408 | 0.010848977 | 0.450684832 | 0.049392566 | -0.108013626 | 0.993542839 |
| Stac2 | 1.33629146 | 2.57E-05 | 0.74509308 | 0.08877026 | 0.316557592 | 0.993542839 |
| Hbb-bs | 1.317534738 | 0.014522964 | 0.465818728 | 0.396458226 | -0.315831722 | 0.993542839 |
| Cdr1os | 1.183239215 | 0.012114379 | 0.227377309 | 0.141798803 | -0.206763925 | 0.993542839 |
| Malat1 | 0.994946694 | 0.040481708 | 0.174640499 | 0.476167284 | -0.191804994 | 0.993542839 |
| Scrt1 | 0.945966784 | 0.000270982 | 0.474566607 | 0.011837481 | -0.043963066 | 0.993542839 |
| Pura | 0.931623445 | 0.005242593 | 0.422288244 | 0.007376539 | 0.000960254 | 0.998243094 |
| Mcur1 | 0.927136058 | 0.038408309 | 0.350639873 | 0.030578665 | 0.111009269 | 0.993542839 |
| Siglech | 0.873307684 | 0.042177996 | 0.169559139 | 0.571822555 | 0.643293675 | 0.905979556 |
| Rfx4 | 0.863068632 | 0.040481708 | -0.265420727 | 0.377878912 | -0.171744731 | 0.993542839 |
| Pcnt | 0.854373675 | 0.018191387 | 0.481423981 | 0.005821068 | 0.079658327 | 0.993542839 |
| Prxl2b | 0.838691986 | 8.73E-07 | 0.359317323 | 0.018653759 | 0.060090014 | 0.993542839 |
| Cep170b | 0.836466447 | 5.95E-05 | 0.418858506 | 0.000463295 | 0.061611933 | 0.993542839 |
| Cdc73 | 0.828544525 | 0.011207416 | 0.289972771 | 0.029718687 | 0.042763317 | 0.993542839 |
| Igfbp5 | 0.799770791 | 0.000378226 | 0.447691153 | 0.170493357 | -0.16646264 | 0.993542839 |
| Igfbp6 | 0.784520777 | 0.048706034 | 1.196144118 | 0.052965631 | 0.454976099 | 0.993542839 |
| Git1 | 0.7745307 | 2.69E-05 | 0.309751035 | 0.022321053 | -0.138541562 | 0.993542839 |
| Med13l | 0.753817267 | 0.020770281 | 0.226529614 | 0.204364059 | 0.109235965 | 0.993542839 |
| Igsf11 | 0.748416712 | 0.031973741 | 0.303404541 | 0.121374286 | 0.286791305 | 0.993542839 |
| Pacrg | 0.74809515 | 0.042697053 | 0.259521965 | 0.060691179 | 0.069369761 | 0.993542839 |
| Zbtb38 | 0.745988092 | 0.015882863 | 0.459558977 | 0.000176064 | 0.139842746 | 0.993542839 |
| Ndrg2 | 0.745156475 | 0.000182159 | 0.395079518 | 0.007093719 | 0.18474536 | 0.993542839 |
| Brsk1 | 0.736782771 | 0.004398243 | 0.289302796 | 0.041490232 | -0.105770315 | 0.993542839 |
| Marf1 | 0.733292563 | 0.043049037 | 0.495089097 | 0.002336089 | 0.070028938 | 0.993542839 |
| Pbx1 | 0.731017594 | 0.008175341 | 0.338828184 | 0.036802357 | 0.007582322 | 0.993542839 |
| Rragd | 0.699615358 | 0.024690038 | 0.414426831 | 0.010567654 | 0.09783182 | 0.993542839 |
| Ctso | 0.697818307 | 0.016432086 | 0.327504982 | 0.069483699 | -0.332553831 | 0.993542839 |
| Scara3 | 0.693027796 | 0.021238579 | 0.234583337 | 0.317602223 | -0.284891566 | 0.993542839 |

| | | | | | | |
|---|---|---|---|---|---|---|
| Nebl | 0.691381521 | 0.040481708 | -0.003790674 | 0.984839549 | 0.093395445 | 0.993542839 |
| B930095G15Rik | 0.688289961 | 0.007378557 | -0.025006027 | 0.888759317 | -0.579682451 | 0.966659749 |
| mt-Co1 | 0.684768114 | 0.003170809 | 0.360613626 | 0.082216684 | 0.127273385 | 0.993542839 |
| Tshz2 | 0.676300275 | 0.030188867 | 0.176386838 | 0.747600454 | -0.245119841 | 0.993542839 |
| Ly6a | 0.660492588 | 0.025055529 | 0.285464315 | 0.235247927 | 0.002855945 | 0.995344851 |
| Itgb8 | 0.656454006 | 0.039566676 | 0.189937613 | 0.323509958 | -0.018848219 | 0.993542839 |
| Rida | 0.650074507 | 0.016321294 | 0.460834412 | 0.003423445 | 0.32448844 | 0.993542839 |
| Camk2n1 | 0.648709099 | 0.002288596 | 0.180446967 | 0.155169209 | -0.191261685 | 0.993542839 |
| Scd1 | 0.647796489 | 0.000539003 | 0.465824511 | 0.000228137 | 0.069941279 | 0.993542839 |
| Bcl7b | 0.638291144 | 0.011387878 | 0.497890527 | 0.000161505 | -0.097010437 | 0.993542839 |
| Zfp106 | 0.636485898 | 0.011701268 | 0.4856607 | 1.16E-09 | 0.018223757 | 0.993542839 |
| Dlgap1 | 0.62996916 | 0.000282345 | 0.452567639 | 1.08E-06 | 0.152447489 | 0.993542839 |
| Tppp3 | 0.62937174 | 0.028107103 | 0.414192009 | 0.038572002 | -0.185734803 | 0.993542839 |
| Gstm1 | 0.623260886 | 0.002197042 | 0.490827092 | 0.019117482 | 0.295566668 | 0.993542839 |
| Slain1 | 0.623044302 | 0.043987673 | 0.341876313 | 0.001344118 | 0.003302808 | 0.994994711 |
| Rabggta | 0.619231137 | 0.026951754 | 0.286026631 | 0.011951871 | -0.27784321 | 0.993542839 |
| Panx2 | 0.617091762 | 0.016432086 | 0.368364375 | 0.023094973 | -0.342290345 | 0.993542839 |
| Serpinb6a | 0.616713036 | 0.040696227 | 0.025960607 | 0.902617066 | -0.528721015 | 0.993542839 |
| Camk2a | 0.612626363 | 9.52E-05 | 0.121380265 | 0.256975378 | -0.261593783 | 0.993542839 |
| mt-Nd4 | 0.607356134 | 0.01052649 | 0.445428681 | 0.023753478 | 0.336583501 | 0.993542839 |
| Fxyd1 | 0.606781561 | 0.017565824 | 0.358197745 | 0.004151468 | 0.120552935 | 0.993542839 |
| Map1a | 0.603699473 | 7.99E-07 | 0.393544348 | 0.000333832 | -0.076575047 | 0.993542839 |
| Gclm | 0.601236072 | 0.006810074 | 0.168680709 | 0.164452066 | -0.057926683 | 0.993542839 |
| Dnajb2 | 0.594351281 | 0.0145729 | 0.411787877 | 0.000487401 | -0.019996444 | 0.993542839 |
| mt-Atp6 | 0.586238538 | 0.015738065 | 0.311468744 | 0.089162048 | 0.14344519 | 0.993542839 |
| Gprc5b | 0.585880159 | 0.012594822 | 0.398615213 | 0.000325224 | 0.233070303 | 0.993542839 |
| Syf2 | 0.582832278 | 0.013982256 | 0.313879902 | 0.007900533 | -0.06471033 | 0.993542839 |
| Naaa | 0.580817073 | 0.039566676 | 0.368183285 | 0.007455111 | 0.188337047 | 0.993542839 |
| Ccdc92 | 0.580736317 | 0.031372954 | 0.488378614 | 0.000255074 | 0.27951658 | 0.993542839 |
| Spag7 | 0.576952393 | 0.011700498 | 0.385772011 | 4.20E-05 | 0.142222542 | 0.993542839 |
| Cops3 | 0.576241867 | 0.002417759 | 0.383219026 | 3.75E-05 | 0.128685932 | 0.993542839 |
| Rps25 | 0.571677677 | 0.002996042 | 0.435783161 | 0.000476647 | 0.149815338 | 0.993542839 |
| Nacc2 | 0.5703027 | 0.006507179 | 0.351700716 | 0.020966786 | -0.009522679 | 0.993542839 |
| Pdp1 | 0.563629177 | 0.034490199 | 0.467824161 | 0.000719222 | -0.218495206 | 0.993542839 |
| Hist1h2bc | 0.553794831 | 0.039604334 | 0.176322329 | 0.078354055 | -0.20637868 | 0.993542839 |
| Plpp3 | 0.553779908 | 0.001113056 | 0.208640317 | 0.040239403 | -0.186475135 | 0.993542839 |
| Ccni | 0.550235671 | 0.00547261 | 0.420150243 | 0.00352261 | -0.061044419 | 0.993542839 |
| Rasgef1a | 0.549908041 | 0.037685634 | 0.199367354 | 0.156669394 | 0.005444647 | 0.993542839 |
| Btbd6 | 0.546254132 | 0.032828132 | 0.220868815 | 0.042443458 | -0.266761405 | 0.993542839 |
| Rasgrf1 | 0.5408718 | 0.000208465 | 0.38559064 | 4.22E-11 | 0.124187232 | 0.993542839 |
| Rcan2 | 0.54002108 | 0.01800354 | 0.471732639 | 0.004818444 | -0.036147219 | 0.993542839 |
| Cyfip2 | 0.539552431 | 0.000310141 | 0.430600097 | 1.26E-08 | 0.170339783 | 0.993542839 |
| Tspyl1 | 0.537970628 | 0.003150448 | 0.472306646 | 1.91E-05 | 0.125652662 | 0.993542839 |
| Rfk | 0.534838638 | 0.013064139 | 0.417214453 | 1.81E-06 | -0.164619742 | 0.993542839 |
| Ccng1 | 0.533640422 | 0.01388522 | 0.426616858 | 0.000293544 | 0.022932981 | 0.993542839 |
| Cplx2 | 0.52971768 | 0.002977712 | 0.095773902 | 0.710858592 | -0.508879968 | 0.993542839 |
| Psme3 | 0.529625116 | 0.046659156 | 0.355797515 | 2.14E-05 | 0.072988945 | 0.993542839 |
| Wscd1 | 0.528989329 | 0.020123855 | 0.368290971 | 0.025661321 | -0.19368747 | 0.993542839 |
| Asah1 | 0.52672731 | 0.048111556 | 0.330804166 | 0.008587324 | -0.336558031 | 0.993542839 |
| Ahcyl2 | 0.525119497 | 0.015955876 | 0.351233419 | 0.001002965 | 0.024670307 | 0.993542839 |
| Shisa5 | 0.520501728 | 0.034664928 | 0.46976905 | 0.000655923 | -0.082363745 | 0.993542839 |
| Zfp365 | 0.520289109 | 0.004984571 | 0.194009623 | 0.036173462 | -0.085761215 | 0.993542839 |
| Wnk1 | 0.519312222 | 0.004676626 | 0.252707648 | 0.027114064 | -0.196762351 | 0.993542839 |
| Ddn | 0.517776401 | 0.0006347 | 0.0387707 | 0.82066857 | -0.063208029 | 0.993542839 |
| Ddx25 | 0.516982172 | 0.049352295 | 0.285244645 | 0.00025523 | 0.03251765 | 0.993542839 |
| Gnai2 | 0.516075195 | 0.00095473 | 0.402308415 | 9.87E-07 | 0.113343249 | 0.993542839 |
| Mllt6 | 0.513994281 | 0.037612454 | 0.235117145 | 0.149422359 | -0.207847032 | 0.993542839 |
| Rpl9 | 0.513407337 | 0.00131415 | 0.473580514 | 0.000237305 | 0.279013683 | 0.993542839 |
| Mid1ip1 | 0.510839349 | 0.010346269 | 0.388159385 | 0.000376915 | -0.125709444 | 0.993542839 |
| Prnp | 0.506987765 | 6.79E-05 | 0.389392173 | 3.07E-05 | -0.048522738 | 0.993542839 |
| Arhgef4 | 0.506788545 | 0.013238105 | 0.28242061 | 0.017356445 | -0.161418687 | 0.993542839 |
| Tnk2 | 0.505642281 | 0.035574291 | 0.295479867 | 0.008587324 | -0.076371133 | 0.993542839 |
| Rpl27a | 0.504320975 | 0.009633751 | 0.43070541 | 0.001765871 | 0.209192358 | 0.993542839 |
| Mzt1 | 0.503449879 | 0.013238105 | 0.452500404 | 1.06E-05 | 0.003099246 | 0.995344851 |
| Rps14 | 0.503315675 | 0.019249448 | 0.385396822 | 0.006444321 | 0.105852366 | 0.993542839 |
| Rnf13 | 0.50172418 | 0.015122571 | 0.420011942 | 7.36E-05 | 0.011186987 | 0.993542839 |
| Eid2 | -0.505933922 | 0.024097186 | -0.219811398 | 0.01096974 | -0.697377988 | 0.988555866 |
| Hist3h2a | -0.50691179 | 0.045283476 | -0.341404815 | 0.001964651 | -0.578152612 | 0.993542839 |
| Pgls | -0.50829498 | 0.046107477 | -0.469419154 | 0.001454876 | -0.936716747 | 0.52600127 |
| Lin7b | -0.509443617 | 0.029910039 | -0.359546133 | 2.09E-07 | -0.76137639 | 0.940150584 |
| Rap2a | -0.512829248 | 0.020123855 | -0.002794187 | 0.988628639 | -0.507576439 | 0.993542839 |
| Pgm2l1 | -0.520584523 | 0.002211112 | -0.462029755 | 0.001489207 | -0.826673812 | 0.993542839 |
| Kitl | -0.526969273 | 0.027508437 | -0.069975123 | 0.687223993 | -1.041875731 | 0.164589982 |

| | | | | | | |
|---|---|---|---|---|---|---|
| Reep2 | -0.528939819 | 0.001630677 | -0.475119445 | 1.50E-08 | -0.973032317 | 0.683203063 |
| Nrbp2 | -0.531983177 | 0.013859315 | -0.480331062 | 0.000443609 | -0.980154897 | 0.566630167 |
| Pacs2 | -0.532048621 | 0.011526164 | -0.458197602 | 0.001455991 | -0.756680272 | 0.813604965 |
| Tspan9 | -0.536488731 | 0.029299808 | -0.381482156 | 0.001039526 | -0.873925905 | 0.532875897 |
| Tubb3 | -0.546110621 | 0.002151982 | -0.340216638 | 1.86E-05 | -1.080136144 | 0.617467376 |
| 1110008P14Rik | -0.550239441 | 0.000660334 | -0.47981086 | 3.73E-06 | -0.820138017 | 0.993542839 |
| Ubfd1 | -0.553709375 | 0.032978742 | -0.317581079 | 0.011837481 | -0.793781627 | 0.731358626 |
| Marcks | -0.559695962 | 0.037963165 | -0.49271269 | 0.008129573 | -0.775128117 | 0.993542839 |
| Ptprs | -0.564451497 | 0.000521309 | -0.403990352 | 2.14E-07 | -0.858317989 | 0.88759934 |
| Ednrb | -0.568496254 | 0.047653207 | -0.423916541 | 0.041422264 | -1.103541859 | 0.250156132 |
| Gtpbp2 | -0.57724922 | 0.032280369 | -0.488977125 | 1.36E-06 | -0.79165447 | 0.665576166 |
| Sidt1 | -0.58573794 | 0.033757441 | -0.413029446 | 0.001228274 | -0.367469767 | 0.993542839 |
| Cacna2d1 | -0.587560179 | 0.009836776 | -0.453337128 | 0.002961017 | -0.876797706 | 0.746080124 |
| Chka | -0.591099244 | 0.011784881 | -0.367429866 | 0.005948969 | -0.675632368 | 0.993542839 |
| Abtb1 | -0.604542257 | 0.03077449 | -0.288434568 | 0.027009073 | -0.582806413 | 0.993542839 |
| Mkrn1 | -0.606500771 | 0.000603607 | -0.395944958 | 5.46E-05 | -0.755554745 | 0.90514278 |
| Xpnpep1 | -0.607694199 | 0.038454558 | -0.36241249 | 0.000456035 | -1.003812075 | 0.442019435 |
| Colgalt1 | -0.610223197 | 0.028009676 | -0.376621722 | 0.002327046 | -0.902114156 | 0.604776757 |
| Sec61a2 | -0.61567428 | 0.04882752 | -0.314038609 | 0.034097364 | -0.787798957 | 0.785664508 |
| Spast | -0.617973948 | 0.031627954 | -0.31744652 | 0.00652867 | -0.627317626 | 0.993542839 |
| Smpd3 | -0.623488558 | 0.017481945 | -0.466276053 | 0.000974651 | -0.936406549 | 0.704422537 |
| Slc7a10 | -0.62396608 | 0.04317125 | -0.422729432 | 0.004823061 | -1.083605814 | 0.313693475 |
| Rab40c | -0.630754853 | 0.027335253 | -0.430158722 | 3.79E-06 | -0.923718496 | 0.52600127 |
| Fabp5 | -0.63445849 | 0.002288596 | -0.30651625 | 0.005578661 | -0.936898558 | 0.805167961 |
| Vopp1 | -0.648869667 | 0.013183781 | -0.238377338 | 0.021542793 | -0.515074949 | 0.993542839 |
| Tuba1a | -0.655339239 | 0.001113056 | -0.474043062 | 0.006032107 | -0.875647387 | 0.993542839 |
| Akap7 | -0.66207944 | 0.021036843 | -0.349658894 | 0.02468514 | -0.907734427 | 0.539788661 |
| Spon1 | -0.666707927 | 0.020929707 | -0.490704299 | 0.000573731 | -0.956573469 | 0.669962604 |
| Olfm2 | -0.669960067 | 0.005499842 | -0.368644286 | 0.000144836 | -0.895838803 | 0.715362667 |
| Btbd3 | -0.670523284 | 4.88E-05 | -0.333352644 | 0.006913768 | -0.703268937 | 0.828427143 |
| Lage3 | -0.672646497 | 0.013238105 | -0.452617266 | 0.001818856 | -0.90419554 | 0.423148525 |
| Fbh1 | -0.673609062 | 0.011175337 | -0.2546809 | 0.010353823 | -0.820235502 | 0.672655957 |
| Foxp4 | -0.674598947 | 0.016321294 | -0.310015504 | 0.018867554 | -0.606342631 | 0.993542839 |
| Fkbp1b | -0.676096606 | 0.009941401 | -0.411676664 | 0.00034929 | -1.121479521 | 0.324775052 |
| Clk4 | -0.676639218 | 0.022781159 | -0.463552489 | 0.001494373 | -0.555147604 | 0.993542839 |
| Fam89b | -0.690834725 | 0.006736937 | -0.498437102 | 4.89E-05 | -0.898594089 | 0.49853379 |
| Daam1 | -0.701366277 | 0.032828132 | -0.308173276 | 0.052938365 | -0.73265596 | 0.736117265 |
| Rnft2 | -0.713915632 | 0.007563272 | -0.359604113 | 0.000568942 | -1.025821292 | 0.440677298 |
| Dpp3 | -0.714828921 | 0.013859315 | -0.433022745 | 0.002006638 | -0.930610259 | 0.550971806 |
| Hbs1l | -0.729555607 | 0.004374324 | -0.388734507 | 0.000154276 | -0.864008775 | 0.481550768 |
| Ireb2 | -0.730609009 | 0.013475266 | -0.443941144 | 0.004287978 | -0.934109014 | 0.479890367 |
| Nrip1 | -0.74482911 | 0.047770303 | -0.389539357 | 0.075914899 | -0.64874065 | 0.993542839 |
| Mta1 | -0.74563993 | 0.002607229 | -0.40694942 | 0.000679104 | -0.724459415 | 0.869121594 |
| Mdm4 | -0.764160558 | 0.016422829 | -0.460101334 | 0.002700627 | -0.987905789 | 0.394193934 |
| Oprl1 | -0.767080316 | 0.010902263 | -0.48686344 | 6.22E-06 | -0.712341215 | 0.90514278 |
| Meis3 | -0.798724174 | 0.02571539 | -0.294464412 | 0.021723112 | -0.703043594 | 0.815735258 |
| Camsap3 | -0.812492509 | 0.042735128 | -0.379215875 | 0.000854646 | -0.942693636 | 0.480415331 |
| Serf1 | -0.820322959 | 0.000978651 | -0.358731142 | 0.00029707 | -1.057874536 | 0.35889639 |
| Cacna1c | -0.840594065 | 0.031094085 | -0.416736621 | 0.003060324 | -0.873797464 | 0.582320142 |
| Fcor | -0.844824762 | 0.0097004 | -0.454779767 | 0.01402636 | -0.70866426 | 0.952066025 |
| Tmem91 | -0.886532585 | 0.043228346 | -0.215077314 | 0.600222666 | -1.183600471 | 0.617467376 |
| Lysmd1 | -0.932488402 | 0.007507405 | -0.484863355 | 0.001194772 | -0.895570908 | 0.520872631 |
| Nt5dc2 | -0.934084429 | 0.031077324 | -0.206939148 | 0.375430883 | -0.709581617 | 0.704589402 |
| Rac3 | -0.952388601 | 0.001267509 | -0.390439312 | 0.02191759 | -1.408494227 | 0.127826557 |
| Tfrc | -1.109731542 | 0.000280062 | -0.444673507 | 0.007951399 | -0.999816303 | 0.416905253 |
| Kcng1 | -1.297371382 | 0.000514509 | -0.449747477 | 0.137773255 | -0.488303705 | 0.993542839 |
| Igfbp3 | -1.475862438 | 0.002288596 | -0.621628477 | 0.123362295 | -0.756041018 | 0.846711539 |
| Col3a1 | -2.344078316 | 0.025055529 | -3.10242279 | 0.005511262 | -1.444586628 | 0.18416142 |
| n = 297 | MEC_s | | MEC_im | | MEC_d | |
| gene | log2FoldChange | padj | log2FoldChange | padj | log2FoldChange | padj |
| H2-Aa | 4.183015662 | 9.49E-08 | 3.941995509 | 1.07E-08 | 2.17880955 | 0.127623655 |
| Cd74 | 3.504471542 | 3.86E-08 | 3.091050948 | 1.80E-11 | 2.456044276 | 0.052399113 |
| Ermn | 2.255928432 | 7.40E-06 | 1.800092032 | 0.000310381 | 1.444204476 | 0.215418753 |
| Lrrc17 | 2.216572625 | 0.003386352 | 2.536486291 | 0.00015929 | 2.44111805 | 0.088407895 |
| H2-K1 | 1.969815752 | 9.15E-12 | 1.545712275 | 9.69E-19 | 1.217978233 | 0.199573866 |
| Arc | 1.930220636 | 5.49E-17 | 1.498634103 | 4.33E-19 | 1.385835462 | 0.165855295 |
| Ifi27l2a | 1.913858118 | 7.53E-05 | 1.078628515 | 0.000365714 | -0.079927306 | 0.993542839 |
| Ppp1r14a | 1.724486939 | 1.17E-08 | 1.747462666 | 0.000247268 | 1.725181349 | 0.058825317 |
| Il33 | 1.642746146 | 3.13E-06 | 1.583691891 | 1.71E-11 | 1.355723188 | 0.107707952 |
| Lars2 | 1.632635167 | 0.017391727 | 0.618321042 | 0.010093158 | -0.00423209 | 0.993571755 |
| Ass1 | 1.505475918 | 4.75E-08 | 1.749052689 | 3.99E-27 | 1.070850743 | 0.35570703 |
| Bc1 | 1.461911216 | 0.022056886 | 1.836640962 | 0.000870094 | 1.780435048 | 0.522349226 |

| | | | | | | |
|---|---|---|---|---|---|---|
| mt-Nd6 | 1.432990248 | 0.013617347 | 0.994311491 | 0.039089693 | 0.980539013 | 0.704422537 |
| B2m | 1.397059982 | 2.15E-09 | 1.093327144 | 4.43E-13 | 1.329472227 | 0.204447175 |
| Tatdn1 | 1.38423179 | 0.006273993 | 1.054890988 | 0.001200628 | 0.681409422 | 0.986773652 |
| Gpr37 | 1.375027243 | 0.000295385 | 1.470574802 | 1.26E-19 | 1.11104677 | 0.184983612 |
| Plekhb1 | 1.364650367 | 9.90E-05 | 1.377305504 | 0.000331509 | 1.408079749 | 0.307206388 |
| Hebp1 | 1.245639333 | 0.003232935 | 0.575893733 | 0.005928917 | 0.068231501 | 0.993542839 |
| Efhd1 | 1.22711866 | 0.002903113 | 1.323521372 | 2.71E-07 | 0.868320382 | 0.504435724 |
| Egr4 | 1.219954706 | 0.000978651 | 1.245956807 | 8.32E-05 | 1.588347684 | 0.282584052 |
| Egr1 | 1.180133231 | 5.78E-10 | 1.55512833 | 2.93E-24 | 1.579075322 | 0.200280631 |
| Lgals3bp | 1.178179714 | 0.003287203 | 1.348458319 | 1.45E-10 | 0.699361063 | 0.82469938 |
| Junb | 1.176735136 | 7.71E-06 | 1.519881098 | 2.53E-16 | 1.12396337 | 0.465383086 |
| Bhlhe40 | 1.174363763 | 3.33E-08 | 1.245699209 | 4.28E-25 | 0.731850786 | 0.810895396 |
| Efemp1 | 1.164788755 | 0.002497205 | 0.616553069 | 0.026180148 | 1.339985938 | 0.184983612 |
| Mobp | 1.160538096 | 0.005471935 | 1.187232219 | 0.004207114 | 0.864970202 | 0.940150584 |
| Neat1 | 1.16048573 | 2.53E-05 | 1.805238386 | 4.53E-10 | 1.631937786 | 0.073570005 |
| Nr4a1 | 1.12997523 | 2.26E-08 | 1.151703194 | 1.78E-11 | 0.86940315 | 0.836346457 |
| Sox9 | 1.116091003 | 0.002211112 | 0.711674709 | 0.001620388 | 0.206778296 | 0.993542839 |
| Cbx7 | 1.110176527 | 0.002613871 | 0.886266697 | 0.010163624 | 0.340271912 | 0.993542839 |
| Gsn | 1.102921416 | 0.013374124 | 0.543927544 | 0.015244175 | 0.121667245 | 0.993542839 |
| Ly86 | 1.101854797 | 0.001500551 | 0.571517819 | 0.006619185 | 1.376297417 | 0.178678744 |
| Ankrd35 | 1.096227627 | 0.000970271 | 0.992958786 | 0.004386077 | 0.345534099 | 0.993542839 |
| Mpeg1 | 1.08813627 | 0.020486911 | 0.986544669 | 7.12E-07 | 0.657168034 | 0.993542839 |
| Ppp1r3c | 1.08715391 | 3.57E-06 | 0.65114542 | 8.42E-07 | 0.195068554 | 0.993542839 |
| mt-Atp8 | 1.086568543 | 0.019290134 | 0.734298199 | 0.019422501 | 0.578508023 | 0.993542839 |
| Zxdb | 1.07018105 | 0.002697798 | 0.7057976 | 2.03E-05 | 0.240144106 | 0.993542839 |
| Tspo | 1.070016356 | 0.007327233 | 0.923267735 | 2.78E-05 | 0.417159232 | 0.993542839 |
| Gfap | 1.069288084 | 0.002592636 | 1.70114274 | 2.46E-05 | 1.808845436 | 0.079943479 |
| Rapgef4 | 1.01746078 | 1.48E-07 | 0.739104629 | 2.03E-06 | 0.650184202 | 0.993542839 |
| Tmbim1 | 1.015870621 | 0.000539003 | 0.661955959 | 0.000353542 | 0.473315712 | 0.993542839 |
| mt-Nd2 | 1.009824269 | 0.002110393 | 0.796781073 | 0.001697633 | 0.72806046 | 0.993542839 |
| Smim1 | 1.008351704 | 0.023326806 | 1.356716225 | 7.56E-09 | 0.682754984 | 0.815549704 |
| Ifi27 | 1.006304409 | 1.48E-07 | 1.35511455 | 2.40E-10 | 0.841198706 | 0.993542839 |
| Plp1 | 0.995444859 | 0.022022994 | 1.185349834 | 0.000551358 | 0.815032591 | 0.993542839 |
| Ankrd33b | 0.995341365 | 0.019290134 | 0.7294673 | 0.000876123 | 0.26105186 | 0.993542839 |
| Ctss | 0.987561748 | 3.80E-05 | 0.932939397 | 1.04E-10 | 1.030748592 | 0.591324978 |
| H2-D1 | 0.981187938 | 8.65E-06 | 0.711776623 | 5.49E-07 | 0.923080431 | 0.493186054 |
| Mettl7a1 | 0.97915151 | 0.003047311 | 0.696303738 | 2.07E-06 | 0.593241118 | 0.993542839 |
| Gstt1 | 0.957752496 | 0.003069566 | 1.413424274 | 0.000745731 | 1.22580286 | 0.16857323 |
| Rps27 | 0.954355889 | 7.06E-05 | 0.911266622 | 1.62E-06 | 0.630232976 | 0.993542839 |
| Plin3 | 0.953472795 | 0.022395806 | 0.921789685 | 0.003090397 | 0.486583027 | 0.993542839 |
| Prr18 | 0.951099065 | 0.03415858 | 1.394570973 | 1.92E-10 | 0.466792127 | 0.993542839 |
| Hapln4 | 0.940047894 | 0.044475764 | 1.142176796 | 0.010439794 | 0.523310043 | 0.993542839 |
| Padi2 | 0.931907508 | 0.009199598 | 0.834187459 | 0.000299834 | 0.304987547 | 0.993542839 |
| Prelp | 0.931438096 | 0.005633612 | 1.442356588 | 1.27E-08 | 0.831292602 | 0.914338763 |
| 2900052N01Rik | 0.929014018 | 0.005672069 | 0.951835095 | 0.001168742 | 0.650978352 | 0.993542839 |
| Jade2 | 0.917517075 | 0.019944242 | 0.961230512 | 1.93E-05 | 0.717926279 | 0.993542839 |
| Nap1l2 | 0.907516446 | 6.18E-06 | 0.802952909 | 1.96E-12 | 0.246467919 | 0.993542839 |
| Macrod1 | 0.903691904 | 0.044018678 | 1.173343648 | 5.49E-05 | 0.713799244 | 0.789916564 |
| Zbtb4 | 0.898242207 | 0.00016795 | 0.517199423 | 0.000735252 | 0.08413916 | 0.993542839 |
| Rapgef3 | 0.895800202 | 0.045460276 | 1.218302388 | 5.18E-11 | 0.608549257 | 0.952813482 |
| mt-Nd3 | 0.893819095 | 0.010346269 | 0.599124455 | 0.006619185 | 0.363681742 | 0.993542839 |
| Plaat3 | 0.884554857 | 0.00026065 | 0.95765953 | 1.02E-07 | 0.683719421 | 0.929751641 |
| Dusp1 | 0.884263656 | 0.000281814 | 0.891749104 | 9.02E-10 | 0.557735996 | 0.993542839 |
| Nfe2l2 | 0.883927851 | 0.011701268 | 0.747276 | 0.000859822 | -0.097269663 | 0.993542839 |
| Nol3 | 0.883624028 | 0.013238105 | 0.776205279 | 1.11E-06 | 0.198733047 | 0.993542839 |
| Ier5 | 0.875282574 | 1.00E-05 | 0.734744389 | 5.08E-05 | 0.638395112 | 0.993542839 |
| Vamp1 | 0.872767461 | 0.006295929 | 0.736393159 | 0.011320058 | 0.241640743 | 0.993542839 |
| Anxa5 | 0.872612255 | 0.001091599 | 0.762649868 | 8.76E-15 | 0.55621816 | 0.993542839 |
| mt-Nd5 | 0.869106205 | 0.007740998 | 0.602839181 | 0.003276787 | 0.464131725 | 0.993542839 |
| Aktip | 0.868229643 | 0.000100453 | 0.694263395 | 2.30E-07 | 0.219189107 | 0.993542839 |
| Ndrg1 | 0.864748002 | 0.000114928 | 0.804756293 | 0.000293544 | 0.340948756 | 0.993542839 |
| Ggact | 0.862608584 | 0.015955876 | 0.909872906 | 1.85E-05 | 0.405809568 | 0.993542839 |
| Gas5 | 0.862060545 | 9.81E-05 | 0.726963724 | 1.27E-05 | 0.492127524 | 0.993542839 |
| Apod | 0.861479753 | 0.034842969 | 1.267938186 | 0.000775905 | 1.41789547 | 0.372214949 |
| Wwox | 0.857617215 | 0.001968236 | 0.893601194 | 0.000200958 | 0.440181072 | 0.993542839 |
| Lgals1 | 0.853186298 | 0.003413199 | 0.712018497 | 8.42E-07 | 0.116303124 | 0.993542839 |
| H2-T22 | 0.847298492 | 0.049560296 | 1.194945795 | 1.52E-05 | 1.159647306 | 0.414591828 |
| Klf10 | 0.846749004 | 0.029355938 | 0.595056704 | 0.000754903 | 0.12710456 | 0.993542839 |
| Qdpr | 0.843232852 | 1.02E-05 | 0.911617871 | 0.000252452 | 0.548651318 | 0.993542839 |
| Rps29 | 0.832627487 | 0.008669366 | 0.744697535 | 0.00014021 | 0.533601933 | 0.993542839 |
| Mbp | 0.830581533 | 7.06E-05 | 0.75871359 | 0.011015032 | 0.481254624 | 0.993542839 |
| mt-Nd4l | 0.827460682 | 0.018249455 | 0.701597718 | 0.01623202 | 0.577862073 | 0.993542839 |
| Gpld1 | 0.827101504 | 0.004386863 | 0.83299606 | 1.67E-07 | 0.414963127 | 0.993542839 |

| | | | | | | |
|---|---|---|---|---|---|---|
| Celf4 | 0.821589881 | 9.65E-05 | 0.554974232 | 0.000332596 | 0.168984976 | 0.993542839 |
| Svip | 0.814475804 | 0.000988658 | 0.505260899 | 4.10E-06 | 0.104239923 | 0.993542839 |
| Evi2a | 0.813208429 | 0.044834059 | 1.421831333 | 1.24E-06 | 1.163256189 | 0.154941158 |
| Kif5b | 0.811103946 | 0.000191527 | 0.625000779 | 1.50E-07 | 0.284498492 | 0.993542839 |
| mt-Co2 | 0.809251097 | 0.001790406 | 0.663828008 | 0.002677055 | 0.463618716 | 0.993542839 |
| Stat1 | 0.798566836 | 0.047817235 | 0.626212559 | 0.003219748 | 0.263600328 | 0.993542839 |
| Ctsz | 0.798560935 | 0.0006291 | 0.782829141 | 4.10E-07 | 0.513323055 | 0.993542839 |
| mt-Nd1 | 0.794517304 | 0.000978651 | 0.671805571 | 0.000944219 | 0.416857825 | 0.993542839 |
| Lgi4 | 0.794143738 | 0.013183781 | 0.76947795 | 3.03E-06 | 0.229205508 | 0.993542839 |
| Scn1a | 0.775099374 | 0.038619803 | 0.755477759 | 1.20E-11 | 0.353096199 | 0.993542839 |
| Id4 | 0.773325058 | 0.000349812 | 0.540308128 | 0.000147946 | 0.085129145 | 0.993542839 |
| Crip1 | 0.769624157 | 0.005474021 | 0.853043862 | 0.002069246 | 0.344843049 | 0.993542839 |
| Fgfr2 | 0.768067092 | 0.012799116 | 0.619614426 | 1.36E-05 | 0.515002738 | 0.993542839 |
| 9930021J03Rik | 0.768013945 | 0.030495671 | 0.515248367 | 0.00427581 | 0.184488652 | 0.993542839 |
| Bcap29 | 0.76739864 | 0.033992548 | 0.696070932 | 6.69E-08 | 0.13961295 | 0.993542839 |
| Rnaset2b | 0.730083175 | 0.015050321 | 0.611588559 | 0.000421608 | 0.07252761 | 0.993542839 |
| Eif3e | 0.729154375 | 0.000108918 | 0.621272197 | 3.65E-10 | 0.228061126 | 0.993542839 |
| Cox6a2 | 0.728769718 | 0.001808821 | 0.756468516 | 3.34E-09 | -0.117251356 | 0.993542839 |
| Rhog | 0.724072186 | 0.018866743 | 0.762457307 | 0.000319712 | 0.507210944 | 0.993542839 |
| Umad1 | 0.716815681 | 0.03008296 | 0.684754499 | 8.99E-08 | -0.049965359 | 0.993542839 |
| Zfas1 | 0.706265291 | 0.000109744 | 0.63962686 | 5.75E-15 | 0.632716567 | 0.993542839 |
| Rpl26 | 0.705780171 | 0.001432597 | 0.639567383 | 0.000160575 | 0.403015875 | 0.993542839 |
| Tomm34 | 0.699493849 | 0.002679565 | 0.733903959 | 2.21E-06 | 0.290569374 | 0.993542839 |
| Cryab | 0.698308134 | 0.014890014 | 0.85322354 | 0.022378928 | 0.694533224 | 0.993542839 |
| Tceal1 | 0.696654361 | 0.040909943 | 0.700298222 | 2.47E-08 | 0.418359362 | 0.993542839 |
| Rps28 | 0.694265111 | 0.004963707 | 0.70285587 | 0.000358976 | 0.427333802 | 0.993542839 |
| Rpl35 | 0.690933058 | 0.007834383 | 0.566899492 | 0.001892676 | 0.263581544 | 0.993542839 |
| Hspa2 | 0.688369387 | 0.013930494 | 0.639288156 | 2.57E-07 | 0.217042094 | 0.993542839 |
| Pex5l | 0.687380839 | 0.001984829 | 0.607812556 | 1.86E-12 | 0.140656167 | 0.993542839 |
| Srebf1 | 0.686204733 | 0.013473384 | 0.636467746 | 3.78E-06 | 0.297065712 | 0.993542839 |
| Wls | 0.684162863 | 0.035448999 | 0.681209476 | 1.21E-05 | 0.011126823 | 0.993542839 |
| Rpl36a | 0.682913394 | 7.06E-05 | 0.624848686 | 2.47E-07 | 0.334537626 | 0.993542839 |
| Ier3 | 0.682754132 | 0.048706034 | 0.681705403 | 0.000307845 | 0.415827844 | 0.993542839 |
| Eif2s3x | 0.679612512 | 0.022395806 | 0.69932326 | 4.82E-05 | 0.303876764 | 0.993542839 |
| Lap3 | 0.67785178 | 0.006795452 | 0.78085287 | 2.00E-09 | 0.493752867 | 0.993542839 |
| Prune2 | 0.672957943 | 0.003170809 | 0.953516885 | 8.84E-17 | 0.155119799 | 0.993542839 |
| Rpl37 | 0.671603 | 0.000400981 | 0.635644739 | 3.10E-05 | 0.417800272 | 0.993542839 |
| Stard10 | 0.670504857 | 0.01800354 | 0.711344777 | 0.008848858 | 0.365050056 | 0.993542839 |
| Cox17 | 0.670443821 | 0.001945581 | 0.69629989 | 5.30E-07 | 0.430465427 | 0.993542839 |
| Zrsr1 | 0.668073791 | 0.029672985 | 0.56897828 | 4.89E-05 | 0.29007916 | 0.993542839 |
| Rps23 | 0.666273764 | 0.000521309 | 0.513557647 | 0.000157646 | 0.247324048 | 0.993542839 |
| Dnajb1 | 0.665687791 | 0.029244789 | 0.654416668 | 9.45E-05 | 0.288085712 | 0.993542839 |
| Rps19 | 0.658968924 | 0.00099746 | 0.659100189 | 2.11E-05 | 0.424745313 | 0.993542839 |
| Fth1 | 0.657955878 | 8.29E-10 | 0.581905039 | 0.000102326 | 0.373754837 | 0.993542839 |
| Rpl37a | 0.652200644 | 0.010224679 | 0.579672352 | 0.001090401 | 0.302510123 | 0.993542839 |
| Rps21 | 0.646799759 | 0.001630677 | 0.573500807 | 8.60E-05 | 0.336707244 | 0.993542839 |
| Itih3 | 0.639353589 | 0.044123376 | 0.913034368 | 1.40E-05 | 0.597013812 | 0.988555866 |
| S100b | 0.631843772 | 0.001932683 | 0.53346567 | 0.000345263 | 0.225875532 | 0.993542839 |
| Fcer1g | 0.624552161 | 0.013312561 | 0.502870696 | 0.003443919 | 0.73096885 | 0.839732955 |
| Pttg1 | 0.622918745 | 0.005474021 | 0.566801189 | 0.00252413 | 0.213056351 | 0.993542839 |
| Pnrc1 | 0.622829287 | 0.018599999 | 0.679667707 | 1.92E-06 | 0.227115386 | 0.993542839 |
| Gatm | 0.622495303 | 0.009941401 | 0.682184692 | 5.33E-07 | 0.744763814 | 0.815048805 |
| Bhlhe41 | 0.617161156 | 0.043335529 | 0.69768997 | 1.81E-05 | 0.631117221 | 0.949705241 |
| Gng11 | 0.611273285 | 0.020123855 | 0.757672179 | 3.79E-05 | 0.536960256 | 0.993542839 |
| Rpl17 | 0.609691427 | 0.00095473 | 0.538186909 | 0.000139127 | 0.296788066 | 0.993542839 |
| Slc5a5 | 0.607798641 | 0.049054813 | 1.168450862 | 1.49E-09 | 0.040065884 | 0.993542839 |
| Rnf7 | 0.606091293 | 0.000647189 | 0.630725322 | 1.09E-07 | 0.291887605 | 0.993542839 |
| Dusp14 | 0.604912042 | 0.047770303 | 0.787643522 | 1.34E-07 | 0.24795003 | 0.993542839 |
| Rps6 | 0.598246407 | 2.53E-05 | 0.671672203 | 5.87E-10 | 0.470497009 | 0.993542839 |
| Tyrobp | 0.587367781 | 0.008827474 | 0.658304855 | 1.73E-05 | 0.639937028 | 0.993542839 |
| Oxr1 | 0.584875793 | 0.012037938 | 0.618372498 | 4.33E-05 | 0.021981635 | 0.993542839 |
| Kcnab2 | 0.58165171 | 0.019359494 | 0.516878506 | 0.007330987 | -0.14900658 | 0.993542839 |
| Phyh | 0.57547192 | 0.001668721 | 0.503306389 | 0.00023936 | 0.080681568 | 0.993542839 |
| Rpl30 | 0.561823393 | 0.001569032 | 0.639476701 | 4.43E-07 | 0.385176652 | 0.993542839 |
| Edil3 | 0.558720492 | 0.007593416 | 0.705421507 | 9.04E-09 | 0.131932109 | 0.993542839 |
| Efhd2 | 0.554579718 | 0.039354321 | 0.658212124 | 0.000147484 | 0.548700885 | 0.993542839 |
| Rpl22l1 | 0.552741854 | 0.002592636 | 0.574751265 | 5.42E-05 | 0.315116832 | 0.993542839 |
| Scn1b | 0.552271605 | 0.007378557 | 0.618140966 | 0.000119088 | 0.06740614 | 0.993542839 |
| Rpl38 | 0.54781684 | 0.002015026 | 0.526100513 | 2.86E-05 | 0.249128519 | 0.993542839 |
| Khdrbs1 | 0.539199922 | 0.007563272 | 0.659805587 | 3.87E-12 | 0.192523869 | 0.993542839 |
| Tmx4 | 0.531605801 | 0.000702816 | 0.51284488 | 5.44E-08 | -0.019594484 | 0.993542839 |
| C1qa | 0.531495291 | 0.039354321 | 0.564338622 | 0.003243241 | 0.64428574 | 0.993542839 |
| Rpl10 | 0.531264364 | 0.000155028 | 0.690002716 | 2.19E-07 | 0.377658322 | 0.993542839 |

| | | | | | | |
|---|---|---|---|---|---|---|
| B230118H07Rik | 0.530332022 | 0.030057156 | 0.593454208 | 1.05E-05 | -0.013061567 | 0.993542839 |
| Tspyl4 | 0.527991524 | 5.25E-06 | 0.676806176 | 7.10E-36 | 0.226852292 | 0.993542839 |
| Rit2 | 0.521103793 | 0.016428564 | 0.506602283 | 2.00E-06 | -0.07075115 | 0.993542839 |
| Mapt | 0.50934293 | 0.008642206 | 0.548195395 | 0.000166372 | 0.064675988 | 0.993542839 |
| Ogfrl1 | 0.500970177 | 0.022887304 | 0.550489457 | 0.000725922 | 0.280210182 | 0.993542839 |
| Slc24a3 | -0.500666112 | 0.023524573 | -0.82445329 | 3.03E-06 | -1.23097609 | 0.251199566 |
| Crym | -0.507602838 | 0.037134763 | -0.546945523 | 0.016819914 | -0.640357597 | 0.993542839 |
| Mgat4b | -0.50877059 | 0.0301224 | -0.579537169 | 2.18E-06 | -1.044444307 | 0.442019435 |
| Adap1 | -0.518778467 | 0.003069566 | -0.69844919 | 1.45E-10 | -1.08421876 | 0.519191726 |
| Klhdc10 | -0.526892523 | 0.04685111 | -0.614702902 | 6.15E-06 | -0.947049473 | 0.479890367 |
| Sirt2 | -0.532402773 | 0.002977712 | -0.620815188 | 1.52E-05 | -1.008268329 | 0.556086383 |
| Chl1 | -0.541350283 | 0.003757451 | -0.549836928 | 8.45E-07 | -0.823613349 | 0.895232727 |
| Gpcpd1 | -0.5516428 | 0.025703418 | -0.523420171 | 0.004143247 | -0.881455725 | 0.57790817 |
| Ptprf | -0.552214363 | 0.046577903 | -0.622323484 | 0.001170895 | -0.986127551 | 0.516861724 |
| Nlgn3 | -0.559683913 | 0.00529548 | -0.506013317 | 1.25E-05 | -0.795740797 | 0.871465753 |
| Ttc9b | -0.56081714 | 1.73E-05 | -0.551086778 | 3.65E-16 | -0.885543757 | 0.987466221 |
| Cx3cl1 | -0.561414799 | 0.000179934 | -0.513642822 | 9.99E-08 | -0.766179747 | 0.993542839 |
| Sidt2 | -0.566960905 | 0.028638925 | -0.642913649 | 1.92E-06 | -1.225114331 | 0.158984606 |
| Car4 | -0.56755298 | 0.044894924 | -0.613209864 | 0.001734475 | -0.863671799 | 0.606369617 |
| Trim46 | -0.568546971 | 0.019249448 | -0.671447583 | 1.65E-14 | -1.136122477 | 0.325535997 |
| Chga | -0.571364567 | 0.001433846 | -0.511507954 | 0.000315214 | -0.842248455 | 0.898104584 |
| Parp6 | -0.573394818 | 0.002717102 | -0.732574742 | 8.67E-11 | -1.036973114 | 0.521778447 |
| Rimbp2 | -0.582227616 | 0.04634046 | -0.514797572 | 0.00050357 | -0.825634135 | 0.881557443 |
| Nfasc | -0.585139836 | 0.009722792 | -0.536975317 | 9.92E-06 | -1.143543783 | 0.400075453 |
| Nsd2 | -0.58706719 | 0.029133243 | -0.55480106 | 0.000212851 | -0.91954248 | 0.498366799 |
| Sh3bp5l | -0.588264631 | 0.043049037 | -0.665091819 | 4.70E-10 | -0.992521995 | 0.28840387 |
| Akap8l | -0.594286795 | 0.002607229 | -0.530907845 | 4.61E-07 | -0.911381544 | 0.702744926 |
| Mmp24 | -0.595864362 | 0.031313396 | -0.613675587 | 8.25E-07 | -1.427960374 | 0.073853991 |
| Klhl23 | -0.595872408 | 0.042998383 | -0.566221297 | 9.38E-05 | -0.819062575 | 0.77914481 |
| Gng2 | -0.596660415 | 0.008175325 | -0.562596272 | 1.35E-06 | -0.869986224 | 0.784284494 |
| Cask | -0.605369432 | 0.033992548 | -0.684521274 | 7.85E-09 | -1.034019853 | 0.347082232 |
| Ache | -0.606587078 | 0.037963165 | -0.71946581 | 3.21E-06 | -1.107393432 | 0.370641325 |
| Irgq | -0.606599885 | 0.001850355 | -0.656319191 | 1.45E-10 | -0.934144116 | 0.556719841 |
| Ttyh3 | -0.618176015 | 0.006591968 | -0.89817993 | 2.07E-06 | -1.217102792 | 0.340258598 |
| Rprml | -0.625857415 | 0.018328997 | -0.536135392 | 5.64E-06 | -0.63637693 | 0.993542839 |
| Flywch2 | -0.643120573 | 0.033294254 | -0.548625253 | 0.000844645 | -1.153073918 | 0.296097344 |
| Myorg | -0.647344386 | 0.031627954 | -0.610626333 | 0.001042329 | -1.147294592 | 0.162913907 |
| Tmem223 | -0.652819861 | 4.63E-05 | -0.521642386 | 1.44E-10 | -1.027720988 | 0.442019435 |
| Lrfn4 | -0.666967441 | 0.047770303 | -0.647202238 | 2.32E-05 | -0.965179885 | 0.484545871 |
| Sprn | -0.670957071 | 0.003687388 | -0.558985699 | 0.000480794 | -0.790624342 | 0.893618261 |
| Csdc2 | -0.671413634 | 0.00458145 | -0.649587682 | 0.000543422 | -1.017268168 | 0.409699752 |
| Gng3 | -0.672950974 | 3.86E-05 | -0.534165645 | 5.05E-16 | -0.846210873 | 0.993542839 |
| Mir9-3hg | -0.67391867 | 0.023915174 | -0.531184626 | 0.019117482 | -0.917902266 | 0.657564323 |
| Zfp821 | -0.679590065 | 0.049127246 | -0.564098607 | 5.22E-07 | -0.841578797 | 0.554606532 |
| Agrn | -0.681206857 | 0.00458145 | -0.722396539 | 3.79E-08 | -1.103220559 | 0.324433351 |
| Ankrd13b | -0.683152295 | 0.010454503 | -0.693189319 | 1.65E-06 | -1.308593767 | 0.179251143 |
| Fnbp1l | -0.685482907 | 0.005628478 | -0.555114608 | 9.93E-05 | -1.22721982 | 0.253541743 |
| Map4k4 | -0.685722981 | 0.019577495 | -0.60088523 | 0.001303118 | -1.046765675 | 0.282076286 |
| Nectin1 | -0.686946002 | 0.031313396 | -0.627446665 | 7.18E-05 | -0.754693778 | 0.715362667 |
| Plppr5 | -0.690410523 | 0.047179792 | -0.574103814 | 0.000386696 | -0.553770378 | 0.993542839 |
| Stmn2 | -0.695828111 | 9.90E-05 | -0.604359526 | 1.78E-07 | -1.327555247 | 0.442019435 |
| Sestd1 | -0.70082071 | 0.030246679 | -0.661391702 | 2.39E-05 | -0.922338067 | 0.559129997 |
| Bdh1 | -0.701807395 | 0.028189417 | -0.802470409 | 0.000197948 | -1.460895325 | 0.079621411 |
| Slc8a2 | -0.706794066 | 0.001932683 | -0.637651483 | 0.000123926 | -0.74317921 | 0.993542839 |
| Arhgdig | -0.714336236 | 8.30E-05 | -0.735685563 | 2.16E-10 | -1.188940651 | 0.432818012 |
| Ccnl2 | -0.718817046 | 0.003163983 | -0.663597927 | 2.16E-05 | -1.100150238 | 0.424642494 |
| Inha | -0.720380623 | 0.043913351 | -1.13291754 | 1.58E-10 | -1.387704618 | 0.070643937 |
| Shisa7 | -0.72139461 | 0.002607229 | -0.626981562 | 5.73E-07 | -0.932280489 | 0.680885348 |
| Hs6st1 | -0.722219871 | 0.008579456 | -0.597016879 | 1.20E-05 | -0.944539283 | 0.538557798 |
| Scn3b | -0.723682814 | 0.004374324 | -0.571849286 | 0.000224475 | -0.818475501 | 0.850687986 |
| Itm2a | -0.728621629 | 0.016219129 | -0.874868553 | 8.82E-10 | -1.230237567 | 0.07580789 |
| Ing4 | -0.73301978 | 0.005346415 | -0.564200266 | 1.00E-05 | -0.820971034 | 0.661491752 |
| Cdkl2 | -0.737457802 | 0.041336761 | -0.663897466 | 0.000513525 | -1.237472188 | 0.199573866 |
| Adam11 | -0.741857699 | 0.002906108 | -0.672977087 | 1.95E-08 | -0.958610992 | 0.655823341 |
| Carmil3 | -0.745278858 | 0.03588093 | -0.690516707 | 5.94E-05 | -1.324743843 | 0.158106894 |
| 1810058I24Rik | -0.760208237 | 0.002038638 | -0.546027883 | 4.87E-06 | -0.990423755 | 0.45949895 |
| Mindy1 | -0.767872423 | 0.021327761 | -0.524017634 | 0.00249654 | -0.778618752 | 0.526849179 |
| Tdg | -0.768671956 | 0.025765148 | -0.757719719 | 7.22E-06 | -1.300661501 | 0.235195451 |
| Gm3764 | -0.76958086 | 0.007509124 | -0.882444598 | 6.14E-06 | -1.295648622 | 0.118078732 |
| Kmt2b | -0.771600685 | 0.023000062 | -0.55490078 | 0.001284865 | -0.812621089 | 0.608884004 |
| Arxes1 | -0.772216573 | 0.020123855 | -0.592525855 | 5.48E-05 | -1.105181498 | 0.400350544 |
| Slc25a27 | -0.782718754 | 0.018752082 | -0.502943738 | 3.09E-05 | -0.808068012 | 0.705185351 |
| Tmem181a | -0.784750957 | 0.000389992 | -0.776447187 | 2.24E-09 | -1.315810693 | 0.144616872 |

| | | | | | | |
|---|---|---|---|---|---|---|
| Plppr2 | -0.799220387 | 0.000250002 | -0.651395376 | 1.16E-07 | -0.970424359 | 0.681723183 |
| Cacna1g | -0.801461368 | 0.002770726 | -0.563594247 | 0.000818416 | -1.061175563 | 0.378741994 |
| Jpt1 | -0.804957212 | 5.14E-06 | -0.506799808 | 4.53E-10 | -0.894659887 | 0.689744667 |
| Cntnap5a | -0.805978523 | 0.04433394 | -0.723035275 | 0.000813712 | -0.952433005 | 0.719737087 |
| Slc16a2 | -0.809702788 | 0.040547532 | -0.81299495 | 0.00053359 | -1.201781713 | 0.151994712 |
| Sulf2 | -0.814950701 | 0.002613871 | -0.833899533 | 7.92E-05 | -0.963945235 | 0.723623811 |
| Pcp4 | -0.815171955 | 2.57E-05 | -0.573127401 | 0.044931104 | -0.64561193 | 0.993542839 |
| Mllt11 | -0.815780115 | 1.92E-06 | -0.603842856 | 4.32E-09 | -1.031215118 | 0.724954309 |
| Penk | -0.817314752 | 0.003047311 | -1.410349976 | 6.01E-15 | -1.019737623 | 0.239571016 |
| Mir124a-1hg | -0.82163735 | 0.001614544 | -0.589662406 | 0.005363817 | -1.030794518 | 0.526849179 |
| Arhgef1 | -0.823062571 | 0.033447813 | -0.828826469 | 4.28E-06 | -1.288508233 | 0.125734729 |
| Dgkd | -0.824597755 | 1.17E-05 | -0.751796091 | 2.01E-11 | -1.185243609 | 0.149930259 |
| Dot1l | -0.826423879 | 0.016631045 | -0.986049708 | 2.49E-05 | -1.413017634 | 0.173484137 |
| Sema6a | -0.828728543 | 0.037612454 | -0.853284256 | 0.000432723 | -1.170658051 | 0.144616872 |
| Eif4e2 | -0.830131646 | 0.008396043 | -0.575319243 | 3.28E-05 | -1.062682646 | 0.279206837 |
| Rimklb | -0.830666073 | 0.022205482 | -0.857691174 | 2.32E-07 | -1.240567408 | 0.088716508 |
| Nasp | -0.831239325 | 0.048706034 | -0.834901913 | 1.28E-08 | -1.072222951 | 0.257813216 |
| Tubb2a | -0.837810582 | 0.000151162 | -0.700469889 | 2.74E-05 | -1.149720857 | 0.715362667 |
| Spata2l | -0.839459823 | 0.013238105 | -0.98294014 | 4.82E-08 | -0.96002085 | 0.685232864 |
| Cckbr | -0.843788083 | 0.028494451 | -0.699542617 | 0.000318451 | -0.899296161 | 0.708363673 |
| Ypel1 | -0.849026542 | 0.040522348 | -0.685277993 | 0.009213653 | -0.918991541 | 0.442019435 |
| Cldn5 | -0.850472664 | 0.020123855 | -0.823723823 | 4.75E-05 | -1.191243549 | 0.154941158 |
| Meg3 | -0.862005349 | 0.017501483 | -0.996659414 | 0.000156176 | -1.531052567 | 0.270720779 |
| Plpp5 | -0.868259386 | 0.022887304 | -0.627362998 | 0.000734077 | -1.005241123 | 0.424716939 |
| Ndn | -0.876950138 | 8.65E-06 | -0.683611356 | 0.008302239 | -0.579542653 | 0.993542839 |
| Zfp580 | -0.877838066 | 0.003344122 | -0.705689171 | 5.33E-05 | -1.142044146 | 0.318301009 |
| Rgl2 | -0.89625473 | 0.002151982 | -0.645288373 | 0.000265585 | -1.281195569 | 0.082770423 |
| Pkia | -0.90878903 | 1.38E-07 | -0.670386174 | 1.62E-13 | -1.007128924 | 0.551475793 |
| Fam57b | -0.909389357 | 2.22E-05 | -0.869344234 | 4.37E-10 | -1.150916567 | 0.305112527 |
| Igsf21 | -0.915081457 | 0.000494217 | -0.950246424 | 1.25E-07 | -0.772414544 | 0.993542839 |
| Bmp1 | -0.936196448 | 0.00084611 | -1.321472695 | 5.82E-11 | -1.527833534 | 0.061954408 |
| Sbk1 | -0.936472724 | 0.000993165 | -0.71667726 | 1.25E-05 | -0.875333266 | 0.663864381 |
| Dpysl4 | -0.941084547 | 4.63E-05 | -1.025110167 | 3.19E-09 | -1.321407892 | 0.21453085 |
| Gpx8 | -0.948950149 | 0.025314375 | -1.763147155 | 4.53E-09 | -1.686052131 | 0.008221945 |
| Igsf9b | -0.954520039 | 0.003470292 | -0.829043245 | 0.000154321 | -1.317282659 | 0.125734729 |
| Clec2l | -0.956018256 | 2.92E-05 | -0.831642782 | 2.92E-08 | -1.355903649 | 0.163455884 |
| Esam | -0.960015921 | 0.023110981 | -0.932318143 | 0.000109965 | -1.082224591 | 0.133225497 |
| Narf | -0.961223918 | 0.006832918 | -0.59294804 | 3.05E-05 | -0.854454019 | 0.556775701 |
| Gdpd5 | -0.969615917 | 0.017897814 | -0.625209401 | 0.000285931 | -0.918338377 | 0.68509699 |
| Pglyrp1 | -0.987145204 | 0.004936233 | -0.741889692 | 0.006137747 | -1.020597284 | 0.328516758 |
| Tmem132a | -0.988854335 | 1.48E-09 | -0.918349228 | 9.80E-11 | -1.140786698 | 0.57790817 |
| Sox11 | -1.001807922 | 0.048706034 | -0.720370607 | 3.03E-05 | -0.382803035 | 0.993542839 |
| Slco1c1 | -1.005366048 | 0.005474021 | -0.98263468 | 1.49E-06 | -1.179561164 | 0.254591155 |
| Hist3h2ba | -1.005860677 | 2.84E-05 | -0.831223872 | 9.88E-09 | -1.097026951 | 0.36792686 |
| Impdh1 | -1.006233098 | 7.28E-05 | -0.591138491 | 4.13E-05 | -0.939312957 | 0.504616615 |
| Rhbdl1 | -1.016924989 | 3.50E-05 | -0.926002945 | 1.21E-11 | -1.468946642 | 0.134188632 |
| Dcakd | -1.017009807 | 0.00104277 | -0.935746385 | 2.85E-10 | -1.245568263 | 0.120455841 |
| Grm4 | -1.020373579 | 0.003275081 | -1.018580106 | 9.97E-09 | -1.022403893 | 0.156304324 |
| Nav3 | -1.044394493 | 0.002625878 | -1.064657472 | 3.84E-06 | -1.299119496 | 0.268832642 |
| Neto1 | -1.044866827 | 0.000588409 | -0.997182855 | 2.09E-09 | -0.839617465 | 0.858634628 |
| Miat | -1.050293871 | 0.000282345 | -1.097892156 | 1.98E-06 | -1.161871261 | 0.448141348 |
| Sppl2b | -1.06754982 | 0.002291593 | -1.066017299 | 4.53E-10 | -1.366800448 | 0.105698283 |
| Erc1 | -1.072253008 | 0.001607052 | -0.98853474 | 6.79E-05 | -1.458289529 | 0.162913907 |
| Nav1 | -1.081357224 | 0.002613871 | -1.150524996 | 2.72E-06 | -1.431783975 | 0.122155429 |
| Focad | -1.089384475 | 0.00122409 | -0.648705359 | 1.66E-05 | -0.891973685 | 0.448141348 |
| Ncan | -1.090958456 | 3.05E-05 | -1.243341905 | 6.73E-09 | -1.660355325 | 0.074947024 |
| Deaf1 | -1.108418308 | 0.000134312 | -0.841793338 | 9.61E-05 | -1.255177934 | 0.299019157 |
| Lhx1os | -1.116631417 | 0.032280369 | -1.547610096 | 0.00013349 | -1.098417782 | 0.423882429 |
| Ccdc28b | -1.130558419 | 3.56E-05 | -1.048774979 | 3.18E-13 | -1.233783683 | 0.171842435 |
| Hmgcs2 | -1.234304984 | 0.005633612 | -1.203854403 | 0.002426019 | -1.302712959 | 0.249573326 |
| Slc16a1 | -1.289349771 | 0.019721157 | -0.870765382 | 0.043590964 | -0.90027574 | 0.676759681 |
| Ddah2 | -1.298965913 | 1.19E-05 | -0.580417001 | 0.002855266 | -1.068897275 | 0.199573866 |
| Ppp1r14c | -1.638351168 | 0.001984829 | -0.962367794 | 0.000121441 | -1.568715382 | 0.083081883 |
| Gm47283 | -1.890138881 | 0.035184648 | -1.949631577 | 0.003946937 | -2.203456701 | 0.116921317 |
| Dpysl5 | -2.149747848 | 3.80E-06 | -1.930016759 | 6.07E-06 | -1.811148567 | 0.14509505 |
| Marcksl1 | -2.151220967 | 1.01E-05 | -2.209543178 | 1.18E-06 | -1.883026828 | 0.186666193 |
| n = 1108 | MEC_s | | MEC_im | | MEC_d | |
| gene | log2FoldChange | padj | log2FoldChange | padj | log2FoldChange | padj |
| Smok2b | 2.69080039 | NA | 6.891944797 | 1.52E-12 | 0.591714905 | 0.993542839 |
| Gm16867 | 3.517751966 | NA | 6.273036808 | 2.14E-14 | 1.27914534 | 0.660210965 |
| Gm15446 | 4.06840004 | 9.28E-05 | 5.881617746 | 1.78E-14 | 3.247199614 | 8.18E-05 |
| Cst7 | 3.416872125 | 0.005633612 | 5.799561964 | 7.62E-09 | 3.292406796 | 0.00729645 |

| | | | | | | |
|---|---|---|---|---|---|---|
| Gsta2 | 3.625799467 | 0.005633612 | 4.210924582 | 0.000349334 | 1.337481373 | 0.813604965 |
| Scnn1a | 1.113754016 | 0.22295652 | 2.770153191 | 6.27E-10 | 0.590595348 | 0.993542839 |
| Egr2 | 3.648669806 | 0.001790406 | 2.266517179 | 2.03E-05 | 1.82511721 | 0.048544447 |
| Spag6 | 0.518508855 | 0.77469308 | 2.252180792 | 2.40E-08 | 1.578166598 | 0.079621411 |
| Gm13530 | 1.874601548 | 0.217240661 | 2.223862659 | 6.02E-10 | 1.161764054 | 0.184983612 |
| Serpinb1a | 1.459993504 | 0.021010927 | 2.203402341 | 3.44E-11 | 1.763911338 | 0.020142081 |
| Galnt6 | 0.603163561 | 0.732765907 | 2.006187631 | 4.62E-05 | 0.752764402 | 0.86014666 |
| Pmaip1 | 2.343785651 | NA | 1.935562799 | 8.82E-07 | 0.578956255 | 0.993542839 |
| Ccdc36 | 1.050705913 | 0.247782387 | 1.858197342 | 0.000682061 | 1.968360796 | 0.056537244 |
| Hapln2 | 1.356908698 | 0.058856564 | 1.849452261 | 0.004983975 | 1.596615262 | 0.105925498 |
| Ifit3 | 2.019674127 | 7.06E-05 | 1.848914271 | 5.43E-06 | 0.92587037 | 0.551429985 |
| Hsf4 | 3.543051619 | 0.000683298 | 1.802376209 | 0.000818416 | 1.31523575 | 0.123169917 |
| Pcdhb8 | 1.40013889 | NA | 1.795837879 | 2.81E-05 | 0.487751668 | 0.993542839 |
| Gm33651 | 2.338230071 | NA | 1.793062998 | 0.000685019 | 0.158560868 | 0.993542839 |
| Gjb1 | 1.788918411 | 0.03261143 | 1.770246145 | 1.92E-08 | 1.173536709 | 0.422664559 |
| Npas4 | 1.675276089 | 0.013312561 | 1.754299408 | 0.000469595 | 1.259259247 | 0.57790817 |
| Tnfaip6 | 0.726532111 | 0.411686858 | 1.719816686 | 0.004582646 | 1.251677938 | 0.150486737 |
| Gpnmb | 3.143305092 | 0.008030733 | 1.680798985 | 0.00015012 | 0.960536962 | 0.638510559 |
| Apol8 | 0.754970093 | 0.565027412 | 1.678515503 | 0.000109048 | 0.039370069 | 0.993542839 |
| Ankrd22 | 1.090032077 | NA | 1.666362836 | 0.001383282 | 0.229361575 | 0.993542839 |
| Nrp1 | 1.44322923 | 0.195183393 | 1.628056533 | 0.001328784 | 1.553268206 | 0.072051016 |
| Tnnt2 | 1.252053689 | 0.054567551 | 1.622446642 | 5.67E-17 | 1.058896737 | 0.606369617 |
| Anxa4 | 0.272952331 | 0.793805165 | 1.59375392 | 2.73E-06 | 0.288254272 | 0.993542839 |
| Pcdhb3 | 0.839044104 | 0.37912849 | 1.579475376 | 4.06E-07 | 1.002933696 | 0.333187521 |
| Hmox1 | -0.527524457 | 0.65511037 | 1.578241787 | 4.03E-05 | 0.235748241 | 0.993542839 |
| Mapk13 | 0.532074691 | NA | 1.574779586 | 0.000466386 | 1.485373507 | 0.206337599 |
| Ntn5 | 1.746066596 | 0.039354321 | 1.563766502 | 0.007864245 | 1.485917072 | 0.150935904 |
| Tmem267 | 1.842439829 | 0.000195117 | 1.560922019 | 6.43E-11 | 1.042149541 | 0.655539463 |
| Pcdhb6 | 1.216780062 | NA | 1.515572997 | 3.34E-05 | 0.5572307 | 0.993542839 |
| Gm40518 | 1.606058247 | 0.009046823 | 1.512187754 | 0.019291698 | 1.839503386 | 0.01602967 |
| Strn3 | 1.396535693 | 0.099726237 | 1.502860617 | 0.000345263 | 0.760217594 | 0.81479986 |
| Tagln | 1.294216806 | 0.151752913 | 1.444432043 | 0.010359425 | 0.59888646 | 0.993542839 |
| Tap2 | 1.593069237 | 0.004374324 | 1.440271684 | 3.79E-06 | 0.241412358 | 0.993542839 |
| Dmp1 | 1.42642139 | 0.040547532 | 1.433186516 | 7.12E-05 | 1.082060647 | 0.460254618 |
| Espn | 0.74317319 | 0.574532388 | 1.425141714 | 0.000272238 | 0.295298116 | 0.993542839 |
| Glt8d2 | 0.411518839 | 0.736604504 | 1.420197587 | 0.000126781 | 0.627190436 | 0.993542839 |
| S1pr5 | 1.855857671 | 0.008175325 | 1.375894091 | 5.62E-05 | 0.87249395 | 0.792577163 |
| Zfp367 | 0.914227691 | 0.159750461 | 1.369170735 | 7.57E-16 | 0.614319822 | 0.993542839 |
| Tmem88b | 1.626438571 | 0.000272913 | 1.364861194 | 2.95E-05 | 0.898335289 | 0.506210118 |
| Pls1 | 0.526221692 | 0.629519068 | 1.36346897 | 9.04E-09 | 0.83703405 | 0.565412228 |
| Aspa | 1.419206122 | 0.002291593 | 1.362485795 | 2.46E-05 | 1.277076708 | 0.120455841 |
| C1ql3 | 0.629546684 | 0.495669634 | 1.361605932 | 0.006963703 | 0.24151763 | 0.993542839 |
| Zc3hav1 | 0.813467553 | 0.193140335 | 1.344717266 | 0.000199781 | 0.742679867 | 0.95811614 |
| Cd82 | 0.889820029 | 0.098108782 | 1.334752581 | 4.13E-06 | 1.046280217 | 0.562024055 |
| Ctxn3 | 0.510862616 | 0.547053657 | 1.328618863 | 0.000791696 | -0.142370534 | 0.993542839 |
| Litaf | 0.882622289 | 0.052078173 | 1.324896583 | 1.61E-09 | 0.818371399 | 0.67405508 |
| Gstp1 | 1.136632481 | 0.105621027 | 1.305028777 | 0.000515673 | 0.719189105 | 0.993542839 |
| Zfp296 | 1.381977486 | NA | 1.295075291 | 0.001598044 | 0.900241042 | 0.715362667 |
| Abca8a | 1.299644668 | 0.176606475 | 1.278073675 | 0.000775905 | 0.727798518 | 0.786915421 |
| Opalin | 1.683484146 | 0.003528462 | 1.262549527 | 0.003773439 | 1.214789633 | 0.498580941 |
| Itgb2 | 0.885574952 | 0.239347525 | 1.259095508 | 6.58E-05 | 0.9333572 | 0.868591714 |
| Nkx6-2 | 1.028227745 | 0.143058809 | 1.257142472 | 0.000651399 | 0.771430364 | 0.738233607 |
| Kank4 | 0.712496999 | 0.339277933 | 1.246571813 | 8.03E-05 | -0.151601947 | 0.993542839 |
| Aif1l | 0.413374875 | 0.662779962 | 1.234147302 | 0.000101782 | 0.765034064 | 0.858634628 |
| Arsg | 1.19281224 | 0.083424852 | 1.229342124 | 0.000180877 | 0.890787281 | 0.482241964 |
| Tmem254a | 0.502055136 | 0.77199044 | 1.228881575 | 0.000235228 | 1.02500305 | 0.48442902 |
| Pde8a | 0.533593028 | 0.551855319 | 1.220577549 | 0.000207816 | 1.057266076 | 0.274235252 |
| Psmb8 | 1.877906766 | 3.50E-05 | 1.207854596 | 1.78E-05 | 0.888366348 | 0.613158534 |
| Plekhh1 | 0.642820665 | 0.452854097 | 1.204883519 | 0.004597198 | 1.07432267 | 0.503069679 |
| Dkkl1 | 0.744647542 | 0.297203972 | 1.199076784 | 0.000771843 | -0.105083025 | 0.993542839 |
| Erbb3 | 0.660667219 | NA | 1.189460293 | 0.006999943 | 0.824965683 | 0.746685932 |
| Pcdhb9 | 1.181159946 | 0.014047239 | 1.17929632 | 1.92E-09 | 1.436477722 | 0.079943479 |
| Smco3 | 0.594982731 | 0.52920493 | 1.172343964 | 0.001170895 | 0.317301506 | 0.993542839 |
| Glra3 | 0.673134158 | 0.618266847 | 1.170101094 | 0.008598043 | -0.28228898 | 0.993542839 |
| Gpat3 | 2.43903754 | 0.082966831 | 1.166335143 | 0.002135504 | 0.374860876 | 0.993542839 |
| Anxa11 | 0.740335095 | 0.149106168 | 1.162850621 | 3.97E-07 | -0.086233637 | 0.993542839 |
| Tmem63a | 0.916199511 | 0.091728428 | 1.156963636 | 0.001250626 | 0.710694221 | 0.993542839 |
| Kazald1 | 1.075475846 | 0.238100445 | 1.156305618 | 0.003305154 | 1.400325126 | 0.053194932 |
| Fbln5 | 0.497844345 | 0.539736543 | 1.151123077 | 0.001561377 | 0.044898098 | 0.993542839 |
| Pcdhb11 | 1.570257207 | 0.214453752 | 1.151119723 | 0.002306018 | -0.134063444 | 0.993542839 |
| Prr5l | 1.449836479 | 0.183011401 | 1.1466929 | 0.006242443 | 0.894201052 | 0.619705337 |
| Hspa1b | 1.031501741 | 0.226687412 | 1.133552847 | 0.00022135 | 0.327200017 | 0.993542839 |
| Trh | -0.499666941 | NA | 1.129727355 | 0.039871306 | -0.770042008 | 0.98655832 |

| | | | | | | |
|---|---|---|---|---|---|---|
| Trim59 | 1.159613134 | 0.094414071 | 1.125190043 | 5.94E-05 | 0.565616133 | 0.993542839 |
| Pcdhb13 | 0.908626487 | 0.493526194 | 1.121154241 | 1.85E-05 | 0.661849412 | 0.993542839 |
| Cox8b | 1.591908609 | 0.158496999 | 1.112293338 | 0.013514914 | 0.384774666 | 0.993542839 |
| Plekhg3 | 0.695339186 | 0.328374681 | 1.103993101 | 0.002504048 | 0.657581313 | 0.914497701 |
| Adssl1 | 1.447157655 | 0.001387144 | 1.096395088 | 4.91E-06 | 1.217025352 | 0.144643344 |
| Irgm1 | 0.955038467 | 0.04168442 | 1.095767719 | 1.59E-05 | 0.595999621 | 0.940150584 |
| Gm14419 | 1.017683504 | 0.240557413 | 1.094122748 | 2.74E-06 | 0.255414517 | 0.993542839 |
| Serpina3n | 1.188679122 | 0.045283476 | 1.084972339 | 0.000861924 | 0.995785658 | 0.562394156 |
| Cfap58 | 0.882004702 | 0.338579303 | 1.079537338 | 0.001517883 | -0.500252729 | 0.993542839 |
| B230206L02Rik | 0.193268746 | 0.902742414 | 1.078879484 | 0.000576233 | 0.617562346 | 0.993542839 |
| Pcdhb14 | 1.559366842 | 0.015436508 | 1.075635973 | 7.08E-07 | 0.417987197 | 0.993542839 |
| Utp14b | 1.533257962 | 0.001579309 | 1.07372668 | 0.000523295 | 0.72998131 | 0.914338763 |
| AW495222 | 1.168783162 | 0.153656005 | 1.066812781 | 9.88E-06 | 0.364062507 | 0.993542839 |
| Rgs16 | 0.912003507 | 0.220707889 | 1.062757029 | 0.001778586 | 0.147405163 | 0.993542839 |
| Tmem98 | 0.541873679 | 0.433561182 | 1.061616295 | 9.05E-05 | 0.49781711 | 0.993542839 |
| Mafb | 0.947679771 | 0.02256716 | 1.060549122 | 1.43E-11 | 0.449944283 | 0.993542839 |
| Ccn1 | 1.417304886 | 0.084508582 | 1.055406092 | 0.001996995 | 1.187480631 | 0.57790817 |
| Deptor | 0.783441113 | 0.093079342 | 1.055083423 | 9.49E-05 | 0.669287758 | 0.993542839 |
| Mthfd2 | 0.399526335 | 0.690354982 | 1.048422024 | 6.10E-07 | -0.050270075 | 0.993542839 |
| Crb3 | 2.343022173 | NA | 1.047477547 | 0.003251324 | 0.109228083 | 0.993542839 |
| Car14 | -0.094782792 | 0.907149841 | 1.046533188 | 0.000362408 | 0.238385683 | 0.993542839 |
| Pkib | 0.626311835 | 0.226283584 | 1.035627915 | 1.45E-10 | 0.302734699 | 0.993542839 |
| Ccl6 | 0.999397004 | 0.203499875 | 1.034134188 | 0.006098916 | 1.410572794 | 0.466493636 |
| Asb16 | 0.973350953 | 0.51805184 | 1.030295709 | 0.002872582 | 1.039384559 | 0.658009065 |
| Alox8 | 0.595696818 | 0.742975796 | 1.025394767 | 0.000205762 | 0.215023236 | 0.993542839 |
| Arhgap15 | 1.681820533 | 0.051689097 | 1.01545608 | 3.71E-05 | 0.829809068 | 0.883573732 |
| Atf5 | 0.59426807 | 0.334820782 | 1.013125215 | 1.80E-06 | -0.067498317 | 0.993542839 |
| Kcnk13 | 0.82248544 | 0.594279783 | 1.01173079 | 0.014130623 | 0.227592358 | 0.993542839 |
| Itgb4 | 1.148918717 | 0.138435584 | 1.008705627 | 0.022650533 | 1.025861996 | 0.480220174 |
| Arsj | -0.324766783 | 0.886352089 | 1.007615765 | 0.009103435 | 0.038659642 | 0.993542839 |
| Lipo3 | 0.405167399 | 0.667891127 | 1.004781804 | 0.001020332 | -0.155998828 | 0.993542839 |
| Sgk1 | 0.395939076 | 0.097931431 | 0.997202104 | 4.47E-08 | 0.574294173 | 0.993542839 |
| Smco4 | 0.339890004 | 0.826514341 | 0.990632196 | 0.002323632 | 0.487964131 | 0.993542839 |
| Ddo | 0.259123326 | 0.849878018 | 0.990488013 | 0.00817617 | 0.247115612 | 0.993542839 |
| 2900041M22Rik | 1.585716636 | 0.115007936 | 0.988843498 | 0.008167617 | 0.565890349 | 0.993542839 |
| Hspa1a | 0.55676181 | 0.337601945 | 0.986030531 | 2.08E-09 | 0.622767492 | 0.828989507 |
| Myo1d | 0.020838753 | 0.990876969 | 0.98227769 | 0.003037308 | 0.69200127 | 0.993542839 |
| Chil1 | 1.278482509 | 0.096290189 | 0.97993597 | 0.000240736 | 0.51469731 | 0.993542839 |
| Fcgr2b | 0.486294373 | 0.623825476 | 0.97387625 | 0.012720764 | 0.769211024 | 0.993542839 |
| Mcam | 1.234632478 | 0.105026208 | 0.97238146 | 0.002553334 | -0.067687998 | 0.993542839 |
| Cmip | 1.171184653 | 0.088041399 | 0.96790534 | 0.027061596 | 0.076510403 | 0.993542839 |
| Acat3 | 1.117360814 | 0.321578098 | 0.964316988 | 0.012888512 | 1.087120197 | 0.480220174 |
| B230209E15Rik | 1.096251773 | 0.459997479 | 0.963543961 | 0.027946482 | 0.183331023 | 0.993542839 |
| Pold1 | 1.187496864 | 0.039991671 | 0.961800513 | 0.000159867 | 0.207000453 | 0.993542839 |
| Zfp503 | 0.393310041 | 0.57916112 | 0.960337238 | 0.000174265 | 1.527935608 | 0.306134629 |
| Tapbpl | 1.442819206 | 0.040696227 | 0.944314383 | 0.001647426 | 0.257460279 | 0.993542839 |
| Dusp5 | 0.112471754 | 0.874813296 | 0.94422803 | 6.23E-08 | -0.165544062 | 0.993542839 |
| Smad9 | 0.605088455 | 0.469915459 | 0.938144227 | 0.001793857 | 0.084349166 | 0.993542839 |
| Depp1 | 2.328142161 | 0.00313805 | 0.930127234 | 0.006348746 | 0.739362931 | 0.993542839 |
| Plaat1 | 0.841614895 | 0.216326306 | 0.928834831 | 1.00E-07 | 0.75979007 | 0.759725354 |
| Tmem243 | 0.452757935 | 0.462237656 | 0.925368387 | 0.001129631 | 0.787874821 | 0.582320142 |
| Nr4a3 | 0.622788525 | 0.101311846 | 0.924343843 | 5.78E-05 | 0.712830535 | 0.88759934 |
| Efcab10 | 0.024775804 | 0.984979144 | 0.921272165 | 1.93E-10 | 0.732373692 | 0.79107705 |
| Syt2 | 0.152649202 | 0.826480535 | 0.905029949 | 0.001637414 | 0.431902711 | 0.993542839 |
| Prtn3 | 1.399524037 | 0.206471432 | 0.904964038 | 0.003567293 | -0.296413878 | 0.993542839 |
| Mansc1 | 0.45260247 | 0.667891127 | 0.897459459 | 0.00027091 | 0.627784204 | 0.983116755 |
| Crhbp | 0.842596756 | 0.107665506 | 0.89146721 | 6.02E-10 | 0.418084596 | 0.993542839 |
| Aass | 0.528860099 | 0.625266764 | 0.89034517 | 0.005797165 | 0.085318112 | 0.993542839 |
| Gm36251 | 1.839690939 | 0.105946354 | 0.888688639 | 0.027189766 | -0.866785101 | 0.712523443 |
| Ccdc82 | 0.088121659 | 0.943169213 | 0.887981665 | 0.000250461 | -0.20746438 | 0.993542839 |
| Fa2h | 0.570775051 | 0.309764433 | 0.887852744 | 0.001553159 | 0.306491311 | 0.993542839 |
| Tuft1 | 0.509670078 | 0.805830114 | 0.883114744 | 0.006512379 | 0.742819784 | 0.993542839 |
| Apold1 | 1.30498531 | 0.208725585 | 0.883034812 | 0.004881893 | -0.190428785 | 0.993542839 |
| Fstl4 | 0.307288442 | 0.848089433 | 0.880265229 | 0.015897318 | 0.083632711 | 0.993542839 |
| Gm4631 | 0.185804513 | 0.826480535 | 0.875375628 | 7.18E-07 | -0.216752105 | 0.993542839 |
| Spg20 | 0.474123917 | 0.378710324 | 0.872040997 | 1.97E-05 | 0.697583935 | 0.76110548 |
| Ar | 0.799742207 | 0.251852682 | 0.871872289 | 0.000758064 | 0.345980252 | 0.993542839 |
| Gngt2 | 0.362861019 | 0.646606502 | 0.871652017 | 0.005394991 | 0.682593146 | 0.901238853 |
| Sowahb | 0.395647979 | 0.620135641 | 0.870372018 | 2.44E-05 | 0.615956596 | 0.993542839 |
| Rbmx | 0.709785782 | 0.117911541 | 0.868773941 | 4.70E-11 | 0.371696913 | 0.993542839 |
| Nek7 | 0.227690514 | 0.634623901 | 0.862967342 | 2.35E-13 | 0.209618663 | 0.993542839 |
| Pir | 0.821194414 | 0.19301296 | 0.859704129 | 0.000334474 | 0.279865581 | 0.993542839 |
| Srek1ip1 | 0.572804837 | 0.252705557 | 0.859631862 | 0.000510253 | 0.631876872 | 0.989210266 |

| | | | | | | |
|---|---|---|---|---|---|---|
| Agmo | -0.069683514 | 0.964380375 | 0.851723601 | 0.029348311 | 0.695807401 | 0.993542839 |
| Sik1 | 0.113536065 | 0.946135497 | 0.850944709 | 0.009254107 | -0.01661952 | 0.993542839 |
| Spty2d1 | 0.667456033 | 0.240220281 | 0.849599198 | 7.12E-05 | 0.175778307 | 0.993542839 |
| 9330179D12Rik | 0.266580578 | 0.892536848 | 0.846083821 | 0.017003395 | 0.608724512 | 0.993542839 |
| Zdhhc1 | 0.870533214 | 0.100397606 | 0.842151417 | 0.000160209 | 0.629570814 | 0.861398184 |
| Ankrd6 | 0.720121586 | 0.115858255 | 0.83967669 | 6.07E-06 | 0.029190134 | 0.993542839 |
| Rffl | 0.405198898 | 0.718815147 | 0.83858495 | 0.006685457 | 0.263331357 | 0.993542839 |
| Mtrf1 | 0.126064571 | 0.936938618 | 0.836079319 | 0.000541262 | 0.389914943 | 0.993542839 |
| Pigz | 1.037378445 | 0.122653656 | 0.834181346 | 0.002551312 | 0.529404313 | 0.993542839 |
| Gstm7 | 0.869992549 | 0.09973548 | 0.831325452 | 0.001013583 | 0.620021765 | 0.891192077 |
| B3galt5 | 0.788185367 | 0.234615902 | 0.831263347 | 0.001647426 | -0.073418856 | 0.993542839 |
| Lzts3 | 0.727641999 | 0.098482067 | 0.830480958 | 0.00015012 | 0.648429361 | 0.914497701 |
| Xrcc3 | 0.508076998 | 0.705529877 | 0.829529277 | 0.02700963 | -0.101416584 | 0.993542839 |
| Maob | 0.828168726 | 0.059706483 | 0.828767865 | 5.91E-07 | 0.295885085 | 0.993542839 |
| Synpr | 0.163797596 | 0.682012507 | 0.82789533 | 0.013163977 | -0.280186119 | 0.993542839 |
| Nfkbiz | 0.890919055 | 0.4639562 | 0.827280426 | 0.010321825 | -0.699600127 | 0.914497701 |
| Pdlim1 | 0.891084344 | 0.229985636 | 0.821249428 | 0.008727814 | 0.460983237 | 0.993542839 |
| Akain1 | 0.488132647 | 0.604063549 | 0.821020876 | 0.034362869 | 0.487552811 | 0.993542839 |
| Pde1c | 0.103876888 | 0.933644581 | 0.82084389 | 0.022509481 | -0.610851769 | 0.946657533 |
| Mylk | 0.09891269 | 0.935936753 | 0.818856807 | 0.000730189 | -0.132256235 | 0.993542839 |
| Tmem88 | 0.026351881 | 0.985280397 | 0.818740913 | 0.005040945 | 0.670273449 | 0.898104584 |
| Nfil3 | 0.653410387 | 0.367206908 | 0.818197536 | 4.25E-05 | 0.363794758 | 0.993542839 |
| Aatf | 0.731243203 | 0.383984294 | 0.816523582 | 0.002336272 | 0.403787984 | 0.993542839 |
| Asah2 | -0.161993793 | 0.920113855 | 0.816196591 | 0.003777171 | 0.238432581 | 0.993542839 |
| Nupr1 | 0.128314712 | 0.894085043 | 0.810190155 | 0.001879417 | 0.685038161 | 0.826923103 |
| Sncg | 1.670545067 | 0.019414816 | 0.809233407 | 0.004653187 | 0.520718948 | 0.993542839 |
| Fundc1 | 0.507542158 | 0.100586779 | 0.804389521 | 3.95E-13 | 0.325885087 | 0.993542839 |
| Cdc42ep1 | 0.058784928 | 0.950361554 | 0.801565352 | 0.000713534 | 0.613853345 | 0.914497701 |
| Haus7 | 0.221190722 | 0.845288626 | 0.799181302 | 0.00013708 | 0.51161527 | 0.993542839 |
| Rdm1 | 0.568023641 | 0.345932798 | 0.799028405 | 0.003594071 | 0.672623321 | 0.987418678 |
| Gm44421 | 0.199143484 | 0.899062283 | 0.795941189 | 0.031833124 | -0.520429254 | 0.993542839 |
| Prrg1 | 0.654474739 | 0.565281461 | 0.793917097 | 0.03584029 | -0.012101772 | 0.993542839 |
| Csrnp1 | 0.694173443 | 0.45295527 | 0.793272323 | 0.003335991 | 0.292962925 | 0.993542839 |
| Metrnl | 0.380404242 | 0.51094747 | 0.793271658 | 9.61E-05 | 0.216220496 | 0.993542839 |
| Ldb3 | -0.137209783 | 0.948224704 | 0.789898269 | 0.018246722 | -0.032422323 | 0.993542839 |
| Chac1 | 0.647985704 | 0.426295209 | 0.78768786 | 1.65E-06 | -0.3256523 | 0.993542839 |
| AC160336.1 | 0.248110453 | 0.869962281 | 0.787177673 | 0.049938037 | 0.58257984 | 0.993542839 |
| Dcn | 0.15996892 | 0.777139605 | 0.786041963 | 0.028025468 | -0.611027161 | 0.993542839 |
| Spin2c | 0.140494098 | 0.921788271 | 0.783579688 | 0.001129158 | 0.361058645 | 0.993542839 |
| Ak8 | 1.263442716 | NA | 0.780330974 | 0.048876484 | 0.350617019 | 0.993542839 |
| Mvp | 0.312399959 | 0.735405595 | 0.779570381 | 0.004771715 | -0.111667703 | 0.993542839 |
| Bcas1 | 0.42527851 | 0.288144076 | 0.776681284 | 0.001017992 | 0.512676587 | 0.993542839 |
| A230056P14Rik | 0.84217429 | 0.553455655 | 0.774169825 | 0.013898606 | 0.252990997 | 0.993542839 |
| Rnf207 | 1.648115891 | 0.089757376 | 0.772412475 | 0.010910033 | 0.341385729 | 0.993542839 |
| Nudt17 | 0.399935576 | 0.67479576 | 0.769957921 | 0.000599826 | -0.245254383 | 0.993542839 |
| Rhpn2 | 0.565417386 | 0.61198661 | 0.768617896 | 0.006724882 | 0.185012563 | 0.993542839 |
| Ppie | 0.629970191 | 0.232506469 | 0.767188328 | 0.000238363 | 0.316404813 | 0.993542839 |
| Resp18 | 0.633618246 | 0.076064261 | 0.766969804 | 0.000169863 | 0.19934415 | 0.993542839 |
| Gm30082 | 1.024842082 | 0.297609224 | 0.766551846 | 0.006752083 | 0.210316837 | 0.993542839 |
| Dph7 | -0.029340336 | 0.98263023 | 0.761983936 | 0.000794974 | 0.629094617 | 0.923087314 |
| Fads6 | 0.868096339 | 0.186196304 | 0.761524261 | 0.005722427 | 0.599332871 | 0.993542839 |
| Eif5a2 | 0.581023217 | 0.094264005 | 0.760515418 | 4.77E-07 | 0.162923258 | 0.993542839 |
| Wdyhv1 | 0.739935729 | 0.175061743 | 0.760373993 | 0.000158757 | 0.266559506 | 0.993542839 |
| Sf3b6 | 0.451213133 | 0.01934644 | 0.757154876 | 1.15E-06 | 0.531930064 | 0.993542839 |
| Fam117a | 0.067346611 | 0.95998109 | 0.751414801 | 0.007964982 | -0.021047087 | 0.993542839 |
| Srcap | 0.773519652 | 0.413755362 | 0.750625249 | 0.049848329 | 0.227157787 | 0.993542839 |
| Spp1 | 0.81788755 | 0.248583457 | 0.74974759 | 0.008695632 | 0.565343791 | 0.993542839 |
| Taf9b | 0.641988181 | 0.203458237 | 0.749182039 | 0.000418915 | 0.362900706 | 0.993542839 |
| 1700047M11Rik | 0.307656276 | 0.838453873 | 0.745050516 | 0.022735703 | 0.669479611 | 0.914338763 |
| Zar1l | -0.115574447 | 0.968733724 | 0.74137678 | 0.045079861 | 0.519491447 | 0.993542839 |
| Tmc6 | 0.688939304 | 0.322799071 | 0.740624112 | 0.020467689 | 0.43148345 | 0.993542839 |
| Pi4kb | 0.097796949 | 0.946527147 | 0.740494633 | 0.0069524 | 0.821495423 | 0.650953551 |
| Medag | 0.420666808 | 0.704432434 | 0.73776251 | 0.028934793 | 0.912516833 | 0.625402311 |
| Btg2 | 0.876447842 | 0.067077039 | 0.737715511 | 2.80E-05 | 0.09687554 | 0.993542839 |
| Pcsk6 | 0.315854684 | 0.593506676 | 0.735755 | 0.000429681 | 0.298067243 | 0.993542839 |
| Sstr2 | -0.073758863 | 0.950603139 | 0.735556347 | 0.002130729 | 0.421229273 | 0.993542839 |
| Eepd1 | 0.364463608 | 0.625230326 | 0.733410043 | 0.000222883 | -0.033238417 | 0.993542839 |
| Fbxl14 | 0.615136638 | 0.266452528 | 0.733055332 | 8.50E-06 | -0.097477742 | 0.993542839 |
| Skap2 | 0.50829907 | 0.356763449 | 0.731664823 | 0.000163412 | 0.359364685 | 0.993542839 |
| Fosl2 | 0.663777646 | 0.051099427 | 0.729284962 | 7.20E-08 | 0.250097371 | 0.993542839 |
| Cchcr1 | 0.594853131 | 0.611370244 | 0.726374319 | 0.046772016 | 0.170229248 | 0.993542839 |
| Prdm16os | 1.06479941 | 0.140444408 | 0.725691193 | 0.027061596 | 0.11706835 | 0.993542839 |
| Rerg | 0.290410069 | 0.804664706 | 0.725520683 | 0.002759854 | -0.348746322 | 0.993542839 |

| | | | | | | |
|---|---|---|---|---|---|---|
| Unc5b | 0.073560555 | 0.956589035 | 0.724067892 | 0.021340457 | 0.582031724 | 0.993542839 |
| Mpp5 | 0.232189652 | 0.708373151 | 0.718110552 | 2.57E-05 | 0.434813617 | 0.993542839 |
| Trmt2b | -0.025506339 | 0.984859892 | 0.717721837 | 0.002960367 | 0.177344753 | 0.993542839 |
| Plvap | 0.70665386 | 0.147724528 | 0.715542597 | 0.000696057 | 0.518834193 | 0.993542839 |
| Gadd45b | 0.082137588 | 0.909764621 | 0.715061626 | 2.29E-05 | 1.065841135 | 0.446079578 |
| Arhgef10 | 0.498181529 | 0.332395404 | 0.714701369 | 0.000176735 | 0.906533413 | 0.327844522 |
| Tmem106c | 0.301886888 | 0.496840995 | 0.711725241 | 6.79E-05 | 0.700481644 | 0.80547115 |
| Dnajc12 | 0.68322215 | 0.12849859 | 0.711156386 | 9.53E-05 | 0.220551908 | 0.993542839 |
| Gm38393 | 1.168011111 | 0.343419291 | 0.711106852 | 0.011674717 | -0.524434623 | 0.993542839 |
| 04.Sep | 0.004636088 | 0.993059155 | 0.709161999 | 0.000216224 | 0.526254369 | 0.993542839 |
| Cyp4v3 | 0.529476718 | 0.366332591 | 0.709055106 | 0.015339372 | 0.101709172 | 0.993542839 |
| Tlr3 | 0.262095214 | 0.723954847 | 0.7084226 | 0.011674717 | 0.440979399 | 0.993542839 |
| 2010204K13Rik | 0.540371238 | 0.270413373 | 0.707903765 | 0.004317603 | 0.322678288 | 0.993542839 |
| Lactb2 | 1.064377795 | 0.122853092 | 0.707330023 | 0.001832917 | 0.467098173 | 0.993542839 |
| Cyb5r1 | 0.304712234 | 0.54979138 | 0.705320524 | 9.63E-08 | 0.007410806 | 0.993542839 |
| Nap1l5 | 0.474766448 | 0.000195117 | 0.704448825 | 1.12E-15 | 0.143692826 | 0.993542839 |
| Anln | 1.176935694 | 0.053668131 | 0.704021426 | 0.005690127 | 0.770084382 | 0.953986287 |
| Kctd4 | 0.502526917 | 0.394827922 | 0.701598788 | 0.001095196 | -0.384875448 | 0.993542839 |
| Gm14418 | 0.565657579 | 0.546235958 | 0.699801066 | 0.002887916 | 0.35243165 | 0.993542839 |
| Hist2h2be | 1.018844396 | 0.146693828 | 0.697480621 | 0.045223317 | 0.136624141 | 0.993542839 |
| Ralbp1 | 0.55305896 | 0.108680681 | 0.694533475 | 1.50E-09 | 0.271724135 | 0.993542839 |
| Eif4ebp1 | 0.454403017 | 0.582029579 | 0.693474874 | 0.004018942 | 0.089185206 | 0.993542839 |
| Utp3 | 0.350400889 | 0.336690137 | 0.69278022 | 2.47E-11 | 0.439197369 | 0.993542839 |
| Frmd6 | 0.592832719 | 0.169048164 | 0.690729676 | 0.000719677 | 0.684483551 | 0.857645378 |
| Srgn | 0.384983576 | 0.475938631 | 0.690324077 | 0.001186021 | 0.382154155 | 0.993542839 |
| Impact | 0.380561873 | 0.021182325 | 0.69012376 | 2.71E-14 | 0.126884367 | 0.993542839 |
| Hhatl | 1.000980261 | 0.0513835 | 0.689723033 | 0.012041549 | 0.192843437 | 0.993542839 |
| Zfp850 | 0.033481331 | 0.988884844 | 0.688420726 | 0.038994686 | 0.040762662 | 0.993542839 |
| Car2 | 0.412151426 | 0.06531488 | 0.686872582 | 0.002223457 | 0.445966148 | 0.993542839 |
| Krt12 | 0.452234735 | 0.330417651 | 0.686791568 | 2.82E-07 | -0.238801414 | 0.993542839 |
| Pcgf5 | 0.710318494 | 0.232035676 | 0.686628042 | 6.20E-05 | 0.406235142 | 0.993542839 |
| Khdrbs2 | -0.516782181 | 0.658352605 | 0.685569371 | 0.0056221 | -0.220499409 | 0.993542839 |
| Atf4 | 0.345254447 | 0.157513442 | 0.684905432 | 3.86E-13 | -0.028875481 | 0.993542839 |
| Rsl1 | 0.133561391 | 0.954804506 | 0.684270842 | 0.043289728 | 0.198675744 | 0.993542839 |
| Slc35g2 | 0.458707652 | 0.52565529 | 0.683761837 | 0.000235228 | 0.292458768 | 0.993542839 |
| Twistnb | 0.44470304 | 0.579837411 | 0.683009878 | 0.000246682 | 0.15565378 | 0.993542839 |
| Gm9989 | 0.500940385 | 0.331423618 | 0.682939467 | 0.017692676 | 0.618954755 | 0.90514278 |
| Gpr62 | 0.166598449 | 0.887812837 | 0.681844592 | 0.002021082 | 0.815800883 | 0.506204991 |
| Lats2 | 0.385445826 | 0.405782739 | 0.681170285 | 0.000421608 | 0.071175582 | 0.993542839 |
| Tnfaip3 | 0.01859314 | 0.992324297 | 0.681141961 | 0.049331508 | -0.13074727 | 0.993542839 |
| Med30 | 0.397059902 | 0.277017107 | 0.680972263 | 6.69E-12 | 0.303009104 | 0.993542839 |
| Prkg2 | 0.307098079 | 0.812183648 | 0.677754823 | 0.013333803 | -0.204314311 | 0.993542839 |
| 2010001A14Rik | 1.05994659 | 0.150516444 | 0.676623182 | 0.003335749 | 0.621606306 | 0.967845527 |
| Tceal9 | 0.280450968 | 0.205870017 | 0.675913308 | 4.43E-13 | 0.324409102 | 0.993542839 |
| Samd4 | 0.71842864 | 0.134092936 | 0.675461826 | 0.000818416 | 0.353327478 | 0.993542839 |
| Dkc1 | 0.294114667 | 0.588694205 | 0.674616343 | 0.000107233 | 0.30356778 | 0.993542839 |
| Ehd4 | 0.628589213 | 0.346568111 | 0.674483931 | 0.00196661 | 0.66089461 | 0.862821352 |
| Cbr3 | 0.964017219 | 0.059537991 | 0.672988663 | 0.047240192 | 0.473228916 | 0.993542839 |
| Klhl15 | 0.490285988 | 0.525706298 | 0.672346848 | 0.004438282 | 0.125642074 | 0.993542839 |
| Pabpn1 | 0.592899036 | 0.217859512 | 0.670458598 | 0.000805945 | 0.561072675 | 0.993542839 |
| Amd2 | -0.018129085 | 0.991965781 | 0.667841145 | 0.035504091 | 0.309497903 | 0.993542839 |
| Cast | 1.020122327 | 0.076064261 | 0.667377024 | 0.006815056 | 0.214629338 | 0.993542839 |
| Clp1 | 0.094277956 | 0.944670088 | 0.665678089 | 0.007974776 | 0.133987255 | 0.993542839 |
| Arhgef6 | 0.63200846 | 0.32063889 | 0.66365799 | 0.016088712 | 0.028424093 | 0.993542839 |
| H2-T23 | 0.826316413 | 0.084995806 | 0.663049548 | 0.012281025 | 0.779064213 | 0.566630167 |
| Fbxo33 | 0.472666428 | 0.172645291 | 0.662300611 | 8.15E-05 | 0.353844151 | 0.993542839 |
| Kiz | 0.088591704 | 0.936394856 | 0.662056291 | 0.001309827 | 0.209295668 | 0.993542839 |
| Dusp23 | 0.098728455 | 0.932676755 | 0.661498777 | 0.00832968 | 0.052741764 | 0.993542839 |
| Flywch1 | 0.495302857 | 0.014595659 | 0.661394832 | 4.89E-05 | 0.325483728 | 0.993542839 |
| Hscb | 0.32378219 | 0.625266764 | 0.65996212 | 0.000118972 | -0.019149462 | 0.993542839 |
| Rbm4b | 0.505776716 | 0.393208925 | 0.656972943 | 0.000176064 | 0.507125592 | 0.993542839 |
| Rnf6 | 0.505660217 | 0.114497021 | 0.655541473 | 2.75E-11 | 0.011157009 | 0.993542839 |
| Iqcb1 | 0.370141187 | 0.450391009 | 0.65330626 | 3.94E-07 | 0.388853215 | 0.993542839 |
| Rpl5 | 0.639731669 | 0.059125025 | 0.65274868 | 0.000574776 | 0.375720129 | 0.993542839 |
| D16Ertd472e | 0.551740533 | 0.448512749 | 0.652060802 | 0.000214158 | 0.019762375 | 0.993542839 |
| Il17ra | 0.455475883 | 0.463881863 | 0.651006 | 0.028025468 | 0.152460489 | 0.993542839 |
| Zfp51 | -0.042357905 | 0.983525033 | 0.650016653 | 0.038531365 | -0.065580983 | 0.993542839 |
| Epdr1 | 0.466464531 | 0.018404008 | 0.64915375 | 1.33E-05 | 0.210149889 | 0.993542839 |
| Frmd8 | 0.223754 | 0.736465621 | 0.648231802 | 0.002147847 | -0.094382914 | 0.993542839 |
| Zfp941 | 0.482647635 | 0.557237821 | 0.647227277 | 0.000551358 | -0.181585014 | 0.993542839 |
| Gnai1 | 0.287992457 | 0.241998742 | 0.647101327 | 1.35E-07 | 0.038275195 | 0.993542839 |
| Myrf | 1.087103962 | 0.043049037 | 0.645432839 | 0.010080063 | 0.596200341 | 0.993542839 |
| Cep76 | 0.585517486 | 0.534017055 | 0.644432884 | 0.043237135 | -0.450454709 | 0.993542839 |

| | | | | | | |
|---|---|---|---|---|---|---|
| Fnbp1 | 0.353369439 | 0.311385058 | 0.642187638 | 2.99E-06 | 0.095985722 | 0.993542839 |
| Efr3a | 0.436133666 | 0.147724528 | 0.641790305 | 0.002572712 | 0.049017826 | 0.993542839 |
| Nipsnap3b | 0.27169168 | 0.676136796 | 0.638942848 | 8.65E-05 | -0.351033004 | 0.993542839 |
| Hsd17b11 | 0.356671708 | 0.321718441 | 0.638239802 | 1.55E-06 | -0.247572543 | 0.993542839 |
| Bmi1 | 0.458421523 | 0.226060441 | 0.637434345 | 3.58E-06 | 0.084585325 | 0.993542839 |
| Bex1 | 0.466488318 | 0.019797667 | 0.637304772 | 2.54E-12 | 0.106293734 | 0.993542839 |
| Tmem184c | 0.310647715 | 0.329246633 | 0.637193175 | 1.74E-08 | -0.104272461 | 0.993542839 |
| Pip4k2a | 0.451213539 | 0.141554382 | 0.635653825 | 5.56E-06 | 0.275375572 | 0.993542839 |
| Nrsn2 | 0.590167908 | 0.059577774 | 0.634723199 | 0.001838556 | 0.136577606 | 0.993542839 |
| Pcdhb17 | 0.604743511 | 0.288144076 | 0.633028606 | 0.00140961 | -0.550605218 | 0.993542839 |
| Cdr2 | 0.515478082 | 0.687931169 | 0.632846557 | 0.04377811 | -0.057881963 | 0.993542839 |
| Atp6v1h | 0.494656797 | 0.009199598 | 0.631919081 | 1.97E-09 | 0.065962369 | 0.993542839 |
| Them6 | 0.190121328 | 0.785765892 | 0.629830092 | 0.000325306 | -0.130874548 | 0.993542839 |
| Zfp729b | 0.199548799 | 0.857805193 | 0.629607091 | 0.005469246 | 0.242611471 | 0.993542839 |
| Vamp5 | 0.152932163 | 0.870662146 | 0.627846821 | 0.04615627 | 0.397396687 | 0.993542839 |
| Cd83 | 0.378020144 | 0.68315399 | 0.627368455 | 0.013791492 | -0.041062431 | 0.993542839 |
| Trem2 | 0.510020346 | 0.080891033 | 0.626850914 | 5.03E-05 | 0.735025982 | 0.875404481 |
| Zfp825 | 0.826097118 | 0.27137124 | 0.625913383 | 0.007399142 | 0.202107114 | 0.993542839 |
| Abt1 | 0.701212046 | 0.158818911 | 0.625486191 | 0.001647426 | 0.140385974 | 0.993542839 |
| Actn3 | 0.457445813 | 0.796829229 | 0.624536057 | 0.048015399 | -0.960570649 | 0.632281631 |
| Cd37 | 0.476520636 | 0.681727618 | 0.624296616 | 0.042823976 | 0.352141235 | 0.993542839 |
| Lonrf1 | 0.337828142 | 0.681727618 | 0.624070287 | 0.002213839 | 0.576757085 | 0.993542839 |
| Cpne4 | 0.418149368 | 0.404051469 | 0.624015235 | 0.000708947 | -0.480887862 | 0.993542839 |
| Acot13 | 0.473523669 | 0.001145993 | 0.623503183 | 6.57E-10 | 0.18206432 | 0.993542839 |
| Cyba | 0.422405262 | 0.366225365 | 0.623092881 | 0.011525979 | 0.668067291 | 0.993542839 |
| Mcub | 0.293219208 | 0.756426573 | 0.622977754 | 0.003187462 | -0.124302078 | 0.993542839 |
| Etv1 | 0.300450398 | 0.448512749 | 0.622164995 | 0.009760277 | -0.215171178 | 0.993542839 |
| Gcat | 0.373836199 | 0.458704493 | 0.620598473 | 0.000253772 | 0.250168869 | 0.993542839 |
| Tpd52l1 | -0.165129072 | 0.763792079 | 0.620130687 | 0.000349895 | 0.864275791 | 0.736141252 |
| Gskip | 0.235683021 | 0.58627475 | 0.619832274 | 0.000124551 | 0.114379087 | 0.993542839 |
| Sox2ot | 0.6832219 | 0.324445661 | 0.619446631 | 0.009510589 | 0.452244331 | 0.993542839 |
| Paip1 | 0.478317129 | 0.145503398 | 0.618899543 | 8.05E-05 | 0.280560634 | 0.993542839 |
| Gad1 | 0.413775472 | 0.092990896 | 0.61705404 | 3.16E-06 | 0.102794382 | 0.993542839 |
| Psmd1 | 0.600446051 | 0.080672292 | 0.616769346 | 0.018254986 | 0.17892361 | 0.993542839 |
| Tmcc3 | 0.758786208 | 0.07823325 | 0.616330725 | 0.000486104 | -0.061456033 | 0.993542839 |
| Arhgap10 | 0.19474222 | 0.901362498 | 0.616183549 | 0.016143192 | 0.128646835 | 0.993542839 |
| Supt16 | 0.052493211 | 0.951345014 | 0.614538867 | 3.75E-05 | 0.161762843 | 0.993542839 |
| Ppl | 0.5238236 | NA | 0.614482049 | 0.049863444 | 1.22781162 | 0.478755413 |
| Slc5a3 | 0.261610077 | 0.751551971 | 0.613013424 | 0.012294614 | 0.498159359 | 0.993542839 |
| Fgfbp3 | 0.964171164 | 0.050127258 | 0.61066532 | 0.027189921 | 0.079029468 | 0.993542839 |
| Crebrf | 0.647624555 | 0.06806501 | 0.609388239 | 0.00094675 | 0.39349473 | 0.993542839 |
| Erbin | 0.467388256 | 0.172645291 | 0.609328859 | 0.00072673 | 0.778174909 | 0.589960018 |
| Stat6 | 0.996383698 | 0.100122283 | 0.60832681 | 0.024707867 | 0.101456935 | 0.993542839 |
| Rbm12b1 | 0.385265059 | 0.684963286 | 0.608244111 | 0.013110027 | 0.12846481 | 0.993542839 |
| Jade1 | 0.595570567 | 0.163634238 | 0.60748476 | 0.000568533 | -0.142193797 | 0.993542839 |
| Nup43 | -0.363316883 | 0.76310176 | 0.60689316 | 0.036470321 | -0.096240466 | 0.993542839 |
| Cmtm5 | 0.462866203 | 0.035574291 | 0.606752177 | 0.000200615 | 0.587845673 | 0.993542839 |
| Tob1 | 0.575712638 | 0.075021429 | 0.605694495 | 2.72E-05 | 0.135704408 | 0.993542839 |
| Usf1 | 0.387384617 | 0.526121093 | 0.605524388 | 0.013092496 | 0.291440027 | 0.993542839 |
| Plek | 0.516620218 | 0.538308925 | 0.605155889 | 0.015616426 | 0.386832802 | 0.993542839 |
| Ephx2 | 0.433885651 | 0.282758891 | 0.60476691 | 0.00024977 | 0.17883605 | 0.993542839 |
| Necab1 | 0.299326876 | 0.253875803 | 0.602077264 | 3.24E-09 | -0.417839997 | 0.993542839 |
| Klf5 | 0.611189721 | 0.189904321 | 0.600815374 | 0.000111759 | -0.16390223 | 0.993542839 |
| Gm14295 | 0.731392441 | 0.158496999 | 0.599897704 | 0.001709659 | -0.102924556 | 0.993542839 |
| Grhpr | 0.374954046 | 0.362083288 | 0.598455518 | 0.002362816 | 0.216413327 | 0.993542839 |
| Polr2k | 0.334929665 | 0.086920501 | 0.597458434 | 2.14E-08 | 0.287681444 | 0.993542839 |
| Amotl1 | 0.675397659 | 0.337476518 | 0.597019781 | 0.005912135 | 0.250902527 | 0.993542839 |
| 1700048O20Rik | 0.201117953 | 0.899196003 | 0.595102403 | 0.021177466 | 0.353174321 | 0.993542839 |
| Tspyl2 | 0.580240542 | 0.211245314 | 0.594616629 | 2.50E-05 | 0.079657239 | 0.993542839 |
| Htr5a | 0.16185474 | 0.890135176 | 0.594368272 | 0.000735252 | 0.131725011 | 0.993542839 |
| Incenp | 0.43380165 | 0.386286228 | 0.594171318 | 0.007522427 | -0.53353947 | 0.95811614 |
| Lrif1 | 0.190933891 | 0.844876471 | 0.593559398 | 0.000473877 | 0.228720877 | 0.993542839 |
| Papss2 | 0.681826438 | 0.341186892 | 0.592733137 | 0.028272496 | 0.165901938 | 0.993542839 |
| Arhgap27os2 | -0.105965616 | 0.944743842 | 0.592597085 | 0.008219503 | -0.016549974 | 0.993542839 |
| Fgfrl1 | 0.257775649 | 0.721246706 | 0.592214311 | 0.00817839 | 0.099739161 | 0.993542839 |
| Clic4 | 0.322886848 | 0.433877458 | 0.59026756 | 0.000659625 | 0.350728207 | 0.993542839 |
| Fn3k | 0.518583606 | 0.132849902 | 0.586678815 | 1.57E-05 | 0.2186235 | 0.993542839 |
| Isoc1 | 0.32028053 | 0.560068322 | 0.58652105 | 0.000111759 | -0.273522403 | 0.993542839 |
| Cog7 | 0.277729578 | 0.609395843 | 0.583188876 | 4.91E-05 | 0.113588739 | 0.993542839 |
| Wipf1 | 0.441963617 | 0.489548733 | 0.582689162 | 0.017173071 | -0.172529176 | 0.993542839 |
| Crebl2 | 0.691134503 | 0.11989504 | 0.582245056 | 0.013340657 | 0.226787798 | 0.993542839 |
| Hrh1 | 0.373764986 | 0.634553005 | 0.581231989 | 0.00058187 | 0.010280565 | 0.993542839 |
| Sh3bp1 | 0.408759926 | 0.637248044 | 0.580883514 | 0.038918212 | 0.141418052 | 0.993542839 |

| | | | | | | |
|---|---|---|---|---|---|---|
| AI987944 | 0.562578505 | 0.561953709 | 0.579413663 | 0.038617172 | -0.070936141 | 0.993542839 |
| Fbxo32 | 0.807076609 | 0.308151219 | 0.578516884 | 0.046328048 | 0.593082126 | 0.993542839 |
| Arl4d | 0.309461959 | 0.626002596 | 0.578413864 | 0.00540765 | 0.159843891 | 0.993542839 |
| Plcl1 | 0.099766196 | 0.917890733 | 0.578249055 | 6.08E-05 | 0.264943764 | 0.993542839 |
| Carhsp1 | 0.426699599 | 0.26493801 | 0.578170522 | 0.010159728 | 0.181903161 | 0.993542839 |
| Dpp9 | 0.26924303 | 0.673280843 | 0.57705917 | 0.009495177 | 0.161247972 | 0.993542839 |
| Zhx1 | 0.562003724 | 0.051439492 | 0.576118686 | 1.01E-07 | 0.15483502 | 0.993542839 |
| Dbt | 0.407831192 | 0.471744471 | 0.576065411 | 0.010754517 | -0.027129057 | 0.993542839 |
| Epm2a | 0.536142128 | 0.321605792 | 0.574380891 | 0.000498311 | -0.105611462 | 0.993542839 |
| Dusp28 | 0.557063995 | 0.200911893 | 0.572670968 | 0.00839982 | 0.267964857 | 0.993542839 |
| Morf4l1 | 0.403360899 | 7.06E-05 | 0.571486858 | 2.13E-14 | 0.214678681 | 0.993542839 |
| Nop58 | 0.495004673 | 0.298150604 | 0.571243955 | 6.71E-05 | 0.13288757 | 0.993542839 |
| Scly | 0.720958571 | 0.266452528 | 0.570602781 | 0.005413847 | -0.116779122 | 0.993542839 |
| Hes1 | 0.182532372 | 0.808765287 | 0.570419391 | 0.011104718 | 0.297782074 | 0.993542839 |
| Msmo1 | 0.320455521 | 0.158496999 | 0.569420671 | 6.30E-09 | -0.07727264 | 0.993542839 |
| Stx4a | 0.382991888 | 0.224230817 | 0.569397988 | 5.41E-08 | 0.30921328 | 0.993542839 |
| Usp5 | 0.329942736 | 0.059577774 | 0.567874909 | 7.80E-13 | 0.092188042 | 0.993542839 |
| Ucma | -0.27387116 | 0.843788145 | 0.567391385 | 0.047216407 | -0.873422156 | 0.655539463 |
| Etv3 | -0.144268474 | 0.879421871 | 0.56646378 | 0.000910732 | 0.231983832 | 0.993542839 |
| Zmat4 | 0.247271568 | 0.774311324 | 0.565917119 | 0.024967269 | 0.285250442 | 0.993542839 |
| Dnajb4 | 0.504253989 | 0.097510766 | 0.565694341 | 0.000200958 | 0.236963596 | 0.993542839 |
| Plekhb2 | 0.418948389 | 0.001945581 | 0.563663239 | 1.06E-17 | 0.182538979 | 0.993542839 |
| Aif1 | 0.687242446 | 0.064041509 | 0.563635881 | 0.001647488 | 0.432779553 | 0.993542839 |
| Pfdn2 | 0.480798933 | 0.018493635 | 0.56205138 | 4.78E-05 | 0.210516171 | 0.993542839 |
| Prcd | 0.084949265 | 0.952189316 | 0.561748132 | 0.029704687 | -0.371401329 | 0.993542839 |
| Ctxn2 | 0.553550296 | 0.12012772 | 0.561570263 | 0.000908817 | 0.006611107 | 0.993542839 |
| Cish | -0.197401343 | 0.880836892 | 0.559755586 | 0.029429589 | 0.248345248 | 0.993542839 |
| Otud1 | 0.447879597 | 0.476181974 | 0.559531716 | 0.003384621 | -0.172352562 | 0.993542839 |
| Isca1 | 0.415338453 | 0.033608198 | 0.559154566 | 2.15E-06 | 0.105189059 | 0.993542839 |
| M6pr | 0.470891692 | 0.028494451 | 0.558621288 | 5.01E-11 | 0.216247945 | 0.993542839 |
| Set | 0.479593398 | 0.008669366 | 0.5575114 | 2.14E-09 | 0.196084602 | 0.993542839 |
| Prcp | -0.07843645 | 0.94010614 | 0.556659496 | 0.003780325 | 0.070129677 | 0.993542839 |
| Amd1 | 0.75267461 | 0.149423768 | 0.556152138 | 0.006999943 | 0.414917273 | 0.993542839 |
| Ublcp1 | 0.26027348 | 0.525706298 | 0.554652581 | 3.95E-08 | 0.001054912 | 0.998000351 |
| Pacsin2 | 0.483793639 | 0.135195576 | 0.554363354 | 0.000223809 | 0.052538188 | 0.993542839 |
| Peli3 | 0.86614648 | 0.110018203 | 0.554340631 | 0.000513866 | 0.030894852 | 0.993542839 |
| Entpd3 | 1.430606411 | 0.09646533 | 0.553265245 | 0.043197109 | 0.375120344 | 0.993542839 |
| Slc16a3 | -0.03346509 | 0.985280397 | 0.553089549 | 0.007330905 | -0.09848719 | 0.993542839 |
| Scrn3 | 0.149410627 | 0.860951108 | 0.551558737 | 0.000451208 | 0.104481042 | 0.993542839 |
| Enpp4 | 0.418281107 | 0.477692707 | 0.551535917 | 0.004066668 | -0.281711732 | 0.993542839 |
| Cpeb1 | 0.437581559 | 0.464082513 | 0.550919804 | 0.018240047 | 0.141901972 | 0.993542839 |
| Ttyh2 | 0.408649195 | 0.525706298 | 0.550820869 | 0.016397891 | 0.310954619 | 0.993542839 |
| Blcap | 0.340616524 | 0.164477058 | 0.550733228 | 4.66E-08 | 0.05582391 | 0.993542839 |
| Fam204a | 0.04076785 | 0.969564186 | 0.549008032 | 0.003754445 | 0.236387444 | 0.993542839 |
| Rsrc1 | 0.347988642 | 0.465658596 | 0.548522423 | 1.64E-06 | 0.178826 | 0.993542839 |
| Rpl35a | 0.487119558 | 0.020087255 | 0.54823136 | 0.000254979 | 0.339429665 | 0.993542839 |
| Zfp65 | 0.540683782 | 0.518621383 | 0.548069701 | 0.011774093 | -0.171419545 | 0.993542839 |
| Fam133b | 0.419614376 | 0.437784835 | 0.546208157 | 0.00034649 | 0.305447366 | 0.993542839 |
| Socs4 | 0.29096748 | 0.699178389 | 0.545145467 | 0.018096785 | 0.342323276 | 0.993542839 |
| Peg13 | 0.45622548 | 0.002607229 | 0.544935755 | 0.00022845 | 0.148390818 | 0.993542839 |
| Ndufb1-ps | 0.37495738 | 0.023154359 | 0.544414205 | 5.34E-06 | 0.192412071 | 0.993542839 |
| Cib2 | 0.548645779 | 0.081461262 | 0.543917938 | 0.000122883 | 0.073968773 | 0.993542839 |
| Nos1 | 0.540726513 | 0.346036027 | 0.543723883 | 0.043257971 | -0.12178535 | 0.993542839 |
| Myh14 | 0.407302915 | 0.309514116 | 0.54326271 | 0.021726153 | 0.149668678 | 0.993542839 |
| Rps3a1 | 0.463619644 | 0.001654585 | 0.541324368 | 3.15E-06 | 0.303577704 | 0.993542839 |
| Zbtb7a | 0.473293247 | 0.219331392 | 0.540050205 | 0.001369702 | 0.028424878 | 0.993542839 |
| Eif2ak3 | 0.447211548 | 0.494766805 | 0.539958811 | 0.012486283 | 0.222590898 | 0.993542839 |
| Flrt1 | 0.067584711 | 0.966529939 | 0.539825833 | 0.015989852 | 0.267626022 | 0.993542839 |
| Gm2415 | 0.874635447 | 0.124034744 | 0.539796505 | 0.02768406 | 0.146310591 | 0.993542839 |
| Rgs2 | 0.283272006 | 0.462740481 | 0.539295627 | 2.86E-05 | -0.023987307 | 0.993542839 |
| Znfx1 | 0.324956235 | 0.497486289 | 0.53869529 | 0.000757004 | 0.321139484 | 0.993542839 |
| Sox8 | 0.14492426 | 0.870772081 | 0.537315084 | 0.032408265 | 0.195747412 | 0.993542839 |
| Sec62 | 0.439279021 | 0.012114379 | 0.535870327 | 8.52E-06 | 0.11411044 | 0.993542839 |
| Rnaset2a | 0.32424109 | 0.430818388 | 0.534508393 | 0.002401789 | 0.012088968 | 0.993542839 |
| Mafk | 0.440476141 | 0.539287654 | 0.53411077 | 0.026293925 | 0.005881712 | 0.993542839 |
| Rnf227 | 0.326511974 | 0.314552791 | 0.533985748 | 0.006764592 | 0.033473273 | 0.993542839 |
| Hspa4l | 0.491245019 | 0.003467891 | 0.533608259 | 4.91E-06 | 0.054265406 | 0.993542839 |
| Eif5b | 0.567319137 | 0.117911541 | 0.532872651 | 0.002682541 | 0.101274896 | 0.993542839 |
| Fdx1 | 0.290713586 | 0.488386892 | 0.532263123 | 3.30E-05 | 0.058308732 | 0.993542839 |
| Gm26881 | 0.511776708 | 0.279612769 | 0.531991706 | 0.000323254 | 0.080024566 | 0.993542839 |
| Mamld1 | 0.331933539 | 0.552178374 | 0.531859389 | 0.00206079 | -0.164165789 | 0.993542839 |
| Map7d2 | 0.311105443 | 0.188656033 | 0.53164382 | 1.96E-10 | 0.05129267 | 0.993542839 |
| Sdf2l1 | 1.156215387 | 0.017421714 | 0.530666848 | 0.027177088 | 0.544078739 | 0.988579962 |

| | | | | | | |
|---|---|---|---|---|---|---|
| E2f1 | 0.031165401 | 0.980074367 | 0.530225213 | 0.002686844 | -0.473772471 | 0.993542839 |
| Stk24 | 0.230636014 | 0.823199538 | 0.528033803 | 0.007538164 | 0.1070868 | 0.993542839 |
| Cry1 | 0.279587219 | 0.647580017 | 0.527072277 | 0.002110718 | 0.383527468 | 0.993542839 |
| Adar | 0.360288599 | 0.135195576 | 0.525297898 | 1.45E-10 | 0.134503058 | 0.993542839 |
| Zkscan14 | 0.302987467 | 0.745438917 | 0.524862575 | 0.019372887 | -0.225824308 | 0.993542839 |
| Ppm1k | 0.180485987 | 0.709847736 | 0.523993195 | 6.34E-06 | 0.120199574 | 0.993542839 |
| Prepl | 0.514231594 | 0.285870977 | 0.523777543 | 0.00014021 | 0.04800721 | 0.993542839 |
| Hlf | 0.331489524 | 0.156715416 | 0.52371192 | 1.40E-05 | 0.316080022 | 0.993542839 |
| Cntfr | 0.4976621 | 0.068274872 | 0.522670537 | 5.28E-06 | -0.173597193 | 0.993542839 |
| Gnpda2 | 0.44925755 | 0.353604273 | 0.522010563 | 0.005912135 | 0.208252742 | 0.993542839 |
| Rab27b | 0.134208906 | 0.869100934 | 0.521901878 | 0.008412169 | -0.214111987 | 0.993542839 |
| Ints12 | 0.907921032 | 0.237382896 | 0.520922577 | 0.014649252 | 0.346206166 | 0.993542839 |
| Pdcd2 | 0.303197256 | 0.62007601 | 0.518335174 | 0.00026797 | 0.070358524 | 0.993542839 |
| Uap1l1 | 0.457954298 | 0.309109824 | 0.517964718 | 0.007538164 | 0.171416201 | 0.993542839 |
| Tmem144 | 0.407664764 | 0.64080298 | 0.517847453 | 0.026131554 | 0.495441902 | 0.993542839 |
| Rps10 | 0.435871891 | 0.006040363 | 0.517410292 | 0.000136757 | 0.252306567 | 0.993542839 |
| Prpsap2 | 0.452514384 | 0.231613104 | 0.517340107 | 2.89E-05 | 0.119610377 | 0.993542839 |
| Phf1 | 0.058255949 | 0.943169213 | 0.51729108 | 0.01552125 | 0.017083001 | 0.993542839 |
| Lrrk2 | 0.598305986 | 0.347014138 | 0.516963656 | 0.005409533 | 0.112890677 | 0.993542839 |
| Ide | 0.643210798 | 0.05096152 | 0.51678809 | 0.031058701 | 0.208497214 | 0.993542839 |
| Rpl14 | 0.458125035 | 0.00099746 | 0.516277676 | 9.10E-05 | 0.240335418 | 0.993542839 |
| Ciart | -0.169343724 | 0.838508115 | 0.515676134 | 0.003463862 | 0.524805998 | 0.993542839 |
| Bmpr1b | 0.276846497 | 0.698470246 | 0.514643921 | 0.04715206 | 0.172764059 | 0.993542839 |
| Mest | -0.13716527 | 0.886646631 | 0.514542231 | 0.009390522 | 0.276593021 | 0.993542839 |
| Snap25 | 0.42454644 | 0.099461411 | 0.514449114 | 0.000643682 | 0.072472693 | 0.993542839 |
| Sertad1 | 0.805316213 | 0.13228519 | 0.513181883 | 0.005775513 | 0.243726034 | 0.993542839 |
| Pcna | 0.41250444 | 0.092119546 | 0.51315127 | 2.12E-09 | 0.050410027 | 0.993542839 |
| G3bp2 | 0.397403288 | 0.010990905 | 0.513031523 | 5.37E-09 | 0.033773982 | 0.993542839 |
| Zwint | 0.458978922 | 0.000168947 | 0.512694977 | 3.40E-11 | 0.071955273 | 0.993542839 |
| Cd68 | 0.302699293 | 0.595833203 | 0.511911459 | 0.003089932 | 0.092462782 | 0.993542839 |
| Ugp2 | 0.478491191 | 0.021419845 | 0.511466038 | 8.40E-08 | -0.011580388 | 0.993542839 |
| Psme2 | 0.357253016 | 0.238276062 | 0.510858042 | 0.000108807 | 0.389883301 | 0.993542839 |
| Spint2 | 0.107748869 | 0.834586265 | 0.510224426 | 3.15E-06 | -0.147925694 | 0.993542839 |
| Kcna6 | 0.289452436 | 0.43943762 | 0.510079761 | 6.45E-05 | -0.064023809 | 0.993542839 |
| Ndufb3 | 0.390892529 | 0.002903113 | 0.509374878 | 1.74E-11 | 0.085208375 | 0.993542839 |
| Nmral1 | 0.215797569 | 0.802103476 | 0.50923033 | 0.006665048 | 0.244001358 | 0.993542839 |
| Ldlr | 0.387805713 | 0.566132935 | 0.507990863 | 0.033782096 | -0.592537446 | 0.95811614 |
| Adrb1 | 0.656256821 | 0.140687301 | 0.507319329 | 0.035701035 | -0.137257007 | 0.993542839 |
| Tmem192 | 0.355550438 | 0.43075221 | 0.507055816 | 0.000725922 | 0.417421106 | 0.993542839 |
| Ccdc91 | 0.385505548 | 0.462740481 | 0.506472669 | 0.000252452 | -0.200943817 | 0.993542839 |
| Kif16b | 0.968209179 | 0.12041738 | 0.505793217 | 0.021516394 | -0.42338835 | 0.993542839 |
| Stx8 | 0.298006291 | 0.461814095 | 0.505200807 | 3.04E-05 | -0.274711694 | 0.993542839 |
| Ankrd45 | 0.641952688 | 0.135195576 | 0.503988793 | 0.013699627 | -0.136030312 | 0.993542839 |
| Ppif | 0.625495803 | 0.096172196 | 0.503958295 | 0.000146713 | -0.327272392 | 0.993542839 |
| Slirp | 0.480211394 | 0.006735929 | 0.503768256 | 1.52E-06 | 0.072075572 | 0.993542839 |
| Ifitm10 | 0.418302136 | 0.238106423 | 0.50364841 | 0.03001631 | -0.091881357 | 0.993542839 |
| Hprt | 0.433180738 | 0.017481945 | 0.503647372 | 9.84E-14 | 0.143163949 | 0.993542839 |
| Gtf2ird2 | 0.29227533 | 0.749023906 | 0.502971672 | 0.006933322 | -0.004094552 | 0.993542839 |
| Fen1 | -0.113166145 | 0.931578253 | 0.502534357 | 0.008808304 | 0.019432846 | 0.993542839 |
| Adi1 | 0.226716123 | 0.691733594 | 0.502473914 | 0.002072665 | 0.377807567 | 0.993542839 |
| Cln5 | 0.206950937 | 0.779248455 | 0.502468255 | 0.035561395 | 0.122867524 | 0.993542839 |
| Cdkn2aip | 1.469719532 | 0.011701268 | 0.502319019 | 0.007585397 | 0.268382487 | 0.993542839 |
| Akap12 | 0.154976328 | 0.816137512 | 0.502268214 | 0.038531365 | -0.136981402 | 0.993542839 |
| Fdft1 | 0.393618106 | 0.081117038 | 0.50222126 | 1.91E-11 | -0.02761107 | 0.993542839 |
| Topors | 0.212062909 | 0.805558369 | 0.502202043 | 0.021726153 | 0.235686319 | 0.993542839 |
| Yrdc | 0.050205608 | 0.944670088 | 0.502198044 | 0.00097339 | 0.250916051 | 0.993542839 |
| Sacm1l | 0.274555257 | 0.336690137 | 0.501352654 | 2.03E-07 | -0.004524829 | 0.993542839 |
| Txndc5 | 0.470175942 | 0.154063408 | 0.501207002 | 0.000223089 | 0.024826717 | 0.993542839 |
| Cnep1r1 | 0.262284578 | 0.675218894 | 0.501089142 | 0.001267713 | 0.123914489 | 0.993542839 |
| Cmc1 | 0.251227304 | 0.636065822 | 0.500754742 | 0.000224045 | 0.278304498 | 0.993542839 |
| Manea | 0.39337608 | 0.399163305 | 0.500297341 | 0.000127697 | -0.018030592 | 0.993542839 |
| Atic | 0.587849985 | 0.241023082 | 0.500262723 | 0.001682881 | -0.253181278 | 0.993542839 |
| Sox6 | -0.462053654 | 0.529778106 | -0.500149506 | 0.045958031 | -1.049075424 | 0.365409496 |
| Oxct1 | -0.412279834 | 0.00458145 | -0.500984465 | 0.000256588 | -1.024347908 | 0.69988617 |
| Mcat | -0.029381729 | 0.974814223 | -0.501120716 | 0.000117983 | -0.705690858 | 0.848996673 |
| Matk | -0.375617929 | 0.063816695 | -0.501121817 | 0.00079839 | -0.966341057 | 0.721979185 |
| Ddah1 | 0.0323457 | 0.951480269 | -0.501722306 | 7.87E-05 | -1.084673539 | 0.3992986 |
| Slc31a1 | -0.72905981 | 0.052867019 | -0.502599777 | 0.014031152 | -1.150284856 | 0.171380409 |
| Slc6a7 | -0.523239566 | 0.170436132 | -0.503065717 | 0.000175138 | -1.191694824 | 0.391744774 |
| Zhx3 | -0.286734365 | 0.431024195 | -0.503733072 | 0.000163412 | -0.758379336 | 0.76110548 |
| Kcnmb4 | -0.277044753 | 0.326308631 | -0.503851617 | 1.84E-06 | -0.525559247 | 0.993542839 |
| Ppfia4 | -0.179172169 | 0.646606502 | -0.504517343 | 0.002323632 | -1.026790104 | 0.39700007 |
| Celsr3 | -0.858842192 | 0.056595078 | -0.505075689 | 0.019597568 | -0.931045323 | 0.547497666 |

| | | | | | | |
|---|---|---|---|---|---|---|
| Panx1 | -0.690891914 | 0.206671409 | -0.505472887 | 0.002659213 | -0.789668404 | 0.719791239 |
| Zfc3h1 | -0.793516599 | 0.07083883 | -0.505770358 | 0.001616405 | -1.00292268 | 0.301943111 |
| Zdhhc8 | -0.376234118 | 0.203325849 | -0.506154464 | 8.20E-05 | -1.010055719 | 0.522741515 |
| Zfp629 | -0.218127727 | 0.775782617 | -0.506573741 | 0.003094499 | -1.047445898 | 0.162913907 |
| Ccn2 | 0.121452723 | 0.876554129 | -0.507622905 | 0.013789642 | 0.026621608 | 0.993542839 |
| Tmem240 | -0.134964637 | 0.795795348 | -0.50780492 | 1.24E-06 | -0.79458417 | 0.742359833 |
| Wdr83 | -0.65309663 | 0.058856564 | -0.508111747 | 0.001001917 | -0.926162087 | 0.464044822 |
| Dcaf15 | -0.791825994 | 0.107913928 | -0.508909833 | 0.002551312 | -0.81046115 | 0.55026974 |
| B3gat1 | -0.397823607 | 0.037636847 | -0.509089933 | 6.51E-06 | -0.997674657 | 0.706091083 |
| Usp28 | -0.599002923 | 0.258593134 | -0.509237071 | 0.004653459 | -0.727638347 | 0.708926839 |
| Grip1 | -0.827497113 | 0.273469952 | -0.509274327 | 0.019903614 | -1.278480049 | 0.074569768 |
| Diras2 | -0.232622496 | 0.330643785 | -0.50938582 | 0.001818538 | -0.603719227 | 0.993542839 |
| Cebpd | 0.067709063 | 0.944670088 | -0.51041044 | 0.012411411 | -0.666959467 | 0.800099317 |
| Abhd5 | -0.584428378 | 0.229114056 | -0.510889969 | 0.021787597 | -0.984396678 | 0.184983612 |
| Fut11 | -0.459950752 | 0.378952702 | -0.511194128 | 0.001495826 | -0.638829451 | 0.90514278 |
| Iba57 | -0.333738366 | 0.788222497 | -0.511369173 | 0.015294998 | -0.637637296 | 0.721979185 |
| Farsa | -0.272331748 | 0.522245112 | -0.511476454 | 5.12E-05 | -0.901694391 | 0.551475793 |
| 1700086O06Rik | -0.406268504 | 0.502313743 | -0.511565092 | 0.002283299 | -0.591641443 | 0.929751641 |
| Akap13 | 0.417155581 | 0.523898825 | -0.51241579 | 0.044066754 | -0.978111589 | 0.35570703 |
| Srrm4 | -0.069914283 | 0.941308731 | -0.513335965 | 0.007093719 | -0.463186552 | 0.993542839 |
| Pxn | 0.028932942 | 0.97448751 | -0.513817305 | 0.009468798 | -0.922293322 | 0.429584827 |
| Cacna1e | -0.464233386 | 0.045366686 | -0.514650523 | 9.97E-06 | -0.777953119 | 0.891074892 |
| Sympk | -0.338338773 | 0.179382118 | -0.514808415 | 8.12E-08 | -0.776350105 | 0.796056265 |
| Trim9 | -0.424397695 | 0.008870156 | -0.515299025 | 5.93E-07 | -1.05304124 | 0.57790817 |
| Pomt2 | -0.554644773 | 0.303198791 | -0.515404635 | 0.004244206 | -1.059803062 | 0.304838758 |
| Mmp17 | -0.32285602 | 0.477692707 | -0.515441293 | 8.07E-05 | -0.700058279 | 0.993542839 |
| Dlgap3 | -0.104139544 | 0.740701801 | -0.51674909 | 2.12E-06 | -0.86097103 | 0.845916653 |
| Sh2b2 | -0.641051186 | 0.43075221 | -0.51699965 | 0.02985136 | -1.206600079 | 0.088407895 |
| Crmp1 | -0.566164019 | 0.082966831 | -0.517682139 | 0.00340039 | -1.070085325 | 0.650953551 |
| Grik1 | -0.438568911 | 0.505206027 | -0.518591206 | 0.012170405 | -1.304336981 | 0.293558199 |
| Gabrg3 | -0.670568566 | 0.295924058 | -0.518784151 | 0.042987408 | -1.156485849 | 0.184983612 |
| Cdh13 | -0.286590961 | 0.41958044 | -0.51916623 | 0.001976307 | -0.975746154 | 0.655539463 |
| Zfp950 | -0.675575111 | 0.119099878 | -0.519416596 | 0.00432723 | -0.419674645 | 0.993542839 |
| Btbd17 | -0.512203202 | 0.09276842 | -0.519964015 | 0.000435089 | -1.041369171 | 0.179469557 |
| Ctnnbip1 | -0.389421551 | 0.1760274 | -0.520001902 | 3.67E-05 | -0.873149312 | 0.587228117 |
| Rapgef1 | -0.261874472 | 0.240958333 | -0.520040729 | 3.19E-06 | -0.892959499 | 0.558777082 |
| Hif1an | -0.218906698 | 0.796549396 | -0.520347648 | 0.001495005 | -0.799844395 | 0.551429985 |
| Trmt1 | -0.122475663 | 0.854226724 | -0.520351386 | 0.000208517 | -0.839649333 | 0.675272363 |
| Tmem63b | -0.435252145 | 0.068274872 | -0.520990713 | 6.15E-06 | -0.988763612 | 0.587228117 |
| Dancr | -0.82670071 | 0.127103111 | -0.521057154 | 0.027136582 | -0.79204778 | 0.708926839 |
| Slco2b1 | -0.112910949 | 0.882004155 | -0.521960769 | 0.010421219 | -0.914097922 | 0.43417881 |
| Csf2ra | 0.350924028 | 0.603586099 | -0.522138036 | 0.001393612 | -0.673950905 | 0.914497701 |
| Kcnk9 | -0.582436853 | 0.170508375 | -0.522326959 | 0.038181763 | -0.784646264 | 0.831110111 |
| Crlf1 | 0.197461095 | 0.835234864 | -0.522450718 | 0.009289282 | -0.929356068 | 0.401745529 |
| Tmem39b | 0.179877178 | 0.887812837 | -0.522460052 | 0.022151706 | -0.731249876 | 0.683865412 |
| Ccdc50 | -0.43275888 | 0.246971017 | -0.522588461 | 2.40E-05 | -0.717360751 | 0.760591695 |
| Kank1 | 0.088789729 | 0.925610978 | -0.523067996 | 0.040046432 | -1.090263811 | 0.319970507 |
| Prss12 | -0.770163109 | 0.322799071 | -0.523129927 | 0.013441057 | -0.49951785 | 0.993542839 |
| Rab36 | -0.607956772 | 0.240526789 | -0.523717643 | 0.000947045 | -0.704463373 | 0.914497701 |
| Klhl5 | -0.072057791 | 0.918800529 | -0.523770784 | 0.002109657 | -0.832645641 | 0.622707517 |
| Tbata | -0.350935972 | 0.680193721 | -0.524000669 | 0.030162228 | -1.301085474 | 0.08440946 |
| Galnt14 | -0.565200681 | 0.45295527 | -0.524015184 | 0.012292931 | -0.412750938 | 0.993542839 |
| Ptprg | -0.424432676 | 0.225175112 | -0.524319719 | 0.000312667 | -0.832163772 | 0.712523443 |
| Gm13446 | 0.228915806 | 0.870221162 | -0.527201985 | 0.005040945 | -0.135533572 | 0.993542839 |
| Eml6 | -0.472001355 | 0.384535769 | -0.527417242 | 0.014130623 | -0.988817493 | 0.424716939 |
| Tgfb1i1 | -0.342359789 | 0.634541386 | -0.527440444 | 0.010161102 | -0.893661366 | 0.53005075 |
| Ttll12 | -0.500085818 | 0.378710324 | -0.528164427 | 0.000443609 | -0.985681707 | 0.345024541 |
| Szt2 | -0.072219817 | 0.938818689 | -0.528962373 | 0.003139113 | -0.694181293 | 0.864636187 |
| Cep131 | -0.363331986 | 0.566653226 | -0.529553232 | 0.000637665 | -0.74556991 | 0.702044135 |
| Phka1 | -0.633245334 | 0.10914466 | -0.529927043 | 0.002782424 | -0.725843035 | 0.645542085 |
| Ajm1 | -0.251091567 | 0.377449745 | -0.529973333 | 1.22E-08 | -0.952346672 | 0.517245429 |
| Guf1 | -0.617150334 | 0.219915429 | -0.530065728 | 0.019264866 | -1.099636615 | 0.187313119 |
| P2ry12 | -0.434574909 | 0.247224763 | -0.530100582 | 0.004823061 | -0.81150642 | 0.708926839 |
| Hexdc | -0.298750903 | 0.641965729 | -0.530485761 | 0.00074005 | -1.116155028 | 0.184651535 |
| D630023F18Rik | -1.117137931 | 0.015044565 | -0.530611158 | 0.003583196 | -0.939833467 | 0.439842065 |
| Abtb2 | -0.25464397 | 0.77607324 | -0.530849541 | 0.00869371 | -0.580045381 | 0.905658464 |
| Rgma | -0.086242875 | 0.878420271 | -0.531412278 | 0.036075307 | -0.975502393 | 0.733580131 |
| Mfsd10 | -0.451190765 | 0.459997479 | -0.531730141 | 0.005258645 | -0.738445705 | 0.790782897 |
| Rnf112 | -0.334222032 | 0.252356789 | -0.534425737 | 0.000402568 | -0.838348696 | 0.969841518 |
| 2300009A05Rik | -0.150500873 | 0.699191461 | -0.534709176 | 7.90E-07 | -0.680254133 | 0.969213241 |
| Oxtr | -0.889869172 | 0.502436934 | -0.534881254 | 0.01560487 | -0.925129858 | 0.414591828 |
| Rasal1 | -0.385919799 | 0.069174002 | -0.535935812 | 1.42E-09 | -0.568454531 | 0.993542839 |
| Cgref1 | -0.643300054 | 0.219982792 | -0.536283773 | 6.30E-05 | -1.034392511 | 0.341519895 |

| | | | | | | |
|---|---|---|---|---|---|---|
| Ndst3 | -0.944532334 | 0.073363006 | -0.536451822 | 0.024346226 | -1.100444184 | 0.407965996 |
| Tmem38a | -0.474845374 | 0.045283476 | -0.536454717 | 1.26E-13 | -0.751149304 | 0.980362413 |
| Camkk2 | -0.380048779 | 0.049191895 | -0.536457903 | 3.75E-10 | -0.806202926 | 0.845001885 |
| Slc41a2 | -0.033100184 | 0.981092457 | -0.536479239 | 0.021433045 | -1.079578872 | 0.187313119 |
| Rapgef6 | -0.226606988 | 0.788222497 | -0.536586192 | 0.002926249 | -0.686127177 | 0.753331908 |
| Mief2 | -0.047718979 | 0.964787595 | -0.536728669 | 0.000148122 | -1.045973452 | 0.276420848 |
| Pheta1 | -1.200422105 | 0.009726155 | -0.536802637 | 0.001058091 | -0.930759131 | 0.344967761 |
| Ankrd10 | -0.391315903 | 0.297794926 | -0.537229473 | 0.001581997 | -1.115551688 | 0.282442752 |
| Cdo1 | -0.481928577 | 0.247782387 | -0.537368634 | 0.000482395 | -0.697901533 | 0.853703438 |
| Epha3 | -0.948849331 | 0.088101727 | -0.537521695 | 0.013694808 | -1.542625098 | 0.008943433 |
| Gfm2 | -0.384694961 | 0.495820679 | -0.53822758 | 0.004045358 | -0.962474125 | 0.418677481 |
| Alcam | 0.119164169 | 0.820510156 | -0.538640205 | 4.48E-08 | -0.273577757 | 0.993542839 |
| Afg3l1 | -0.324361088 | 0.58730628 | -0.540296571 | 0.002633112 | -0.657087589 | 0.833210591 |
| Rab43 | -0.399440811 | 0.526732832 | -0.541453299 | 0.012261334 | -0.711270876 | 0.661491752 |
| Mchr1 | -0.200978749 | 0.845988693 | -0.541561153 | 0.006620959 | -1.426238063 | 0.162913907 |
| Pkdcc | -0.258619136 | 0.761454672 | -0.541808401 | 0.023664731 | -0.996034042 | 0.493900582 |
| Agt | -0.521509437 | 0.125054301 | -0.542222314 | 0.016663557 | -0.843674138 | 0.668491626 |
| Gramd1b | -0.400183542 | 0.103654294 | -0.542253128 | 8.42E-07 | -0.805293951 | 0.752700131 |
| Nat8f1 | -0.463456788 | 0.175626579 | -0.542877217 | 0.000264025 | -0.994163834 | 0.248565899 |
| Fbxw5 | -0.150972507 | 0.617042621 | -0.543811289 | 1.15E-10 | -0.889978821 | 0.661661871 |
| 1810055G02Rik | -0.362296751 | 0.448565591 | -0.544173432 | 0.000737919 | -1.012485069 | 0.384513629 |
| Lims2 | -0.334356401 | 0.535360709 | -0.54447927 | 0.010984287 | -0.945177561 | 0.286694005 |
| B230307C23Rik | -1.000441028 | 0.383802165 | -0.544889473 | 0.035880902 | -1.21553796 | 0.215418753 |
| Hectd2 | -0.542032462 | 0.51997736 | -0.545277255 | 0.032318421 | -0.522706583 | 0.993542839 |
| Arhgap33 | -0.226536208 | 0.541278577 | -0.545555 | 0.000289088 | -0.964734739 | 0.661512067 |
| Sephs1 | -0.65899018 | 0.117621546 | -0.545854539 | 0.000946195 | -1.007510786 | 0.41492578 |
| Snhg10 | -0.363952119 | 0.659370497 | -0.546059354 | 0.027480542 | -0.722105567 | 0.79107705 |
| Mettl17 | -0.550909373 | 0.447238852 | -0.547472609 | 0.002531407 | -0.645202671 | 0.788381343 |
| Snapc4 | -0.353655868 | 0.543527334 | -0.547909532 | 0.014342359 | -1.016275409 | 0.340227357 |
| Plxna1 | -0.551997339 | 0.150280011 | -0.54798604 | 0.002223457 | -1.053682302 | 0.268832642 |
| Inhbb | -1.129841461 | 0.158899163 | -0.54907059 | 0.04382866 | -1.012918344 | 0.355381522 |
| Zfand2a | -0.10750513 | 0.832762217 | -0.550320316 | 1.12E-05 | -0.637280068 | 0.993542839 |
| Ermard | -0.156927502 | 0.887344475 | -0.550647938 | 0.007730632 | -0.817355754 | 0.53005075 |
| Tab1 | -0.573083854 | 0.182467761 | -0.551044616 | 0.000153677 | -0.806567314 | 0.45949895 |
| Slc17a7 | -0.367475674 | 0.017481945 | -0.551530477 | 9.10E-11 | -1.03433026 | 0.828701453 |
| Rimkla | -0.261352696 | 0.770843306 | -0.552208298 | 0.00382006 | -1.109772136 | 0.227083554 |
| Mob1b | -0.072617543 | 0.925926629 | -0.552288098 | 3.15E-06 | -0.683918631 | 0.936045285 |
| Tmem117 | -0.287973207 | 0.816137512 | -0.552477744 | 0.039959832 | -0.724905263 | 0.929751641 |
| Gm1976 | -0.178161676 | 0.855240576 | -0.552573168 | 0.000427576 | -0.529451075 | 0.993542839 |
| 2700081O15Rik | -0.407117345 | 0.284144605 | -0.552576828 | 4.98E-05 | -0.971116151 | 0.306244669 |
| Fbxl19 | -0.446764055 | 0.271492763 | -0.552681834 | 4.29E-05 | -0.942559604 | 0.304838758 |
| A330008L17Rik | -0.564761076 | 0.699405851 | -0.552919726 | 0.035503951 | 0.090779767 | 0.993542839 |
| Arhgef7 | -0.460992365 | 0.056243432 | -0.553004819 | 9.54E-09 | -0.866547021 | 0.727825365 |
| Rap2b | -0.376353016 | 0.298969118 | -0.553469825 | 3.03E-06 | -0.701281085 | 0.940570955 |
| Sfswap | -0.320661652 | 0.397342355 | -0.553533639 | 2.09E-05 | -1.080564609 | 0.261034447 |
| Smox | -0.373827048 | 0.271492763 | -0.554460288 | 4.88E-05 | -0.819541641 | 0.612369982 |
| Slc19a1 | -0.414316936 | 0.36121298 | -0.5547201 | 0.008228297 | -0.877506426 | 0.499928377 |
| Wbp1 | -0.472102112 | 0.087415107 | -0.555087739 | 1.37E-06 | -0.953746499 | 0.45949895 |
| Dap | -0.584918494 | 0.278072193 | -0.555237569 | 0.006718069 | -0.671553326 | 0.777821 |
| Ascc2 | -0.03490025 | 0.973475731 | -0.55562057 | 0.009717936 | -0.400370708 | 0.993542839 |
| Pdzd2 | -0.729455845 | 0.183011401 | -0.556344731 | 0.047772747 | -1.386482314 | 0.070736914 |
| Gda | -0.338876467 | 0.145701668 | -0.556560829 | 0.000317942 | -0.768333554 | 0.993542839 |
| Lhfpl4 | -0.331822885 | 0.200911893 | -0.55805402 | 3.21E-13 | -0.926787476 | 0.605843331 |
| Ankrd16 | -0.537265639 | 0.315286697 | -0.558279414 | 0.00817617 | -0.940068618 | 0.448360032 |
| 9430015G10Rik | -0.070092062 | 0.96356596 | -0.558686969 | 0.00637411 | -0.718026378 | 0.681338109 |
| Sdc3 | -0.454157311 | 0.078883144 | -0.558941676 | 0.001326053 | -1.13123138 | 0.440677298 |
| Astn2 | -0.29834252 | 0.764098124 | -0.559036874 | 0.021876233 | -0.14658178 | 0.993542839 |
| Mapk7 | -0.828883508 | 0.193140335 | -0.559368668 | 0.017230177 | -0.969410351 | 0.504435724 |
| Grik2 | -0.23307807 | 0.695916002 | -0.560014676 | 0.00027727 | -0.536411404 | 0.993542839 |
| Sugp2 | -0.141915286 | 0.788953496 | -0.560584959 | 3.73E-09 | -0.956277614 | 0.558601332 |
| 1110017D15Rik | -0.419146515 | 0.777139605 | -0.562153283 | 0.037138434 | -0.875078273 | 0.442019435 |
| Lhpp | -0.250684156 | 0.698585117 | -0.563212553 | 0.001167978 | -0.853531542 | 0.551475793 |
| Alg6 | -0.171057194 | 0.886466511 | -0.563282713 | 0.009486434 | -0.736105904 | 0.711807045 |
| Wdr5 | -0.238620822 | 0.744084548 | -0.563771655 | 0.002671196 | -0.812270044 | 0.683203063 |
| Abcb6 | -0.436869457 | 0.42964689 | -0.563783708 | 0.000225565 | -1.022549722 | 0.366560931 |
| Itpr2 | -0.296032408 | 0.642026421 | -0.563905596 | 0.010391856 | -0.776115064 | 0.81939952 |
| Pik3r3 | -0.454695823 | 0.181342299 | -0.564639605 | 5.45E-06 | -0.927305858 | 0.559235611 |
| Rtel1 | -1.074772428 | 0.071387638 | -0.564783657 | 0.034938995 | -0.799054155 | 0.810895396 |
| Tax1bp3 | -0.52742183 | 0.225047698 | -0.564978653 | 0.007399142 | -0.861149548 | 0.487685487 |
| Hgh1 | -0.401976956 | 0.670833621 | -0.565145163 | 0.002242556 | -0.722981526 | 0.655539463 |
| Cdk16 | -0.41410477 | 0.043987673 | -0.56571547 | 4.13E-06 | -1.045847534 | 0.492520769 |
| Fem1a | -0.054432882 | 0.939450278 | -0.566062925 | 1.39E-06 | -0.898254522 | 0.500668007 |
| Vps37c | -0.500220835 | 0.387530907 | -0.566402706 | 0.000487744 | -0.880771319 | 0.439842065 |

| | | | | | | |
|---|---|---|---|---|---|---|
| Mmp16 | -1.0037724 | 0.172645291 | -0.566717355 | 0.025479188 | -1.11280075 | 0.457385996 |
| 2700049A03Rik | -0.286464567 | 0.816788729 | -0.566939301 | 0.033704644 | -0.81953524 | 0.74445641 |
| Tigd5 | -0.468733097 | 0.620140501 | -0.567556198 | 0.012220191 | -0.629952079 | 0.84752452 |
| Scn3a | -0.656239378 | 0.271037157 | -0.568659436 | 0.012459465 | -0.564913309 | 0.993542839 |
| 3110039M20Rik | -0.171021786 | 0.814831517 | -0.569455078 | 0.000443675 | -0.841918345 | 0.605843331 |
| Atp13a2 | -0.354859506 | 0.204712901 | -0.574296166 | 5.73E-07 | -1.082283883 | 0.424642494 |
| Fam219b | -0.380891677 | 0.506575804 | -0.574512135 | 8.72E-05 | -0.713787898 | 0.810531232 |
| Wsb1 | -0.486196281 | 0.143641898 | -0.574724955 | 0.003629331 | -0.974178186 | 0.51705363 |
| Chpf2 | -0.426264225 | 0.23597416 | -0.574915577 | 0.000170303 | -0.923972665 | 0.670470868 |
| Tspoap1 | -0.541192918 | 0.140054282 | -0.575620917 | 0.001005745 | -1.046251934 | 0.448360032 |
| P3h1 | 0.213086178 | 0.857805193 | -0.575719756 | 0.024367594 | -0.993952326 | 0.268832642 |
| Prex1 | -0.433383092 | 0.070508408 | -0.577567369 | 3.79E-07 | -0.708620927 | 0.898348809 |
| Trit1 | -0.468497226 | 0.544297126 | -0.579749667 | 0.00556526 | -0.943258437 | 0.329229158 |
| Dmpk | -0.240585008 | 0.800819574 | -0.580059727 | 0.005792274 | -0.733336565 | 0.918033192 |
| Lmo3 | -0.474084148 | 0.211143524 | -0.581190232 | 0.002920496 | -0.5241804 | 0.993542839 |
| Lrrc3 | -0.196012774 | 0.870772081 | -0.581480506 | 0.012567598 | -1.046331562 | 0.711807045 |
| Ago4 | 0.086410523 | 0.955331783 | -0.583534917 | 0.033467712 | -0.90227357 | 0.613158534 |
| Prep | -0.572219659 | 0.177168005 | -0.583772408 | 0.006825284 | -1.196269439 | 0.194839114 |
| Tyk2 | -0.664324766 | 0.521933557 | -0.584835159 | 0.017511678 | -0.72867441 | 0.898104584 |
| Pmm2 | -0.432931981 | 0.494766805 | -0.585028543 | 0.006090887 | -1.028568645 | 0.171842435 |
| Slc4a7 | -0.416511324 | 0.542807881 | -0.585119365 | 0.004646753 | -0.852999469 | 0.752700131 |
| Wwc1 | -0.363892224 | 0.208048161 | -0.585180506 | 1.00E-05 | -0.677569012 | 0.993542839 |
| Dvl2 | -0.307039259 | 0.738694292 | -0.585562558 | 0.023845501 | -1.035219856 | 0.165927265 |
| Usp6nl | -0.488739927 | 0.419522486 | -0.58629394 | 0.002203576 | -0.89065789 | 0.425234843 |
| Spaca6 | -0.366886929 | 0.634623901 | -0.586539096 | 0.022577276 | -0.986899544 | 0.395359805 |
| Ephb2 | -1.042940101 | 0.057806404 | -0.58685776 | 0.010888699 | -0.741045009 | 0.727825365 |
| Mov10 | -0.23329328 | 0.844487367 | -0.586857919 | 0.033731436 | -0.859270107 | 0.865366478 |
| Lcorl | -0.393240169 | 0.536833287 | -0.587105311 | 0.012551291 | -0.582193719 | 0.95811614 |
| Mbd6 | -0.641706009 | 0.098980344 | -0.587355265 | 0.011983175 | -0.86752333 | 0.476137166 |
| Cfp | -0.422284149 | 0.678475555 | -0.588018263 | 0.019638338 | -0.726028162 | 0.887385118 |
| Sh3pxd2b | 0.163656333 | 0.866171631 | -0.58822435 | 0.012958063 | -1.088373269 | 0.215418753 |
| Gm39469 | -0.089831974 | 0.950361554 | -0.588394808 | 0.02725007 | -1.107106176 | 0.480415331 |
| Tysnd1 | -0.510568156 | 0.262230407 | -0.588498227 | 9.08E-06 | -0.836893605 | 0.455540245 |
| Ramp3 | -0.291880389 | 0.67434971 | -0.588570645 | 0.017967985 | -0.262574632 | 0.993542839 |
| Ncam1 | -0.46647079 | 0.044167695 | -0.589091001 | 9.47E-05 | -1.008475723 | 0.651644024 |
| Usp40 | -1.062262494 | 0.156401924 | -0.589713248 | 0.018077803 | -0.724728283 | 0.725906701 |
| Acvr2a | -0.682147184 | 0.076154255 | -0.589722292 | 9.39E-05 | -0.845298757 | 0.727825365 |
| Tmem121b | -0.631568273 | 0.086610186 | -0.590082238 | 0.002109789 | -1.147965025 | 0.598135223 |
| Heg1 | -0.841224797 | 0.068691824 | -0.59022895 | 0.008316577 | -1.060351466 | 0.379400692 |
| Miip | -1.264716782 | 0.009584846 | -0.590286698 | 0.008792846 | -1.083942968 | 0.414591828 |
| Sarm1 | -0.606874848 | 0.297609224 | -0.590467395 | 0.000221772 | -0.897046007 | 0.458222912 |
| Shroom2 | -0.475282939 | 0.038925056 | -0.590835555 | 0.000207202 | -0.928270325 | 0.492520769 |
| Numbl | -0.367471294 | 0.146872045 | -0.591148047 | 2.82E-10 | -0.828320694 | 0.847029174 |
| Adra2a | 0.028688742 | 0.978077736 | -0.591308057 | 0.000241617 | -0.662527904 | 0.993542839 |
| Vwa5b2 | -0.209553276 | 0.754781572 | -0.591996803 | 0.002869415 | -0.649761172 | 0.86359932 |
| Ipcef1 | -0.486448284 | 0.26493801 | -0.592688133 | 0.003037308 | -0.554945824 | 0.993542839 |
| Trmu | -0.691934863 | 0.267943887 | -0.592873631 | 0.001328113 | -1.098539588 | 0.120455841 |
| Snhg5 | -0.542748534 | 0.455485074 | -0.593688169 | 0.004597198 | -1.016558342 | 0.201763205 |
| Clic1 | -0.406803895 | 0.30275585 | -0.593999196 | 0.002099464 | -0.763278051 | 0.67132303 |
| Bcan | -0.346889466 | 0.141554382 | -0.595074769 | 0.000386582 | -1.284351001 | 0.223860837 |
| Carmil1 | -0.3057797 | 0.699593518 | -0.595478376 | 0.003174283 | -0.960916839 | 0.623433955 |
| Bhlhe22 | -0.549920212 | 0.134092936 | -0.595579652 | 0.030958782 | -0.776778964 | 0.90514278 |
| Trp53inp2 | -0.478765665 | 0.01800354 | -0.597021009 | 1.13E-05 | -0.956658259 | 0.565412228 |
| 09.Mar | -0.344186804 | 0.45424603 | -0.597056862 | 7.08E-05 | -1.011436452 | 0.442019435 |
| Supt20 | -0.246625272 | 0.744697748 | -0.598634665 | 0.000910872 | -0.462741007 | 0.993542839 |
| Znrf1 | -0.346020355 | 0.090474508 | -0.600833826 | 1.28E-09 | -0.86035142 | 0.752700131 |
| Zfp784 | -0.360665655 | 0.857805193 | -0.600931964 | 0.048412851 | -0.073066045 | 0.993542839 |
| Spred3 | -0.167847501 | 0.838453873 | -0.602792603 | 0.002331905 | -0.788419205 | 0.617082316 |
| Abcb1a | -0.118207778 | 0.883163442 | -0.604145314 | 0.001820218 | -0.952517094 | 0.230084888 |
| 1500026H17Rik | -0.156073608 | 0.86668022 | -0.604260612 | 0.024593577 | -0.814291435 | 0.838213606 |
| Sox17 | -0.683952733 | 0.315286697 | -0.604868413 | 0.038961075 | -0.386221762 | 0.993542839 |
| Rfx3 | -0.317154975 | 0.565281461 | -0.605156149 | 0.007118468 | -0.746729675 | 0.675711796 |
| Hps4 | -0.517575845 | 0.403922921 | -0.606628677 | 0.00074251 | -1.079405447 | 0.16514846 |
| 2610008E11Rik | -0.743589968 | 0.130558121 | -0.606694137 | 0.000979136 | -0.774971848 | 0.612202342 |
| Tunar | -0.350293461 | 0.69571614 | -0.607020154 | 0.003335991 | -0.759402223 | 0.691499826 |
| Eml5 | -0.334805777 | 0.41854679 | -0.607191699 | 9.63E-05 | -0.814385315 | 0.504435724 |
| Tctn3 | -0.281450581 | 0.727185004 | -0.607546317 | 0.000856875 | -0.828230085 | 0.548463319 |
| Fam102b | -0.565292604 | 0.184936928 | -0.607674013 | 0.02835079 | -0.924958258 | 0.581792313 |
| Ttc13 | -0.272804841 | 0.600205597 | -0.607914444 | 4.34E-05 | -1.09554038 | 0.256821027 |
| Rnf165 | -0.133115203 | 0.818980228 | -0.60799314 | 0.000634261 | -1.057734889 | 0.33853316 |
| Apc | -0.509774469 | 0.089391351 | -0.608728718 | 0.002778387 | -1.219066114 | 0.353913196 |
| A730063M14Ril | 0.007189858 | 0.995159917 | -0.609323038 | 0.028796514 | -1.044569249 | 0.171308801 |
| Gm2a | -0.408516993 | 0.179393614 | -0.61082179 | 0.005256335 | -0.904516962 | 0.61271111 |

| | | | | | | |
|---|---|---|---|---|---|---|
| Smpd2 | -0.010499899 | 0.990327384 | -0.610949698 | 0.000385424 | -0.84397308 | 0.451775056 |
| Grik5 | -0.443519231 | 0.000702478 | -0.611263196 | 9.25E-19 | -1.011381538 | 0.661491752 |
| Ppp1r18 | -0.471014317 | 0.410227457 | -0.611591424 | 0.003311271 | -0.485483087 | 0.993542839 |
| Alms1 | -0.203661144 | 0.876554129 | -0.612045991 | 0.02437541 | -0.350726937 | 0.993542839 |
| Stk17b | -0.266107257 | 0.705990885 | -0.612132104 | 0.037195502 | -0.770249052 | 0.727825365 |
| Arhgap29 | -0.122401917 | 0.884609978 | -0.612733173 | 0.010705325 | -0.713222927 | 0.738568045 |
| Zfyve27 | -0.171082836 | 0.721246706 | -0.613140001 | 2.73E-05 | -0.984147601 | 0.472868398 |
| Popdc3 | -0.023370507 | 0.990876969 | -0.615485689 | 0.038908462 | -0.652018953 | 0.993542839 |
| Nsmf | -0.378520439 | 0.105321351 | -0.616126708 | 3.11E-09 | -0.941768322 | 0.805167961 |
| Fbxo41 | -0.464764305 | 0.069589804 | -0.61700057 | 2.12E-09 | -0.889308706 | 0.638510559 |
| Purg | -0.630559795 | 0.064770068 | -0.618740877 | 0.001058965 | -1.144288049 | 0.204447175 |
| Kdm4b | -0.255683358 | 0.613053483 | -0.618827553 | 4.13E-09 | -1.043896329 | 0.232215573 |
| Cmtm3 | -0.473168763 | 0.330454601 | -0.619048245 | 0.02437541 | -0.612902152 | 0.914497701 |
| Mdm1 | -1.676945441 | 0.114460243 | -0.619100064 | 0.048896532 | -0.817099693 | 0.565412228 |
| Ndc1 | -0.794729995 | 0.063722808 | -0.619617939 | 4.65E-05 | -1.12589386 | 0.162913907 |
| Bgn | 0.477261295 | 0.57398609 | -0.620017521 | 0.028532728 | -1.097939224 | 0.200280631 |
| Slc30a10 | -0.441497258 | 0.433289316 | -0.620183919 | 0.022152172 | -0.872092235 | 0.420244076 |
| Tbcd | -0.389777444 | 0.34590193 | -0.620756412 | 1.09E-07 | -0.864194069 | 0.702744926 |
| 0610040J01Rik | 0.320003979 | 0.790232298 | -0.62156199 | 0.049181558 | -0.700932184 | 0.914497701 |
| Rep15 | -1.352485009 | 0.014177438 | -0.62246648 | 0.018071791 | -1.069771895 | 0.223830609 |
| Elfn1 | -0.462563814 | 0.545080266 | -0.622472414 | 0.000385231 | -0.790000385 | 0.661512067 |
| Gm28151 | -0.492184604 | 0.744084548 | -0.622631154 | 0.041859456 | -0.462446937 | 0.993542839 |
| Slc29a3 | -0.236375785 | 0.777784833 | -0.622657822 | 0.002865901 | -0.886751872 | 0.361261997 |
| Cd2ap | -0.285379949 | 0.645937808 | -0.624689346 | 0.000709675 | -0.794691656 | 0.767456916 |
| Galns | 0.028447655 | 0.985746783 | -0.625515275 | 0.007887472 | -0.516480488 | 0.993542839 |
| Gpr173 | -0.670201178 | 0.326451801 | -0.625619088 | 0.000396249 | -0.795327072 | 0.557037243 |
| Ptprb | -0.465938917 | 0.452992048 | -0.626160427 | 0.007284357 | -1.216904451 | 0.200280631 |
| Fam241a | -0.01649389 | 0.992196944 | -0.626197502 | 0.03889043 | -0.387350122 | 0.993542839 |
| Cidea | -0.308153573 | 0.770773253 | -0.627059286 | 0.039089693 | -0.845701591 | 0.898104584 |
| Wnt7b | -0.390093156 | 0.224204465 | -0.627367262 | 0.001164401 | -0.84811154 | 0.715362667 |
| Zfp811 | 0.431444175 | 0.73598328 | -0.627376541 | 0.043979388 | -0.648120984 | 0.771757636 |
| Frmd4a | -0.666674431 | 0.075744845 | -0.629362604 | 0.003760965 | -1.109340835 | 0.139657782 |
| Rfc5 | -0.759021712 | 0.253306262 | -0.630092347 | 0.001647426 | -0.695118293 | 0.689744667 |
| Fscn1 | -0.421141722 | 0.06225414 | -0.630106332 | 1.97E-05 | -1.091707145 | 0.499928377 |
| Gdpd2 | 0.079276038 | 0.964815151 | -0.632501962 | 0.048329176 | -0.87119722 | 0.727638346 |
| Pcyt1b | -0.116252312 | 0.878843747 | -0.632591126 | 0.000814149 | -0.932982803 | 0.486642813 |
| 5430416N02Rik | -0.649755024 | 0.18075732 | -0.633253735 | 0.000196536 | -1.042629414 | 0.305037962 |
| Atp10a | -0.598847333 | 0.368934331 | -0.634837422 | 0.002001157 | -0.630392139 | 0.942706292 |
| Dcbld2 | -0.332528232 | 0.517731848 | -0.635002268 | 6.31E-06 | -0.904395155 | 0.472868398 |
| D630045J12Rik | 0.040416472 | 0.964815151 | -0.635207257 | 1.27E-05 | -1.049844615 | 0.324775052 |
| Pigo | -0.38273577 | 0.53648286 | -0.635442333 | 5.48E-05 | -0.896448211 | 0.480220174 |
| Garnl3 | -0.119232889 | 0.816835979 | -0.63552462 | 0.000454583 | -0.848680627 | 0.90514278 |
| Cd276 | -1.221149074 | 0.141987213 | -0.635721302 | 0.032433475 | -0.831796909 | 0.550971806 |
| Slco1a4 | -0.525384854 | 0.147724528 | -0.636248067 | 0.000273977 | -0.943241685 | 0.231636794 |
| Mast1 | -0.474660285 | 0.030246679 | -0.636326722 | 1.73E-09 | -1.080791322 | 0.428402898 |
| Kctd15 | -0.356629449 | 0.634171195 | -0.636529753 | 0.00611049 | -0.960683499 | 0.35570703 |
| Cacna1i | -0.506151623 | 0.183011401 | -0.636882512 | 3.00E-05 | -0.82474873 | 0.610842744 |
| Hs3st4 | -0.595900258 | 0.102685451 | -0.636885306 | 0.002255077 | -0.670050906 | 0.993542839 |
| 5830448L01Rik | -0.556384785 | 0.610424898 | -0.63711915 | 0.031632693 | -0.634843347 | 0.993542839 |
| Cep89 | -0.243507777 | 0.774726519 | -0.637947366 | 0.009378206 | -1.225891512 | 0.139657782 |
| Plod1 | -0.435211474 | 0.383802165 | -0.638546985 | 0.002461371 | -0.822474641 | 0.55026974 |
| Cfap69 | -0.478047228 | 0.64080298 | -0.640686126 | 0.006619185 | -0.672802238 | 0.783606847 |
| Sipa1 | -0.280711903 | 0.679709777 | -0.641040834 | 0.00520342 | -1.143555658 | 0.143948382 |
| Zkscan17 | -0.726334935 | 0.23128754 | -0.641441098 | 0.004924038 | -0.839798647 | 0.557634413 |
| Syngap1 | -0.630475855 | 0.080672292 | -0.641605049 | 0.002640931 | -0.909480727 | 0.651644024 |
| Tac1 | -0.356280745 | 0.564659219 | -0.642924369 | 0.000637665 | -0.90554678 | 0.642474421 |
| Abca3 | -0.447739864 | 0.0706622 | -0.643274406 | 4.98E-06 | -1.080659254 | 0.46219062 |
| Rhbdl3 | -0.514883436 | 0.591821291 | -0.644231584 | 0.004929688 | -0.958777754 | 0.271077972 |
| Kctd5 | -0.548859058 | 0.164544197 | -0.645011028 | 2.51E-05 | -1.228713161 | 0.149930259 |
| Gdf11 | -0.556906644 | 0.23597416 | -0.645290689 | 0.000609636 | -0.748731623 | 0.655539463 |
| Slc2a6 | -0.29144815 | 0.58106819 | -0.645898236 | 6.66E-06 | -1.352530645 | 0.268832642 |
| Zfp579 | -0.030367943 | 0.977489239 | -0.646273925 | 5.31E-06 | -1.000303043 | 0.365409496 |
| Tspan4 | -0.44229474 | 0.132849902 | -0.646434823 | 3.75E-05 | -1.037029415 | 0.206337599 |
| Cdc7 | -0.939275255 | 0.215434877 | -0.647903806 | 0.009284853 | -0.898615662 | 0.261034447 |
| Sult1a1 | 0.022841152 | 0.978804679 | -0.648743046 | 0.000265108 | -0.873283807 | 0.482241964 |
| Lpcat3 | -0.339408531 | 0.565281461 | -0.650960098 | 0.000794974 | -0.560689886 | 0.957602283 |
| Ust | -0.595499593 | 0.129064883 | -0.651421465 | 0.000233301 | -1.038713078 | 0.204447175 |
| Limd2 | -0.344411299 | 0.25200889 | -0.652212539 | 1.50E-09 | -0.560942117 | 0.993542839 |
| Obsl1 | -0.911441176 | 0.159065979 | -0.653301388 | 0.000880141 | -0.675972217 | 0.945315481 |
| Sh3bp4 | -0.729952733 | 0.055805192 | -0.654108076 | 0.000412661 | -0.668759724 | 0.993542839 |
| Kank3 | -0.616864408 | 0.23129921 | -0.65439718 | 0.000651601 | -0.932428822 | 0.285054031 |
| Igfbp7 | -0.249624348 | 0.657694994 | -0.654503478 | 0.000257431 | -1.253859892 | 0.120455841 |
| Bach2 | 0.412500329 | 0.696386709 | -0.654537589 | 0.038908462 | -0.931021304 | 0.392980578 |

| | | | | | | |
|---|---|---|---|---|---|---|
| Rin1 | -0.578109584 | 0.194479741 | -0.655426211 | 1.35E-05 | -0.34824958 | 0.993542839 |
| Dock6 | -0.26735069 | 0.741274047 | -0.656091523 | 0.011481344 | -0.880518161 | 0.480220174 |
| Syce2 | -0.795154322 | 0.268459897 | -0.657042491 | 0.007871793 | -1.110495504 | 0.324775052 |
| Hcn4 | -0.28680625 | 0.865531666 | -0.657609921 | 0.036185897 | -0.57768226 | 0.993542839 |
| Shmt2 | -0.354577684 | 0.557069181 | -0.660792192 | 2.67E-05 | -0.80763961 | 0.602533675 |
| 4930570G19Rik | -0.285693182 | 0.845988693 | -0.66118158 | 0.005334591 | -0.663785485 | 0.708926839 |
| Gm28050 | -1.333929729 | 0.00718833 | -0.662349767 | 0.00079759 | -0.336747395 | 0.993542839 |
| Rab3d | -0.670562277 | 0.412018221 | -0.662933242 | 0.004143247 | -1.375803413 | 0.188587928 |
| Zcchc7 | -0.716689163 | 0.212709813 | -0.663296892 | 0.000651995 | -1.004177228 | 0.293558199 |
| Fkbp5 | -0.454249274 | 0.297794926 | -0.664337261 | 0.000245953 | -0.538747488 | 0.993542839 |
| Chfr | -0.793881344 | 0.089986387 | -0.665261097 | 5.64E-05 | -1.224293051 | 0.179310236 |
| Adgrf5 | -0.462484286 | 0.394827922 | -0.666863129 | 0.000430278 | -0.638711597 | 0.852409217 |
| Iqce | -0.092010571 | 0.924917225 | -0.667945772 | 0.000276576 | -0.853090577 | 0.582320142 |
| Pcdh11x | -0.041906114 | 0.981092457 | -0.669213084 | 0.049072178 | -0.156282203 | 0.993542839 |
| Csmd2 | -0.756676766 | 0.150516444 | -0.66988617 | 0.000588444 | -0.755715725 | 0.993542839 |
| Abcd1 | -0.428580038 | 0.495866108 | -0.670993091 | 0.000326482 | -1.022825633 | 0.230084888 |
| Sdk2 | -0.652361737 | 0.43075221 | -0.67127037 | 0.012889584 | -0.757642583 | 0.993542839 |
| Uck2 | -1.574899035 | 0.001654585 | -0.671343533 | 0.001495826 | -1.121202942 | 0.305112527 |
| Slc1a6 | -0.510406696 | 0.237382896 | -0.673526251 | 0.000873012 | -0.692552443 | 0.724954309 |
| Fam189a2 | -0.055807511 | 0.957898224 | -0.674047765 | 0.008827818 | -0.847408348 | 0.420366805 |
| Megf11 | -0.434936696 | 0.355872186 | -0.678096208 | 0.003377881 | -1.405207149 | 0.029304431 |
| Ogfr | -0.492150455 | 0.175061743 | -0.678099639 | 2.13E-05 | -1.010826639 | 0.287037664 |
| Gm12371 | -0.45921756 | 0.754820056 | -0.679178836 | 0.03589756 | -0.636362121 | 0.993542839 |
| Fam189b | -0.474029247 | 0.203527529 | -0.679437242 | 8.61E-06 | -1.043595385 | 0.375588387 |
| Dlg2 | -0.434294708 | 0.029799226 | -0.680661827 | 1.78E-07 | -1.093713593 | 0.548463319 |
| Vat1l | -0.689479699 | 0.078606748 | -0.680884635 | 0.000107724 | -1.093590717 | 0.445982173 |
| Ocel1 | -0.742613687 | 0.162409096 | -0.681877651 | 0.00032733 | -1.046604034 | 0.279604201 |
| Zdhhc12 | -0.823207512 | 0.258593134 | -0.682140532 | 0.003243241 | -1.323646705 | 0.079943479 |
| Snhg11 | -0.592671463 | 0.202507216 | -0.682624644 | 0.020557542 | -1.16394428 | 0.613158534 |
| Scmh1 | -0.383000535 | 0.301440827 | -0.683271353 | 4.35E-05 | -1.178314622 | 0.133225497 |
| Trib2 | -0.700604749 | 0.056469132 | -0.683688661 | 0.007410326 | -0.783951349 | 0.993542839 |
| Ick | -0.53475166 | 0.399694016 | -0.684075368 | 0.00053359 | -0.672991473 | 0.89196315 |
| Fam163b | -0.622718929 | 0.089391351 | -0.684166394 | 0.000461814 | -0.908389298 | 0.776116412 |
| Brinp3 | -0.671811842 | 0.156483354 | -0.68463286 | 0.00377465 | -0.962103577 | 0.883573732 |
| Nit2 | -0.711233838 | 0.193140335 | -0.685322536 | 0.000550258 | -0.997229902 | 0.305112527 |
| Lrrc45 | -0.574568613 | 0.114978712 | -0.685802262 | 6.20E-05 | -1.095697646 | 0.348436139 |
| Mterf1a | -0.170862758 | 0.905382356 | -0.685908092 | 0.00533297 | -0.682073159 | 0.702044135 |
| Atp11b | -0.773951794 | 0.122238623 | -0.686595431 | 0.00021509 | -1.192690375 | 0.13599948 |
| Ptprr | -0.42891342 | 0.3734325 | -0.688455542 | 0.000281564 | -0.780941916 | 0.706091083 |
| Efcab6 | -0.241890648 | 0.884067647 | -0.688579655 | 0.044201489 | -0.952572362 | 0.342004625 |
| A230051N06Rik | -0.477701891 | 0.788507882 | -0.69079017 | 0.010008584 | -1.110426719 | 0.121081622 |
| A330102I10Rik | -1.037515465 | 0.22942492 | -0.691858523 | 0.006415963 | -1.314858725 | 0.063360951 |
| Bbc3 | -0.56207709 | 0.448512749 | -0.692417496 | 0.000248821 | -1.127716267 | 0.14509505 |
| Zfp174 | -0.180918272 | 0.923111754 | -0.693292937 | 0.041454447 | -0.461538642 | 0.993542839 |
| Vcam1 | -0.191063471 | 0.725800153 | -0.693607523 | 0.003060324 | -0.81940725 | 0.448141348 |
| Wdr90 | -1.263034682 | 0.064261478 | -0.694538152 | 0.016819914 | -0.949714381 | 0.290877569 |
| Cartpt | -0.670474814 | 0.334365306 | -0.695659362 | 0.031653774 | -1.313743161 | 0.272421392 |
| Glra2 | -1.31127417 | 0.120029366 | -0.69785772 | 0.016461303 | -0.854455187 | 0.906596277 |
| Hsd11b1 | -0.474094466 | 0.247782387 | -0.697883436 | 0.001228274 | -1.21269382 | 0.102511937 |
| Cyp11a1 | 0.482695126 | 0.704823134 | -0.697912295 | 0.01623202 | -0.860491402 | 0.846711539 |
| 8430429K09Rik | -0.711775871 | 0.096909033 | -0.699681609 | 0.000850682 | -1.089435069 | 0.235195451 |
| Fam193b | -0.474845519 | 0.252759779 | -0.699933373 | 1.13E-05 | -1.108031324 | 0.338503594 |
| B4galt2 | -0.464101764 | 0.039779349 | -0.701531875 | 3.96E-11 | -1.198623946 | 0.187221494 |
| Bak1 | -0.599444459 | 0.458248846 | -0.701620066 | 0.001554828 | -1.077770652 | 0.165855295 |
| Epb41l2 | -0.31698764 | 0.216690701 | -0.702846349 | 5.36E-07 | -0.982908011 | 0.480220174 |
| Gm5914 | -0.320570652 | 0.649813046 | -0.704902956 | 0.000818416 | -0.980504359 | 0.428402898 |
| Igflr1 | -0.533565042 | 0.53102009 | -0.705004017 | 0.036105171 | -1.047360584 | 0.46139675 |
| 5031434O11Rik | -0.674374585 | 0.401607301 | -0.706463515 | 0.011393961 | -0.745948197 | 0.952599775 |
| Rps6kb2 | -0.557422147 | 0.290530149 | -0.708042723 | 0.001590826 | -1.2032083 | 0.184983612 |
| Raver2 | -0.418009496 | 0.388146392 | -0.708921662 | 1.10E-05 | -0.865713732 | 0.471587545 |
| Slc45a4 | -0.319135932 | 0.458968581 | -0.709649682 | 1.22E-08 | -0.938217573 | 0.524703798 |
| Tek | -0.95262266 | 0.085976554 | -0.710719182 | 0.004795774 | -0.789761042 | 0.586695619 |
| Echdc2 | 0.151078303 | 0.873157988 | -0.712962672 | 0.002091383 | -0.965035941 | 0.279618923 |
| Gpsm2 | -0.289055424 | 0.785781078 | -0.71314849 | 0.014472703 | -0.62269559 | 0.946657533 |
| Thbs3 | -0.581133583 | 0.321709292 | -0.714669603 | 0.00054713 | -0.568246887 | 0.993542839 |
| Pantr1 | -0.26826359 | 0.186027228 | -0.715543104 | 1.40E-07 | -0.847491651 | 0.702744926 |
| Nectin3 | -0.642839979 | 0.325510182 | -0.716992406 | 0.006344717 | -0.907647834 | 0.437548745 |
| Tmem198b | -0.699474124 | 0.370640249 | -0.717608353 | 0.004032152 | -0.81977316 | 0.565412228 |
| Lama2 | 0.003266277 | 0.997072382 | -0.717843194 | 0.007794176 | -0.916198196 | 0.492520769 |
| Myo5b | -1.102185351 | 0.028052502 | -0.718524181 | 0.012962368 | -0.696054868 | 0.993542839 |
| Hip1 | -0.429324822 | 0.377449745 | -0.719904994 | 0.000881549 | -0.857897561 | 0.593780342 |
| Mycl | -0.900245461 | 0.212076143 | -0.720725759 | 0.001344118 | -0.731625839 | 0.833210591 |
| Fmnl3 | -0.073817886 | 0.955331783 | -0.722604125 | 0.013423822 | -0.604638266 | 0.930503377 |

| | | | | | | |
|---|---|---|---|---|---|---|
| Fmo1 | -1.025203856 | 0.033757441 | -0.723640882 | 0.028535071 | -1.281176997 | 0.169777283 |
| Rgs5 | -0.511220497 | 0.188598132 | -0.724927148 | 0.000493278 | -1.036745347 | 0.42727242 |
| Stx2 | -0.741370697 | 0.135195576 | -0.726261377 | 4.70E-05 | -1.316778888 | 0.066989675 |
| 9330020H09Rik | -0.193925003 | 0.848593627 | -0.728746996 | 0.000381605 | -1.108881479 | 0.156304324 |
| Aldh1b1 | -0.838958745 | 0.409824013 | -0.729025974 | 0.005477895 | -1.015710067 | 0.712523443 |
| Mdga1 | -0.426219249 | 0.609395843 | -0.729648523 | 0.001126241 | -0.498451294 | 0.993542839 |
| Cdk5r2 | -0.357317501 | 0.13135773 | -0.730803645 | 3.75E-05 | -1.0819454 | 0.528616571 |
| Gpam | -0.530124873 | 0.10402716 | -0.731611944 | 0.000794974 | -1.085417154 | 0.223860837 |
| Sfi1 | 0.003748259 | 0.997072382 | -0.736534006 | 0.010925475 | -1.098358881 | 0.214884113 |
| Cplx3 | 0.048710143 | 0.952189316 | -0.740483957 | 0.000735252 | -0.493535611 | 0.993542839 |
| C230057M02Rik | -0.771837517 | 0.228476422 | -0.741359596 | 0.001359149 | -0.658804897 | 0.790782897 |
| Nup210 | -0.426104188 | 0.52205735 | -0.741594685 | 6.79E-06 | -0.932182614 | 0.416271582 |
| Sema6c | -0.64076966 | 0.211137463 | -0.741749651 | 0.003236098 | -1.168802685 | 0.133225497 |
| Gm42067 | -0.37523073 | 0.362036304 | -0.74286733 | 7.23E-06 | -1.18854367 | 0.150294672 |
| Phtf2 | -0.249737717 | 0.773987264 | -0.743535065 | 0.015013147 | -1.179267509 | 0.548463319 |
| Id1 | -0.173269839 | 0.686113234 | -0.744045487 | 1.64E-07 | -0.98066104 | 0.348436139 |
| Afap1l2 | -0.665879254 | 0.271492763 | -0.744063582 | 0.004182058 | -0.85132571 | 0.55026974 |
| Ints6l | -0.111511995 | 0.893075846 | -0.744935164 | 0.001872688 | -0.93883338 | 0.304838758 |
| Adam17 | -0.33077735 | 0.709165071 | -0.745951724 | 0.000835737 | -0.846335388 | 0.373193584 |
| P2ry13 | 0.054804572 | 0.962253496 | -0.746961788 | 0.008363489 | -0.580883971 | 0.993542839 |
| Cyren | -0.91325454 | 0.112935488 | -0.747441069 | 1.62E-05 | -1.02757104 | 0.176091408 |
| Bcl6 | -0.087889654 | 0.904891074 | -0.74750616 | 9.97E-09 | -0.754402403 | 0.623433955 |
| Sertad4 | -0.30113699 | 0.758021812 | -0.748114418 | 0.000541262 | -1.352325021 | 0.204447175 |
| Glis2 | -0.515966363 | 0.39927175 | -0.748415199 | 0.001719843 | -0.822340012 | 0.46940436 |
| Thsd4 | -0.917949578 | 0.111408905 | -0.749247476 | 0.016217074 | -1.162475343 | 0.210888125 |
| Kif22 | -0.422118391 | 0.384084283 | -0.749252502 | 0.000179239 | -0.806126254 | 0.968479194 |
| Atp11c | -0.66329388 | 0.346849038 | -0.75096093 | 0.003517688 | -0.764836912 | 0.838213606 |
| 6330409D20Rik | 1.023593596 | 0.453329899 | -0.75152882 | 0.026838529 | -0.753265701 | 0.736117265 |
| Nhsl1 | 0.132824659 | 0.884969445 | -0.753883163 | 0.000205762 | -0.908463456 | 0.724979396 |
| Thpo | -0.791964672 | 0.418658857 | -0.754059545 | 0.013898606 | -1.047607252 | 0.235195451 |
| Ccdc141 | -1.116592465 | 0.075744845 | -0.755630427 | 0.011930571 | -0.488284506 | 0.993542839 |
| Slc6a20a | -0.129525959 | 0.91046804 | -0.756332397 | 0.001687084 | -1.007083949 | 0.330196193 |
| Ecm1 | -0.679163342 | 0.280310098 | -0.758788135 | 0.003078938 | -1.110480201 | 0.165855295 |
| Nectin2 | -0.540685468 | 0.452403192 | -0.761564754 | 0.001113299 | -1.143369564 | 0.166395649 |
| Thbd | 0.477813222 | 0.378921328 | -0.765524077 | 0.012161979 | -0.984163008 | 0.509418999 |
| Plppr1 | -0.597697353 | 0.378710324 | -0.766407372 | 0.008395768 | -1.375143153 | 0.189079328 |
| Sirt4 | -0.761884922 | 0.213096483 | -0.768124318 | 9.47E-05 | -0.67474982 | 0.776116412 |
| Palmd | 0.081038782 | 0.951213698 | -0.768777561 | 0.001831604 | -1.216740287 | 0.123169917 |
| Mirg | -0.40569188 | 0.464092819 | -0.770100961 | 1.98E-08 | -1.108224303 | 0.26348138 |
| Rad52 | -0.39843165 | 0.612670796 | -0.771719426 | 2.44E-05 | -0.909290038 | 0.35092649 |
| Slc7a6 | -0.740264248 | 0.233469764 | -0.77783577 | 7.67E-06 | -0.85846521 | 0.565412228 |
| Trpc4 | -0.378732368 | 0.611119261 | -0.779083041 | 0.00017058 | -1.18263673 | 0.217708863 |
| D430019H16Rik | -0.430867189 | 0.108063089 | -0.779281462 | 2.75E-05 | -1.073006089 | 0.384674356 |
| Pak6 | -0.508227633 | 0.184903152 | -0.781076049 | 3.34E-06 | -1.04282123 | 0.420366805 |
| 2810029C07Rik | -0.638366251 | 0.479209985 | -0.78112657 | 0.005041439 | -1.270452015 | 0.083278006 |
| Lratd2 | -0.848777462 | 0.566818603 | -0.781697398 | 0.038908462 | -0.666289268 | 0.969592356 |
| Slc16a7 | -0.648027414 | 0.186966414 | -0.78711071 | 3.17E-05 | -1.28612944 | 0.079139514 |
| Actn2 | -0.547548759 | 0.618796079 | -0.787508256 | 0.020201076 | -0.954567575 | 0.345024541 |
| Gmip | -0.48359658 | 0.781762865 | -0.789037967 | 0.018369867 | -0.807843515 | 0.5924171 |
| Cdh5 | -0.430774585 | 0.52205735 | -0.792322939 | 0.016002147 | -1.123028261 | 0.240109114 |
| Samd14 | -0.405753597 | 0.195872765 | -0.796480725 | 1.76E-06 | -0.889887963 | 0.612202342 |
| Myo7a | -0.698739912 | 0.361672552 | -0.80382234 | 1.95E-05 | -0.663402306 | 0.82469938 |
| Gm11627 | -0.953918702 | 0.099255357 | -0.805243401 | 9.04E-05 | -0.86676237 | 0.448411251 |
| Mapkbp1 | -0.598282134 | 0.135572425 | -0.805806179 | 2.20E-10 | -1.372674852 | 0.079943479 |
| Cd59a | -0.363772559 | 0.59815063 | -0.806115794 | 1.73E-05 | -0.916561943 | 0.27305958 |
| Prss36 | -0.509830184 | 0.474253754 | -0.808053756 | 1.55E-06 | -0.930157139 | 0.328516758 |
| Gm21781 | -1.247287511 | 0.29875803 | -0.809230734 | 0.01787627 | -0.579099604 | 0.993542839 |
| Homer3 | -0.422174704 | 0.350863804 | -0.80939924 | 0.00188067 | -0.746181826 | 0.57790817 |
| Apbb1ip | 0.466685914 | 0.650871792 | -0.811585843 | 0.003202498 | -0.656354725 | 0.965386891 |
| Shisa6 | -0.86062658 | 0.05387109 | -0.814138435 | 3.52E-05 | -1.289630433 | 0.068917542 |
| Zbtb5 | -0.813434697 | 0.296790717 | -0.815000142 | 0.000384279 | -1.305964784 | 0.05717001 |
| Ppfibp1 | -0.803032943 | 0.099424344 | -0.818784242 | 0.000248195 | -1.018537235 | 0.463705236 |
| Ppil6 | -1.324031969 | 0.151752913 | -0.818954856 | 0.007410325 | -1.139384554 | 0.156304324 |
| Zfp512b | -0.453042207 | 0.258943259 | -0.819570151 | 3.15E-06 | -1.246923938 | 0.156304324 |
| Nkiras2 | -0.675205682 | 0.291010128 | -0.819895753 | 0.003798381 | -1.398288446 | 0.050403738 |
| Rflnb | -1.19440457 | 0.025055529 | -0.822764863 | 0.012347746 | -1.284880639 | 0.173600143 |
| Sema5b | -1.003833741 | 0.109578433 | -0.825830317 | 0.000273274 | -0.85281416 | 0.499359773 |
| Tubd1 | -0.336087936 | 0.796829229 | -0.826684106 | 0.005387942 | -1.086549347 | 0.511900147 |
| Pdgfrb | -0.587293478 | 0.300359507 | -0.82808578 | 0.000360005 | -0.7214522 | 0.704589402 |
| Sox12 | -1.02124497 | 0.052441699 | -0.82918492 | 3.34E-06 | -1.175685933 | 0.105925498 |
| Nphp3 | 0.241861465 | 0.869962281 | -0.831925615 | 0.005061574 | -0.745021575 | 0.753331908 |
| Angptl4 | -1.206572159 | 0.045283476 | -0.832248257 | 0.011458564 | -0.923258728 | 0.649800646 |
| Ccdc57 | -0.292571393 | 0.81785846 | -0.835112317 | 0.003022093 | -0.914737248 | 0.582320142 |

| | | | | | | |
|---|---|---|---|---|---|---|
| Ackr1 | -0.591853641 | 0.081461262 | -0.837289599 | 5.67E-07 | -1.359638313 | 0.144616872 |
| Pde3b | -0.066392453 | 0.96842841 | -0.837581989 | 0.000658844 | -0.258925064 | 0.993542839 |
| Il16 | -0.443507416 | 0.40049309 | -0.839065768 | 0.001437218 | -0.725342241 | 0.743906977 |
| Gm19531 | -0.522371907 | 0.482196453 | -0.839180621 | 0.000229374 | -0.656157715 | 0.969841518 |
| Tubb2b | -0.572683491 | 0.090951901 | -0.842781806 | 0.001078269 | -1.642022936 | 0.122845847 |
| AI480526 | -0.508928822 | 0.523898825 | -0.844162659 | 6.50E-05 | -1.13661311 | 0.136779782 |
| Ryr3 | -1.370671291 | 0.017481945 | -0.844227314 | 0.000604946 | -0.972009429 | 0.442019435 |
| Snhg20 | -0.592846145 | 0.059577774 | -0.844266275 | 2.33E-05 | -1.164111745 | 0.282442752 |
| Gm43597 | -0.636548195 | 0.377978289 | -0.846609359 | 9.51E-06 | -0.990681365 | 0.374866046 |
| Rgs9 | -0.385276265 | 0.66182316 | -0.848813453 | 0.012775813 | -0.887056089 | 0.448141348 |
| Adamts17 | -0.442976198 | 0.762988733 | -0.848838132 | 0.020659853 | -0.624089052 | 0.993542839 |
| Kcne1l | -0.174223428 | 0.862554147 | -0.849323171 | 0.006942601 | -0.579396471 | 0.993542839 |
| Dab2 | -0.491907597 | 0.384198943 | -0.849725695 | 0.014387779 | -1.042341132 | 0.287037664 |
| Nfya | -0.388466455 | 0.645400215 | -0.850302218 | 4.89E-05 | -0.564653469 | 0.993542839 |
| Meak7 | -0.595503527 | 0.54708064 | -0.851711435 | 0.000116751 | -1.130204344 | 0.362676994 |
| Neu4 | -1.15705125 | 0.029062552 | -0.855875687 | 0.002482563 | -1.508646041 | 0.020799188 |
| Sh3rf3 | -0.279079431 | 0.79168295 | -0.856093142 | 0.000640049 | -1.202068936 | 0.284181522 |
| Tgfb2 | -0.091702787 | 0.937650567 | -0.858353369 | 0.004798623 | -0.945026967 | 0.410588099 |
| Pigc | -0.442822711 | 0.561953709 | -0.858600325 | 1.36E-05 | -0.983242128 | 0.279604201 |
| Adamtsl2 | -0.926542048 | 0.404711799 | -0.864139254 | 0.008457008 | -1.113037209 | 0.093755915 |
| Amer1 | -0.949399287 | 0.378710324 | -0.867007802 | 0.01233318 | -0.809273444 | 0.554575284 |
| Leng8 | -0.557763555 | 0.240557413 | -0.868471009 | 0.000493278 | -1.29402759 | 0.184651535 |
| Eme2 | -0.419991057 | 0.66109888 | -0.869024392 | 4.73E-05 | -1.301476411 | 0.14509505 |
| Eogt | -0.518549947 | 0.458395635 | -0.86931355 | 0.000939006 | -0.426059128 | 0.993542839 |
| Hapln1 | -0.778221819 | 0.200911893 | -0.869326353 | 0.009942644 | -1.095187064 | 0.215418753 |
| Mycn | -0.449956481 | 0.59420189 | -0.869741033 | 0.000304803 | -1.053745756 | 0.556086383 |
| Trpc3 | -0.809925365 | 0.103654294 | -0.87067738 | 0.000291147 | -0.449586218 | 0.993542839 |
| Ntf3 | -0.392645874 | 0.682700201 | -0.871805557 | 0.000439911 | -0.578819666 | 0.993542839 |
| Robo4 | -1.06869702 | 0.345932798 | -0.871937326 | 0.00520342 | -1.397305702 | 0.072150965 |
| 2310039H08Rik | -0.701688896 | 0.080013985 | -0.875304759 | 3.78E-07 | -1.014866385 | 0.416318665 |
| Nckap5l | -0.861353324 | 0.200156962 | -0.879152385 | 4.17E-05 | -0.825209539 | 0.603872653 |
| Fzd6 | -0.624415833 | 0.512545525 | -0.880126142 | 0.016482023 | -0.796402108 | 0.646744301 |
| Mirt1 | -0.962545813 | 0.365594375 | -0.881367762 | 0.006216331 | -1.125004259 | 0.184651535 |
| Col6a1 | 0.043138977 | 0.97243427 | -0.88469046 | 0.020365208 | -0.95354159 | 0.593819527 |
| Pcsk5 | -1.344361927 | 0.025055529 | -0.88603223 | 0.01990521 | -0.990372489 | 0.79972996 |
| Myt1 | -1.040941346 | 0.147724528 | -0.886884025 | 0.001270007 | -1.090590867 | 0.176091408 |
| Gja4 | -0.964173871 | 0.143083661 | -0.890344185 | 0.007864245 | -1.519414626 | 0.093755915 |
| Brwd3 | -1.132309037 | 0.270413373 | -0.89136931 | 0.007000428 | -1.36609594 | 0.053194932 |
| Dynlt1a | -1.336455369 | 0.121980037 | -0.891446793 | 0.042205111 | -1.698048246 | 0.105593769 |
| Snhg9 | -0.365989557 | 0.156715416 | -0.895622878 | 2.85E-09 | -1.428038414 | 0.120455841 |
| Anks6 | -0.130013473 | 0.942905259 | -0.896390821 | 0.007115237 | -1.140423739 | 0.161173688 |
| Pald1 | -0.62466845 | 0.454041884 | -0.897750481 | 0.001561377 | -0.877408321 | 0.691499826 |
| Gm27032 | -0.874373655 | 0.058905214 | -0.902810873 | 0.000179329 | -1.134465344 | 0.288202869 |
| Ccdc39 | -0.392973836 | 0.757049131 | -0.902852502 | 0.000813451 | -0.843940383 | 0.492099813 |
| Kdr | -0.747420676 | 0.280649253 | -0.905725727 | 0.013794521 | -0.799853356 | 0.906596277 |
| Dok3 | 1.071998106 | 0.226687412 | -0.909199142 | 0.003318184 | -0.650217582 | 0.90514278 |
| Igf2 | -0.798148938 | 0.315840413 | -0.917747717 | 0.003425487 | -1.068618588 | 0.426223943 |
| C1ql1 | -0.814041399 | 0.066783224 | -0.918104105 | 2.17E-06 | -1.187920612 | 0.120455841 |
| Myo19 | -0.003286309 | 0.998340434 | -0.925099304 | 0.01576935 | -1.142203471 | 0.169819532 |
| 4933439C10Rik | -0.22069747 | 0.831224605 | -0.92827523 | 0.000134277 | -1.257272904 | 0.120455841 |
| Catsperz | -0.602055213 | 0.432512862 | -0.930618502 | 4.77E-07 | -0.788298076 | 0.607196454 |
| Stard9 | -0.330214264 | 0.803347585 | -0.933759018 | 0.005478325 | -0.840949876 | 0.607196454 |
| Cntn6 | -0.656291135 | 0.410227457 | -0.93599335 | 0.000119088 | -1.032373376 | 0.810531232 |
| Ecscr | -0.527426166 | 0.470929869 | -0.94826815 | 0.0018546 | -0.610364683 | 0.993542839 |
| Zfp46 | -1.058855569 | 0.013473384 | -0.961517639 | 2.22E-06 | -1.251731522 | 0.318301009 |
| Adamts10 | -0.662226107 | 0.143641898 | -0.963571137 | 0.000384279 | -1.266379855 | 0.222875678 |
| Vcan | -0.437019442 | 0.588979009 | -0.964501143 | 0.000822554 | -1.767176081 | 0.005245142 |
| Slc40a1 | 0.067115673 | 0.963908403 | -0.968213158 | 0.008560058 | -0.243716809 | 0.993542839 |
| Traf3 | -0.785184205 | 0.050868122 | -0.972447611 | 2.75E-05 | -1.452795054 | 0.058825317 |
| Kcnk10 | -0.230708987 | 0.854226724 | -0.977587017 | 0.002426019 | -1.243919946 | 0.064866767 |
| Spef1 | -1.109098277 | 0.032741415 | -0.98266031 | 2.90E-06 | -1.190914099 | 0.143948382 |
| Pdgfra | -0.550708631 | 0.093474125 | -0.985662277 | 2.32E-08 | -1.44542608 | 0.066212538 |
| Car8 | -0.727812909 | 0.052420073 | -0.988291715 | 0.000119714 | -1.155129203 | 0.068837016 |
| Itga1 | -0.465176448 | 0.589796554 | -0.992265377 | 0.011955109 | -1.17786781 | 0.176893683 |
| Fabp7 | -0.540106867 | 0.433561182 | -0.993351532 | 0.016819914 | -1.330399295 | 0.566099353 |
| Dok4 | -0.914701657 | 0.126889335 | -1.003813356 | 2.25E-05 | -1.056722842 | 0.493900582 |
| Tet3 | -0.760619754 | 0.108712706 | -1.007733335 | 0.000235228 | -1.222624059 | 0.148749078 |
| Atp13a5 | -0.687422879 | 0.340313732 | -1.010199468 | 0.000412346 | -0.865479519 | 0.499928377 |
| Rab34 | -0.505253446 | 0.19759029 | -1.010877556 | 3.43E-07 | -1.228535088 | 0.14403429 |
| Slc22a8 | -0.14273411 | 0.860191182 | -1.018867662 | 0.001023293 | -1.365250418 | 0.041981486 |
| Car15 | -0.53019464 | 0.569628243 | -1.033076554 | 3.93E-07 | -0.871060538 | 0.468766725 |
| Dhrs13 | -2.17179472 | 0.001945581 | -1.0345415 | 0.000814149 | -1.294407856 | 0.056984159 |
| Dynlt1f | -1.180553624 | 0.115007936 | -1.034813119 | 0.006356292 | -1.285409136 | 0.217319509 |

| | | | | | | |
|---|---|---|---|---|---|---|
| Etv4 | -1.002924365 | 0.137437781 | -1.04844045 | 8.14E-05 | -1.216329535 | 0.287037664 |
| 04.Mar | -1.335121919 | 0.021327761 | -1.054200679 | 8.42E-06 | -1.230221336 | 0.121671394 |
| Pkp2 | -0.493587355 | 0.671300323 | -1.055327762 | 0.00027091 | -0.713294172 | 0.993542839 |
| Igsf3 | -0.746599256 | 0.226529853 | -1.069024018 | 0.005896434 | -1.093360564 | 0.338503594 |
| Pcdh18 | 0.201526286 | 0.905430476 | -1.073552706 | 0.00098457 | -0.905609936 | 0.416271582 |
| Ndnf | 0.106231187 | 0.876954951 | -1.08092312 | 0.001399768 | -0.876088402 | 0.549889564 |
| Notch4 | -0.91019208 | 0.286514864 | -1.08287698 | 0.000680564 | -1.053658569 | 0.244529146 |
| Gpx7 | -1.086175764 | 0.087928842 | -1.084198323 | 3.22E-06 | -0.843971158 | 0.51168305 |
| Syndig1l | -0.529168763 | 0.398506323 | -1.087198673 | 0.005742258 | -0.966914691 | 0.789365697 |
| Pip5kl1 | -0.157859659 | 0.941308731 | -1.092055371 | 1.62E-06 | -1.528237942 | 0.025071994 |
| Col26a1 | -1.856724614 | 0.004705084 | -1.096761054 | 0.001480968 | -1.877699286 | 0.004361519 |
| Ctla2a | -0.740704816 | 0.066270374 | -1.098969384 | 4.74E-05 | -1.26230065 | 0.056537244 |
| Fbxw9 | -1.238830891 | 0.002213206 | -1.107095774 | 2.53E-07 | -1.434513434 | 0.14403429 |
| Shisal1 | 0.025490678 | 0.9783404 | -1.121340873 | 2.51E-10 | -1.054735062 | 0.284181522 |
| Cnpy1 | -1.342248841 | 0.010244429 | -1.127461976 | 0.049789925 | -0.784864708 | 0.85552675 |
| Ebf4 | -0.235388486 | 0.867726781 | -1.13001807 | 1.70E-05 | -1.023525439 | 0.537400201 |
| Elovl2 | -0.972172535 | 0.078361197 | -1.132924767 | 0.001446342 | -1.858550612 | 0.001910659 |
| Col16a1 | -0.799164024 | 0.081114957 | -1.133859564 | 1.49E-05 | -1.182082118 | 0.154941158 |
| Ifitm1 | -0.233955699 | 0.873157988 | -1.134994532 | 9.04E-05 | -1.642074125 | 0.008574694 |
| Sap25 | -0.750041229 | 0.279269831 | -1.135257273 | 0.000100298 | -0.940211373 | 0.702044135 |
| Tie1 | -0.82447809 | 0.312483201 | -1.143203667 | 0.000845489 | -0.84091522 | 0.702044135 |
| Smim3 | -0.558333336 | 0.41958044 | -1.145326391 | 4.96E-07 | -0.815251604 | 0.548463319 |
| Adam19 | -1.732564141 | 0.006064926 | -1.145516264 | 9.38E-05 | -1.244704455 | 0.154941158 |
| Eva1b | -0.620323906 | 0.37175461 | -1.152372676 | 0.002920496 | -1.478737545 | 0.144616872 |
| Npnt | -0.160998984 | 0.894085043 | -1.170899987 | 5.03E-05 | -1.339519339 | 0.072051016 |
| Ccdc3 | -0.145579546 | 0.876322727 | -1.171747793 | 2.73E-06 | -0.9853772 | 0.366017902 |
| Zfp41 | -1.380852059 | 0.126897769 | -1.177813152 | 3.30E-05 | -1.32041289 | 0.162913907 |
| Vav2 | -0.619562484 | 0.431298879 | -1.187997653 | 0.000800356 | -1.595034637 | 0.008221945 |
| Insyn2b | -0.942541238 | NA | -1.193520548 | 0.008937564 | -0.577689045 | 0.993542839 |
| Gm5577 | 0.093107163 | 0.96842841 | -1.199216722 | 0.000882508 | -0.92818604 | 0.727825365 |
| Ptpro | -0.657204185 | 0.187021239 | -1.201483446 | 2.05E-09 | -1.450725619 | 0.11149893 |
| Col22a1 | -2.300048627 | 0.053560462 | -1.205555068 | 0.001210838 | -1.325035429 | 0.159216978 |
| Lama4 | -1.182723423 | 0.10076685 | -1.213701857 | 0.00069132 | -1.219589365 | 0.242348538 |
| Sla | -1.577954266 | 0.019589423 | -1.232056329 | 0.002535786 | -1.004057049 | 0.963286002 |
| Sema5a | -0.42225791 | 0.378180547 | -1.233768522 | 2.03E-05 | -1.261620771 | 0.144090638 |
| Rem2 | -0.660073172 | 0.4639562 | -1.248662124 | 0.000240736 | -1.009811202 | 0.52600127 |
| Serpinh1 | -0.527012533 | 0.312076691 | -1.260672846 | 1.67E-05 | -1.218396381 | 0.122195887 |
| Cspg4 | -0.773079637 | 0.112929364 | -1.264417838 | 1.24E-07 | -1.182532953 | 0.400222874 |
| Ccl17 | -0.644043555 | 0.52565529 | -1.276472591 | 1.93E-05 | -1.578021632 | 0.122393321 |
| Dcx | -2.114215853 | 0.001916363 | -1.282779202 | 1.87E-06 | -1.385251283 | 0.160901822 |
| Grin2d | -0.907110301 | 0.078606748 | -1.290702514 | 5.18E-09 | -1.574887475 | 0.058825317 |
| Cenpa | -2.094187067 | 0.037862512 | -1.30018278 | 0.000179745 | -1.05298369 | 0.221127986 |
| Gm27004 | -1.079977925 | 0.223314347 | -1.315987727 | 6.66E-06 | -1.108418763 | 0.57790817 |
| Clmp | -0.331836165 | 0.650064587 | -1.31792821 | 4.94E-07 | -1.511794966 | 0.089309572 |
| Ncaph | -1.802462716 | 0.192282764 | -1.32884492 | 0.001553919 | -1.220967805 | 0.452078746 |
| Gm20033 | -1.920226593 | 0.045283476 | -1.344572966 | 0.000641456 | -1.39051485 | 0.313767783 |
| Kmt5c | -0.759710471 | 0.247782387 | -1.344983564 | 2.14E-07 | -1.335199722 | 0.052362561 |
| Traf4 | -0.806727207 | 0.251843499 | -1.366311226 | 1.22E-07 | -1.374033755 | 0.064866767 |
| Kif21b | -1.053416404 | 0.091728428 | -1.376124751 | 2.51E-05 | -1.661903885 | 0.120455841 |
| Ap1g2 | -1.368485185 | 0.122653656 | -1.387147629 | 0.00056012 | -1.332412049 | 0.176893683 |
| Hrk | -0.772804409 | 0.116461671 | -1.395648103 | 3.08E-07 | -1.344049576 | 0.079417314 |
| Igsf1 | -1.103624299 | 0.195183393 | -1.412224261 | 7.79E-07 | -1.875588463 | 0.00510989 |
| Cd38 | -0.764077698 | 0.510537427 | -1.412953409 | 0.000236522 | -1.132123014 | 0.442019435 |
| Gm20501 | -0.438791052 | 0.566653226 | -1.426930879 | 9.05E-05 | -1.249228869 | 0.282442752 |
| Slc26a6 | -0.814772773 | 0.175433742 | -1.47009303 | 1.45E-10 | -1.368137929 | 0.053194932 |
| Cnbd2 | -0.806615631 | 0.415249217 | -1.477598939 | 1.21E-08 | -2.11279454 | 4.00E-05 |
| Bglap3 | -1.746036473 | 0.001984829 | -1.482112513 | 0.000121602 | -1.445387675 | 0.029495229 |
| Col1a2 | -0.882883686 | 0.135247129 | -1.508332044 | 1.61E-06 | -1.374152647 | 0.056537244 |
| AI506816 | -0.84160172 | 0.295924058 | -1.526082171 | 0.000222883 | -0.953375317 | 0.440677298 |
| Col18a1 | -0.877883581 | 0.211245314 | -1.526320969 | 2.95E-05 | -1.35381858 | 0.120455841 |
| Enpp6 | -1.983749256 | 0.013484847 | -1.548207022 | 0.000534647 | -1.751116154 | 0.1546397 |
| Muc3a | -0.974290592 | 0.42964689 | -1.565364927 | 4.78E-06 | -1.067642336 | 0.223860837 |
| Col5a1 | -0.977517572 | 0.120824424 | -1.570000125 | 1.41E-05 | -1.582616257 | 0.154941158 |
| Tmem163 | -0.920266798 | 0.025322435 | -1.578792788 | 3.73E-09 | -1.019765133 | 0.407128671 |
| Gm3435 | -0.145242313 | NA | -1.60989479 | 2.66E-05 | -1.696866379 | 0.018509708 |
| AW551984 | -1.751375691 | 0.119942339 | -1.610014005 | 3.28E-05 | -1.558091944 | 0.058231153 |
| E130006D01Rik | -1.895162044 | NA | -1.633057607 | 0.00014296 | -1.162414219 | 0.383865425 |
| Ism1 | -0.551943738 | 0.671353901 | -1.640771798 | 4.35E-05 | -0.754341177 | 0.95811614 |
| Hcrtr1 | -0.815550079 | NA | -1.646049081 | 0.000106496 | -1.668976128 | 0.081245568 |
| Mia | -1.520185174 | 0.036923628 | -1.652150632 | 2.80E-05 | -0.974015856 | 0.522741515 |
| Fam184b | -0.939554059 | 0.200761298 | -1.72689811 | 9.69E-07 | -1.130808284 | 0.354283275 |
| C030013G03Rik | -2.523633649 | NA | -1.747638902 | 0.000149138 | -1.036725313 | 0.708926839 |
| Tet1 | -1.582515725 | 0.019414816 | -1.777290728 | 2.24E-06 | -1.930515952 | 0.067728655 |

| | | | | | | |
|---|---|---|---|---|---|---|
| Col1a1 | -0.602410237 | 0.522245112 | -1.788163241 | 6.82E-07 | -1.676147481 | 0.033636292 |
| Nts | -1.672346546 | 0.016428564 | -1.912016046 | 4.53E-08 | -1.554861523 | 0.780092679 |
| 1500035N22Rik | -2.080763679 | NA | -1.952171754 | 4.65E-05 | -1.404650803 | 0.13599948 |
| Robo3 | -1.675440443 | 0.105026208 | -2.209219041 | 1.61E-07 | -2.198590031 | 0.005538947 |
| Met | -1.133841053 | 0.369114931 | -2.250705682 | 3.21E-08 | -1.280014002 | 0.414591828 |
| Emid1 | -2.084061597 | 0.010105346 | -2.38962339 | 1.76E-09 | -2.782981955 | 6.23E-06 |
| C1qtnf6 | -2.948467469 | 0.000590276 | -2.446439644 | 1.50E-07 | -2.397295186 | 0.011560784 |
| Mkrn3 | -3.326821467 | 0.005533311 | -3.674791183 | 4.01E-05 | -3.788330371 | 6.23E-06 |
| Chrna3 | -2.811948834 | NA | -5.139840315 | 3.89E-06 | -1.524206573 | 0.207054318 |
| n = 37 | MEC_s | | MEC_im | | MEC_d | |
| gene | log2FoldChange | padj | log2FoldChange | padj | log2FoldChange | padj |
| Xist | 5.963581281 | 0.001630677 | 7.56039505 | 1.91E-05 | 4.073556502 | 2.08E-05 |
| Defb1 | 3.131561727 | 0.012424732 | 4.149242614 | 6.43E-11 | 3.453914238 | 0.000519773 |
| Fosb | 1.912389493 | 0.014798119 | 2.635892323 | 1.63E-19 | 1.981761456 | 0.01937533 |
| Tcap | -0.364810949 | 0.76269534 | -1.17635288 | 6.32E-05 | -1.212075141 | 0.047607156 |
| Cacna1h | -0.156580393 | 0.89641451 | -0.972780087 | 2.68E-09 | -1.289846661 | 0.038734655 |
| Igfbp2 | -0.076877199 | 0.952757129 | -0.566296298 | 0.04590162 | -1.297264591 | 0.039844811 |
| Nanos3 | -1.501554158 | 0.019454516 | -0.892966987 | 4.89E-05 | -1.321177794 | 0.020799188 |
| Mmp15 | -0.9389478 | 0.053143452 | -1.012291542 | 6.51E-06 | -1.335683291 | 0.048630507 |
| Csrp2 | -0.670910339 | 0.25923341 | -0.874125284 | 0.007093719 | -1.349882789 | 0.034271287 |
| Unc13b | -0.66343114 | 0.081461262 | -0.533307078 | 0.000259077 | -1.398997814 | 0.034062608 |
| Zkscan2 | -0.333290798 | 0.754820056 | -0.564628072 | 0.0150851 | -1.409291413 | 0.041479197 |
| Spink10 | -0.798599704 | 0.107755698 | -0.709194962 | 0.000191976 | -1.42286916 | 0.024358057 |
| Afap1 | -0.607201393 | 0.158496999 | -1.012663344 | 6.63E-05 | -1.428093827 | 0.034432384 |
| Slc44a5 | -0.569344756 | 0.437518054 | -0.707500256 | 4.53E-05 | -1.440957801 | 0.030487199 |
| Iglon5 | -1.074038421 | 0.041899927 | -0.878460852 | 8.82E-06 | -1.487571445 | 0.019098929 |
| Id3 | -0.156009141 | 0.622823314 | -0.795989138 | 1.81E-06 | -1.504699839 | 0.03066901 |
| A230057D06Rik | -0.517270033 | 0.619218724 | -0.769811284 | 0.007814922 | -1.507800932 | 0.013875166 |
| Calb2 | -0.474763428 | 0.247224763 | -1.086427701 | 0.00074251 | -1.508541505 | 0.048544447 |
| Sox4 | -0.516176962 | 0.390806961 | -0.704208714 | 6.20E-06 | -1.52218216 | 0.021427857 |
| Zfp282 | -1.43351523 | 0.002383805 | -0.931596769 | 2.65E-05 | -1.53512496 | 0.015191189 |
| Tmem169 | -0.74774915 | 0.267438387 | -0.702779546 | 0.000735124 | -1.595289865 | 0.015191189 |
| Prom1 | -0.802995454 | 0.084009904 | -1.771376244 | 6.46E-19 | -1.605760049 | 0.003123286 |
| Srgap1 | -0.386905321 | 0.659324344 | -0.557203967 | 0.012729154 | -1.632435569 | 0.007008625 |
| Fn1 | -0.545162462 | 0.084152104 | -0.842739429 | 0.009645973 | -1.675130104 | 0.006086541 |
| Crybb1 | -0.729894468 | 0.294500427 | -1.229156311 | 4.53E-05 | -1.709078173 | 0.029771345 |
| Gp1bb | -1.220366966 | 0.031415755 | -1.716592153 | 4.22E-07 | -1.715870676 | 0.021427857 |
| Parvb | -0.691080081 | 0.204712901 | -0.773113184 | 0.005908508 | -1.73565521 | 0.026797842 |
| Rgs11 | -0.918705278 | 0.159065979 | -1.108946816 | 0.003946937 | -1.738698978 | 0.041539125 |
| Faap100 | -0.428799174 | 0.511089229 | -0.935697496 | 4.74E-05 | -1.751348453 | 0.006697647 |
| Nptx2 | -1.040459002 | 0.034490199 | -1.050083522 | 0.000576861 | -1.787622005 | 0.024716696 |
| Rras2 | -1.009476205 | 0.030881651 | -1.306247704 | 1.62E-06 | -1.794407972 | 0.013875166 |
| Fam71e1 | -0.632407534 | 0.397317453 | -0.919449182 | 5.64E-05 | -1.852290416 | 0.00729645 |
| Slc29a4 | -1.399675941 | 0.043492662 | -2.24087514 | 1.24E-10 | -1.949628242 | 0.018356888 |
| Lpl | -1.117983784 | 0.127361654 | -1.081939913 | 0.013513038 | -2.032681301 | 0.036044927 |
| Gm43305 | -1.936190919 | 0.000764241 | -2.245940522 | 4.87E-11 | -2.316589263 | 3.08E-07 |
| Lsm11 | -1.48723731 | 0.002310315 | -1.491078641 | 3.88E-06 | -2.399651333 | 0.008574694 |
| Cd24a | -1.996826938 | 0.004281791 | -2.151018799 | 2.44E-08 | -3.906120328 | 1.33E-07 |
| n = 5 | MEC_s | | MEC_im | | MEC_d | |
| gene | log2FoldChange | padj | log2FoldChange | padj | log2FoldChange | padj |
| P3h3 | 0.148750542 | 0.843343574 | -0.481785796 | 0.05304548 | -1.407383331 | 0.041981486 |
| Dchs1 | -0.553820905 | 0.496849836 | -0.390786644 | 0.202364856 | -1.491093142 | 0.018498267 |
| Lypd1 | -0.308972593 | 0.410227457 | -0.307687715 | 0.011619207 | -1.509091685 | 0.040467838 |
| Cdh8 | -0.419905723 | 0.496575986 | -0.38931038 | 0.081272148 | -1.531472997 | 0.015718225 |
| Ctsk | -0.666246414 | 0.504209111 | -0.843918109 | 0.145519028 | -2.045105716 | 0.008221945 |
